# A prostaglandin alpha F2 analog protects from statin-induced myopathic changes in primary human muscle cells

**DOI:** 10.1101/271932

**Authors:** Stefanie Anke Grunwald, Oliver Popp, Stefanie Haafke, Nicole Jedraszczak, Ulrike Grieben, Kathrin Saar, Giannino Patone, Wolfram Kress, Elisabeth Steinhagen-Thiessen, Gunnar Dittmar, Simone Spuler

**Affiliations:** Muscle Research Unit, Experimental and Clinical Research Center, a joint cooperation between the Charité Medical Faculty and the Max Delbrück Center for Molecular Medicine, Berlin 13125, Germany; Charité Universitätsmedizin Berlin, Berlin 13125, Germany; Mass Spectrometry Core Facility, Max Delbrück Center for Molecular Medicine in the Helmholtz Society, Berlin 13125, Germany; Mass Spectrometry Facility, Berlin Institute of Health, Berlin 13125, Germany; Interdisciplinary Lipid Metabolic Center, Charité Universitätsmedizin Berlin, Berlin 13353, Germany; Genetics and Genomics of Cardiovascular Diseases, Max Delbrück Center for Molecular Medicine in the Helmholtz Society, Berlin 13125, Germany; Institute for Human Genetics, Julius-Maximilians-University of Würzburg, Würzburg 97074, Germany

**Author notes:** **Corresponding author**: Stefanie Grunwald and Simone Spuler Stefanie Grunwald, Simone Spuler, Lindenberger Weg 80, 13125 Berlin, Germany.

**Keywords:** statin, cholesterol, signal transduction, eicosanoids, primary human muscle cell, therapy

## Abstract

Statin-related muscle side effects are a constant healthcare problem since patient compliance is dependent on side effects. Statins reduce plasma cholesterol levels and can prevent secondary cardiovascular disease. Although statin-induced muscle damage has been studied, preventive or curative therapies are yet to be reported.

We exposed primary human muscle cell populations (n=25) to a lipophilic (simvastatin) and a hydrophilic (rosuvastatin) statin and analyzed their expressome. Data and pathway analyses included GOrilla, Reactome and DAVID. We measured mevalonate intracellularly and analyzed eicosanoid profiles secreted by human muscle cells. Functional assays included proliferation and differentiation quantification.

More than 1800 transcripts and 900 proteins were differentially expressed after exposure to statins. Simvastatin had a stronger effect on the expressome than rosuvastatin, but both statins influenced cholesterol biosynthesis, fatty acid metabolism, eicosanoid synthesis, proliferation, and differentiation of human muscle cells. Cultured human muscle cells secreted ω-3 and ω-6 derived eicosanoids and prostaglandins. The ω-6 derived metabolites were found at higher levels secreted from simvastatin-treated primary human muscle cells. Eicosanoids rescued muscle cell differentiation.

Our data suggest a new aspect on the role of skeletal muscle in cholesterol metabolism. For clinical practice, the addition of omega-n fatty acids could be suitable to prevent or treat statin-myopathy.

## 1. Introduction

Patient non-compliance to medication due to side effects is a major healthcare concern. Statin side effects on skeletal muscle causes about 70% of patients to discontinue treatment^1-7^. Millions of patients ingest statins to reduce blood cholesterol levels thereby preventing cardiovascular diseases. Most statin-intolerant patients experience muscle cramps and myalgia, with occasional asymptomatic elevation of creatine kinase and weakness. Although skeletal muscle damage has been studied in mice^8^, it is unknown whether the same or different mechanisms underlie the various myopathic phenotypes and how we can circumvent or treat statin-induced changes in skeletal muscle. Also, the effect of lipophilic and/or hydrophilic statins on muscle is unclear. Removal of cholesterol might affect membrane structure and lipid rafts that are essential for signalling and cellular compartmentalization. Given that, cholesterol-lowering agents including proprotein convertase subtilisin / kexin type 9 – inhibitors (PCSK9-inhibitors), would lead to the same spectrum of muscle complaints. Alternatively, statins might influence pathways downstream of the hydroxymethylglutaryl-coenzyme A (HMG-CoA) reduction into mevalonate by the reductase HMGCR that is inhibited by statins and causing a distinct myopathy^9,10^.

Mevalonate can be metabolized into squalene for cholesterol biosynthesis^11^ or into metabolites, i.e. farnesyl-pyrophosphate and geranylgeranyl-pyrophosphate, involved in isoprenoid synthesis, protein prenylation, and ubiquinone synthesis. Consequently, hypotheses for statin-related side effects include a decrease in ubiquinone synthesis, cholesterol, and vitamin D levels, and a loss of isoprenoid metabolites. Altering vitamin D levels and protein ubiquination remain an intriguing yet controversial hypothesis for causing statin-induced side effects^10,12-15^. Interventions into isoprenoid synthesis have been associated with ATP depletion and DNA damage and therefore suggested for cancer therapy^16,17^. Since ubiquinone is critical for the mitochondrial respiratory chain, mitochondrial impairment has been reported in vitro as well as in statin-treated patients^18^. Ubiquinone supplementation is argumentatively beneficial to patients^19^. However, most studies have been performed in non-human muscle or human non-muscle cell lines or animals and only focus on mevalonate depletion.

In this study, we asked whether there are key signalling pathways in muscle that are significantly interfered by statins and may have a statin myopathy management potential. We chose an integrated approach to study the effects of a lipophilic and a hydrophilic statin in many individual primary human myoblast cells at molecular, cellular, and functional levels^20^. We developed a new assay for mevalonate detection and showed that statins have a significant effect on gene and protein expression profiles. Our model for signal transduction pathway suggests statin-induced metabolic changes upstream of mevalonate, particularly in mitochondrial ß-oxidation and eicosanoid synthesis. Eicosanoids have profound effects on inflammation, cell proliferation and differentiation, and acts as pain mediators. Primary human muscle cells secrete eicosanoids and prostaglandins in vitro. Arachidonic acid derived eicosanoid levels changed time dependently. Furthermore, statins directly influence the proliferative and regenerative features of skeletal muscle cells. Eicosanoid-species substitution to statin-treated primary human muscle cells rescue these effects. Statin myopathy, therefore, appears as a distinct and potentially treatable myopathy driven by HMG-CoA accumulation rather than mevalonate depletion.

## 2. Results

In primary human muscle cell populations obtained from 25 patients we investigated key signalling pathways that are significantly interfered by statins in muscle cells and their impact on cellular and molecular levels.

### Simvastatin more than rosuvastatin extensively alters myotube expressome

Transcriptome data obtained from statin-treated primary human myotubes group clearly separate from untreated and DMSO-treated samples on a PCA analysis plot (Figure 1A). A concern about studying primary human samples derived from individuals of different age and gender usually has been the potentially high variation between samples. Clinical patient information is provided in Supplementary Table S1. Here, however, we can show that primary human samples behave very similar among various conditions but different for both statins (Figure 1A). We found that simvastatin had a dramatic effect on primary human muscle cells, much more than rosuvastatin. 1807 genes (FDR<0.05) were differentially expressed after exposure to simvastatin (Figure 1B, Supplementary Table S2, Supplementary Data File 1). Simvastatin had a particularly strong impact on RNA metabolism (Table 1). The number of genes affected by rosuvastatin was only 68 (FDR<0.05) (Supplementary Table S3, Supplementary Data File 2) with a main impact on lipid and cholesterol metabolism.

**Table 1.** Biological process clustering. Genes and proteins differentially expressed in statin-treated primary human myotubes at mRNA and protein level were clustered into biological processes using GOrilla. The GO Term Top10 are different for each statin but similar at both expression levels. Lipid and cellular metabolic processes are most commonly modified by both statins at both expression levels. The p-values correspond to number of genes / proteins clustered into a GO node. (Supplementary Tables S2-S6)

| Expression level | GO Term – Top 10 | P-value |
| --- | --- | --- |
| **Simvastatin**<br>**Transcriptome** | • SRP-dependent cotranslational protein targeting to membrane | 5.65 E-29 |
|  | • Cotranslational protein targeting to membrane | 1.60 E-28 |
|  | • Protein targeting to ER | 1.27 E-26 |
|  | • rRNA processing | 5.10 E-26 |
|  | • Establishment of protein localization to endoplasmic reticulum | 7.75 E-26 |
|  | • rRNA metabolic process | 3.00 E-24 |
|  | • Protein localization to endoplasmic reticulum | 4.42 E-24 |
|  | • Translational initiation | 1.88 E-23 |
|  | • Nuclear-transcribed mRNA catabolic process, nonsense-mediated decay | 8.48 E-23 |
|  | • Viral transcription | 1.03 E-21 |
| **Simvastatin**<br>**Proteome** | • Negative regulation of cellular process | 2.30E-04 |
|  | • Regulation of biological quality | 3.91E-04 |
|  | • Signal transduction | 2.69E-05 |
|  | • RNA metabolic process | 7.27E-04 |
|  | • Small molecule metabolic process | 5.35E-04 |
|  | • Regulation of localization | 2.37E-04 |
|  | • Immune system process | 4.85E-04 |
|  | • Macromolecule biosynthetic process | 8.40E-04 |
|  | • Multicellular organismal process | 2.72E-04 |
|  | • Positive regulation of response to stimulus | 8.05E-04 |
| **Rosuvastatin**<br>**Transcriptome** | • Secondary alcohol biosynthetic process | 3.29 E-31 |
|  | • Sterol biosynthetic process | 5.28 E-30 |
|  | • Cholesterol biosynthetic process | 1.47 E-29 |
|  | • Secondary alcohol metabolic process | 9.54 E-27 |
|  | • Cholesterol metabolic process | 6.92 E-26 |
|  | • Sterol metabolic process | 1.47 E-25 |
|  | • Alcohol biosynthetic process | 9.80 E-25 |
|  | • Steroid biosynthetic process | 4.95 E-23 |
|  | • Organic hydroxy compound biosynthetic process | 7.60 E-22 |
|  | Regulation of cholesterol biosynthetic process | 1.21 E-21 |
| **Rosuvastatin**<br>**Proteome** | Small molecule metabolic process<br>Oxidation-reduction process<br>Small molecule catabolic process<br>Cellular lipid metabolic process<br>Monocarboxylic acid metabolic process<br>Small molecule biosynthetic process<br>Lipid metabolic process<br>Regulation of lipid metabolic process<br>Lipid biosynthetic process<br>Organic hydroxy compound metabolic process | 1.73E-04<br>7.32E-05<br>6.13E-04<br>9.49E-07<br>9.03E-06<br>1.26E-05<br>3.11E-09<br>7.54E-05<br>2.90E-08<br>6.49E-08 |
| **Common GO-**<br>**terms at**<br>**transcriptome**<br>**and proteome**<br>**level influenced**<br>**by both statins** | **Lipid metabolic process**<br>Cholesterol metabolic process<br>Sterol metabolic process<br>Cholesterol biosynthetic process<br>Isoprenoid biosynthetic process<br>Triglyceride metabolic process<br>Fatty acid metabolic process<br>Arachidonic metabolism | 8.07 E-11<br>5.65 E-10<br>1.65 E-09<br>1.37 E-08<br>1.15 E-06<br>1.67 E-04<br>8.52 E-05<br>3.33 E-04 |

**Figure 1.**
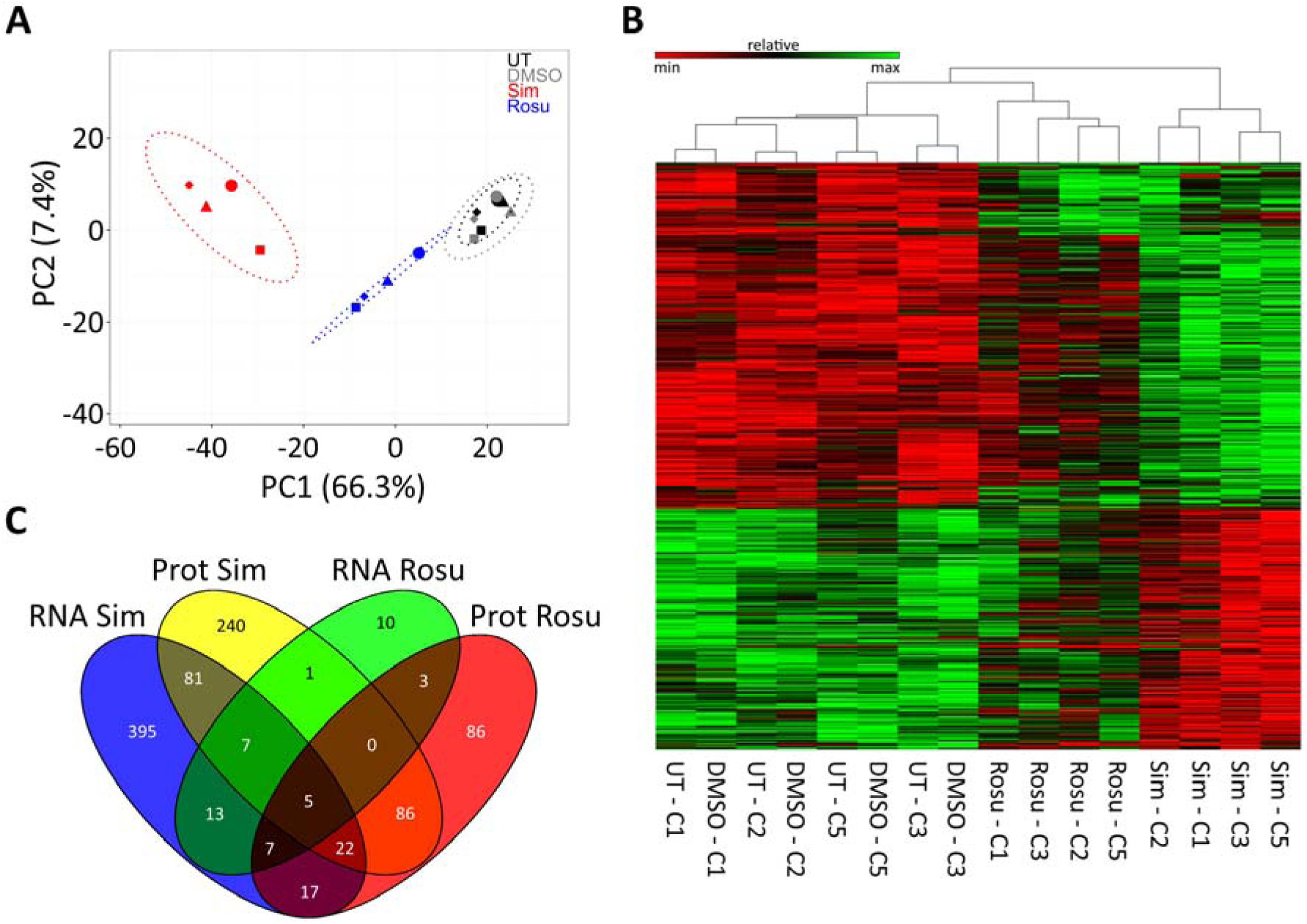
Gene expression profiling of primary human myotubes exposed to statins. Four primary human myotube cell populations from four different statin naïve donors were left untreated (UT), treated with DMSO, rosuvastatin (Rosu) or simvastatin (Sim). (A) Principal components analysis (PCA) of RNA-Seq samples. The PCA plot shows a clear grouping. (B) Heatmap of RNA-Seq samples represents log2 transformed normalized count data of differently expressed genes. Samples of each statin group cluster hierarchically together. Untreated and DMSO treated samples are equally clustered confirming that DMSO has no significant effect on RNA expression data. (C) Venn diagram displaying overlaps between proteome and transcriptome expression data from simvastatin and rosuvastatin groups. For cell populations used see Supplementary Table S1. See also Supplementary Figure S1 and Supplementary Data File 1 and 2.

When transcriptome and proteome of identical samples are compared, there may or may not be a good overlap. We found that on proteome level 3255 skeletal muscle proteins were significantly identified after exposure to statins (Supplementary Figure S1, Supplementary Data File 3; Supplementary Tables S5, S6). Proteins belonging to lipid metabolic, small molecule metabolic, and cellular, developmental as well as RNA metabolic processes were particularly affected (Supplementary Data File 4). A Venn diagram (Figure 1C) demonstrates that 19% of genes and proteins in simvastatin and 6% in rosuvastatin treated myotubes were significantly differentially expressed at proteome <u>and</u> transcriptome levels when compared to controls. Five gene products are differently regulated by both statins at mRNA as well as protein levels: Calsyntenin-2 (CLSTN2), lanosterol 14-alpha demethylase (CYP51A1), squalene synthase (FDFT1), isopentenyl-diphosphate delta-isomerase 1 (IDI1), and tubulin beta-2B chain (TUBB2B) (Supplementary Table S7). All five are involved in cell adhesion and cholesterol biosynthesis. The degree of similarity between RNA-Seq and proteome-data is visualized in a circus plot (Supplementary Video)^21^.

The significant effect of statins on the transcriptome and proteome of primary human muscle cells was unexpected and widespread. Because both, simvastatin and rosuvastatin, induce “statin-myopathy” we focussed our further analysis on pathways in muscle cells that were influenced by both statins. These were fatty acid metabolism, fatty acid import, and lipid metabolic processes (Table 1, Supplementary Data File 4).

### Muscle cells possess a similar repertoire for cholesterol biosynthesis as liver

We next asked how the muscle was equipped to synthesize and metabolize cholesterol. We found key enzymes for cholesterol biosynthesis (HMGCS1, HMGCR, CYP51A1, DHCR7) as well as cholesterol transport (LDLR, ABCA1, ABCG1) expressed in human muscle cells and significantly altered by different exposure to statins. We further analysed the transcript variants of 3-hydroxy-3-methylglutaryl-CoA reductase (HMGCR) of human muscle cells and compared them to human liver (HepG2 cells) (Figure 2A). HMGCR transcript variants (ENST00000287936 and ENST00000343975 that excludes exon 13) have an impact on cholesterol metabolism regulation and statin efficacy^22^. The transcripts in primary human myotubes and HepG2 cells were identical (Figure 2A). In our OMICs data set (Supplementary Table S4) the ENST00000343975 was also detectable at low levels. We then tested to what extent statins added to primary human muscle cells would influence cholesterol and mevalonate levels, or expression of cholesterol dependent gene expression (Figure 2B-D). To accomplish this, we redeveloped a simple assay for mevalonate determination in human muscle cells. Cholesterol and mevalonate levels in primary human myotubes were significantly lowered by statins in a concentration dependent manner. (Figure 2B and C). On mRNA level as assessed by qPCR, we found that statins increased HMGCS and HMGCR (Figure 2D), the same effects that have been described for liver including LDL receptor^23^. Eight protein coding transcript variants are known for LDLR. In our RNASeq data set, we found six of them differentially expressed (between statin and controls) with similar fold changes (see Supplementary Table S4). ENST00000558518, ENST00000558013, ENST00000535915, and ENST00000545707 are the most supported transcript variants throughout databases. In our data set, ENST00000545707 (682 amino acids) had the highest number of uniquely mapped and total transcript reads (FPKM 7.8-19.4). This variant misses LDL receptor class A3-5 and the EGF-like 3 domain that are linked to familiar hypercholesterolemia, i.e. rs121908028, rs121908029, rs28942083, rs748944640 (UniProt identifier P01130-2). LDLR binding partner PCSK9 has only one protein coding transcript variant which is 2fold upregulated in rosuvastatin-treated primary human myotubes. Our data show, that primary human myotubes possess the same repertoire for cholesterol biosynthesis and expression regulation feedback mechanisms as liver cells, an important condition for our further analyses.

**Figure 2.**
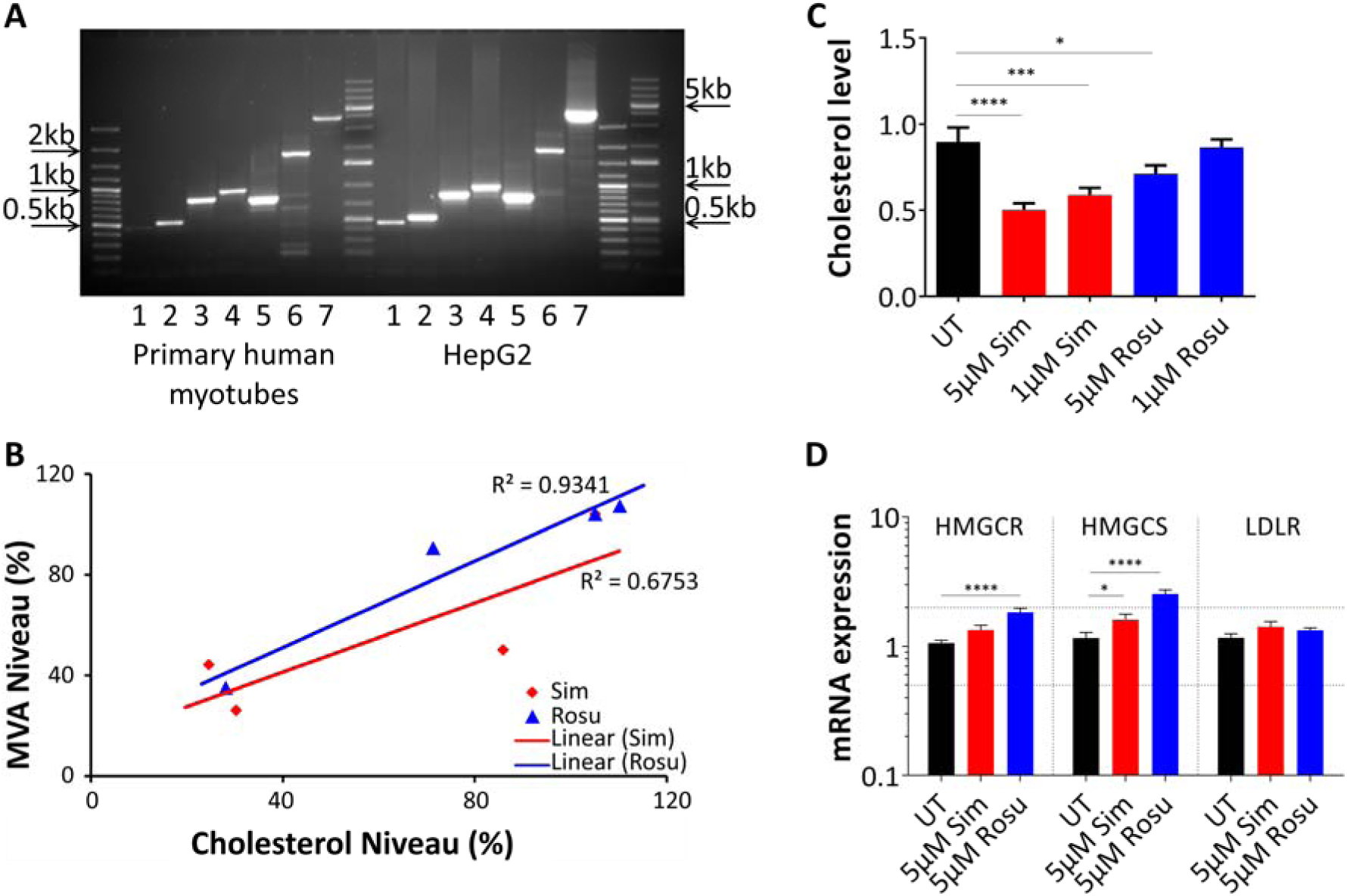
Statin effect in primary human myotubes on mevalonate and cholesterol dependent gene expression level. (A) HMGCR transcript variants amplified for exon 1-20 in primary human myotubes and liver HepG2 cells. Primary human muscle cells and HepG2 cells share the identical HMGCR transcripts. Unspecific bands in lane 6 were sequenced. [1, Exon1-5; 2, Exon 3-8; 3, Exon 6-11; 4, Exon 9-14; 5, Exon 12-17; 6, Exon 15-20; 7, Exon 1-20] (B) Mevalonate levels in primary human myoblasts after 72h statin treatment at different concentrations in delipidated medium. Mevalonate levels correlate to cholesterol levels. Data were normalized to total protein level and relativized to DMSO treated samples. (error bars = mean with SD; n=3) (C) Relative cholesterol levels were determined under delipidated medium conditions in primary human myoblasts after 72 h statin treatment at different concentrations. (D) Relative mRNA expression of HMGCR, HMGCS, and LDLR were examined in primary human myoblasts undergoing differentiation under statin treatment (5 µM) for 72 h-96 h. The results were normalized to two reference genes and calculated relatively to DMSO-treated samples. (error bars = mean with SEM; n=6). For cell populations used see Supplementary Table S1. [UT, untreated (black); Sim, simvastatin (red); Rosu, rosuvastatin (blue); HMGCR, 3-hydroxy-3-methylglutaryl-CoA reductase; HMGCS, 3-hydroxy-3-methylglutaryl-CoA synthase 1; LDLR, low density lipoprotein receptor]

### Statins interfere with fatty acid metabolism

To further analyze pathways significantly affected by both statins, we used Gene Ontology enRIchment anaLysis (GOrilla) and the Database for Annotation, Visualization and Integrated Discovery (DAVID) analysis (Supplementary Data File 4). Within the gene ontology terms *fatty acid metabolism, fatty acid import, and lipid metabolic processes*, we identified significant effects on genes involved in lipid, fatty acid, and arachidonic/ eicosanoid metabolism as well as on retinoic acid response of the retinoid X receptor’s (RXR) pathway and carnitine shuttling (Figure 3A). Carnitine palmitoyltransferase I (CPT1) was up to 12-fold upregulated at protein level. Based on these findings, we qualitatively analysed gene and protein interactions known or predicted to HMGCR and included the quantitative abstraction level for pathway modelling. We modelled a regulatory network showing the statin-induced alterations on HMGCR-associated pathways in human muscle. (Figure 3B, Supplementary Table S8). Among the general impact of statins on acetyl-CoA dependent metabolic processes we found particularly interesting upregulation of fatty acid desaturases and elongases. Both, fatty acid desaturases and elongases catalyse precursor molecules for prostaglandin and eicosanoid synthesis, molecules involved in inflammation, mitochondrial function, and mediating pain; a common finding in statin-induced myopathy. We propose that the statin effect in muscle may be driven by HMG-CoA and acetyl-CoA accumulation rather than a loss of mevalonate downstream metabolites.

**Figure 3.**
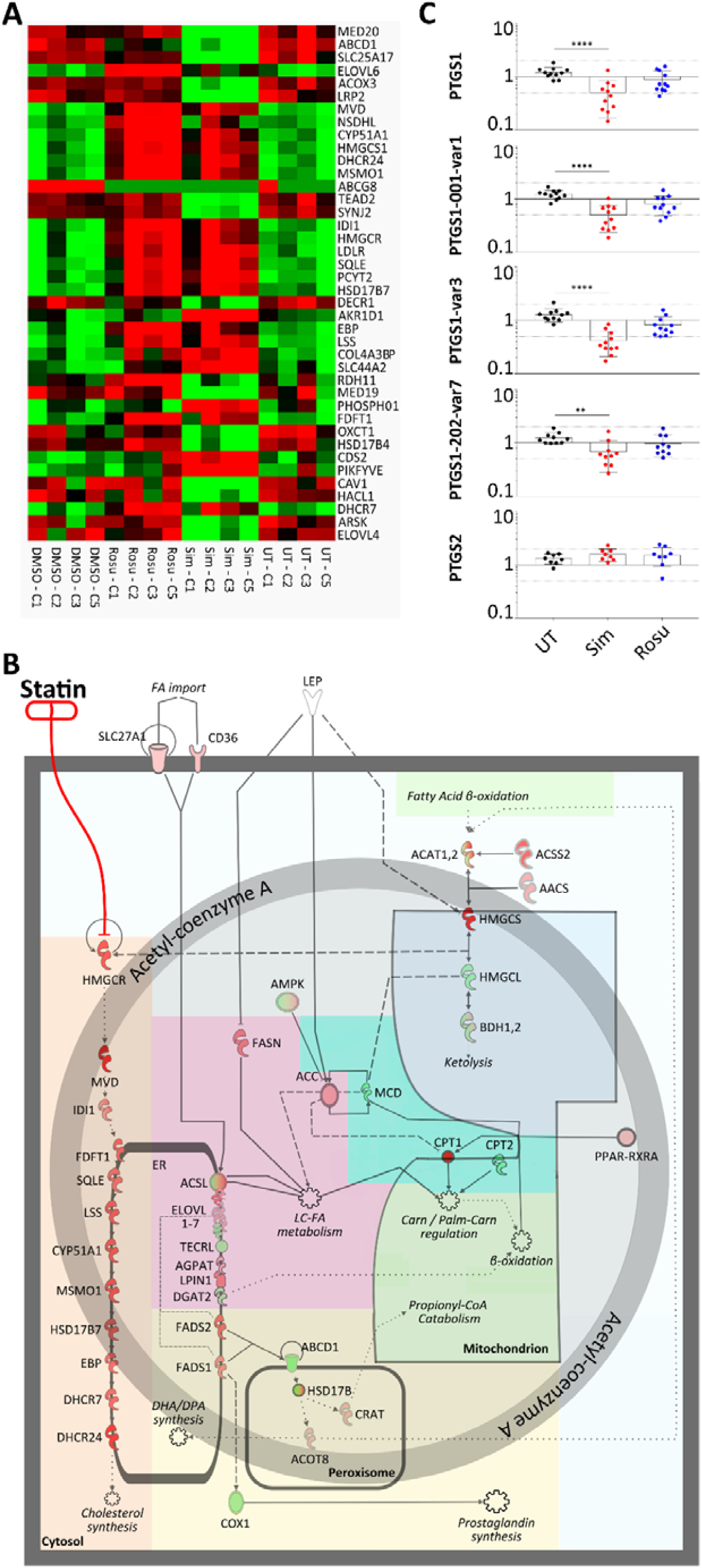
Statin influence on lipid metabolic processes and on PTGS1 mRNA expression in primary human myotubes. (A) RNA expression of primary human myotubes treated with statins compared to controls. Heatmap displays gene enrichment in lipid metabolic processes at RNA level. (Green – downregulated; Red – upregulated; n=4) (B) Signalling pathway model of statin-induced alterations in muscle lipid metabolism. Statin-induced changes in cholesterol biosynthesis (orange) are a direct effect of HMGCR inhibition. This also affects HMG-CoA synthesis and ketone body metabolism (blue) and especially Ac-CoA. Ac-CoA directly affects fatty acyl and triglyceride biosynthesis (magenta) and impairs eicosanoid synthesis (yellow). Also, Ac-CoA influences carnitine-palmitoyl transferases (turquois) and mitochondrial beta oxidation (green). See also Supplementary Table S8. (C) Transcript variant expression study of PTGS1 and PTGS2 in primary human myotubes statin-treated for 96h. For PTGS1, we analysed three different variants. PTGS1 variant 3 shows the lowest expression values (mean fold change = 0.41). Thus, it mainly reflects the low expression level detected for all variants. (error bar = mean with SD; n=11) See Supplementary Table S9 for PCR primer details. For cell populations used see Supplementary Table S1. [UT, untreated (black); Sim, 5 µM simvastatin (red); Rosu, 5 µM rosuvastatin (blue); Ac-CoA, acetyl coenzyme A; HMG-CoA, 3-hydroxy-3-methylglutaryl coenzyme A; HMGCR, 3-hydroxy-3-methylglutaryl-CoA reductase; PTGS, prostaglandin-endoperoxide synthase (**** p<0.0001); Error bar = mean with SD]

Indeed, we found that a number of genes involved in eicosanoid synthesis, such as the ATP-binding cassette sub-family D member 1, the elongation of very long chain fatty acids 4 and 6 as well as the aldo-keto reductase family 1 member D1, fatty acid desaturase, phospholipase A2 group III, and prostaglandin-endoperoxide synthase 1 (PTGS1) were significantly different expressed (Figure 3A,B). PTGS1 is constitutively expressed in a wide range of tissues and PTGS2 is the inducible isoform. Significant reduction of PTGS1 (Figure 3C), but not PTGS2, the key enzyme in ω-3 and ω-6 fatty acid metabolism converting eicosapentaenoic acid and arachidonic acid to eicosanoids including prostaglandins, points a disturbance in eicosanoid biosynthesis.

### Eicosanoid AL-8810 reverses the effect off simvastatin on primary human muscle cells

If an impairment in eicosanoid biosynthesis would be responsible for statin related side effects on muscle, reconstitution with eicosanoids might be beneficial. We first identified parameters on cellular level that would be possible to quantify and measured the effect of statins on proliferation and differentiation in primary human muscle cells. Myoblast proliferation decreased in response to both, simvastatin and rosuvastatin, in a concentration-dependent manner (Figure 4A, Supplementary Figure S2B). We found no evidence for significant statin toxicity (Supplementary Figure S2A) or oxidative stress as measured by nitric oxide concentration (Supplementary Figure S3A,B). Rescue experiments with mevalonate and geranylgeranyl pyrophosphate demonstrate that the effect on proliferation was indeed statin specific (Supplementary Figure S2B). In addition, both statins markedly delayed primary human myoblast differentiation as quantified by fusion index, and by myosin heavy chain expression on mRNA and protein level (Figure 4B and C).

**Figure 4.**
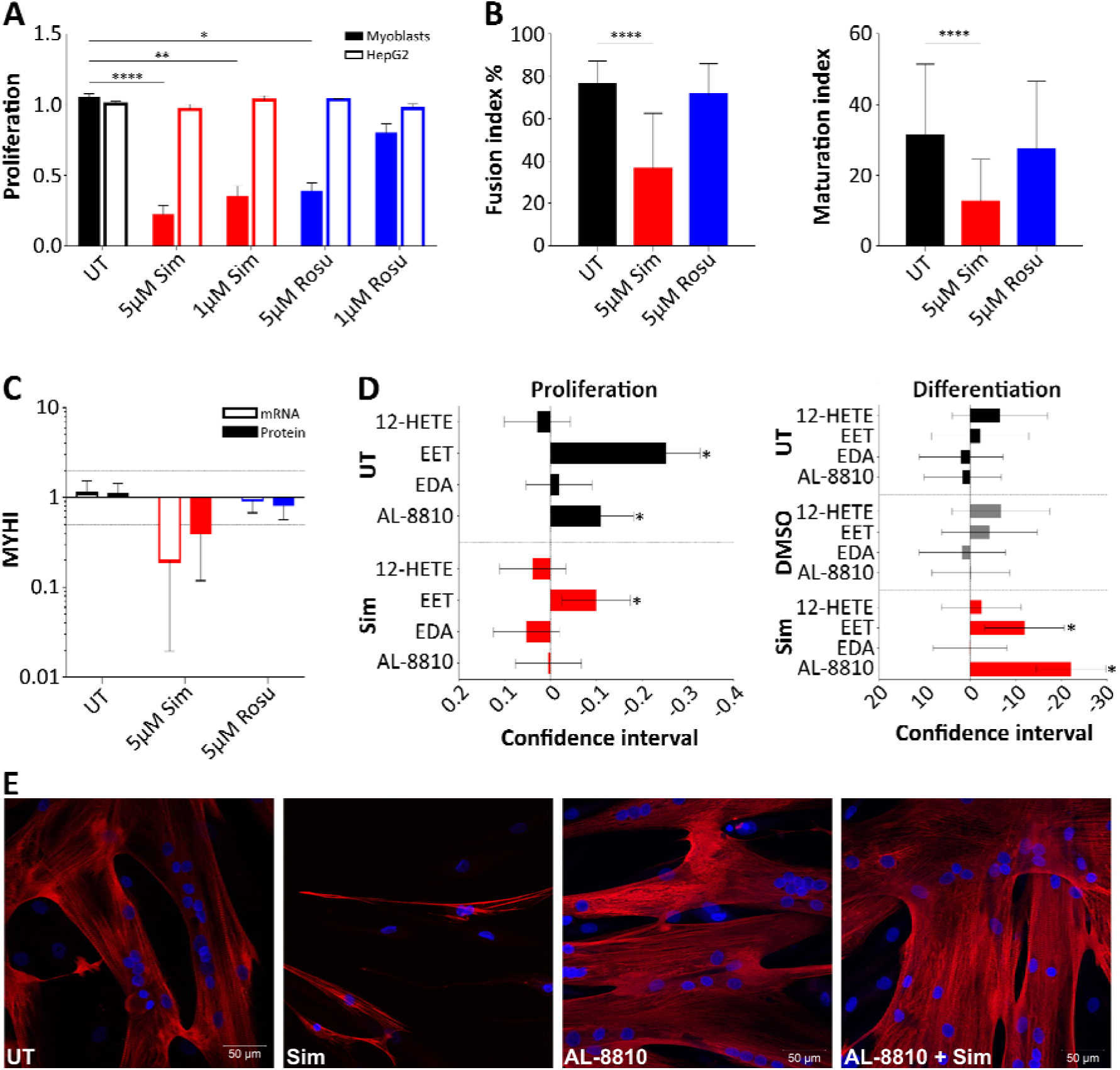
Statin influence on fatty acid metabolism of primary human myotubes. (A) Cell proliferation (BrdU) test: Primary human myoblasts and liver HepG2 cells were treated with statins for 72 h. At 5 µM concentration, statins negatively influence primary human myoblast but not HepG2 proliferation. The proliferation data were relatively calculated to DMSO-treated samples to quench the effect of DMSO. (error bars = mean with SEM; n=8 for myoblasts; n=4 for HepG2) (B) Differentiation of primary human myoblasts under statin treatment for 96 h. Primary human myoblast differentiation into myotubes is delayed under treatment with 5 µM simvastatin or 5 µM rosuvastatin. Myotube fusion and maturation index (number of nuclei per myotube) were determined by counting MYHI (red) positively immunolabelled myotubes and HOECHST (blue) stained nuclei using the cell image analysis software CellProfiler v.2.1.1. (error bars = mean with SD; n=8, 2-4 immunofluorescent pictures counted each) (C) MYHI mRNA and protein expression were examined in primary human myotubes treated with 5 µM statin for 84 h to 96 h under differentiation condition relative to DMSO-treated samples. Dashed lines represent 2 fold and 0.5 fold expression changes. (error bars = mean with SD; n=4) (D) Proliferation of primary human myoblasts under simvastatin and/or eicosanoid species treatment relative to DMSO. (5)6-EET ameliorates proliferation for simvastatin-treated samples. (95% confidence interval; n=8). (D,E) Primary human myoblast differentiation into myotubes under treatment with simvastatin and/or fatty acids. Myotube fusion index was determined by counting MYHI (red) positively immunolabelled myotubes and HOECHST (blue) stained nuclei using the cell image analysis software CellProfiler v.2.1.1. Error bars = mean with SD. (95% confidence interval; n=8, 2-4 immunofluorescent pictures counted each) (E) Primary human statin-treated myotubes regain normal morphology when AL-8810 is added to the medium. See also Supplementary Figure S5. For cell populations used see Supplementary Table S1. [UT, untreated (black); Sim, simvastatin (red); Rosu, rosuvastatin (blue); MYHI, myosin heavy chain 1]

In a time-series experiment, we analysed eicosanoid profile changes in the supernatant from statin-treated primary human myoblasts under fusion conditions compared to DMSO control. Because we found primary human myoblast fusion under statin treatment still occurring but delayed, we analysed the first 6 h to 36 h of myoblast fusion onset as well as 96 h matching the expressome analyse time point. (Supplementary Figure S4, Supplementary Table S10) Long-chain ω-6 polyunsaturated fatty acids (PUFA) arachidonic acid derived eicosanoids (PGF2a, 8-iso-PGF2a, PGD2, LXB4, LXA4) showed increased levels in the supernatant from simvastatin-treated primary human myoblasts compared to DMSO control. At 96 h, the pro-inflammatory PGD2, PGF2a and 8-iso-PGF2a levels in simvastatin decreased again matching low PTGS1 mRNA expression levels after 96 h whereas the inflammatory resolving LXA4 continued to increase. LXB4 was detected only from 6 h to 36 h. Long-chain ω-3 eicosapentaenoic acid derived PGE3 and the PUFA 13-HODE were completely absent in simvastatin-treated primary human myoblasts at any time point. PGE3 showed higher levels after rosuvastatin treatment compared to controls after 96 h. Inflammatory resolving ω-3 docosahexaenoic acid derived eicosanoid RvD2 levels were only detectable in the supernatant from simvastatin-treated primary human myoblasts and dropped down to zero at 96 h. This suggests that long-chain ω-6 PUFA metabolites were affected in simvastatin-treated primary human muscle cells.

We then tested six different eicosanoid species at concentrations reported in the literature (Supplementary Table S11) for their effect on proliferation and differentiation of primary human muscle cells during statin exposure. We identified AL-8810, an 11β-fluoro receptor analog of prostaglandin 2α, but not prostaglandin 2α, to significantly reverse the effect of simvastatin on proliferation and differentiation (Figure 4D). AL-8810 had no effect on rosuvastatin treated myotubes (Supplementary Figure S5A-C). 5,6-EET, another arachidonic acid metabolite, increased proliferation independently from the statin effect (8Z,14Z). EDA, found in human milk, showed a negative trend on proliferation without impact on myotube fusion. Other eicosanoids, such as 12-HETE, did not have a significant effect on statin-treated primary human muscle cells.

Thus, eicosanoid species from arachidonic acid influence primary human muscle cell proliferation and fusion. Depending on statin type, eicosanoids have potential to ameliorate statin-induced changes in primary human myotubes.

## 3. Discussion

In our comprehensive expressome analysis and molecular and cellular assessment of muscle related statin effects we discovered that statins have profound and unexpected effects on the transcriptome and proteome of primary human muscle cells. Some cellular functions such as RNA synthesis and turnover were much more influenced by the lipophilic simvastatin than by the hydrophilic rosuvastatin, but cholesterol and fatty acid metabolism, mitochondrial metabolism as well as eicosanoid synthesis pathways were severely affected irrespective of the class of statin that was delivered. The phenotype could be rescued by the addition of eicosanoids.

We were surprised to find a dramatic effect of statins on cholesterol and eicosanoid synthesis in primary human muscle cells. We investigated the cellular gene set to synthesise cholesterol and found that muscle and liver cells (HepG2) share the same repertoire for cholesterol biosynthesis (i.e. HMGCS1, HMGCR, CYP51A1, DHCR7) and transport (i.e. ABCA1, ABCG1, LDLR). Besides SCARB1, cholesterol biosynthesis proteins have been positively detected in human skeletal muscle (https://www.proteinatalas.org) [53]. It possibly points towards a so far underestimated role of skeletal muscle for their own cholesterol homeostasis^24^. It might be possible that muscle synthesizes cholesterol to the same or a larger degree as liver. The human body consists of more than 25% muscle mass, whereas the liver only makes up 2-3%. If cholesterol synthesis is an integral part of muscle function, it would be interesting to speculate whether muscular cholesterol is also secreted into the blood circulation or whether skeletal muscle only produces cholesterol for its own needs. It will be the role of future experiments employing *in vivo* microdialysis^25^ together with metabolomic plus lipidomic analysis that will provide the necessary answers.

Genes involved in cholesterol metabolism like HMGCR, HMGCS1, LDLR, and PCSK9 are increased under statin treatment in primary human muscle cells^26^. If HMG-CoA synthesis is increased but its reduction inhibited by statins, HMG-CoA levels increase and accumulate^27^. Therefore, HMG-CoA might reconvert into ketone metabolites, namely acetoacetyl-CoA and Ac-CoA. Acetoacetyl-CoA synthetase and Ac-CoA acetyltransferase expression are increased suggesting ketolysis or Ac-CoA formation. Muscle is not known to be able to produce ketone bodies whereas liver would be. In rat hepatocytes, high Ac-CoA concentrations trigger insulin resistance *in vivo*^28^. Excess Ac-CoA has been also observed in diabetes patients as insulin negatively regulates gluconeogenese by suppressing Ac-CoA [34]. Indeed, clinical observations associate statins with an increased risk for diabetes mellitus type-2^29-31^. Ac-CoA is a central metabolite. Multiple metabolic effects are possible and are intermingled between each other. Increased Ac-CoA would shuttle into the tricarboxylic acid cycle, carnitine-palmitoyl-transferases (CPT1) as well as fatty acid and triglyceride biosynthesis (Figure 5). All pathways appear affected by statins in muscle as we find increased protein levels of Ac-CoA carboxylase alpha, a very prominent upregulation of CPT1, and an influence on key enzymes for fatty acid synthesis and mitochondrial ß-oxidation (Supplementary Table S8, Figure 5). On the one hand, Ac-CoA carboxylase alpha, an ACC family member, converts Ac-CoA into malonyl-CoA (Mal-CoA), the primary substrate for fatty acid biosynthesis. Mal-CoA inhibits CPT1 for ß-oxidations. On the other hand, Ac-CoA can be metabolised by ACC and fatty acid synthase via transformation of palmitic acid and various fatty acids into acyl-CoA for ß-oxidation. Active CPT1 shuttles acyl-CoA into the inner mitochondrial vortex for ß-oxidation^31^. In rodents, high CPT1 levels have been shown to induce mitochondrial overload by acyl-carnitine accumulation with excess ß-oxidation leading to insulin resistance^32^ and mitochondrial stress^33^. High CPT1 values in statin-treated primary human myotubes may hint towards altered ß-oxidation and mitochondrial impairment. Insulin resistance under statin treatment would be interesting for further studies.

In the cytosol, fatty acid synthases, desaturases, and elongases synthesize long-chain fatty acids by using Mal-CoA and Ac-CoA. Fatty acid desaturases catalyse the formation of arachidonic acid and eicosapentanoic acid, the precursor of prostaglandins (PG) and eicosanoids. These molecules have multiple effects including influences on cell proliferation and differentiation^34^, inflammation, mitochondrial function, and being pain mediators, a very frequent complaint of statin-users. The degree of statin influence on prostaglandin and eicosanoid synthesis was striking at molecular level, i.e. on fatty acid desaturases, elongases, de-/carboxylases (see Supplementary Table S8, Figure 5). We concentrated our further investigation on the formation of prostaglandins (PG) and eicosanoids. As the next important enzyme being significantly downregulated, PTGS1 (COX1) converts ω-6 derived arachidonic acid into biological active eicosanoids and PGs, i.e. PGE2, PGD2, and PGF2a. PTGS is interesting because nonsteroidal anti-inflammatory drugs such as ibuprofen inhibit PTGS and are relevant for pain management^35^. PTGS-inhibitors influence on human skeletal muscle is controversially reported^34,36-38^. In older patients performing resistance exercise under PTGS-inhibition treatment, Type I fiber size increased up to 28% compared to control group^38^. Beneficial effects on skeletal muscle performance have been reported for PTGS up-regulation at mRNA level^39^. Decreased satellite cell fusion after PTGS1 inhibition has been observed in rats^40^. We found that PTGS1 expression is associated with reduced differentiation of simvastatin-treated primary human myotubes. PTGS1 is suggested as the only source for PGF2a^41^ derived from arachidonic acid. PGF2a acts pro-inflammatory but at the same time suggested to promote murine myoblasts fusion <u>and</u> proliferation^42,43^. Its inhibition is proposed to significantly diminish murine myoblast fusion as suggested by Horsely et al^42^. Contrary to the data in animals, Brewer et al. found no significant effect in human skeletal muscle with PGF2a metabolite inhibition and any negative effect diminished over time^34^. In our study, treatment with PGF2a did not significantly change myoblast proliferation and differentiation under statin treatment but PGF2a secretion initially increased under simvastatin treatment together with PGD2, another proinflammatory PG. In a clinical study, eccentric resistance exercise initially revealed high PGF2a levels in human skeletal muscle biopsy specimens^44^. PGD2 metabolites are also linked to DMD suggesting a role in muscle cell pathology^45^. Our findings support data obtained from human skeletal muscle^36^ rather than from mice and rats^40,42^. Inflammation resolving metabolites derived from ω-6 arachidonic acid derived (lipoxins LXA4, LXB4), ω-3 docosahexaenoic acid (RvD2), and ω-3 eicosapentanoic acid (PGE3) were found secreted after statin treatment. A specific role for these metabolites in muscle cells is still unknown. For PGE3, anti-proliferative activities have been reported for tumour cells^46^. Peroxidation of ω-6 fatty acids via arachidonate 15-lipoxygenase produces 13-HODE. Its complete absence for simvastatin samples was surprising. Beside its activating capability for peroxisome proliferator-activated receptors and G protein coupled receptor, 13-HODE is speculated for mitochondria dysfunction and altered calcium homeostasis in epithelia cells^47^. There is currently no information for 13-HODE in skeletal muscle cells but may be another intriguing candidate to be studied in myoblast fusion. Taken together, lipid profiles from supernatant of statin-treated primary human muscle cells under fusion condition show increase in ω-6 arachidonic acid metabolites. However, supplements with ω-3 eicosapentanoic acid and docosahexaenoic acid, but not ω-6, are reported to improve skeletal muscle damage, stiffness and regeneration^48-50^. *In vivo* studies in elderly human probands and insulin-resistant non-diabetic patients, ω-3 PUFA supplementation had no effect on mitochondrial function^48,51^ but improved mitochondrial ADP kinetics and insulin sensitivity in young, healthy adults^52^. Pre-treatment with ω-3 fatty acid eicosapentanoic acid but not docosahexaenoic acid stimulates fatty acid ß-oxidation and glucose transport in human myotubes^53^. Here, we showed that AL-8810, a PGF2a-receptor agonist^34^, positively influences statin-induced effects on muscle cell proliferation and differentiation. We speculate that AL-8810 quenches excess levels of PGF2a, initially secreted by simvastatin treated primary human myoblasts at a critical time point for fusion. We believe that our bench side findings could be translated into the clinic. Very recently, a clinical study showed that eicosapentanoic acid and docosahexaenoic acid added to statin therapy to prevent enlargement of carotid plaques, also significantly reduced severe and less severe musculoskeletal side effects, in particular myalgia^54^. Our study provides an explanation and supports the notion that ω-3 fatty acids should be added to statin therapy either as a prevention or as a treatment of statin myopathy. Patients may be less inclined to discontinue treatment when statin muscular side effects including pain are reduced.

## 4. Materials and Methods

### 4.1. Primary human myoblasts and cell populations

Muscle biopsy specimens for myoblast isolation were obtained from M. vast. lat. after IRB approval by the regulatory agencies (EA1/203/08 and EA2/051/10, Charité Universitätsmedizin Berlin, in compliance with the Declaration of Helsiniki) at the Helios Hospital Berlin Buch, Berlin, Germany, for diagnostic or orthopedic reasons. All study participants provided informed consent. Human muscle biopsy material is restricted regarding availability and quantity, also for ethical reasons. Experiments were conducted from a total of 25 different primary human myoblast cultures (Supplementary Table S1). All cells used in this study were maintained at 37 °C in 5% CO2. For all experiments, except for eicosanoid substitutions, we only used primary muscle cells obtained from statin naïve individuals. Statin-myopathy patients had myalgias (>3 of 10 on a visual analogue scale) and/ or creatine kinase levels elevated to > 250 IU/l under therapy. Isolation was performed as described^55^. The percentage of fibroblasts in myoblast cultures was always below 5% as assessed by anti-desmin staining. Differentiation into myotubes was induced using Opti-MEM (Invitrogen, Germany).

We controlled for confounding factors including cell culture, sample handling, and biopsy-stress variations. At least two to a maximum of four cell populations were used simultaneously in an experimental setup by the same experimenter. In an experimental setup, controls and both statins were completed simultaneously.

### 4.2. Pharmacological reagents

The active form of simvastatin was purchased from ARC (Missouri, USA), rosuvastatin from LKT Laboratories (Minnesota, USA), fatty acid species from Cayman Chemical (Michigan, USA). Geranylgeranyl-pyrophosphate, mevalonolactone, and mevalonic acid were from Sigma Aldrich (Germany). Both statins were dissolved in DMSO and tested for their functionality at different concentrations (1-50µM). (Supplementary Table S11)

### 4.3. Treatment of myoblast and myotubes

Primary human myoblasts were cultured in skeletal muscle growth medium (ProVitro, Berlin, Germany) in 96 well plates (3 × 10^3 cells/ well). Test substances were applied 24 h after plating (Supplementary Table S11). Every 24 h, the cells were refed with fresh medium in the presence or absence of test compounds. All treatments were performed in parallel for each cell population at the same time of the day to exclude circadian rhythm effects. Proliferation was quantified by the colorimetric BrdU assay (Roche, Germany). Toxicity was assessed using the ToxiLight™ Non-destructive Cytotoxicity BioAssay Kit (Lonza, Cologne, Germany), and the Cytotoxicity Detection Kit (Roche, Germany). Amplex® Red Cholesterol Assay Kit (Invitrogen, Germany) was used to measure cholesterol. Mevalonate and cholesterol levels were determined under delipidated medium conditions. The cells were examined after 24 h, 48 h, 72 h, and 96 h. For differentiation into myotubes, primary human myoblasts at a 70% confluence were serum starved with Opti-MEM (Invitrogen, Germany). Selected eicosanoid species were added to myoblast cultures under proliferation and differentiation condition with and without statins. Statins were titrated to a concentration effective for both simvastatin and rosuvastatin using proliferation, toxicity, and mevalonate detection tests as measures. Eicosanoid concentrations were used as suggested by the literature. (Supplementary Table S11, Supplementary Figure S2)

### 4.4. Mevalonic acid in statin-treated primary human myotubes

Together with Lipidomix GmbH (Berlin, Germany), we developed a new and easy method to detect mevalonate intracellular in primary human muscle cells. Cells were lysed by three freeze and thaw cycles and mechanically using 21, 23, and 26 Gauge needles [Lysis buffer: 20 mM Tris-HCl pH 7,4; 150 mM NaCl; 1 mM EDTA; 1 mM PMSF; 1 x Complete Protease-Inhibitor (Roche, Germany)]. Protein concentration was determined using BCA kit (Biorad, Germany). After lysis, proteins were precipitated with 1/10 volume 4 M HClO4 for 10-20 min at 4°C. When pH was neutralised with 5 N KOH, samples were mixed and incubated with 1/5 volume 6 M HCl at 4°C for at least 15 h to promote conversion of mevalonate into mevalonolactone. After centrifugation at 13,000 x g for 2 min, further analysis using HPLC/MS/MS was performed by Dr. Michael Rothe at Lipidomix GmbH, Berlin, Germany. (Supplementary Table S12, Supplementary Figure S6)

### 4.5. RNA analyses

RNA was isolated using the NucleoSpin® RNA/Protein Kit (Marcherey-Nagel, Germany). RNA quantity and purity were determined using a NanoDrop ND-1000 spectrophotometer (Thermo Scientific, USA). For qPCR, the mRNA was reverse transcribed using the QuantiTect reverse transcription kit (Qiagen, Germany). qPCR was performed according to MIQE guidelines^56^ using SYBR Green I in the Mx3000P instrument (Stratagene, Germany) (Primer sequences in Supplementary Table S4) with two reference genes. C_T_-value replicates were checked for consistency and PCR primer efficacy (%) was determined.

### 4.6. RNA sequencing

RNA quality was determined using Agilent 2100 bioanalyzer (Agilent Technologies, USA). The RIN factor was >8 for all samples. cDNA libraries for paired-end sequencing were prepared using the TruSeq Stranded mRNA library preparation kit (Illumina, USA). RNA sequencing was performed at Illumina HiSeq platform (Illumina, USA).

Pre-processed raw RNA-Seq data were imported into CLC Genomics Workbench (v. 7.5.1) and processed using a standardized workflow. The workflow defines the following steps: (1) trimming sequences (2) creating sequence quality check report (3) RNA-Seq analysis with reporting of read tracks, gene expression tracks, transcript expression tracks, fusion tracks, and un-mapped reads. All samples passed the RNA-seq quality check (Supplementary Figure S1). Trimmed reads were aligned to human reference genome (GRCh37) and mapped back to the human transcriptome (v. 19). Mapped read counts were statistically analyzed using the R packages EdgeR and DESeq for matched sample data^57^. Samples without any and with DMSO treatment were considered as reference group. Gene expression data from simvastatin and rosuvastatin treated samples with FDR<0.05 (Bonferroni) were considered as statistically significantly different from control groups. Data are stored at Gene Expression Omnibus - GSE107998.

Transcriptional profiling can be influenced by unintentional factors. We controlled for confounding factors including individual differences, muscle biopsy variations, and sample composition. The muscle cell population were obtained from age-matched donors from the same ethnic group during an orthopaedic surgery. Gender specific genes were removed for RNAseq statistic. qPCR analyses verified RNAseq expression results of selected candidate genes.

### 4.7. Deep proteome analysis

Proteins were prepared according to Sapcariu et al^58^. Samples were measured by LC-MS/MS on a Q-Exactive orbitrap mass spectrometer (Thermo Scientific, Germany). For the data analysis, the MaxQuant software package version 1.4.1.2 was used, with a multiplicity of 3 for dimethylation analysis (modified epsilon amino groups of lysine plus modified N-terminal amino groups)^59^. Carbamidomethylation was set as a fixed modification while oxidized methionine and acetylated N-termini were set as variable modifications. An FDR of 0.01 was applied for peptides and proteins search was performed using a human Uniprot database (August 2013). Normalized ratios of the protein groups output were used to determine proteins which are obviously regulated. (Supplementary Figure S1)

### 4.8. Cell quantification

Myotube fusion and maturation were quantified using an LSM 700 AxioObserver Z1 (Zeiss, Germany) and an N-Achroplan 10x/0.25 M27 objective. Image files were exported and converted in batch into black-white images with adjusted gray-scale using IrfanView (v.4.38). Separate image files for MYHCf and HOECHST 33258 were loaded into the CellProfiler software (v 2.1.1)^60,61^. A pipeline for calculating multinucleated cells was used and slightly modified^62^. A myotube was defined having more than three nuclei. The fusion index was calculated by the number of nuclei in myotubes divided by the total number of nuclei counted in each image. The maturation index is described by the average number of nuclei per myotube.

### 4.9. Data visualization for RNA-Seq and deep proteome

For analyses with mapped count data apart from the EdgeR or DESeq package, mapped read counts were exported from CLC Genomics Workbench (v 7.5.1) and normalized for their library size using the trimmed mean of M-values normalization approach (TMM)^63^. Differential gene expression data were plotted as dendogram clustered heatmaps using GENE-E (v 3.0.206)^64^. Plot.ly was used to visualize quality check data. Principal component analysis was performed in discriminate gene expression data for each treatment group with ClustVis (release June 2016)^65^. Venny 2.0 was used to generate the Venn diagram^66^. The Circos-Plot for omics data was generated using the Bioconductor package OmicCircos^67^.

### 4.10. Pathway analyses

Statistically significant, differential gene expression data from simvastatin and rosuvastatin treated myotubes and controls were further analysed with GOrilla (release March 2013), Reactome (v 53), and DAVID (v 6.7) to identify enriched GO terms and pathway clustering^68-70^. Gene lists were ranked regarding their expression value. In addition, the web-based Ingenuity pathway analysis application (IPA; Ingenuity Systems, Redwood City, CA) was used.

### 4.11. Statistics

All statistics for RNA-Seq and proteome data analyses were performed using RStudio (v 0.98.1006) for executing R packages (v 3.1.1). Statistical analyses for proliferation, toxicity, and qPCR data were performed with GraphPad Prism (v 8.0) using the One-way ANOVA corrected with Holm-Sidak / Dunn’s method. Two-way ANOVA for multiple testing was used for all data obtained from statin, rescue experiments, and fatty acid species treated primary human muscle cells. Each comparison stands for its own. Data obtained from western blots with protein from primary human myotubes were analysed for statistical significance using two-tailed Mann-Whitney test. Values of p are shown by asterisks using the standard convention: * p≤0.05; ** p ≤ 0.01; *** p ≤ 0.001; **** p ≤ 0.0001

## Supporting information

NatMetab_Supplementary Data_Grunwald-Spuler

Supplementary_TablesS2-S6_DEG_Transcriptome_Proteome

Supplementary-Table_S10_Eicosanoid-profile

Supplementary-Table S13_Eicosanoid-species-detection-transitions

Data-file-S1_Full-Transcriptome_Simvastatin

Data-file-S2_Full-Transcriptome_Rosuvastatin

Data-file-S3_Proteome_Simvastatin-Rosuvastatin

Data-File-S4_GO-Terms_Proteome_Simvastatin_Rosuvastatin

Circos-Plot_Grunwald_Spuler

## Data availability statement

All primary data are available upon science-based request.

Bulk RNAseq data are available at NCBI GEO accession: GSE107998 https://www.ncbi.nlm.nih.gov/geo/query/acc.cgi?acc=GSE107998

## Author contributions

S.A.G, O.P., S.P., N.J., K.S., G.P., G.D. performed experiments and analysed data. U.G., E.S-T. recruited patients. W.K. performed genetic analysis. S.A.G., E.S-T., S.S. designed and organized the study and analysed data. S.A.G. and S.S. wrote the manuscript.

## Acknowledgments

We thank all patients for participating in the study. We thank Friedrich C. Luft for discussing the manuscript and Susanne Wissler for help with Endnote®. For his expertise in eicosanoid and beneficial discussions in this field we particularly thank Wolf-Hagen Schunck. This work was funded by the German Research Society (DFG) KFO192/2. None of the authors have any competing interests.

