## Supplementary material for "A prostaglandin alpha F2 analog protects from statin-induced myopathic changes in primary human muscle cells": NatMetab_Supplementary Data_Grunwald-Spuler

**Content**

1. **Supplementary Experimental Procedures**
2. **Supplementary Figures**
3. **Supplementary Tables**
4. **Supplementary Data Files description**
5. **Supplementary References**
1. **Supplementary Experimental Procedures**

**Buffers and antibodies**

Opti-MEM, GlutaMax, and Amplex® Red Cholesterol Assay Kit were purchased from Invitrogen (Germany). 2-(N-morpholino)ethanesulfonic acid buffered saline (MBS) buffer was composed of 25 mM Mes, pH 6.5, and 0.15 M NaCl. Antibodies against cyclophilin A (#ab41684) and GAPDH (#ab9484) were obtained from Abcam (Germany) and MYHI was delivered by Sigma Aldrich (Germany). Primary immunoblot antibodies were labelled with IRDye 800 (Rockland) and ECL™ IgG, HRP linked (GE-Helthcare). For immunofluorescence, secondary antibody were Alexa 568 and Alexa 488 (Invitrogen, Germany). Nuclei were counterstained with Hoechst 33258 (Sigma Aldrich, Germany). Confocal images were obtained using Zeiss LSM 700 (Jena, Germany). Pre-stained protein marker PageRuler™ Plus from Thermo Scientific (Germany) was used in Western blots.

**Delipidation of fetal calf serum (FCS)**

The FCS delipidation protocol was followed as described by Dr. Christoph Thiele in lipidomicnet.org and provides a simple method to reduce cholesterol, triglycerides and most other lipids but not proteins^1^. The delipidated FCS was added to skeletal muscle basal medium (ProVitro, 500 mL) that was supplemented with fetuin (50 µg), human rec. EGF (10 ng), human rec. bFGF (1 ng), insulin (10 μg), dexamethason (400 ng), and gentamicin (50 μg).

**Mevalonolactone HPLC/MS/MS measurements**

Mevalonic acid is unstable after cell lysis and was converted into its stable lactone (MVAL) as described in the experimental procedure in this paper. Dr. Michael Rothe at Lipidomix GmbH, Berlin, Germany established a method for measuring MVAL in muscle cell lysates. MVAL samples were centrifuged with 3000 U/min, 5 min. 50 µL supernatant were pipetted into an Eppendorf tube, 50 µL MVAL-D3 (CDN Isotopes, Pointe-Claire, Canada) 50 ng/mL as internal standard were added and mixed vigorously. The mixture was filtered through a syringe filter (RC, 0.45 µm, Phenomenex, Aschaffenburg, Germany) into an autosampler glass vial. The HPLC/MS/MS analysis was performed using an Agilent 6490/ 1290 HPLC-Triplequad-mass spectrometer (Agilent Technologies, Santa Clara, USA). A Phenomenex Synergy Hydro-RP 150 x 2.1 mm, 3.5 µm was used as stationary phase. The solvent system consisted of methanol (solvent A) and 10 mM ammonium acetate in water (solvent B). The gradient started at 100 % B, hold for 1 min, 62.5 % B after 4 min and 5 % B after 4.1 min.

The injection volume was 2 µL. The mass spectrometer was equipped with a Jetstream electrospray source, operated in positive mode. Drying gas temperature and flow were 140°C / 14 L/min, sheath gas 400 °C / 10 L/min. Capillary and nozzle voltage were optimized at 2500 V / 300 V. The mass spec parameters were optimized for each transition of target compound and internal standard as shown in SuuportingTable S3. Calibration was calculated with respect to the internal standard and was linear from 0.2 to 64 ng/mL. Calibration curve shows linear regression from 1-64 ng/mL MVA (Supplementary Figure S6). Detected peaks from calibrator as well as MVAL at lower and higher rates are presented in Supplementary Figure S6B-D.

**Conventional PCR conditions**

HMGCR transcription variants were identified using mRNA from HepG2 and primary human myotubes. (Supplementary Table S9). For primers 1-6 the PCR Taq "all inclusive" kit (PeqLab) was applied. The Advantage 2 PCR kit (Clontech) was used for PCRs with primerpair 7.

**Real-time PCR setup**

Selected RNA-seq expression data were verified using qPCR. (Supplementary Table S9) For PTGS1 with seven transcript variants we designed primers which mainly amplifies variants one, three, and seven referred to ENSEMBL database version GRCH38. The real-time PCR reaction was prepared using KAPA SYBR FAST qPCR Mastermix Universal (PeqLab, Germany). The PCR cycle conditions were 10 min at 95°C, 40 cycles at 95°C for 10 sec, 57°C for 20 sec, 72°C for 20 sec (single acquisition). The annealing temperature for MYHI was 58°C. The melting curve was performed at 95°C for 1 min, 45°C for 1 min, followed by increasing the temperature to 95°C using a ramp rate at 0.1°C/sec (continuous acquisition). Each sample was set up in duplicates and repeated twice with a difference in C_T_-values less than or equal 0.25. The r²-factor for each efficiency-test was >0.9.

qPCR results were analysed using MxPro (v. 4.1). The relative quantification was calculated in Excel using the ΔΔC_T_-method, which can be used for PCR efficiencies greater or equal 1.98 and lower or equal 2.01. For *HMGCR*, *LDLR*, *MYHI, PTGS1, and PTGS2* with an efficiency out of this range, the efficiency corrected ΔΔC_T_-method after Michael Pfaffl was applied^2^.

**RNA sequencing data quality check and mapping**

For RNA sequencing a total of 16 cDNA libraries derived from four individual myoblast cell populations treated with statins, DMSO and control were sequenced using paired end sequencing (101 base pairs per read). We received a mean total of ~25 699 957 fragment counts (±6.43%). After trimming of nucleotides below a quality score of 20, 89.5% (±1.15%) of the fragments were successfully aligned to the human genome hg19 with 91.3% (±1.02%) mapped to exon regions. Supplementary Figure S1A summarizes the basic quality distribution of all RNA-seq reads showing an equal quality for all samples sequenced.

**Deep proteome sample preparation**

Proteins were prepared according to Sapcariu et al^3^. Briefly, cells were fixed in situ by adding 400 µL ice-cold methanol followed by 400 µL ice-cold water. Afterwards the cells were collected from the plate by using a rubber policeman and transferred into a reaction vial. After the addition of 400 µL ice-cold chloroform and mixing, the protein containing phase was isolated and washed with methanol. After drying the pellet, proteins were resuspended by sonication in denaturation buffer (6 M urea, 2 M thiourea, 10 mM HEPES-KOH pH 8.0). Insoluble material was removed by ultra-centrifugation at 100’000 x g. Protein concentrations were determined by Bradford assay and 10 µg of each sample were taken for in-solution digestion.

For tryptic in-solution digestion, samples were reduced with 1 mM tris(2-carboxyethyl) phosphine (TCEP) and free sulfhydryl groups carbamido-methylated using 5.5 mM choloroacetamide. Proteins were digested with 0.5 µg sequencing grade endopeptidase LysC (Wako Chemicals GmbH, Germany) for 3 h, diluted with four volumes of 50 mM ammonium-bicarbonate and 1 µg sequencing grade trypsin (Promega, Germany). After 10 h incubation the reaction was stopped with trifluoroacetic acid (TFA) to a final concentration of 1%. Peptides were purified by using C18 stage-tips (3 M)^4^. The digest was performed in an automated fashion on an xt PAL (CTC analysis)^5^.

The eluted peptides were lyophilized in a speed-vac and reconstituted with 100 µL 100 mM triethylammonium bicarbonate buffer (TEAB) and dimethyl-labelled in solution using an automated setup^5,6^. The pooled samples were fractionated using strong-anion exchange (SAX) material (3M) on stage tips producing 4 fractions (elution at pH 11, pH 6 and pH 3) including the flow-through. In order to remove residual background noise, a further clean-up using strong-cation exchange (SCX) chromatography was performed by loading the sample on SCX stage-tips, washing with 0.1% TFA in 5% acetonitrile and eluting with 500 mM ammonium acetate buffer, 0.5% acetic acid in 50% acetonitrile followed by C18 stage-tipping of the combined elution fractions.

We measured all samples by LC-MS/MS on a Q-Exactive orbitrap mass spectrometer (Thermo Scientific, Germany) connected to a Proxeon nano-LC system (Thermo Scientific, Germany) in data-dependent acquisition mode selecting the top 10 peaks for HCD fragmentation. A volume of 5 μL was injected and a three-hour gradient (solvent A: 5% acetonitrile, 0.1% formic acid; solvent B: 80% acetonitrile, 0.1% formic acid) was applied eluting peptides at 4% to 76% acetonitrile gradient using an in-house prepared nano-LC column (0.074 mm x 250 mm, 3 µm Reprosil C18, Dr Maisch GmbH, Germany) at a flow rate of 0.25 μL/min. MS acquisition was performed at a resolution of 70’000 in the scan range from 300 to 1700 m/z. Dynamic exclusion was set to 30 sec and the normalized collision energy was specified to 26.

**Western Blot**

Proteins from human muscle cells were obtained with NucleoSpin® RNA/Protein Kit (Marcherey-Nagel, Germany) with minor changes. After protein precipitation of the flow through, the pellet was washed with MBS buffer and stored in protein solving buffer including 1x Complete EDTA-free protease inhibitor cocktail (Roche, Germany). Proteins were separated in 8-16% Tris-Glycine gradient gels and electrophoretically transferred to nitrocellulose membranes (Whatman, Dassel, Germany). After blocking in 5% milk/ TBS/ 0.05% Tween-20, membranes were incubated overnight at 4°C in antibody solutions (MYHI 1:1000; cyclophilin A 1:5000; GAPDH 1:10000). Immunoblots were incubated for 1 h with secondary antibodies (IgG HRP-linked 1:2000, IRDye 800 1:5000). Protein bands were determined with infra-red scanner (LI-COR, Germany) and ECL (SuperSignal West Dura ECL solution, Thermo Scientific, Rockford, USA; ChemiSmart 5000, VILBER LOURMAT, Germany). Quantitation was performed using Image Studio Lite ver5.2 and ImageJ 1.49v.

**Immunofluorescence**

After 84 h - 96 h differentiation, cells were fixed with 3.7% PFA and permeabilised with 0.2% Triton X-100. After blocking with 3% BSA in PBS cells were incubated with MYHI antibody (Sigma, Germany, 1:1000 in 1% BSA). After washing, cells were labelled with secondary antibody Alexa 568 (Invitrogen, Germany, 1:500 in 1% BSA in PBS) and HOECHST 33258, and mounted with 90% glycerol (w/v) in PBS with 0.25% DABCO (w/v).

**Eicosanoid profile**

Statins or DMSO was applied to primary human myoblasts 24 h after plating (Supplementary Table S11) and switched to differentiation conditions. Supernatant was collected after 6 h, 12 h, 18 h, 36 h, and 96 h and directly transferred to dry ice. Protein content was assessed from adherent primary human muscle cells for each sample. Eicosanoid profiles were measured by Dr. Michael Rothe at Lipidomix GmbH, Berlin, Germany.

Sample preparation for analysis of PUFA-derived lipid mediators and metabolites

The supernatants were spiked with an internal standard consisting of 14,15-EET-d8, 14,15-DHET-D11, 15-HETE-d8, 20-HETE-d6, LTB4-d4, PGE2-d2 10 ng each (Cayman Chemical, Ann Arbor, USA), 500 µL methanol was added and shaked vigorously. The samples were brought to pH 6 with 500 µL 1 M sodium acetate buffer. After centrifugation, the obtained supernatant was added to the Bond Elute Certify II columns (Agilent Technologies, Santa Clara, USA) for Solid phase Extraction, which were preconditioned with 3 mL methanol, followed by 3 mL of 0.1 mol/L sodium acetate buffer containing 5 % methanol (pH 6). The SPE-columns were then washed with 3 mL methanol/H2O (50/50, vol/vol). For elution 2 mL of n-hexane:ethyl acetate 25:75 with 1 % acetic acid was used. The extraction was performed with a SPE Vacuum Manifold. The eluate was evaporated on a heating block at 40 °C under a stream of nitrogen to obtain a solid residue.

LC/ESI-MS/MS

The residues were dissolved in 70 µL acetonitrile and analyzed using an Agilent 1290 HPLC system with binary pump, autosampler and column thermostat with a stationyry phase Agilent Zorbax Eclipse plus C18, 2.1 x 150 mm, 1.8 µm (Agilent, Santa Clara, CA) column using a solvent system of aqueous acetic acid (0.05 %) and acetonitrile/methanol 50:50 %. The elution gradient was started with 5 % acetonitrile, which was increased within 0.5 minutes to 55, 14.5 minutes to 69 %, 14.6 minutes to 95 % and held there for 5.4 minutes. The flow rate was set at 0.3 mL/min, the injection volume was 7.5 µL. The HPLC was coupled with an Agilent 6490 Triplequad mass spectrometer (Agilent Technologies, Santa Clara, USA) with electrospray ionisation source. The source parameters were: Drying gas: 250 °C/10 L/min, Sheath gas: 400 °C/10 L/min, Capillary voltage: 4500 V, Nozzel voltage: 1500 V and Nebulizer pressure: 30 psi. Analysis was performed with Multiple Reaction Monitoring in negative mode. For details see Supplementary Table S13.

2. **Supplementary Figures**

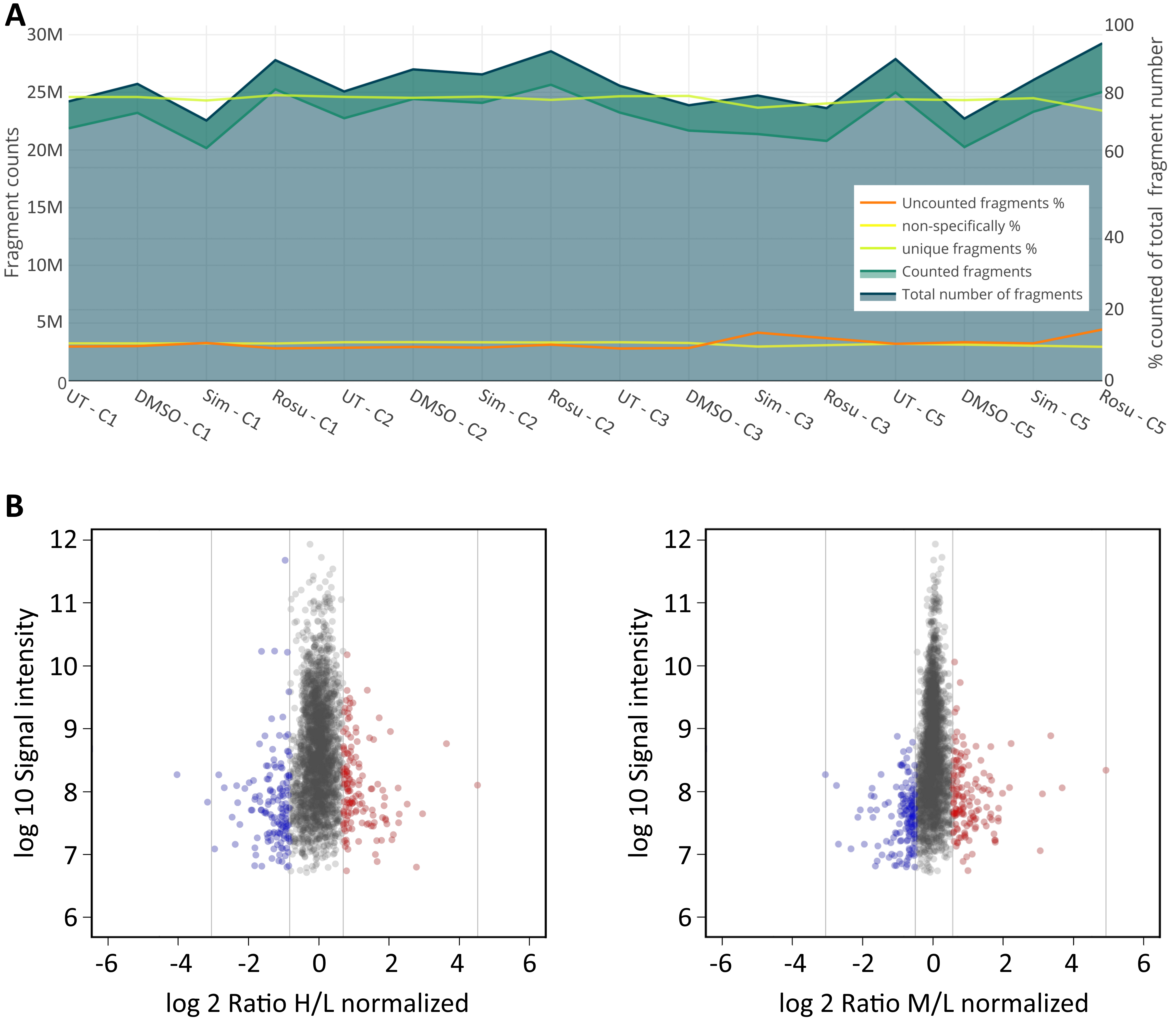

**Supplementary Figure S1.** Transcriptome and proteome analyses (A) Quality statistic of RNA-seq reads. (related to Figure 1) Length distribution of RNA-seq fragments and mapping of fragments to the human transcriptome for all 16 samples. (B) . MA-plot of differently expressed proteins from deep proteome data. (related to Figure 1 and Supplementary Video) The multiplex stable isotope dimethyl-labelling method (heavy (H), medium (M), light (L)) was applied with DMSO as control labelled with light stable isotopes. Simvastatin was labelled with heavy and rosuvastatin with medium stable isotopes. Data were analysed with MaxQuant software package version 1.4.1.2. and plotted as log10 of the intensities over log2 of the ratios. For each experiment, a ratio heavy over light or medium over light is given. The ratios reflect the abundance of the proteins in the respective experiments. (n=4) For cell populations used see Supplementary Table S1. [UT, untreated; Sim, 5 µM simvastatin; Rosu, 5 µM rosuvastatin; plotted using: Plotly Technologies Inc. Collaborative data science. Montréal, QC, 2015. https://plot.ly.]

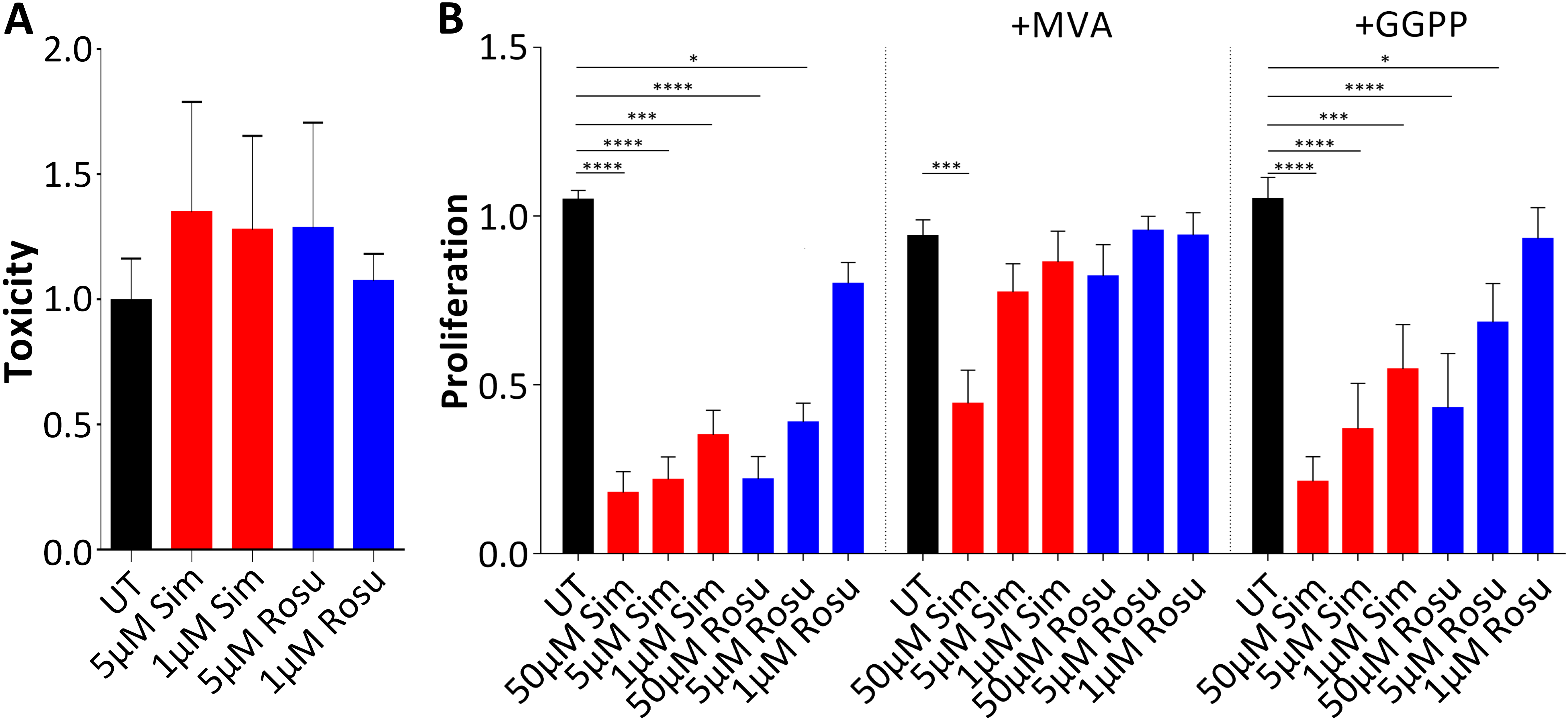

**Supplementary Figure S2.** (related to Figure 4) Statin cytotoxicity and concentration dependent impact on human myoblast proliferation. (A) No cytotoxicity was detected in human myoblasts after 72 h statin treatment. Data were calculated relatively to DMSO-treated samples. (n=6) (B) Cell proliferation (BrdU) test in human myoblasts treated with statins, mevalonate and geranylgeranyl-pyrophosphate for 72h. Myoblast proliferation is negatively affected by statins in a concentration dependent manner. Mevalonate supplementation shows full rescue of statins negative cell proliferation effect (except for 50µM simvastatin) and geranylgeranyl-pyrophosphate to some extent. (n=6-8) For cell populations used see Supplementary Table S1. [UT, untreated (black); Sim, simvastatin(red); Rosu, rosuvastatin (blue)]

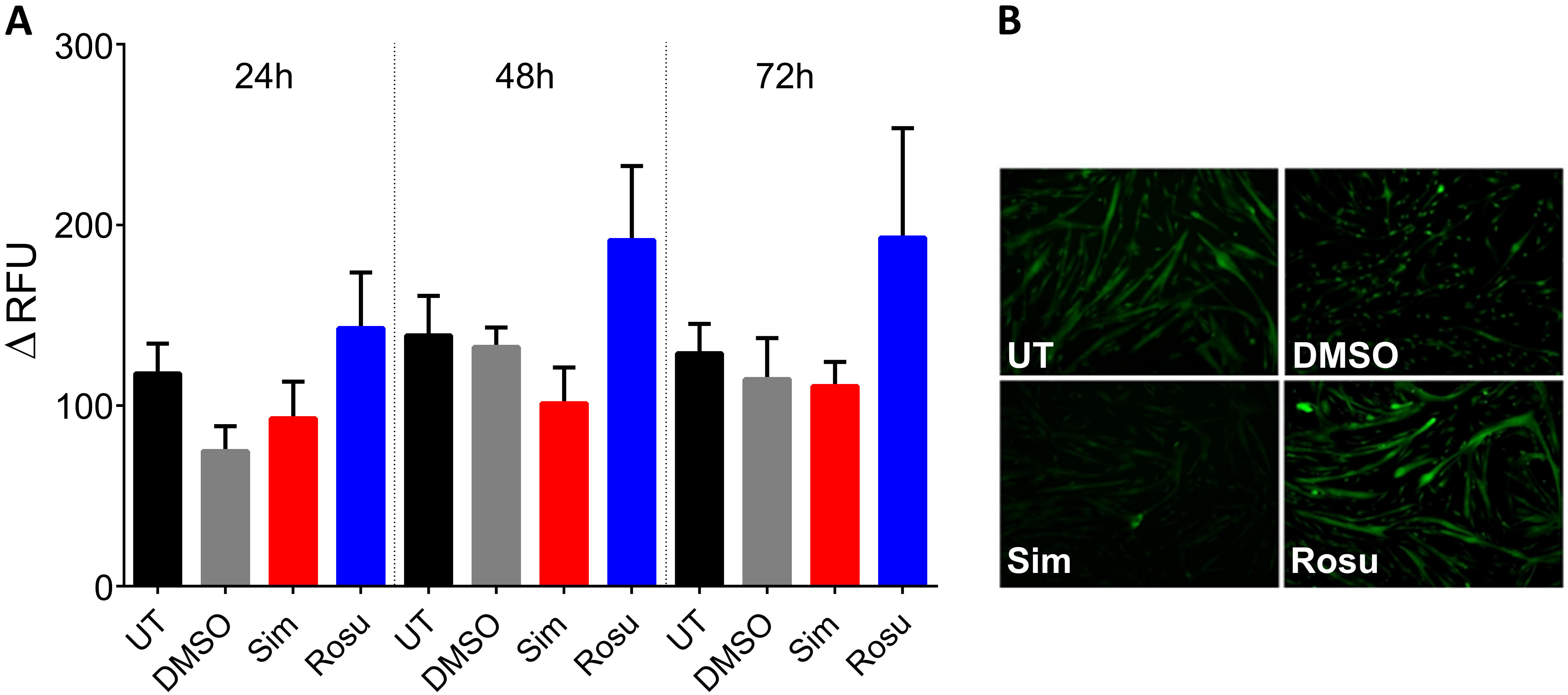

**Supplementary Figure S3.** Nitric oxide (NO) levels in myotubes detected with DAF-2DA. (related to Figure 4 and Supplementary Figure S1) (A) NO quantification in primary human myotubes (n=4, 3 replicates each) treated with statins for 24 h to 72 h under differentiation conditions. Results were normalized to protein content. Error bars = mean with SD. (B) Microscopic observation of intracellular NO in primary human myotubes after 72h statin-treatment. For cell populations used see Supplementary Table S1. [UT, untreated (black); Sim, 5 µM simvastatin (red); Rosu, 5 µM rosuvastatin (blue); DAF-2DA 4,5-Diaminofluorescein diacetate]

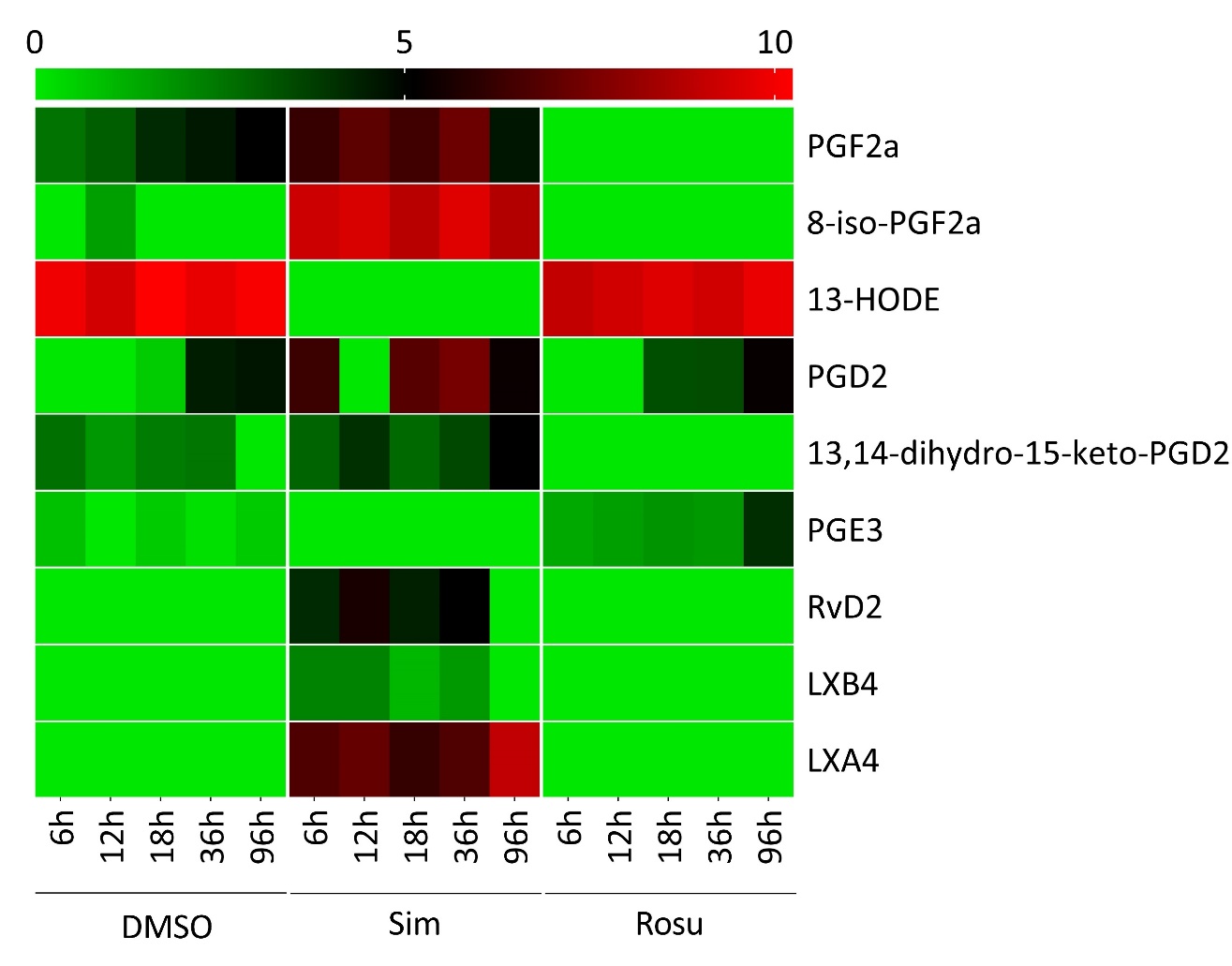

**Supplementary Figure S4:** Long-chain ω-3 and ω-6 PUFAs and their metabolites in the supernatant from statin-treated primary human muscle cells and DMSO controls under fusion conditions. Primary human myoblasts supernatants were analysed at myoblast fusion onset (after 6 h-96 h). Except for PGE3, the supernatant derived from rosuvastatin-treated primary human myoblasts showed eicosanoid profiles comparable to DMSO control. Levels of arachidonic acid derived eicosanoid species PGF2a, 8-iso-PGF2a, PGD2, LXB4, and LXA4 were higher whereas eicosapentaenoic acid derived PGE3 and PUFA 13-HODE levels were absent in simvastatin-treated samples. Concerning ω-3 docosahexaenoic acid derived eicosanoids, RvD2 levels were only detected in the supernatant from simvastatin-treated primary human myoblasts. (5 time points with n=1 each) [Sim, simvastatin; Rosu, rosuvastatin; log2 transformed values; AA, arachidonic acid; DHA, docosahexaenoic acid; 13-HODE, 13-hydroxyoctadecadienoic acid; EPA, eicosapentaenoic acid; PUFA, poly unsaturated fatty acid; PGD2, prostaglandin D2; PGE3, prostaglandin E3; PGF2a, prostaglandin F2 alpha; LXA4/ B4, lipoxin A4/ B4; RvD2, resolvin D2] See Supplementary Table S13 for full eicosanoid profile. For cell population used see Supplementary Table S1.

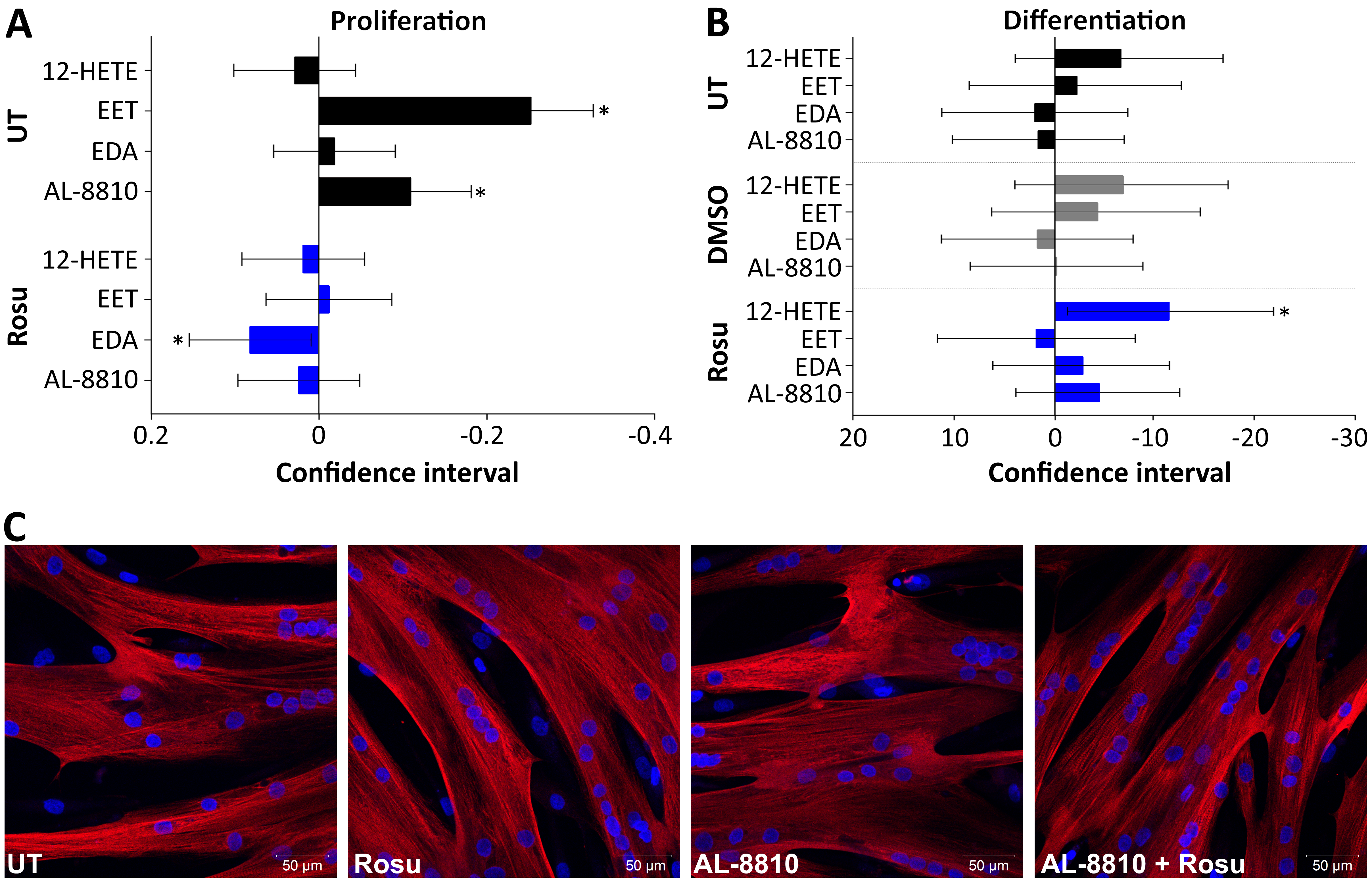

**Supplementary Figure S5.** Proliferation and differentiation of myoblasts under rosuvastatin and/or eicosanoid species treatment. (related to Figure 4) (A) Proliferation of human myoblasts under rosuvastatin and/or fatty acid treatment. Data were calculated relatively to DMSO. (95% confidence interval; n=4); (B and C) Differentiation of primary human myoblasts into myotubes under treatment with rosuvastatin and/or fatty acids. Myotube fusion index were determined by counting MYHI (red) positively immunolabelled myotubes and HOECHST (blue) stained nuclei using the cell image analysis software CellProfiler v.2.1.1. Human rosuvastatin-treated myotubes showed normal morphology with and without eicosanoid species added to the medium. (95% confidence interval; n=8 with 2-4 immunofluorescent pictures counted each) For cell populations used see Supplementary Table S1. [UT, untreated (black); Rosu, 5 µM rosuvastatin (blue); MYHI, myosin heavy chain 1]

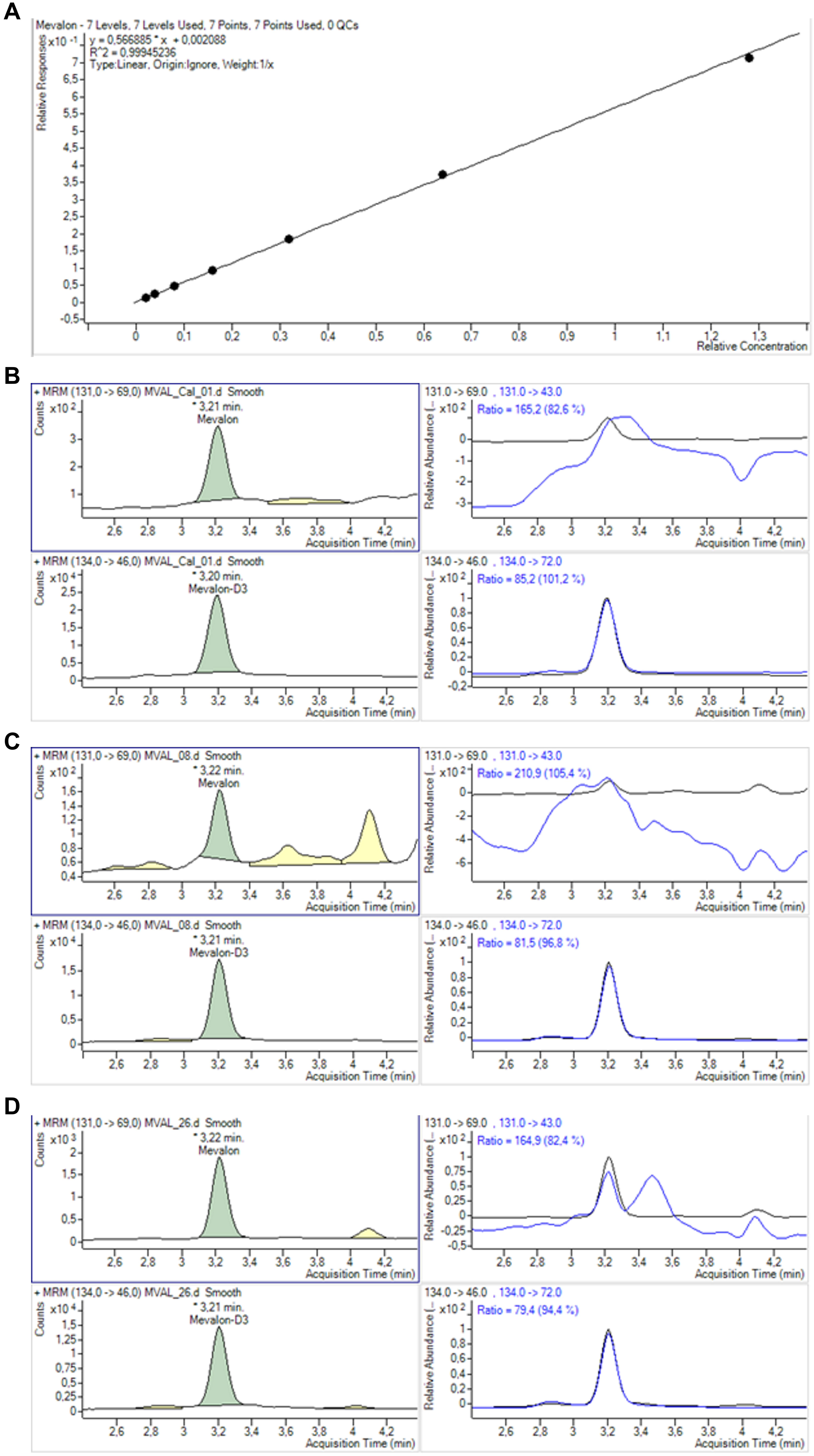

**Supplementary Figure S6.** (related to Figure 2) Specification for mevalonolactone detection in primary human myoblasts. (A) Calibration was calculated with respect to the internal standard and was linear from 0.2 to 64 ng/mL. (B) Lower limit of quantification was calculated from signal to noise ratio (peak-to-peak) of the lowest calibrator as 0.2 ng/mL. Mevalonolactone detected in matrix at (C) lower range (0.34 ng/mL, S/N = 17) and (D) higher range (11.5 ng/mL, S/N = 95). [MVAL, mevalonolactone]

3. **Supplementary Tables**

**Supplementary Table S1.** Source of primary human myoblasts: Clinical information and myopathological findings. Patients C1-C19 were statin-naïve. For statin-treated patients (S1-S6), we excluded myotonic dystrophy type 2 by molecular genetic analysis. (W Kress, Institute for Human Genetics, Würzburg, Germany)

| **Number of cell population** | **Sex of donor** | **Age of donor years** | **Medical history of donor** | **Myopathological findings in muscle biopsy specimens*^a^*** | **Cell population used in Figure** |
| --- | --- | --- | --- | --- | --- |
| C1 | m | 67 | No neuromuscular symptoms, hip replacement surgery | Normal | 1; 3; S2; S3; |
| C2 | m | 76 | No neuromuscular symptoms, hip replacement surgery | Normal | 1; 2; 3; S2; S3; |
| C3 | f | 71 | No neuromuscular symptoms, hip replacement surgery | Normal | 1; 2; 3; S2; S3; |
| C4 | f | 74 | No neuromuscular symptoms, hip replacement surgery | Mild type 2 fiber atrophy | 2; 3; 4; |
| C5 | f | 59 | No neuromuscular symptoms, hip replacement surgery | Normal | 1; 3; 4; S2; S3; S4; |
| C6 | m | 78 | No neuromuscular symptoms, hip replacement surgery | Normal | 3; 4; S4; |
| C7 | f | 82 | No neuromuscular symptoms, hip replacement surgery | Normal | 4; |
| C8 | m | 67 | No statins, myalgias of unknown origin, CK 464 U/L | Normal | 4; |
| C9 | m | 35 | No statins, myalgias of unknown origin, CK 300 U/L | Normal | 2; 4; |
| C10 | f | 58 | No neuromuscular symptoms, hip replacement surgery | Normal | 3; 4; S4; |
| C11 | m | 79 | No neuromuscular symptoms, hip replacement surgery | Normal | 2; 4; S1; |
| C12 | f | 43 | No statins, myalgias of unknown origin, CK normal | Normal | 2; 4; S1; |
| C13 | m | 80 | No neuromuscular symptoms, hip replacement surgery | Normal | 2; 3; 4; S1; S4; |
| C14 | f | 87 | No neuromuscular symptoms, hip replacement surgery | Type 2 fiber atrophy | 4; S1; |
| C15 | f | 75 | No neuromuscular symptoms, hip replacement surgery | Normal | 4; S1; |
| C16 | m | 58 | No neuromuscular symptoms, hip replacement surgery | Normal | 4; |
| C17 | m | 64 | No neuromuscular symptoms, hip replacement surgery | Not available | 4; |
| C18 | w | 80 | No neuromuscular symptoms, hip replacement surgery | Not available | 4; S1; |
| C19 | m | 51 | No statins, myalgias of unknown origin, CK normal | Normal | S4 |
| S1 | m | 55 | Statin myopathy; Simvastatin 10 mg/d;  CK 895 U/L | Normal | 4; |
| S2 | m | 55 | Statin myopathy; Rosuvastatin 10 mg;  CK 1435 U/L | Normal | 3; 4; S4; |
| S3 | m | 77 | Statin-Patient without side effects; Simvastatin 10 mg; CK 195 U/L | Normal | 4; |
| S4 | f | 20 | Statin myopathy; Simvastatin 10 mg/d;  CK 6260 U/L | Normal | 3; 4; S4; |
| S5 | f | 56 | Statin myopathy; Pravastatin 10 mg;  CK 780 U/L | Some internal nuclei | 3; 4; S4; |
| S6 | m | 77 | Statin myopathy; Pravastatin 5 mg;  CK 1900 U/L | Mild neurogenic atrophy | 3; 4; S4; |

*^a^*All biopsies were obtained from M. vastus lateralis

**Supplementary Table S2 – S6: Supplementary_TablesS2-S6_DEG_Transcriptome_Proteome.pdf**

Supplementary Table S2 RNA-seq Simvastatin DEG FDR<0.05

Supplementary Table S3 RNA-seq Rosuvastatin DEG FDR<0.05

Supplementary Table S4 HMGCR LDLR PCSK9 Transcript variants

Supplementary Table S5 Proteome Simvastatin

Supplementary Table S6 Proteome Rosuvastatin

**Supplementary Table S7: Genes significantly differently expressed in both statin groups and at RNA and protein level.** [FC = fold change; n=4]

| **Gene name** | **Biological process** | **RNA Sim (FC)** | **Protein Sim (FC)** | **RNA Rosu (FC)** | **Protein Rosu (FC)** |
| --- | --- | --- | --- | --- | --- |
| CLSTN2 | Cell adhesion | 0.552 | 0.397 | 0.635 | 0.306 |
| CYP51A1 | Cholesterol biosynthetic process | 2.148 | 2.232 | 2.925 | 3.190 |
| FDFT1 |  | 1.997 | 1.632 | 2.567 | 2.341 |
| IDI1 |  | 2.098 | 2.135 | 2.483 | 1.582 |
| TUBB2B | Microtubule-based process | 1.673 | 0.353 | 1.896 | 1.570 |

**Supplementary Table S8.** Expression data at RNA and protein level from features shown in Figure 5. [ND = not detected, FC = fold change; n=4]

| **Feature** | **RNA (FC)** | | **Protein (FC)** | |
| --- | --- | --- | --- | --- |
|  | **Simvastatin** | **Rosuvastatin** | **Simvastatin** | **Rosuvastatin** |
| ACC | 1.2 | 1 | ND | ND |
| ACOT8 | 2 | 1 | ND | ND |
| ACSL | 0.8-1.8 | 0.8-1.2 | 0.7-1.4 | 0.8-1.4 |
| ACSS2 | 2.6 | 2.8 | 1.4 | 1.5 |
| AGPAT | 0.8-1.3 | 0.9-1.1 | 0.5-1 | 0.7-1.3 |
| AMPK | 0.7-1.4 | 0.7-1.2 | ND | ND |
| CD36 | 0.4 | 1 | 0.6 | 1 |
| CPT1 | 2.1 | 1.1 | 4.8 | 12.8 |
| CPT2 | 0.9 | 1 | 0.8 | 0.8 |
| DGAT2 | 0.6 | 1 | ND | ND |
| ELOVL | 0.5-2.2 | 0.9-2.3 | ND | ND |
| FADS1 | 2 | 1.8 | ND | ND |
| FADS2 | 1.8 | 2.5 | 0.8 | 1.8 |
| FASN | 1.3 | 2 | 1.2 | 1.3 |
| HMGCR | 2.3 | 2.9 | ND | ND |
| HMGCS | 3.26 | 3.91 | 1.0532 | 1.7971 |
| HSD17B | 0.21-2.9 | 0.8-2.7 | 0.9-1.2 | 0.9-1.1 |
| LPIN1 | 1.8 | 1.9 | ND | ND |
| MCEE | 1.6 | 0.9 | ND | ND |
| PPAR | 1.3-1.7 | 1.0-1.5 | ND | ND |
| PTGS1 | 0.43 | 0.87 | ND | ND |
| PTGS2 | 2.75 | 1.16 | ND | ND |
| RXRA | 2.2 | 1 | ND | ND |
| SLC25A20 | 0.6 | 0.9 | ND | ND |
| SLC27A1 | 1.4 | 1.1 | ND | ND |

**Supplementary Table S9.** Sequence information and PCR conditions for real time PCR primer pairs and HMGCR transcript variant analyses in HepG2 and primary human muscle cells (based on NM_000859.2). Cyclophilin A, GAPDH, and RPL13a act as reference genes. PTGS1-001-var1 is based on ENST00000632012.6 and PTGS1-202-var7 on ENST00000540753.5. [GAPDH, glyceraldehyde 3-phosphate dehydrogenase; HMGCR, 3-hydroxy-3-methylglutaryl-CoA reductase; PTGS, prostaglandin-endoperoxide synthase; RPL13a, ribosomal protein L13a]

| **Gene** | **Position** | **Sequences** | **Expected length** | **Efficiency** |
| --- | --- | --- | --- | --- |
| Cyclophilin A  NM_021130.3 | Exon 1 | CGCCGAGGAAAACCGTGTACTATT | 115bp | 1.99 |
|  | Exon 1/2 | GACCTTGTCTGCAAACAGCTCAAAG |  |  |
| GAPDH  NM_002046.5 | Exon 2 | GAAGGTGAAGGTCGGAGTC | 226bp | 1.99 |
|  | Exon 4 | GAAGATGGTGATGGGATTTC |  |  |
| RPL13a  NM_012423.3 | Exon 6 | CGTGCGTCTGAAGCCTACA | 227bp | 1.99 |
|  | Exon 8 | GGAGTCCGTGGGTCTTGAG |  |  |
| HMGCR  NM_000859.2 | Exon 13 | TGTTGGAGTGGCAGGACCCCTT | 127bp | 1.96 |
|  | Exon 13/14 | GGCACCTCCACCAAGACCTATTGC |  |  |
| HMGCS  NM_002130.6 | Exon 5/6 | TGCCCAGTGGCAGAAAGAGGGA | 128bp | 1.99 |
|  | Exon 6 | GGAAGTCATTCAGCAACATCCGAGC |  |  |
| LDLR  NM_000527.4 | Exon 8 | CTGCAGCCAGCTCTGCGTGA | 108bp | 2.05 |
|  | Exon 8/9 | GCGATGGAGCCCACAGCCTT |  |  |
| MYHI  NM_005963.3 | Exon 29 | TAGTTTCACAGCTCTCGAGGGGC | 101bp | 2.18 |
|  | Exon 29/30 | TGCCAGGGCACTCTTGGCCTTTATC |  |  |
| PTGS1 all variants  NM_000962.3 | Exon 5/6 | GGAACCAAAGGGAAGAAGCA | 138bp | 2.08 |
|  | Exon 6 | GAACTGGTGGGTGAAGTGTT |  |  |
| PTGS1-001-var1  NM_000962.3 | Exon 9 | CCAGGAGTACAGCTACGA | 105bp | 2.08 |
|  | Exon 9/10 | ATCCGGCCAGCAATCTG |  |  |
| PTGS1-var3  NM_001271164.1 | Exon 4 | GCTGGTTCTGGGAGTTTGT | 76bp | 2.14 |
|  | Exon 4/5 | GCTTCTTCCCTGTGAGTACC |  |  |
| PTGS1-202-var7  NM_001271368.1 | Exon 3 | GCAGGCACCAAGGGAAA | 147bp | 2.00 |
|  | Exon 3 | CATCCCGGCTCCTAAATGT |  |  |
| PTGS2  NM_000963.3 | Exon 3/4 | TGAGTTATGTGTTGACATCCAG | 137bp | 1.96 |
|  | Exon 4 | ATCATCAGGCACAGGAGGAA |  |  |
| **HMGCR PCR primers and conditions** | | | | |
| **Primer number** | **Position** | **Sequences** | **Annealing temperature** | **Elongation time** |
| 1 | Exon 1 | GGGTTCGGTGGCCTCTAGTGAGAT | 61,5°C | 40sec |
|  | Exon 5 | TGTGCTTGCTCTGGAAAGGTCAA |  |  |
| 2 | Exon 3 | ACAATAACACGATGCATAGCC | 55°C |  |
|  | Exon 8 | AAACTCGGGCAAAATGGCTG |  |  |
| 3 | Exon 6 | ACGTTTACCCTCGATGCTCTTGT |  |  |
|  | Exon 11 | TTCATTAGGCCGAGGTTCCCTG |  |  |
| 4 | Exon 9 | TGTTCATGCTCACAGTCGCTGG |  |  |
|  | Exon 14 | TCCCATCTGCAAGGACTCGGCT |  |  |
| 5 | Exon 12 | GCGTGGTGTATCTATTCGCCGACA |  |  |
|  | Exon 17 | TGCATGGGCGTTGTAGCCTCCT |  |  |
| 6 | Exon 15 | TCAGGGGATGCCATGGGGATGA | 54°C | 2min |
|  | Exon 20 | TAGGCTCGGCAAGCAAGCCA |  |  |
| 7 | Exon 1 | TTCCGCTCCGCGACTGCGTT | 68°C | 3min |
|  | Exon 20 | CAGTCACCAACCTCCTGGCCACA |  |  |

**Supplementary-Table S10:** Full Eicosanoid profile secreted from statin treated primary human myoblasts under fusion conditions. (5 time points with n=1 each) See Supplementary-Table_S10_Eicosanoid-profile.xlsx

**Supplementary Table S11.** Statins and eicosanoid species used in primary human myoblasts under proliferation and differentiation conditions. Eicosanoid concentrations were used as suggested by the literature. [HMG-CoA, 3-hydroxy-3-methylglutaryl coenzyme A; HMGCR, 3-hydroxy-3-methylglutaryl-CoA reductase]

| **Substance** | **Concentration** | **Site of metabolic/ pharmacological effects** |
| --- | --- | --- |
| **Statins** | | |
| GGPP | 2,5µM | Isoprenoid, precursor for protein geranylation and prenylation |
| Mevalonic acid | 25µM | Reduced form of HMG-CoA |
| Rosuvastatin | 1µM – 50µM | Hydrophilic statin, HMGCR inhibitor |
| Simvastatin | 1µM – 50µM | Hydrophobic statin; HMGCR inhibitor |
| **Eicosanoid species** | | |
| AL-8810 | 1 µM | Prostaglandin F2 alpha agonist^7^ |
| (8Z,14Z) EDA | 5 µM | Modulates effects on pro-inflammatory mediators^8^ |
| 5(6)-EET | 0.1 µM | Modulates calcium release and contractility^9^ |
| Prostaglandin F2alpha | 1 µM | Cell survival, via Prostaglandin F receptor^7^ |
| 20-HETE | 1 µM | Pro-inflammatory; promotes proliferation^10,11^ |
| (±)12-HETE | 1 µM | Pro-inflammatory, involved in myogenesis^12,13^ |

**Supplementary Table S12.** Mevalonolactone mass spec parameters optimized for each transition of target compound and internal standard. Calibration was calculated with respect to the internal standard and was linear from 0.2 to 64 ng/mL. [MVAL, Mevalonolactone; CE, collision energy; CAV, cell accelerator voltage]

| Compound | Precursor | MS1 Res | Product | MS2 Res | Dwell | CE *^a^* | CAV *^b^* |
| --- | --- | --- | --- | --- | --- | --- | --- |
| MVAL | 131 | wide | 69 | wide | 75 | 4 | 1 |
| MVAL | 131 | wide | 43 | wide | 75 | 23 | 8 |
| MVAL-D3 | 134 | wide | 72 | wide | 75 | 4 | 1 |
| MVAL-D3 | 134 | wide | 46 | wide | 75 | 23 | 8 |

**Supplementary Table S13:** Multiple Reaction Monitoring mode and transitions of eicosanoid species. See Supplementary-Table S13_Eicosanoid-species-detection-transitions.xlsx

4. **Supplementary data files 1 – 4**

**Supplementary Data File S1** (related to Figure 1-3, 5, Supplementary Figure S5). Differential expressed genes at transcriptome levels from simvastatin treated myotubes relative to DMSO controls. (n=4) **Data-file-S1_Transcriptome_DEG_Simvastatin.txt**

**Supplementary Data File S2** (related to Figure 1-3, 5, Supplementary Figure S5). Differential expressed genes at transcriptome levels from rosuvastatin treated myotubes relative to DMSO controls. (n=4) **Data-file-S2_Transcriptome_DEG_Rosuvastatin.txt**

**Supplementary Data File S3** (related to Figure 1-3, 5, Supplementary Figure S6). Proteome data from simvastatin and rosuvastatin treated myotubes compared to DMSO controls. (n=4) **Data-file-S3_Proteom_Simvastatin-Rosuvastatin.txt**

**Supplementary Data File S4** (related to Figure 4) GO-Terms for significantly different genes and proteins in statin-treated primary human muscle cells (obtained with GOrilla). **Data-File-S4_GO-Terms_Proteom_Simvastatin_Rosuvastatin.xlsx**

**Supplementary Video** (related to Figure 1 and 3) OmicCircos plot of differentially expressed genes and proteins in statin-treated human muscle cells. (n=4) **Circos-Plot_Grunwald_Spuler.mp4**

5. **Supplementary References**

1	Thiele, C., Hannah, M. J., Fahrenholz, F. & Huttner, W. B. Cholesterol binds to synaptophysin and is required for biogenesis of synaptic vesicles. *Nat. Cell Biol.* **2**, 42-49, doi:10.1038/71366 (2000).

2	Pfaffl, M. W. A new mathematical model for relative quantification in real-time RT-PCR. *Nucleic Acids Res.* **29**, e45 (2001).

3	Sapcariu, S. C. *et al.* Simultaneous extraction of proteins and metabolites from cells in culture. *MethodsX* **1**, 74-80, doi:10.1016/j.mex.2014.07.002 (2014).

4	Rappsilber, J., Mann, M. & Ishihama, Y. Protocol for micro-purification, enrichment, pre-fractionation and storage of peptides for proteomics using StageTips. *Nat. Protoc.* **2**, 1896-1906, doi:10.1038/nprot.2007.261 (2007).

5	Kanashova, T. *et al.* Differential proteomic analysis of mouse macrophages exposed to adsorbate-loaded heavy fuel oil derived combustion particles using an automated sample-preparation workflow. *Anal. Bioanal. Chem.* **407**, 5965-5976, doi:10.1007/s00216-015-8595-4 (2015).

6	Boersema, P. J., Raijmakers, R., Lemeer, S., Mohammed, S. & Heck, A. J. Multiplex peptide stable isotope dimethyl labeling for quantitative proteomics. *Nat. Protoc.* **4**, 484-494, doi:10.1038/nprot.2009.21 (2009).

7	Horsley, V. & Pavlath, G. K. Prostaglandin F2(alpha) stimulates growth of skeletal muscle cells via an NFATC2-dependent pathway. *J. Cell Biol.* **161**, 111-118, doi:10.1083/jcb.200208085 (2003).

8	Pereira, D. M., Correia-da-Silva, G., Valentao, P., Teixeira, N. & Andrade, P. B. Anti-inflammatory effect of unsaturated fatty acids and Ergosta-7,22-dien-3-ol from Marthasterias glacialis: prevention of CHOP-mediated ER-stress and NF-kappaB activation. *PLoS One* **9**, e88341, doi:10.1371/journal.pone.0088341 (2014).

9	Nieves, D. & Moreno, J. J. Epoxyeicosatrienoic acids induce growth inhibition and calpain/caspase-12 dependent apoptosis in PDGF cultured 3T6 fibroblast. *Apoptosis* **12**, 1979-1988, doi:10.1007/s10495-007-0123-3 (2007).

10	Lu, S. *et al.* Effect of 20-hydroxyeicosatetraenoic acid on biological behavior of human villous trophoblasts and uterine vascular smooth muscle cells. *Mol Med Rep* **9**, 1889-1894, doi:10.3892/mmr.2014.2017 (2014).

11	McCarthy, E. T., Sharma, R. & Sharma, M. Protective effect of 20-hydroxyeicosatetraenoic acid (20-HETE) on glomerular protein permeability barrier. *Kidney Int.* **67**, 152-156, doi:10.1111/j.1523-1755.2005.00065.x (2005).

12	Siangjong, L., Gauthier, K. M., Pfister, S. L., Smyth, E. M. & Campbell, W. B. Endothelial 12(S)-HETE vasorelaxation is mediated by thromboxane receptor inhibition in mouse mesenteric arteries. *Am. J. Physiol. Heart Circ. Physiol.* **304**, H382-392, doi:10.1152/ajpheart.00690.2012 (2013).

13	Zhou, Y. P. *et al.* Apoptosis in insulin-secreting cells. Evidence for the role of intracellular Ca2+ stores and arachidonic acid metabolism. *J. Clin. Invest.* **101**, 1623-1632, doi:10.1172/jci1245 (1998).
