## Supplementary_TablesS2-S6_DEG_Transcriptome_Proteome for "A prostaglandin alpha F2 analog protects from statin-induced myopathic changes in primary human muscle cells"

Supplementary Table 2 RNA-seq Simvastatin DEG FDR<0.05 ____ 2
Supplementary Table 3 RNA-seq Rosuvastatin DEG FDR<0.05 __ 54
Supplementary Table 4 HMGCR LDLR PCSK9 Transcript variants 57
Supplementary Table 5 Proteome Simvastatin ____________________ 58
Supplementary Table 6 Proteome Rosuvastatin ____________________ 73

Supplementary Table S2. Differentially expressed genes at RNA-level from simvastatin-treated primary human myotubes (FDR<0.05)

| Gene name | Controls normalized mean counts | Simvastatin normalized mean counts | foldChange | pval | padj | Protein name |
|---|---|---|---|---|---|---|
| SEPT4 | 1501.069803 | 801.4972929 | 0.533950714 | 0.00116405 | 0.025355381 | Septin-4 |
| SEPT8 | 1441.801849 | 869.1346625 | 0.60281145 | 0.001020561 | 0.022970752 | Septin-8 |
| SEPT11 | 15446.9098 | 7040.109901 | 0.455761702 | 1.61E-05 | 0.000918517 | Septin-11 |
| 7SK_5 | 15.37250235 | 2.453813586 | 0.159623562 | 0.000193119 | 0.006512156 | |
| A4GALT | 337.0734749 | 647.7282426 | 1.921623299 | 3.99E-05 | 0.001884916 | Lactosylceramide 4-alpha-galactosyltransferase |
| AARD | 16.34273774 | 2.936168593 | 0.179661978 | 0.000385314 | 0.011128664 | Alanine and arginine-rich domain-containing protein |
| AARSD1 | 677.076378 | 414.8116488 | 0.61265119 | 0.001650077 | 0.03285031 | Alanyl-tRNA-editing protein Aarsd1 |
| AATK | 107.0697753 | 36.78265486 | 0.343539106 | 6.61E-05 | 0.002811 | Serine/threonine-protein kinase LMTK1 |
| ABAT | 275.9205401 | 151.5957045 | 0.549417975 | 0.000800454 | 0.019197327 | 4-aminobutyrate aminotransferase, mitochondrial |
| ABCA8 | 499.8487363 | 269.8631157 | 0.539889563 | 0.000170156 | 0.005920437 | ATP-binding cassette sub-family A member 8 |
| ABCB10 | 299.3483499 | 481.8190029 | 1.609559575 | 0.002820587 | 0.049042586 | ATP-binding cassette sub-family B member 10, mitochondrial |
| ABCB4 | 21.30582529 | 54.3599761 | 2.551413773 | 0.001657132 | 0.032895175 | Multidrug resistance protein 3 |
| ABCB9 | 138.8827064 | 54.94028798 | 0.395587683 | 6.29E-05 | 0.002705609 | ATP-binding cassette sub-family B member 9 |
| ABCC3 | 137.1951044 | 343.6263325 | 2.504654478 | 8.40E-07 | 8.14E-05 | Canalicular multispecific organic anion transporter 2 |
| ABCD1 | 1969.667397 | 781.8006913 | 0.396920156 | 2.28E-07 | 2.71E-05 | ATP-binding cassette sub-family D member 1 |
| ABCE1 | 2086.066838 | 1184.803548 | 0.567960492 | 0.000182863 | 0.006245683 | ATP-binding cassette sub-family E member 1 |
| ABCF2 | 3515.65228 | 1991.402957 | 0.566439112 | 0.00014128 | 0.00514598 | ATP-binding cassette sub-family F member 2 |
| ABCG1 | 138.7478852 | 55.14821277 | 0.397470655 | 0.002052767 | 0.038212676 | ATP-binding cassette sub-family G member 1 |
| ABHD3 | 42.20842311 | 122.648491 | 2.905782352 | 0.000288392 | 0.008829437 | cDNA FLJ52972, weakly similar to Homo sapiens abhydrolase domain containing 3 (ABHD3), mRNA |
| ABI3 | 10.8281812 | 49.88598563 | 4.607051239 | 1.35E-05 | 0.000789097 | ABI gene family member 3 |
| ABR | 1871.163621 | 3757.169846 | 2.007932286 | 1.91E-06 | 0.000158586 | cDNA FLJ54747, highly similar to Active breakpoint cluster region-related protein |
| ABTB1 | 519.6104715 | 881.5494112 | 1.69655821 | 0.000522543 | 0.014119048 | Ankyrin repeat and BTB/POZ domain-containing protein 1 |
| AC002066.1 | 40.10237568 | 8.568605257 | 0.213668271 | 3.95E-05 | 0.001871068 | |
| AC002117.1 | 90.87222631 | 239.7803571 | 2.63865393 | 0.001155924 | 0.025214773 | |
| AC002463.3 | 29.14019146 | 7.816251739 | 0.268229251 | 0.000664623 | 0.016712823 | |
| AC004041.2 | 122.0963524 | 227.1615798 | 1.860510779 | 0.000465097 | 0.012894214 | |
| AC004112.4 | 167.4996187 | 298.5961525 | 1.782667655 | 0.00066058 | 0.016638049 | |
| AC005013.1 | 45.71762421 | 16.34237587 | 0.357463367 | 0.002491365 | 0.044461503 | |
| AC005027.3 | 328.3891791 | 84.26356492 | 0.256596655 | 2.81E-13 | 1.68E-10 | |
| AC005592.2 | 101.4891551 | 50.66029425 | 0.499169534 | 0.001527709 | 0.031079739 | |
| AC006460.2 | 90.9547424 | 38.62806085 | 0.424695402 | 0.002486593 | 0.044445622 | |
| AC007228.11 | 280.2546707 | 459.0730292 | 1.638056658 | 0.001895232 | 0.035919228 | |
| AC007796.1 | 106.3847416 | 490.3962437 | 4.609648305 | 0.000581903 | 0.015266 | |
| AC009041.2 | 48.90001985 | 97.3395313 | 1.990582654 | 0.001914774 | 0.036163306 | |

| Gene name | Controls normalized mean counts | Simvastatin normalized mean counts | foldChange | pval | padj | Protein name |
|---|---|---|---|---|---|---|
| AC009133.20 | 116.0774353 | 490.4602582 | 4.225285105 | 1.13E-07 | 1.50E-05 | |
| AC009492.1 | 186.5152469 | 16.8019044 | 0.090083276 | 4.18E-06 | 0.000304706 | |
| AC010226.4 | 92.95251997 | 229.1465679 | 2.465200168 | 5.14E-07 | 5.42E-05 | |
| AC010970.2 | 1182.337019 | 17389.68498 | 14.70789184 | 1.15E-55 | 1.20E-51 | |
| AC011530.4 | 323.0503702 | 592.1147703 | 1.832886834 | 0.000135315 | 0.004957719 | |
| AC011747.7 | 19.38196496 | 53.30554582 | 2.750265308 | 0.000934512 | 0.0215805 | |
| AC016747.3 | 85.14028949 | 41.37769307 | 0.485994273 | 0.001134485 | 0.024886665 | |
| AC018647.3 | 840.9429372 | 222.1116231 | 0.26412211 | 0.000145896 | 0.005289343 | |
| AC018890.6 | 932.1042053 | 527.5690089 | 0.565997885 | 0.00030059 | 0.009077806 | |
| AC022596.6 | 1.570127796 | 16.46521327 | 10.4865434 | 0.000110439 | 0.00423591 | |
| AC025335.1 | 401.0579155 | 157.4717785 | 0.392640994 | 0.000274099 | 0.008475257 | |
| AC037459.4 | 184.3927076 | 97.63396508 | 0.529489297 | 0.00060751 | 0.015714443 | |
| AC073115.7 | 163.4556742 | 19.74747111 | 0.120812393 | 5.27E-12 | 2.22E-09 | |
| AC073283.4 | 255.649435 | 125.9739252 | 0.492760429 | 0.000522894 | 0.014119048 | |
| AC079922.3 | 27.86471568 | 79.13921936 | 2.840122981 | 0.000795911 | 0.019147424 | |
| AC093388.3 | 5525.579029 | 9657.408071 | 1.747763994 | 0.001791017 | 0.034326162 | |
| AC093673.5 | 190.6130609 | 103.2857499 | 0.541860822 | 0.000614285 | 0.015791705 | |
| AC096669.2 | 49.16574591 | 9.813631133 | 0.199603015 | 0.000509679 | 0.013870572 | |
| AC104532.2 | 77.36551653 | 177.1309296 | 2.289533341 | 1.21E-05 | 0.000716899 | |
| AC104695.3 | 24.22380896 | 62.70945326 | 2.588752799 | 0.000445332 | 0.012502266 | |
| AC104809.3 | 2.761599056 | 17.0356372 | 6.168758336 | 0.000733063 | 0.018083005 | |
| AC112229.4 | 12.52733469 | 1.427396597 | 0.113942561 | 0.000561114 | 0.014858114 | |
| AC112721.2 | 10.69546318 | 1.242447262 | 0.116165821 | 0.002192679 | 0.040358276 | |
| AC135048.13 | 96.78695739 | 300.3283372 | 3.102983556 | 5.76E-07 | 5.97E-05 | |
| AC144652.1 | 86.5688519 | 194.6452779 | 2.248444719 | 6.27E-05 | 0.002701219 | |
| AC144831.1 | 43.9560005 | 138.8545317 | 3.158943718 | 1.92E-07 | 2.37E-05 | |
| AC147651.3 | 32.32569508 | 118.0579551 | 3.652139722 | 0.000740096 | 0.018227584 | |
| ACAT2 | 1064.936878 | 2495.672704 | 2.343493549 | 0.00032769 | 0.00974611 | Acetyl-CoA acetyltransferase, cytosolic |
| ACOT7 | 1099.701356 | 514.9548654 | 0.468267919 | 1.22E-06 | 0.000112115 | Cytosolic acyl coenzyme A thioester hydrolase |
| ACOT8 | 726.6825833 | 1476.644862 | 2.032035576 | 2.30E-06 | 0.000185534 | Acyl-coenzyme A thioesterase 8 |
| ACOX3 | 800.8442561 | 382.449077 | 0.47755737 | 6.09E-06 | 0.000417313 | Peroxisomal acyl-coenzyme A oxidase 3 |
| ACSS2 | 664.1220082 | 1701.472947 | 2.561988499 | 5.15E-10 | 1.37E-07 | Acetyl-coenzyme A synthetase, cytoplasmic |
| ACTA2 | 22761.40606 | 4056.236354 | 0.178206757 | 5.88E-11 | 2.01E-08 | Actin, aortic smooth muscle |
| ACTB | 40532.69565 | 18879.60271 | 0.465787 | 6.54E-05 | 0.002788137 | Actin, cytoplasmic 1 |
| ACTG2 | 7.637200695 | 0.255427941 | 0.033445231 | 0.000797921 | 0.019180936 | Actin, gamma-enteric smooth muscle |
| ACTN1 | 15057.17784 | 5539.320826 | 0.367885728 | 5.89E-08 | 8.77E-06 | Alpha-actinin-1 |
| ACVR2A | 646.3692657 | 295.2762609 | 0.456822867 | 1.20E-06 | 0.000110603 | Activin receptor type-2A |
| ADAM19 | 6253.971067 | 1277.082167 | 0.204203402 | 3.82E-09 | 8.49E-07 | Disintegrin and metalloproteinase domain-containing protein 19 |
| ADAMTS12 | 604.0498105 | 202.1974087 | 0.334736317 | 1.24E-06 | 0.000113657 | A disintegrin and metalloproteinase with thrombospondin motifs 12 |
| ADAMTS5 | 2076.507554 | 1172.948636 | 0.564866058 | 0.001733692 | 0.033660222 | A disintegrin and metalloproteinase with thrombospondin motifs 5 |
| ADAMTS6 | 200.9877148 | 102.8478976 | 0.511712358 | 0.000644598 | 0.016399421 | A disintegrin and metalloproteinase with thrombospondin motifs 6 |
| ADAMTSL5 | 421.3024452 | 852.7596751 | 2.024103313 | 4.51E-05 | 0.00207994 | ADAMTS-like protein 5 |
| ADCK3 | 1532.047944 | 2670.684743 | 1.74321225 | 0.002212938 | 0.040613531 | Chaperone activity of bc1 complex-like, mitochondrial |
| ADCY9 | 2104.941801 | 4441.556678 | 2.110061511 | 1.75E-06 | 0.000149495 | Adenylate cyclase type 9 |

| Gene name | Controls normalized mean counts | Simvastatin normalized mean counts | foldChange | pval | padj | Protein name |
|---|---|---|---|---|---|---|
| ADD3 | 1452.359067 | 2551.071004 | 1.756501585 | 0.001693736 | 0.033260952 | Gamma-adducin |
| ADHFE1 | 81.3313592 | 186.4513799 | 2.292490642 | 6.77E-05 | 0.002866978 | Hydroxyacid-oxoacid transhydrogenase, mitochondrial |
| ADIRF | 244.5050701 | 449.4335625 | 1.838135963 | 0.000217057 | 0.007084757 | Adipogenesis regulatory factor |
| ADORA1 | 2519.294154 | 965.9907407 | 0.383437059 | 1.11E-06 | 0.000104028 | Adenosine receptor A1 |
| ADPRM | 288.3029375 | 158.2930789 | 0.549051218 | 0.001090348 | 0.024174178 | Manganese-dependent ADP-ribose/CDP-alcohol diphosphatase |
| ADRB2 | 160.1047743 | 72.52916085 | 0.453010606 | 0.000469453 | 0.012956242 | Beta-2 adrenergic receptor |
| ADSSL1 | 142.3357557 | 303.2545061 | 2.130557459 | 0.000199961 | 0.006631129 | Adenylosuccinate synthetase isozyme 1 |
| ADTRP | 176.3949384 | 37.04463945 | 0.210009651 | 2.03E-09 | 4.72E-07 | Androgen-dependent TFPI-regulating protein |
| AFAP1L2 | 86.36572127 | 34.77695728 | 0.402670837 | 0.0004621 | 0.01283868 | Actin filament-associated protein 1-like 2 |
| AFF3 | 51.19885195 | 15.50605457 | 0.302859419 | 7.06E-05 | 0.002969886 | AF4/FMR2 family member 3 |
| AGAP2 | 493.7382513 | 173.1149163 | 0.350620832 | 9.97E-06 | 0.000616519 | Arf-GAP with GTPase, ANK repeat and PH domain-containing protein 2 |
| AGTRAP | 433.5597821 | 749.8331341 | 1.729480374 | 0.000439822 | 0.012392294 | Type-1 angiotensin II receptor-associated protein |
| AHCY | 3760.858933 | 2297.490917 | 0.610895266 | 0.000916361 | 0.021314624 | Adenosylhomocysteinase |
| AHNAK2 | 6675.719832 | 11828.20409 | 1.771824521 | 0.000112877 | 0.004308154 | Protein AHNAK2 |
| AHRR | 1889.345119 | 3870.141862 | 2.048403875 | 1.36E-05 | 0.000792085 | Aryl hydrocarbon receptor repressor |
| AHSA1 | 2862.898195 | 1760.533253 | 0.614947907 | 0.001251613 | 0.026831613 | cDNA FLJ56675, highly similar to Activator of 90 kDa heat shock protein ATPase homolog 1 |
| AIF1L | 6729.528856 | 3246.372626 | 0.482407119 | 0.000113408 | 0.00431784 | Allograft inflammatory factor 1-like |
| AK7 | 9.941095502 | 26.9241959 | 2.708373126 | 0.00288021 | 0.049717978 | Adenylate kinase 7 |
| AKAP1 | 2030.585746 | 3136.765895 | 1.544759142 | 0.002903636 | 0.049983675 | A-kinase anchor protein 1, mitochondrial |
| AKR1B10 | 57.41667791 | 222.2221745 | 3.870341903 | 2.44E-06 | 0.000193734 | Aldo-keto reductase family 1 member B10 |
| AKR1B15 | 7.835874087 | 38.3680399 | 4.896459473 | 4.02E-06 | 0.000294404 | Aldo-keto reductase family 1 member B15 |
| AKR1C1 | 1505.194865 | 3352.855232 | 2.227522369 | 1.28E-07 | 1.67E-05 | Aldo-keto reductase family 1 member C1 |
| AKR1D1 | 18.32676602 | 127.1514989 | 6.938021619 | 3.60E-07 | 3.94E-05 | 3-oxo-5-beta-steroid 4-dehydrogenase |
| ALDH18A1 | 1039.909217 | 616.6819031 | 0.593015133 | 0.00066771 | 0.0167769 | Delta-1-pyrroline-5-carboxylate synthase |
| ALDH1A2 | 79.26977315 | 166.8887942 | 2.105327006 | 0.000680942 | 0.017067988 | Retinal dehydrogenase 2 |
| ALDH1B1 | 3597.22129 | 2150.071759 | 0.597703501 | 0.000959805 | 0.021936581 | Aldehyde dehydrogenase X, mitochondrial |
| ALDH3A2 | 1879.205285 | 3243.634662 | 1.726067231 | 0.00022725 | 0.007332806 | Fatty aldehyde dehydrogenase |
| ALDH3B1 | 973.9996554 | 2413.710434 | 2.478143006 | 1.13E-06 | 0.000105839 | Aldehyde dehydrogenase family 3 member B1 |
| ALOX12B | 0.981038388 | 36.56572871 | 37.27247492 | 1.84E-10 | 5.71E-08 | Arachidonate 12-lipoxygenase, 12R-type |
| ALOX5 | 7.457312551 | 29.53977219 | 3.961181992 | 0.00064471 | 0.016399421 | Arachidonate 5-lipoxygenase |
| ALPL | 523.480778 | 260.2716466 | 0.497194276 | 0.000290768 | 0.00888469 | Alkaline phosphatase, tissue-nonspecific isozyme |

| Gene name | **Controls** normalized mean counts | **Simvastatin** normalized mean counts | **foldChange** | **pval** | **padj** | **Protein name** |
|---|---|---|---|---|---|---|
| ALS2 | 2241.146452 | 1374.649276 | 0.613368785 | 0.001220393 | 0.026362187 | cDNA FLJ31851 fis, clone NT2RP7000624, highly similar to Alsin |
| ALS2CL | 401.4220705 | 734.7925492 | 1.830473716 | 8.87E-05 | 0.003555356 | ALS2 C-terminal-like protein |
| AMMECR1 | 763.5766688 | 358.9905095 | 0.470143372 | 1.46E-06 | 0.000128946 | AMME syndrome candidate gene 1 protein |
| AMOT | 44.4887868 | 109.6603387 | 2.464898385 | 0.000184783 | 0.006295571 | Angiomotin |
| AMOTL2 | 5467.528827 | 2700.182706 | 0.493857973 | 0.001665716 | 0.032981381 | Angiomotin-like protein 2 |
| AMPD3 | 113.3405606 | 623.2921306 | 5.499285755 | 3.07E-10 | 8.73E-08 | AMP deaminase 3 |
| AMY2B | 145.4491284 | 309.3641406 | 2.12695768 | 1.13E-05 | 0.000683175 | cDNA FLJ38286 fis, clone FCBBF3008153, highly similar to ALPHA-AMYLASE 2B (EC 3.2.1.1) |
| ANGPT1 | 137.6426258 | 26.72906093 | 0.19419174 | 1.99E-08 | 3.44E-06 | cDNA FLJ56604, highly similar to Angiopoietin-1 |
| ANGPTL1 | 246.5851098 | 63.55978181 | 0.257760016 | 0.000594339 | 0.015496645 | Angiopoietin-related protein 1 |
| ANGPTL2 | 1786.173049 | 3489.206277 | 1.953453658 | 0.002619062 | 0.046288949 | Angiopoietin-related protein 2 |
| ANGPTL7 | 73.72492359 | 31.70739863 | 0.430077063 | 0.002327807 | 0.042369083 | Angiopoietin-related protein 7 |
| ANK3 | 2548.010618 | 4257.306429 | 1.670835435 | 0.000506861 | 0.013814846 | Ankyrin-3 |
| ANKRD12 | 1330.211574 | 2176.776447 | 1.636413703 | 0.000926603 | 0.021477573 | Ankyrin repeat domain-containing protein 12 |
| ANKRD13A | 2399.686445 | 929.4523566 | 0.387322418 | 4.34E-05 | 0.002022043 | Ankyrin repeat domain-containing protein 13A |
| ANKRD23 | 274.3185069 | 476.2253743 | 1.73603079 | 0.000930767 | 0.021525968 | Ankyrin repeat domain-containing protein 23 |
| ANKRD30BL | 10121.40241 | 132016.0571 | 13.04325742 | 9.47E-52 | 5.89E-48 | |
| ANKRD34A | 302.9031593 | 59.06765108 | 0.195005068 | 1.32E-18 | 1.71E-15 | Ankyrin repeat domain-containing protein 34A |
| ANKRD36C | 122.3499653 | 266.3062177 | 2.176594142 | 0.001397433 | 0.028998366 | Ankyrin repeat domain-containing protein 36C |
| ANKRD40 | 1195.847465 | 2115.52178 | 1.769056542 | 0.000134714 | 0.004941509 | Ankyrin repeat domain-containing protein 40 |
| ANKRD44 | 808.413957 | 369.5841949 | 0.457171962 | 3.35E-05 | 0.001630053 | Serine/threonine-protein phosphatase 6 regulatory ankyrin repeat subunit B |
| ANKRD65 | 45.09646341 | 132.7062101 | 2.942718787 | 3.80E-06 | 0.000279754 | |
| ANKRD9 | 269.6805753 | 743.7331508 | 2.757829888 | 2.33E-10 | 7.05E-08 | |
| ANKS1A | 10783.5345 | 4678.614175 | 0.433866482 | 0.002325903 | 0.042359517 | Ankyrin repeat and SAM domain-containing protein 1A |
| ANXA11 | 2627.335247 | 4502.85694 | 1.713849402 | 0.000194543 | 0.006535054 | Annexin |
| ANXA4 | 938.9998627 | 2169.113838 | 2.310025724 | 8.15E-08 | 1.15E-05 | Annexin |
| ANXA6 | 17866.23006 | 10401.66675 | 0.582197067 | 0.000230111 | 0.007405343 | Annexin |
| AP000688.14 | 13.33403777 | 68.57292173 | 5.142697429 | 2.01E-07 | 2.44E-05 | |
| AP001055.1 | 72.28421367 | 30.64187147 | 0.423908208 | 0.000782473 | 0.018895814 | |
| AP001372.2 | 6.448156801 | 28.5538655 | 4.428221333 | 0.000231079 | 0.007421302 | |
| AP002856.5 | 148.5572189 | 47.29351673 | 0.318352195 | 1.06E-06 | 0.000100381 | |
| AP006621.8 | 72.03607343 | 213.0063045 | 2.95693941 | 0.000132387 | 0.004873415 | |
| AP3B2 | 16.26854427 | 53.27014271 | 3.274425899 | 2.24E-05 | 0.001178117 | AP3B2 protein |
| APEX1 | 3404.507448 | 1983.386991 | 0.582576781 | 0.000306279 | 0.009213844 | DNA-(apurinic or apyrimidinic site) lyase |
| APLP1 | 335.4707474 | 179.5990878 | 0.535364377 | 0.002466616 | 0.044248303 | cDNA FLJ56046, highly similar to Amyloid-like protein 1 (APLP)(APLP-1) |
| APRT | 847.6512726 | 533.8079895 | 0.629749529 | 0.002655454 | 0.046746204 | Adenine phosphoribosyltransferase |
| AQP1_2 | 2939.22898 | 295.9004468 | 0.100672812 | 1.09E-09 | 2.70E-07 | |
| AQP5 | 40.05185069 | 2.935639097 | 0.073295966 | 0.001420232 | 0.029316215 | Aquaporin-5 |

| Gene name | **Controls** normalized mean counts | **Simvastatin** normalized mean counts | **foldChange** | **pval** | **padj** | **Protein name** |
|---|---|---|---|---|---|---|
| AQPEP | 7.793181831 | 25.51510592 | 3.274029334 | 0.001398756 | 0.029006471 | Aminopeptidase Q |
| ARHGAP21 | 1642.409793 | 2663.821951 | 1.621898483 | 0.001002183 | 0.022671935 | Rho GTPase-activating protein 21 |
| ARHGAP23P1 | 71.88127957 | 30.0360612 | 0.417856518 | 0.001111344 | 0.024517354 | |
| ARHGAP27 | 217.730661 | 377.4318702 | 1.733480569 | 0.000994239 | 0.022524972 | Rho GTPase-activating protein 27 |
| ARHGAP28 | 549.4803874 | 273.6419107 | 0.498001233 | 0.000194267 | 0.006532815 | Rho GTPase-activating protein 28 |
| ARHGAP4 | 240.9456229 | 420.967212 | 1.747146127 | 0.001820371 | 0.034747991 | Rho GTPase-activating protein 4 |
| ARHGAP42 | 485.3593017 | 297.0810737 | 0.612084846 | 0.002400217 | 0.043306934 | Rho GTPase-activating protein 42 |
| ARHGAP9 | 258.0231084 | 102.5360605 | 0.397390998 | 8.37E-05 | 0.003399083 | Rho GTPase-activating protein 9 |
| ARHGEF25 | 184.2171195 | 88.64311613 | 0.481188265 | 0.000151611 | 0.005452028 | Rho guanine nucleotide exchange factor 25 |
| ARHGEF3 | 249.531042 | 522.6448696 | 2.094508424 | 3.83E-05 | 0.001822482 | Rho guanine nucleotide exchange factor 3 |
| ARHGEF6 | 6208.3063 | 2672.519239 | 0.430474772 | 3.41E-06 | 0.000256127 | cDNA FLJ50776, highly similar to Rho guanine nucleotide exchange factor 6 |
| ARID5A | 789.7796034 | 2352.599688 | 2.978805325 | 5.60E-13 | 3.01E-10 | AT rich interactive domain 5A (MRF1-like), isoform CRA_a |
| ARL4A | 643.7964735 | 166.4109795 | 0.258483832 | 3.08E-15 | 2.33E-12 | ADP-ribosylation factor-like protein 4A |
| ARL4D | 1464.397882 | 659.9830624 | 0.450685616 | 2.24E-05 | 0.001177188 | ADP-ribosylation factor-like protein 4D |
| ARL6IP6 | 209.6817887 | 341.6944025 | 1.6295855 | 0.002864386 | 0.049533915 | ADP-ribosylation factor-like protein 6-interacting protein 6 |
| ARMCX2 | 4408.770833 | 2013.12223 | 0.456617571 | 1.96E-07 | 2.39E-05 | Armadillo repeat-containing X-linked protein 2 |
| ARNT2 | 1263.837411 | 411.9694709 | 0.325967144 | 3.45E-09 | 7.71E-07 | Aryl hydrocarbon receptor nuclear translocator 2 |
| ARRDC2 | 123.2240219 | 393.6532614 | 3.194614616 | 1.72E-09 | 4.02E-07 | Arrestin domain-containing protein 2 |
| ARSK | 375.9947416 | 212.385519 | 0.564863003 | 0.000451768 | 0.01262596 | Arylsulfatase K |
| ASB14 | 294.9563422 | 1057.849413 | 3.586460983 | 0.001106556 | 0.02444639 | Ankyrin repeat and SOCS box protein 14 |
| ASB16-AS1 | 253.6496725 | 522.6282892 | 2.060433526 | 6.63E-06 | 0.000449074 | |
| ASS1 | 2481.572342 | 1496.95415 | 0.603228093 | 0.000788732 | 0.019004093 | Argininosuccinate synthase |
| ATAT1 | 716.1350097 | 427.6001064 | 0.597094264 | 0.000937044 | 0.021606889 | Alpha-tubulin N-acetyltransferase |
| ATF3 | 117.6409372 | 585.8133453 | 4.979672548 | 5.29E-07 | 5.56E-05 | Cyclic AMP-dependent transcription factor ATF-3 |
| ATG16L2 | 278.9971331 | 596.3812529 | 2.13758918 | 8.64E-06 | 0.000556537 | Autophagy-related protein 16-2 |
| ATHL1 | 146.650107 | 325.9122074 | 2.222379609 | 2.72E-05 | 0.001368674 | Acid trehalase-like protein 1 |
| ATIC | 1451.883601 | 887.421669 | 0.611220946 | 0.001387079 | 0.028821964 | Phosphoribosylaminoimidazolecarboxamide formyltransferase |
| ATL3 | 437.7793437 | 755.6150076 | 1.726017955 | 0.002063751 | 0.038371217 | Atlastin-3 |
| ATP2A1 | 1109.5748 | 2962.468168 | 2.669912987 | 0.000218112 | 0.007111733 | Sarcoplasmic/endoplasmic reticulum calcium ATPase 1 |

| Gene name | **Controls** normalized mean counts | **Simvastatin** normalized mean counts | foldChange | pval | padj | Protein name |
|---|---|---|---|---|---|---|
| ATP2A3 | 56.77661272 | 167.7626843 | 2.954785013 | 2.34E-05 | 0.001212845 | Sarcoplasmic/endoplasmic reticulum calcium ATPase 3 |
| ATP2B4 | 486.795428 | 1450.598518 | 2.979893471 | 4.04E-05 | 0.001909211 | Plasma membrane calcium-transporting ATPase 4 |
| ATP5G1 | 1145.043651 | 660.3767968 | 0.576726308 | 0.000469834 | 0.012956242 | ATP synthase F(0) complex subunit C1, mitochondrial |
| ATP7B | 439.3453229 | 157.0182296 | 0.357391376 | 1.42E-09 | 3.45E-07 | WND/140 kDa |
| ATP8B1 | 261.8548001 | 513.7458825 | 1.961949456 | 4.10E-05 | 0.001930388 | Probable phospholipid-transporting ATPase IC |
| ATP9A | 989.2826907 | 1927.420963 | 1.948301513 | 2.63E-05 | 0.001325752 | Probable phospholipid-transporting ATPase IIA |
| AURKA | 242.3981355 | 141.3859375 | 0.583279806 | 0.002642106 | 0.046608812 | Aurora kinase A |
| AVIL | 62.94305565 | 132.2057186 | 2.100401979 | 0.000198424 | 0.006600877 | Advillin |
| B3GALT2 | 419.0357316 | 41.50998458 | 0.099060728 | 4.87E-07 | 5.19E-05 | Beta-1,3-galactosyltransferase 2 |
| B3GALT4 | 30.79108492 | 101.5330414 | 3.297481777 | 0.000883379 | 0.020738395 | Beta-1,3-galactosyltransferase 4 |
| B4GALNT3 | 273.1513278 | 640.3402503 | 2.34426922 | 0.00043819 | 0.012368724 | Beta-1,4-N-acetylgalactosaminyltransferase 3 |
| B4GALT5 | 1746.148557 | 3173.872958 | 1.817642001 | 0.000109705 | 0.004218164 | Beta-1,4-galactosyltransferase 5 |
| BACE2 | 984.5118861 | 475.9309338 | 0.483418169 | 6.92E-05 | 0.002918698 | Beta-secretase 2 |
| BACH2 | 140.5633676 | 315.1365015 | 2.241953268 | 2.42E-06 | 0.000193306 | Transcription regulator protein BACH2 |
| BAG5 | 260.4611614 | 426.5219187 | 1.637564374 | 0.002616703 | 0.046273552 | BAG family molecular chaperone regulator 5 |
| BAIAP2L2 | 73.50328099 | 211.202162 | 2.873370538 | 1.32E-06 | 0.000118871 | Brain-specific angiogenesis inhibitor 1-associated protein 2-like protein 2 |
| BBC3 | 479.3534259 | 1043.51421 | 2.176920313 | 3.31E-06 | 0.000249492 | Bcl-2-binding component 3 |
| BBS9 | 775.710776 | 376.8003545 | 0.485748511 | 9.33E-06 | 0.000589785 | Protein PTHB1 |
| BCAS1 | 11084.84388 | 3810.87696 | 0.343791667 | 2.64E-07 | 3.05E-05 | Breast carcinoma-amplified sequence 1 |
| BCKDK | 2355.023485 | 1429.768758 | 0.607114437 | 0.001066415 | 0.02372811 | [3-methyl-2-oxobutanoate dehydrogenase [lipoamide]] kinase, mitochondrial |
| BCL2 | 111.1415349 | 377.0169874 | 3.392224047 | 1.88E-06 | 0.000157978 | B-cell CLL/lymphoma 2, isoform CRA_b |
| BCL3 | 256.0213582 | 454.1375227 | 1.773826707 | 0.000296727 | 0.008996087 | B-cell lymphoma 3 protein |
| BCL6 | 1479.253635 | 2519.641869 | 1.703319707 | 0.000295303 | 0.008972928 | B-cell lymphoma 6 protein |
| BCOR | 381.0940401 | 631.858875 | 1.658013006 | 0.001235162 | 0.026547825 | BCL-6 corepressor |
| BCR | 1722.953316 | 517.1467476 | 0.300151341 | 1.12E-07 | 1.49E-05 | Breakpoint cluster region protein |
| BDH2 | 704.692093 | 408.9000676 | 0.58025352 | 0.000461841 | 0.01283868 | 3-hydroxybutyrate dehydrogenase type 2 |
| BEST1 | 99.46302774 | 179.8844717 | 1.808556161 | 0.001201542 | 0.025991081 | Bestrophin-1 |
| BHLHE40 | 1693.253951 | 4040.941624 | 2.386494727 | 7.81E-06 | 0.000513192 | Class E basic helix-loop-helix protein 40 |
| BICD2 | 2984.350377 | 1810.271994 | 0.606588291 | 0.000932578 | 0.021551843 | Protein bicaudal D homolog 2 |
| BID | 1004.168421 | 614.7327578 | 0.61218093 | 0.001028865 | 0.023118758 | BH3-interacting domain death agonist |
| BMF | 349.5805963 | 865.3878143 | 2.475502998 | 4.27E-08 | 6.70E-06 | Bcl-2-modifying factor |
| BMP2 | 46.35847883 | 102.2900777 | 2.206502032 | 0.000603167 | 0.015674275 | Bone morphogenetic protein 2 |

| Gene name | Controls normalized mean counts | Simvastatin normalized mean counts | foldChange | pval | padj | Protein name |
|---|---|---|---|---|---|---|
| BOLA2B | 406.6075336 | 239.855007 | 0.58989317 | 0.001301126 | 0.027496984 | |
| BOP1 | 1419.498077 | 858.4622709 | 0.604764659 | 0.001035224 | 0.023200051 | Ribosome biogenesis protein BOP1 |
| BRSK1 | 227.915843 | 84.24640711 | 0.369638223 | 4.21E-08 | 6.65E-06 | Serine/threonine-protein kinase BRSK1 |
| BTBD16 | 121.5326562 | 29.82580377 | 0.245413905 | 1.15E-05 | 0.000690149 | |
| BTBD6 | 635.3769388 | 2063.475417 | 3.247639772 | 3.03E-14 | 2.19E-11 | BTB/POZ domain-containing protein 6 |
| BVES-AS1 | 147.0281654 | 254.3491303 | 1.729934734 | 0.001697407 | 0.03329101 | |
| BYSL | 949.6010544 | 521.2007981 | 0.54886291 | 0.000167359 | 0.005842717 | Bystin |
| C10orf10 | 74.62254498 | 428.6322821 | 5.744005142 | 4.88E-09 | 1.03E-06 | |
| C10orf2 | 381.3981145 | 218.2867586 | 0.572333083 | 0.000845016 | 0.020019103 | |
| C10orf32 | 787.0631335 | 460.6862298 | 0.585323096 | 0.000468011 | 0.012940413 | |
| C10orf71 | 589.7708573 | 2015.983105 | 3.418248087 | 1.10E-09 | 2.70E-07 | |
| C10orf71-AS1 | 7.652637071 | 32.61361685 | 4.261748799 | 0.000165195 | 0.005812854 | |
| C11orf35 | 14.96276763 | 54.25788502 | 3.62619312 | 9.56E-06 | 0.000599326 | |
| C11orf52 | 39.98888669 | 95.24271753 | 2.381729661 | 0.000282582 | 0.008668626 | Uncharacterized protein C11orf52 |
| C11orf74 | 577.05915 | 360.3325327 | 0.624429112 | 0.002755182 | 0.048120542 | |
| C11orf96 | 5.219158067 | 22.48178019 | 4.307549207 | 0.001547904 | 0.031387935 | |
| C12orf39 | 38.96026888 | 83.72228134 | 2.148914362 | 0.000697753 | 0.017433175 | Spexin |
| C12orf5 | 442.3504326 | 258.9374402 | 0.585367214 | 0.000889276 | 0.020853427 | |
| C14orf1 | 205.5251562 | 475.3701752 | 2.312953722 | 1.72E-06 | 0.000147767 | Probable ergosterol biosynthetic protein 28 |
| C16orf11 | 2.433401491 | 38.27431239 | 15.72872891 | 8.06E-09 | 1.54E-06 | |
| C16orf59 | 60.45750139 | 28.6843656 | 0.474455029 | 0.002143428 | 0.039592326 | |
| C1orf106 | 26.43247807 | 1.199953809 | 0.045396947 | 8.21E-09 | 1.56E-06 | |
| C1orf127 | 8.619023052 | 57.89107195 | 6.716662851 | 5.59E-08 | 8.45E-06 | |
| C1orf198 | 11848.50561 | 5995.178536 | 0.505986049 | 6.02E-06 | 0.000414115 | Uncharacterized protein C1orf198 |
| C1orf21 | 255.3990775 | 768.2375884 | 3.007988893 | 2.02E-11 | 7.57E-09 | |
| C1orf213 | 57.75386002 | 181.3878054 | 3.140704455 | 0.000706899 | 0.017605118 | |
| C1orf226 | 76.49945828 | 29.55848746 | 0.386388193 | 0.000118291 | 0.00445466 | |
| C1orf95 | 603.470001 | 85.3822802 | 0.141485542 | 7.96E-16 | 6.88E-13 | Uncharacterized membrane protein C1orf95 |
| C1QBP | 2124.296427 | 1223.967773 | 0.576175602 | 0.000276684 | 0.008538214 | Complement component 1, q subcomponent binding protein, isoform CRA_a |
| C1QL4 | 600.7841547 | 125.0711323 | 0.208179812 | 0.000253949 | 0.00800339 | Complement C1q-like protein 4 |
| C1QTNF5 | 217.8801646 | 105.1539861 | 0.482623034 | 0.001008211 | 0.022758652 | Complement C1q tumor necrosis factor-related protein 5 |
| C1QTNF6 | 132.5747225 | 262.9644185 | 1.983518528 | 0.000524808 | 0.014137253 | Complement C1q tumor necrosis factor-related protein 6 |
| C1QTNF9B-AS1 | 28.24728541 | 66.22703464 | 2.344545101 | 0.000659858 | 0.016633338 | |
| C20orf166 | 2454.084207 | 4610.457418 | 1.878687538 | 0.001218138 | 0.026331751 | |
| C20orf166-AS1 | 3232.527609 | 1616.409185 | 0.500044974 | 0.000345858 | 0.010168492 | |
| C20orf27 | 617.7075994 | 347.698275 | 0.562884891 | 0.000264383 | 0.008223902 | UPF0687 protein C20orf27 |
| C2CD3 | 3042.237621 | 1230.833434 | 0.404581623 | 4.67E-09 | 9.94E-07 | C2 domain-containing protein 3 |
| C2CD4A | 1.037118087 | 13.33366578 | 12.85645863 | 0.00023206 | 0.007441718 | C2 calcium-dependent domain-containing protein 4A |
| C2orf82 | 85.7347033 | 156.404388 | 1.824283306 | 0.001599939 | 0.032135878 | Uncharacterized protein C2orf82 |
| C3orf58 | 1072.839457 | 442.3936668 | 0.412357752 | 3.17E-06 | 0.000240218 | Deleted in autism protein 1 |

| Gene name | Controls normalized mean counts | Simvastatin normalized mean counts | foldChange | pval | padj | Protein name |
|---|---|---|---|---|---|---|
| C6orf1 | 196.693483 | 331.7527953 | 1.686648639 | 0.00168361 | 0.033103901 | Uncharacterized protein C6orf1 |
| C6orf222 | 18.59563002 | 1.455897473 | 0.078292452 | 6.30E-06 | 0.000429769 | |
| C7 | 21.7735007 | 106.9366923 | 4.911322885 | 2.16E-06 | 0.000176277 | Complement component C7 |
| C7orf41 | 9860.281145 | 5712.354176 | 0.579329746 | 0.000858652 | 0.020249604 | |
| C9orf96 | 28.06523791 | 66.2212805 | 2.359548161 | 0.000555681 | 0.014773515 | cDNA FLJ60521, highly similar to Protein kinase-like protein C9orf96 |
| CACHD1 | 77.77953034 | 33.6821398 | 0.433046325 | 0.001434535 | 0.02951234 | VWFA and cache domain-containing protein 1 |
| CACNA1S | 6900.603146 | 3540.523056 | 0.513074434 | 0.000341017 | 0.010045161 | Voltage-dependent L-type calcium channel subunit alpha-1S |
| CACNG4 | 61.08876485 | 16.35592732 | 0.267740351 | 0.002468387 | 0.044254554 | Voltage-dependent calcium channel gamma-4 subunit |
| CADM4 | 268.8680003 | 104.3975682 | 0.388285583 | 2.46E-05 | 0.001261659 | Cell adhesion molecule 4 |
| CALD1 | 18673.71863 | 7644.444352 | 0.409369152 | 1.60E-07 | 2.01E-05 | Caldesmon |
| CALM2 | 2946.523578 | 1896.845137 | 0.643756986 | 0.002488577 | 0.044445622 | Calmodulin |
| CAMK2D | 4046.544912 | 7575.571908 | 1.872108693 | 1.99E-05 | 0.001070183 | Calcium/calmodulin-dependent protein kinase (CaM kinase) II delta, isoform CRA_e |
| CAMKK1 | 134.3475335 | 289.9966239 | 2.158555624 | 1.66E-05 | 0.000936421 | Calcium/calmodulin-dependent protein kinase kinase 1 |
| CAPN3 | 337.680559 | 940.2378191 | 2.784400209 | 1.72E-06 | 0.000147767 | Calpain-3 |
| CAPN9 | 122.4747485 | 53.07662901 | 0.433367936 | 4.36E-05 | 0.002026544 | Calpain-9 |
| CAPS | 28.20870851 | 81.48490668 | 2.888643648 | 8.60E-05 | 0.003467861 | Calcyphosine, isoform CRA_b |
| CARHSP1 | 628.162202 | 1285.558751 | 2.046539487 | 3.08E-06 | 0.00023721 | Calcium-regulated heat stable protein 1 |
| CASC10 | 29.81002904 | 7.869126261 | 0.2639758 | 6.47E-05 | 0.002770285 | |
| CASC7 | 1362.547799 | 2226.670987 | 1.634196605 | 0.000920429 | 0.02135038 | |
| CASC8 | 204.0118134 | 363.4118108 | 1.78132729 | 0.002845329 | 0.049362415 | |
| CASS4 | 2050.343935 | 418.7520539 | 0.204235029 | 6.09E-09 | 1.21E-06 | Cas scaffolding protein family member 4 |
| CAST | 3076.812912 | 5134.305568 | 1.668709055 | 0.000500972 | 0.013705582 | cDNA FLJ56102, highly similar to Homo sapiens calpastatin (CAST), transcript variant 8, mRNA |
| CAV1 | 6404.573722 | 3638.982468 | 0.568184961 | 9.23E-05 | 0.003688584 | Caveolin |
| CBLB | 814.0957309 | 1459.367582 | 1.792624045 | 0.000121355 | 0.004540248 | Cas-Br-M (Murine) ecotropic retroviral transforming sequence b, isoform CRA_a |
| CBR3 | 27.30704792 | 77.44871907 | 2.836217203 | 0.000128814 | 0.004770124 | Carbonyl reductase [NADPH] 3 |
| CBWD5 | 438.8153261 | 264.7350976 | 0.603295012 | 0.001820846 | 0.034747991 | COBW domain-containing protein 5 |
| CBX7 | 268.2367571 | 662.4205181 | 2.469536708 | 9.32E-08 | 1.27E-05 | Chromobox protein homolog 7 |
| CCDC110 | 23.59728708 | 54.40662947 | 2.305630698 | 0.001772581 | 0.034037863 | Coiled-coil domain-containing protein 110 |
| CCDC127 | 589.9990429 | 1005.845009 | 1.704824815 | 0.000449117 | 0.012563167 | |
| CCDC28A | 120.6553567 | 212.0093479 | 1.757148243 | 0.001564684 | 0.031666281 | |
| CCDC3 | 1618.859158 | 3154.330108 | 1.948489522 | 0.002643154 | 0.046608812 | Coiled-coil domain-containing protein 3 |
| CCDC39 | 1991.096456 | 3691.518583 | 1.854012935 | 0.000101494 | 0.003966184 | Coiled-coil domain-containing protein 39 |
| CCDC60 | 12.29010944 | 37.86160892 | 3.080656774 | 0.000783023 | 0.018895814 | |

| Gene name | Controls normalized mean counts | Simvastatin normalized mean counts | foldChange | pval | padj | Protein name |
|---|---|---|---|---|---|---|
| CCDC63 | 15.96708456 | 64.92448817 | 4.066145445 | 3.19E-06 | 0.000240991 | |
| CCDC69 | 93.68365575 | 403.5140778 | 4.307198246 | 2.41E-10 | 7.22E-08 | |
| CCDC74A | 40.18439588 | 98.63349353 | 2.454522243 | 0.000295087 | 0.008972928 | |
| CCDC85B | 848.4007804 | 523.4992572 | 0.617042404 | 0.001545875 | 0.031367251 | Coiled-coil domain-containing protein 85B |
| CCNA1 | 75.2622511 | 18.66026783 | 0.247936616 | 0.000133559 | 0.004904931 | Cyclin-A1 |
| CCNG2 | 1599.116058 | 3262.814477 | 2.040386288 | 1.39E-06 | 0.000124627 | cDNA FLJ56533, highly similar to Cyclin-G2 |
| CCR7 | 127.3041756 | 515.9082118 | 4.052563157 | 7.12E-07 | 7.04E-05 | C-C chemokine receptor type 7 |
| CCRN4L | 55.33425732 | 23.15504942 | 0.418457761 | 0.000579939 | 0.015249011 | Nocturnin |
| CCT2 | 7767.367989 | 4146.765777 | 0.533870133 | 4.63E-05 | 0.002117606 | T-complex protein 1 subunit beta |
| CCT3 | 6647.504714 | 3758.045485 | 0.565331752 | 0.000153602 | 0.005498197 | cDNA, FLJ78822, highly similar to T-complex protein 1 subunit gamma |
| CD109 | 1271.737073 | 2265.10992 | 1.781114955 | 0.000196286 | 0.006561974 | CD109 antigen |
| CD180 | 21.39204624 | 3.845424022 | 0.179759523 | 0.000491991 | 0.013483592 | CD180 antigen |
| CD320 | 305.804742 | 172.7947049 | 0.565049135 | 0.000696088 | 0.017405559 | CD320 antigen |
| CD34 | 779.7130639 | 168.113486 | 0.215609426 | 2.15E-15 | 1.71E-12 | cDNA FLJ53734, highly similar to Hematopoietic progenitor cell antigen CD34 |
| CD5 | 3.709415325 | 19.51473278 | 5.260864873 | 0.00063512 | 0.016220068 | T-cell surface glycoprotein CD5 |
| CD7 | 11.6621326 | 80.40367906 | 6.89442333 | 1.68E-10 | 5.27E-08 | T-cell antigen CD7 |
| CD79B | 28.25334665 | 89.70532566 | 3.175033625 | 1.57E-06 | 0.000137322 | B-cell antigen receptor complex-associated protein beta chain |
| CDC42EP1 | 3489.153563 | 2121.803694 | 0.608114162 | 0.001590273 | 0.032017489 | Cdc42 effector protein 1 |
| CDC42EP4 | 80.70776885 | 221.2152264 | 2.740940923 | 1.15E-06 | 0.000107166 | Cdc42 effector protein 4 |
| CDC73 | 3779.92605 | 2067.121185 | 0.546868155 | 7.58E-05 | 0.003138679 | Parafibromin |
| CDH1 | 88.36627542 | 15.45284106 | 0.174872608 | 0.001989169 | 0.037117624 | Cadherin-1 |
| CDH11 | 979.981363 | 474.8421363 | 0.484542007 | 3.68E-06 | 0.000273009 | Cadherin-11 |
| CDH15 | 16094.76946 | 9312.645477 | 0.578613164 | 0.002602984 | 0.046109578 | Cadherin-15 |
| CDH8 | 442.2641452 | 135.5662463 | 0.306527779 | 0.000369883 | 0.010762937 | Cadherin-8 |
| CDHR3 | 18.06632431 | 61.08293657 | 3.381038419 | 8.67E-06 | 0.000557269 | Cadherin-related family member 3 |
| CDHR4 | 0.242778076 | 13.24407065 | 54.55216895 | 2.25E-05 | 0.001179057 | Cadherin-related family member 4 |
| CDKN2D | 247.9193353 | 427.7083436 | 1.725191555 | 0.000772444 | 0.018722267 | Cyclin-dependent kinase 4 inhibitor D |
| CDON | 1110.278921 | 518.234549 | 0.466760685 | 0.000122391 | 0.004570345 | Cell adhesion molecule-related/down-regulated by oncogenes |
| CEBPG | 675.4123356 | 1249.434094 | 1.849883439 | 4.63E-05 | 0.002117606 | CCAAT/enhancer-binding protein gamma |
| CECR6 | 56.49155204 | 106.8363323 | 1.89119131 | 0.002672789 | 0.046921569 | |
| CELA3A | 2.098393693 | 14.68695994 | 6.999144147 | 0.001705914 | 0.033415709 | Chymotrypsin-like elastase family member 3A |
| CELSR1 | 374.6543024 | 102.7106179 | 0.274147707 | 1.25E-07 | 1.64E-05 | Cadherin EGF LAG seven-pass G-type receptor 1 |
| CENPN | 338.1099856 | 152.0902422 | 0.449824757 | 8.79E-06 | 0.000563909 | Centromere protein N |
| CEP78 | 928.3401244 | 351.0725113 | 0.378172291 | 3.10E-06 | 0.000237672 | Centrosomal protein of 78 kDa |
| CFH | 1033.92844 | 630.5018293 | 0.609811864 | 0.001470297 | 0.030111653 | Complement factor H |
| CFL1 | 14617.12832 | 8736.139967 | 0.597664587 | 0.000502391 | 0.013720262 | Cofilin-1 |
| CFLAR | 2081.763474 | 3232.719234 | 1.552875374 | 0.002869554 | 0.049589078 | CASP8 and FADD-like apoptosis regulator |
| CHAC2 | 92.55100585 | 33.47582649 | 0.361701379 | 0.000772822 | 0.018722267 | Cation transport regulator-like protein 2 |

| Gene name | **Controls** normalized mean counts | **Simvastatin** normalized mean counts | **foldChange** | **pval** | **padj** | **Protein name** |
|---|---|---|---|---|---|---|
| CHAMP1 | 580.7106774 | 351.1148161 | 0.604629516 | 0.001674274 | 0.033045663 | Chromosome alignment-maintaining phosphoprotein 1 |
| CHCHD10 | 732.9708071 | 453.4080185 | 0.618589464 | 0.002128075 | 0.039378875 | Coiled-coil-helix-coiled-coil-helix domain-containing protein 10, mitochondrial |
| CHCHD6 | 330.4987931 | 177.9607124 | 0.538461005 | 0.000182646 | 0.006245683 | Coiled-coil-helix-coiled-coil-helix domain-containing protein 6, mitochondrial |
| CHD3 | 7272.950015 | 3105.735319 | 0.427025528 | 8.61E-06 | 0.00055541 | Chromodomain-helicase-DNA-binding protein 3 |
| CHKB-CPT1B | 31.46034741 | 99.46083603 | 3.161466552 | 0.001016311 | 0.022907447 | Protein CHKB-CPT1B |
| CHST10 | 341.5843841 | 194.5184094 | 0.569459315 | 0.002236759 | 0.040975626 | Carbohydrate sulfotransferase 10 |
| CHST2 | 292.475522 | 88.99032584 | 0.304265893 | 2.44E-10 | 7.24E-08 | Carbohydrate sulfotransferase 2 |
| CILP | 203.0078957 | 19.26821334 | 0.094913615 | 4.87E-14 | 3.36E-11 | Cartilage intermediate layer protein 1 |
| CLDN5 | 77.5449287 | 419.682612 | 5.412121967 | 2.96E-05 | 0.001469693 | Claudin-5 |
| CLIC1 | 3119.61204 | 5632.248829 | 1.805432457 | 7.17E-05 | 0.00301075 | Chloride intracellular channel protein 1 |
| CLIC5 | 23.65036342 | 67.2424229 | 2.843187722 | 8.45E-05 | 0.003414061 | Chloride intracellular channel protein 5 |
| CLIP1 | 2355.32696 | 3846.375727 | 1.633053836 | 0.000911392 | 0.021219885 | CAP-Gly domain-containing linker protein 1 |
| CLIP3 | 1516.545612 | 538.8313456 | 0.355301774 | 0.000162019 | 0.005726991 | CAP-Gly domain-containing linker protein 3 |
| CLU | 393.5787564 | 1079.451872 | 2.742657865 | 2.35E-06 | 0.000188319 | Clusterin beta chain |
| CMC2 | 630.3832017 | 367.7135378 | 0.583317476 | 0.000607543 | 0.015714443 | COX assembly mitochondrial protein 2 homolog |
| CMYA5 | 805.4377326 | 1704.096388 | 2.115739454 | 0.001272356 | 0.027071071 | Cardiomyopathy-associated protein 5 |
| CNN2 | 5569.35916 | 1669.382606 | 0.29974411 | 0.000411789 | 0.011773071 | Calponin-2 |
| CNNM4 | 261.9376656 | 546.9970605 | 2.088271877 | 4.15E-06 | 0.000303135 | Metal transporter CNNM4 |
| CNTNAP3B | 15.32824094 | 1.182349725 | 0.077135382 | 0.001064096 | 0.023710443 | Contactin-associated protein-like 3B |
| COA4 | 2270.424789 | 1169.753592 | 0.515213539 | 1.35E-05 | 0.000789097 | Cytochrome c oxidase assembly factor 4 homolog, mitochondrial |
| COL14A1 | 592.6835435 | 229.2921022 | 0.386871046 | 1.11E-06 | 0.000104028 | Collagen alpha-1(XIV) chain |
| COL16A1 | 4129.282644 | 1396.096679 | 0.338096662 | 0.000157381 | 0.005601238 | Collagen alpha-1(XVI) chain |
| COL24A1 | 93.49519236 | 202.7625682 | 2.168695128 | 0.000211896 | 0.006974851 | Collagen alpha-1(XXIV) chain |
| COL26A1 | 28.14037618 | 4.288429859 | 0.152394191 | 7.89E-06 | 0.000513507 | Collagen alpha-1(XXVI) chain |
| COL4A2 | 3618.281369 | 2022.144867 | 0.558868883 | 0.001302098 | 0.027496984 | Canstatin |
| COL4A5 | 655.3150686 | 179.5770652 | 0.274031643 | 6.10E-11 | 2.06E-08 | Collagen alpha-5(IV) chain |
| COL4A6 | 43.72680193 | 16.21030106 | 0.370717737 | 0.000534561 | 0.014359292 | Collagen alpha-6(IV) chain |
| COL6A4P1 | 16.27847036 | 2.42531271 | 0.148988981 | 3.13E-05 | 0.001536688 | |
| COL8A2 | 115.0507359 | 36.58328929 | 0.317975274 | 0.000260346 | 0.008130856 | Collagen type VIII alpha 2 |
| COMP | 302.6078011 | 33.52963691 | 0.110802289 | 4.34E-06 | 0.000314868 | Cartilage oligomeric matrix protein |
| CORO6 | 5619.717432 | 9242.563205 | 1.644666893 | 0.000899951 | 0.021032219 | Coronin-6 |
| CORO7 | 6654.203152 | 2793.826412 | 0.419858899 | 0.000583208 | 0.015283285 | Coronin |
| COTL1 | 1180.733925 | 378.7356214 | 0.320762886 | 5.98E-10 | 1.54E-07 | Coactosin-like protein |

| Gene name | **Controls** normalized mean counts | **Simvastatin** normalized mean counts | foldChange | pval | padj | Protein name |
|---|---|---|---|---|---|---|
| CPEB2 | 839.2004349 | 1632.986044 | 1.945883219 | 7.89E-06 | 0.000513507 | Cytoplasmic polyadenylation element-binding protein 2 |
| CPLX1 | 72.83120274 | 261.7753888 | 3.594275242 | 2.81E-09 | 6.37E-07 | Complexin-1 |
| CPNE8 | 126.2014286 | 309.5803005 | 2.453064944 | 9.61E-06 | 0.000601489 | Copine-8 |
| CPS1 | 331.5241424 | 194.9448223 | 0.588026021 | 0.00129562 | 0.02743469 | Carbamoyl-phosphate synthase [ammonia], mitochondrial |
| CPT1A | 211.539592 | 447.4280547 | 2.115103138 | 0.001144836 | 0.025042602 | Carnitine O-palmitoyltransferase 1, liver isoform |
| CPT1C | 91.62970142 | 31.34647072 | 0.342099453 | 7.09E-06 | 0.000473001 | Carnitine O-palmitoyltransferase 1, brain isoform |
| CRAT | 1165.939726 | 2065.017546 | 1.77111861 | 0.000105734 | 0.004106058 | Carnitine O-acetyltransferase |
| CREBRF | 399.4436682 | 759.4801821 | 1.901344902 | 0.000114869 | 0.004362786 | CREB3 regulatory factor |
| CREG2 | 92.27227991 | 302.8825353 | 3.28248674 | 5.02E-11 | 1.77E-08 | Protein CREG2 |
| CRYM | 22.45647866 | 183.394149 | 8.166647666 | 0.000127422 | 0.00472983 | Ketimine reductase mu-crystallin |
| CSF1 | 907.9596935 | 1942.621977 | 2.139546492 | 0.001039543 | 0.023275782 | Macrophage colony-stimulating factor 1 |
| CSPG4 | 393.8798244 | 2422.287206 | 6.14981285 | 3.89E-18 | 4.84E-15 | Chondroitin sulfate proteoglycan 4 |
| CSPG4P13 | 27.99316883 | 125.6258015 | 4.487730643 | 1.46E-05 | 0.000836824 | |
| CTA-221G9.11 | 15.2302967 | 88.83037737 | 5.832478454 | 9.28E-05 | 0.003699742 | |
| CTAGE5 | 287.9872622 | 560.2888993 | 1.945533615 | 4.51E-05 | 0.00207994 | cTAGE family member 5 |
| CTB-129O4.1 | 8.142859524 | 34.37480219 | 4.221465701 | 0.000137256 | 0.005017007 | |
| CTC-264K15.6 | 108.0342768 | 275.535932 | 2.550449174 | 1.54E-07 | 1.96E-05 | |
| CTC-325H20.4 | 293.3306451 | 107.1584898 | 0.365316381 | 1.32E-06 | 0.000118871 | |
| CTD-2001E22.1 | 84.81494747 | 3.753729825 | 0.044257881 | 5.89E-05 | 0.002586064 | |
| CTD-2044J15.2 | 58.10586252 | 116.2286332 | 2.000290989 | 0.000813713 | 0.019470273 | |
| CTD-2083E4.7 | 40.71170481 | 167.3095666 | 4.109618288 | 7.64E-05 | 0.003159103 | |
| CTD-2201E18.3 | 288.6943431 | 477.0599957 | 1.652474345 | 0.001604469 | 0.032178343 | |
| CTD-2228K2.7 | 841.0502763 | 2581.990525 | 3.069959785 | 7.45E-12 | 3.05E-09 | |
| CTD-2545M3.8 | 800.7699968 | 2185.806208 | 2.729630502 | 5.38E-06 | 0.000382217 | |
| CTD-2552K11.2 | 15.74295371 | 57.46385718 | 3.650131877 | 3.08E-06 | 0.00023721 | |
| CTD-2587H24.5 | 2.984524968 | 25.56226472 | 8.564935792 | 5.28E-06 | 0.00037684 | |
| CTD-2636A23.2 | 82.66326051 | 203.0636929 | 2.456516857 | 0.001912316 | 0.036138822 | |
| CTD-2651B20.1 | 78.34404798 | 227.4223747 | 2.902867296 | 1.46E-07 | 1.87E-05 | |
| CTD-3184A7.4 | 64.79132964 | 122.0050629 | 1.883046135 | 0.001390059 | 0.028864602 | |
| CTDSP1 | 687.3962744 | 1114.013308 | 1.620627503 | 0.001232032 | 0.026506209 | Carboxy-terminal domain RNA polymerase II polypeptide A small phosphatase 1 |
| CTGF | 3668.441027 | 966.6150932 | 0.263494789 | 3.35E-13 | 1.89E-10 | Connective tissue growth factor |
| CTNND1 | 7126.125978 | 4194.388113 | 0.588593034 | 0.000395252 | 0.011385346 | Catenin delta-1 |
| CTPS1 | 919.6497972 | 461.5767172 | 0.501904876 | 1.83E-05 | 0.000992178 | CTP synthase |
| CTXN1 | 280.3847706 | 149.4914635 | 0.533165418 | 0.000310295 | 0.009280806 | Cortexin-1 |
| CXCL2 | 7.039813168 | 25.88769542 | 3.677327054 | 0.000561864 | 0.014858114 | C-X-C motif chemokine 2 |
| CYFIP2 | 444.4635533 | 132.3707593 | 0.297821404 | 3.15E-06 | 0.000239054 | Cytoplasmic FMR1-interacting protein 2 |
| CYP19A1 | 2.724933064 | 33.3262142 | 12.23010379 | 5.80E-06 | 0.00040205 | Aromatase |
| CYP1B1 | 1118.111313 | 8312.551105 | 7.434457563 | 2.60E-06 | 0.000204918 | Cytochrome P450 1B1 |
| CYP27A1 | 230.8223957 | 492.9432516 | 2.135595422 | 0.000513539 | 0.013951231 | Sterol 26-hydroxylase, mitochondrial |
| CYP27C1 | 34.15067508 | 8.456602374 | 0.247626214 | 0.000437415 | 0.012368724 | Cytochrome P450 27C1 |
| CYP51A1 | 1586.891945 | 3361.102152 | 2.118040968 | 1.26E-06 | 0.000114514 | Lanosterol 14-alpha demethylase |
| CYR61 | 3596.144311 | 1365.94882 | 0.379837043 | 0.000702429 | 0.017521859 | Protein CYR61 |

| Gene name | Controls normalized mean counts | Simvastatin normalized mean counts | foldChange | pval | padj | Protein name |
|---|---|---|---|---|---|---|
| CYSLTR1 | 17.96942623 | 84.91064959 | 4.725284409 | 7.46E-05 | 0.003100194 | Cysteinyl leukotriene receptor 1 |
| CYTH1 | 799.8939522 | 1791.452351 | 2.239612321 | 7.34E-08 | 1.06E-05 | cDNA FLJ54415, highly similar to Cytohesin-1 |
| CYTIP | 43.67801896 | 213.2850761 | 4.883121562 | 6.82E-05 | 0.002883672 | Cytohesin-interacting protein |
| DBN1 | 14191.39111 | 7658.150997 | 0.539633566 | 4.59E-05 | 0.002110623 | Drebrin |
| DBNDD2 | 52.23375615 | 288.7869085 | 5.528740985 | 3.11E-07 | 3.51E-05 | Dysbindin domain-containing protein 2 |
| DCHS1 | 166.5631416 | 385.9696762 | 2.317257423 | 6.07E-06 | 0.00041713 | Protocadherin-16 |
| DCLK3 | 82.99306071 | 10.47464691 | 0.126211117 | 0.00232592 | 0.042359517 | Serine/threonine-protein kinase DCLK3 |
| DCX | 133.2121096 | 17.48441957 | 0.131252479 | 1.72E-05 | 0.00095754 | Neuronal migration protein doublecortin |
| DDAH1 | 2762.609593 | 939.4556577 | 0.340060955 | 5.17E-05 | 0.002338639 | N(G),N(G)-dimethylarginine dimethylaminohydrolase 1 |
| DDN | 44.16480165 | 491.1732869 | 11.12137423 | 5.67E-20 | 9.29E-17 | Dendrin |
| DDOST | 4348.287104 | 2444.750933 | 0.562233099 | 0.00010552 | 0.004102868 | Dolichyl-diphosphooligosaccharide--protein glycosyltransferase 48 kDa subunit |
| DDX20 | 606.5012816 | 353.5535533 | 0.582939499 | 0.000707693 | 0.017605118 | cDNA FLJ50269, highly similar to Probable ATP-dependent RNA helicase DDX20 (EC 3.6.1.-) |
| DDX5 | 13454.72185 | 8485.230789 | 0.630650777 | 0.00172522 | 0.033540427 | cDNA FLJ59339, highly similar to Probable ATP-dependent RNA helicase DDX5 (EC 3.6.1.-) |
| DDX55 | 424.1262383 | 240.6987571 | 0.56751678 | 0.000550632 | 0.014651798 | ATP-dependent RNA helicase DDX55 |
| DEC1 | 1.027192005 | 18.3511569 | 17.86536189 | 4.79E-06 | 0.000344833 | Deleted in esophageal cancer 1 |
| DECR1 | 1902.36231 | 1135.644011 | 0.596965155 | 0.000646064 | 0.016405316 | cDNA FLJ50204, highly similar to 2,4-dienoyl-CoA reductase, mitochondrial (EC 1.3.1.34) |
| DEFB103A | 671.5425838 | 55.04982761 | 0.081975185 | 9.70E-05 | 0.003824035 | Beta-defensin 103 |
| DEFB103B | 957.0152157 | 53.29099556 | 0.055684585 | 9.83E-07 | 9.35E-05 | |
| DEK | 6329.225533 | 2965.438544 | 0.468531028 | 4.40E-07 | 4.75E-05 | cDNA FLJ53031, highly similar to Protein DEK |
| DENND2C | 279.5350766 | 600.5133447 | 2.148257571 | 2.00E-06 | 0.00016465 | DENN domain-containing protein 2C |
| DENND3 | 87.00594368 | 166.4949094 | 1.913603857 | 0.000829516 | 0.019727004 | DENN domain-containing protein 3 |
| DENND4C | 549.6261459 | 986.2303298 | 1.794365747 | 0.000133023 | 0.004891021 | DENN domain-containing protein 4C |
| DES | 108219.4395 | 231124.4729 | 2.135701996 | 2.98E-07 | 3.40E-05 | Desmin |
| DESI2 | 2942.811771 | 1760.012125 | 0.598071593 | 0.000608409 | 0.015716214 | Desumoylating isopeptidase 2 |
| DFNA5 | 568.8308379 | 1048.92618 | 1.844003718 | 0.000183774 | 0.006268065 | Non-syndromic hearing impairment protein 5 |
| DGKI | 208.8876438 | 467.3376081 | 2.237267842 | 3.30E-05 | 0.001609437 | Diacylglycerol kinase iota |
| DGKZ | 2493.946052 | 4028.584728 | 1.615345579 | 0.000978205 | 0.022226475 | Diacylglycerol kinase zeta |
| DHCR24 | 2540.165571 | 6567.214492 | 2.585348989 | 1.38E-07 | 1.79E-05 | cDNA FLJ53870, highly similar to 24-dehydrocholesterol reductase (EC1.3.1.-) |
| DHCR7 | 1753.126494 | 3281.287292 | 1.871677431 | 4.72E-05 | 0.002151101 | 7-dehydrocholesterol reductase |

| Gene name | **Controls** normalized mean counts | **Simvastatin** normalized mean counts | foldChange | pval | padj | Protein name |
|---|---|---|---|---|---|---|
| DHRS7C | 11.97363638 | 157.6458494 | 13.16607957 | 0.000194078 | 0.006532815 | Dehydrogenase/reductase SDR family member 7C |
| DHX29 | 1905.46895 | 1124.724851 | 0.590261443 | 0.000592516 | 0.015468619 | ATP-dependent RNA helicase DHX29 |
| DIAPH1 | 1041.543489 | 2466.086033 | 2.367722576 | 3.27E-08 | 5.40E-06 | cDNA FLJ61549, highly similar to Protein diaphanous homolog 1 |
| DICER1-AS1 | 50.2865322 | 112.505673 | 2.237292334 | 0.0001453 | 0.00527387 | |
| DIO2-AS1 | 8.820794805 | 47.85718487 | 5.425495766 | 2.21E-07 | 2.63E-05 | |
| DIRAS1 | 586.1788857 | 1099.35532 | 1.875460455 | 0.000529482 | 0.014247452 | GTP-binding protein Di-Ras1 |
| DIRAS3 | 23.26356379 | 5.82151323 | 0.250241678 | 0.00028009 | 0.008612436 | GTP-binding protein Di-Ras3 |
| DIXDC1 | 582.4170424 | 1276.862664 | 2.192351135 | 2.57E-07 | 2.99E-05 | Dixin |
| DLC1 | 729.100399 | 385.6883862 | 0.528992148 | 4.23E-05 | 0.001985755 | Rho GTPase-activating protein 7 |
| DLGAP1-AS1 | 122.0559766 | 208.6957534 | 1.709836414 | 0.002437134 | 0.043820511 | |
| DLX3 | 22.66368682 | 86.19155579 | 3.803068604 | 1.89E-06 | 0.000158152 | Homeobox protein DLX-3 |
| DMBT1 | 841.9561845 | 366.5052118 | 0.435302001 | 7.82E-06 | 0.000513192 | Deleted in malignant brain tumors 1 protein |
| DNAAF3 | 8.406849728 | 27.8797655 | 3.316315434 | 0.001323991 | 0.027845882 | Dynein assembly factor 3, axonemal |
| DNAH7 | 109.4490661 | 35.1732179 | 0.321366085 | 2.85E-06 | 0.000223493 | Dynein heavy chain 7, axonemal |
| DNAJB4 | 3496.536563 | 2099.102097 | 0.600337522 | 0.001004888 | 0.022716172 | DnaJ homolog subfamily B member 4 |
| DNAJC12 | 113.9920657 | 48.00437062 | 0.42112028 | 2.60E-05 | 0.001315189 | |
| DNM1P35 | 6.542607968 | 38.58987056 | 5.898239777 | 3.83E-06 | 0.000280967 | |
| DPYSL2 | 603.066268 | 1687.250644 | 2.797786468 | 3.24E-05 | 0.001585723 | Dihydropyrimidinase-related protein 2 |
| DPYSL5 | 7767.416629 | 2332.68459 | 0.300316656 | 0.001494516 | 0.030484201 | Dihydropyrimidinase-related protein 5 |
| DST | 17084.80468 | 27936.72119 | 1.635179431 | 0.000747248 | 0.018344996 | Dystonin |
| DTNA | 2784.578366 | 1725.45019 | 0.619645046 | 0.001853473 | 0.035240917 | cDNA FLJ77455, highly similar to Homo sapiens dystrobrevin, alpha (DTNA), transcript variant 5, mRNA |
| DUOX1 | 10.50990972 | 35.44508676 | 3.372539603 | 0.00029897 | 0.009046457 | Dual oxidase 1 |
| DUSP14 | 1384.582087 | 630.1295519 | 0.45510451 | 2.92E-07 | 3.35E-05 | Dual specificity protein phosphatase 14 |
| DUSP15 | 10.01788885 | 42.80497736 | 4.272854093 | 5.10E-05 | 0.002310511 | Dual specificity protein phosphatase 15 |
| DUSP16 | 253.3592983 | 430.8086471 | 1.700386171 | 0.001005599 | 0.022716172 | Dual-specificity protein phosphatase 16 |
| DUSP5P1 | 11.96441676 | 35.83146671 | 2.994836057 | 0.002336029 | 0.042444228 | |
| DUSP6 | 767.3771576 | 314.3217092 | 0.409605246 | 0.000132084 | 0.004868005 | Dual-specificity protein phosphatase 6 |
| DUSP8 | 112.3881185 | 407.3876942 | 3.624828849 | 1.04E-07 | 1.39E-05 | Dual specificity protein phosphatase 8 |
| DVL1 | 1513.906211 | 2618.062605 | 1.729342668 | 0.000195044 | 0.006537749 | Segment polarity protein dishevelled homolog DVL-1 |
| DYNLL1 | 5415.849481 | 2908.437834 | 0.537023387 | 3.57E-05 | 0.00172 | Dynein light chain 1, cytoplasmic |
| DYSF | 6246.88902 | 10503.99001 | 1.681475367 | 0.000318343 | 0.009494117 | Dysferlin |
| DZIP1 | 465.7111566 | 285.4971306 | 0.613034768 | 0.002450763 | 0.044014687 | Zinc finger protein DZIP1 |
| EARS2 | 799.9542183 | 454.3710366 | 0.567996301 | 0.000262024 | 0.008167779 | Probable glutamate--tRNA ligase, mitochondrial |
| EBNA1BP2 | 2647.474894 | 1199.946318 | 0.453241812 | 3.17E-07 | 3.55E-05 | EBNA1 binding protein 2, isoform CRA_d |

| Gene name | Controls normalized mean counts | Simvastatin normalized mean counts | foldChange | pval | padj | Protein name |
|---|---|---|---|---|---|---|
| EBP | 479.1687988 | 943.5735433 | 1.969188198 | 0.000224572 | 0.007253947 | 3-beta-hydroxysteroid-Delta(8),Delta(7)-isomerase |
| ECHDC3 | 279.1343979 | 140.4668184 | 0.50322289 | 8.16E-05 | 0.003330814 | Enoyl-CoA hydratase domain-containing protein 3, mitochondrial |
| ECM1 | 1827.975652 | 826.884913 | 0.452350069 | 0.000559199 | 0.014829021 | Extracellular matrix protein 1 |
| ECM2 | 233.9910159 | 95.11912299 | 0.406507586 | 0.00063729 | 0.01626213 | Extracellular matrix protein 2 |
| EDNRB | 79.67488321 | 235.8129784 | 2.959690292 | 0.001959564 | 0.03670834 | Endothelin B receptor |
| EEF1A2 | 2299.138446 | 4542.800405 | 1.97587075 | 0.000198625 | 0.006600877 | Elongation factor 1-alpha 2 |
| EEF1E1 | 467.3313379 | 285.0501402 | 0.609952976 | 0.001713899 | 0.033466747 | Eukaryotic translation elongation factor 1 epsilon-1 |
| EEF2K | 518.5420927 | 890.6142981 | 1.717535202 | 0.000448535 | 0.012560013 | Eukaryotic elongation factor 2 kinase |
| EEPD1 | 166.9006845 | 89.73719411 | 0.537668221 | 0.000965685 | 0.022006301 | Endonuclease/exonuclease /phosphatase family domain-containing protein 1 |
| EFCAB4B | 207.2458444 | 454.7161773 | 2.194090688 | 1.79E-06 | 0.00015255 | EF-hand calcium-binding domain-containing protein 4B |
| EFCC1 | 76.17167915 | 21.50316751 | 0.28229872 | 0.002873593 | 0.049631303 | EF-hand and coiled-coil domain-containing protein 1 |
| EFHD1 | 796.5373001 | 396.2368676 | 0.497449231 | 0.000313771 | 0.009366766 | EF-hand domain-containing protein D1 |
| EFHD2 | 1212.26877 | 2080.129466 | 1.715897924 | 0.000215474 | 0.007058261 | EF-hand domain-containing protein D2 |
| EFNB2 | 375.7792582 | 158.0284899 | 0.420535425 | 9.79E-05 | 0.003856169 | Ephrin-B2 |
| EFR3B | 100.0689872 | 53.09271921 | 0.530561173 | 0.002025799 | 0.037778486 | |
| EGFLAM | 183.7185096 | 71.34259689 | 0.388325581 | 0.000856403 | 0.020211881 | Pikachurin |
| EGLN3 | 8983.300733 | 4657.525108 | 0.518464788 | 0.000608823 | 0.015716214 | Egl nine homolog 3 |
| EHD1 | 4228.257627 | 2367.845199 | 0.560004949 | 0.000622732 | 0.015995614 | EH domain-containing protein 1 |
| EID1 | 1824.901289 | 602.3232231 | 0.330057974 | 2.60E-07 | 3.02E-05 | EP300-interacting inhibitor of differentiation 1 |
| EIF2AK3 | 265.7073797 | 550.6179303 | 2.072271876 | 1.74E-05 | 0.000960417 | Eukaryotic translation initiation factor 2-alpha kinase 3 |
| EIF4A1 | 9914.524192 | 4609.532877 | 0.464927291 | 3.47E-07 | 3.84E-05 | Eukaryotic initiation factor 4A-I |
| EIF5A | 10254.10635 | 6156.196544 | 0.600364024 | 0.000538673 | 0.014421372 | Eukaryotic translation initiation factor 5A-1 |
| ELF1 | 762.5073238 | 1221.5526 | 1.60202081 | 0.001683072 | 0.033103901 | ETS-related transcription factor Elf-1 |
| ELF4 | 343.6616231 | 640.1153947 | 1.862632752 | 9.50E-05 | 0.003768897 | ETS-related transcription factor Elf-4 |
| ELOVL4 | 110.0914044 | 56.88594884 | 0.516715625 | 0.001452175 | 0.029816084 | Elongation of very long chain fatty acids protein 4 |
| EMC1 | 2204.275358 | 1379.474142 | 0.625817522 | 0.001644755 | 0.032795992 | ER membrane protein complex subunit 1 |
| EMILIN1 | 809.8285264 | 426.6139609 | 0.526795423 | 0.000331015 | 0.009815591 | EMILIN-1 |
| EMILIN2 | 15.85509768 | 65.16143797 | 4.109809936 | 1.01E-05 | 0.00061844 | EMILIN-2 |
| EMP3 | 1406.210814 | 6937.608805 | 4.933548182 | 6.55E-25 | 1.70E-21 | Epithelial membrane protein 3 |
| EN2 | 259.9814443 | 628.4468235 | 2.417275684 | 7.50E-08 | 1.08E-05 | Homeobox protein engrailed-2 |

| Gene name | Controls normalized mean counts | Simvastatin normalized mean counts | foldChange | pval | padj | Protein name |
|---|---|---|---|---|---|---|
| ENC1 | 764.0654952 | 190.8616686 | 0.249797524 | 7.51E-13 | 3.89E-10 | Ectoderm-neural cortex protein 1 |
| ENOX1 | 634.1359761 | 285.5176558 | 0.45024674 | 7.30E-06 | 0.000485042 | Ecto-NOX disulfide-thiol exchanger 1 |
| ENPP6 | 27.75457007 | 2.783331641 | 0.100283724 | 8.84E-07 | 8.51E-05 | Ectonucleotide pyrophosphatase/phosphodiesterase family member 6 soluble form |
| EPB41 | 80.82919466 | 192.5994479 | 2.382795582 | 2.43E-05 | 0.00125152 | Protein 4.1 |
| EPB41L1 | 597.0035585 | 1391.215525 | 2.330330372 | 0.000139513 | 0.005087548 | Band 4.1-like protein 1 |
| EPB41L2 | 1757.777099 | 951.63754 | 0.541386926 | 0.002666638 | 0.046863519 | Band 4.1-like protein 2 |
| EPB41L4B | 17.04924633 | 83.18027565 | 4.878824204 | 3.52E-05 | 0.001700779 | Band 4.1-like protein 4B |
| EPHA2 | 308.0488821 | 129.1556618 | 0.419270023 | 7.06E-06 | 0.000472605 | Ephrin type-A receptor 2 |
| EPHB2 | 111.8008772 | 36.80053218 | 0.32916139 | 0.001360635 | 0.028405318 | Ephrin type-B receptor 2 |
| EPHX1 | 785.5086343 | 1379.246369 | 1.755864047 | 0.000170937 | 0.005940983 | Epoxide hydrolase 1 |
| ERBB3 | 3832.027338 | 5976.129872 | 1.559521722 | 0.002819195 | 0.049042586 | Receptor tyrosine-protein kinase erbB-3 |
| ERC2 | 118.4742989 | 21.76073438 | 0.183674726 | 3.62E-08 | 5.89E-06 | ERC protein 2 |
| ERF | 1108.138163 | 630.3325388 | 0.568821253 | 0.00137859 | 0.028679451 | Ets2 repressor factor, isoform CRA_b |
| ERVK3-1 | 619.9422227 | 364.8966344 | 0.58859781 | 0.000741951 | 0.018256377 | |
| ESRP1 | 51.78085449 | 3.763003406 | 0.072671713 | 0.001058152 | 0.02361181 | Epithelial-splicing regulatory protein 1 |
| ESYT2 | 2651.66282 | 5162.942352 | 1.947058394 | 6.75E-06 | 0.000453727 | Extended synaptotagmin-2 |
| EVA1C | 23.46064214 | 94.54293057 | 4.029852636 | 1.46E-08 | 2.59E-06 | Chromosome 21 open reading frame 63, isoform CRA_c |
| EVC | 1155.22426 | 2054.673371 | 1.778592644 | 0.000180452 | 0.006204143 | Ellis-van Creveld syndrome protein |
| EXOC3L4 | 66.809767 | 28.04967559 | 0.419843937 | 0.001184668 | 0.025679647 | Exocyst complex component 3-like protein 4 |
| EXOG | 2969.030588 | 1332.137854 | 0.448677713 | 6.72E-05 | 0.00284589 | Nuclease EXOG, mitochondrial |
| EXPH5 | 92.95101237 | 32.0749026 | 0.345073193 | 0.000466901 | 0.012921181 | Exophilin-5 |
| EYS | 302.5505873 | 121.5176108 | 0.401643943 | 1.21E-05 | 0.000716899 | Protein eyes shut homolog |
| EZR | 1142.983903 | 4347.667515 | 3.803787177 | 9.70E-18 | 1.16E-14 | Ezrin |
| FADS1 | 1530.490151 | 3090.74646 | 2.019448774 | 1.88E-06 | 0.000157978 | Fatty acid desaturase 1 |
| FADS3 | 9199.6304 | 4959.67356 | 0.539116611 | 0.001254188 | 0.026849811 | Fatty acid desaturase 3 |
| FAM13A-AS1 | 62.94418308 | 156.95475 | 2.493554485 | 1.59E-05 | 0.000907078 | |
| FAM170B-AS1 | 6.52709408 | 25.23581491 | 3.866317016 | 0.001555394 | 0.03149875 | |
| FAM178B | 24.02555859 | 95.60906616 | 3.979473184 | 4.68E-07 | 5.04E-05 | |
| FAM198B | 4045.617824 | 1536.106179 | 0.379696315 | 6.99E-08 | 1.01E-05 | Protein FAM198B |
| FAM207A | 522.8553834 | 284.8941615 | 0.544881377 | 0.000408185 | 0.011702312 | |
| FAM20A | 94.89055841 | 32.43099695 | 0.341772643 | 5.22E-05 | 0.002358542 | Protein FAM20A |
| FAM20C | 605.1020943 | 1485.354096 | 2.454716502 | 1.53E-08 | 2.68E-06 | Extracellular serine/threonine protein kinase FAM20C |
| FAM216A | 529.7782937 | 185.9212823 | 0.350941676 | 2.42E-09 | 5.54E-07 | |
| FAM219A | 1099.747201 | 1724.694467 | 1.568264476 | 0.002508765 | 0.044720716 | |
| FAM230A | 41.17381983 | 4.193639877 | 0.101852097 | 2.92E-08 | 4.86E-06 | |
| FAM47E-STBD1 | 253.8234325 | 1037.343843 | 4.086871858 | 1.62E-17 | 1.82E-14 | Protein FAM47E-STBD1 |
| FAM53B | 470.0660081 | 1022.669113 | 2.175586185 | 3.55E-07 | 3.90E-05 | Protein FAM53B |
| FAM60A | 466.24663 | 200.9435126 | 0.430981158 | 4.24E-07 | 4.60E-05 | Protein FAM60A |
| FAM63A | 318.7881416 | 571.2729918 | 1.792014561 | 0.000250094 | 0.007900081 | Protein FAM63A |
| FAM63B | 350.9228796 | 581.0412671 | 1.655752021 | 0.002352563 | 0.042620157 | Protein FAM63B |
| FAM66A | 79.90837874 | 34.73501943 | 0.434685573 | 0.001017012 | 0.022907447 | |
| FAM66D | 229.7093532 | 65.0311545 | 0.283101901 | 0.001513956 | 0.030820093 | |
| FAM76A | 275.7221181 | 458.9026545 | 1.664366492 | 0.001633267 | 0.032629665 | |
| FAM84A | 232.2016423 | 52.60578721 | 0.226552176 | 1.27E-14 | 9.38E-12 | |
| FAM90A11P | 22.92818337 | 3.51492302 | 0.153301418 | 4.94E-06 | 0.000354005 | |

| Gene name | Controls normalized mean counts | Simvastatin normalized mean counts | foldChange | pval | padj | Protein name |
|---|---|---|---|---|---|---|
| FAM90A18P | 43.43188445 | 13.79944815 | 0.317726212 | 5.61E-05 | 0.00249356 | Putative protein FAM90A18P/FAM90A19P |
| FAM90A21P | 147.4932471 | 49.19684001 | 0.333553169 | 3.89E-07 | 4.23E-05 | |
| FAM90A3P | 489.9129707 | 144.1651143 | 0.294266784 | 2.25E-06 | 0.000182356 | |
| FAM98A | 2453.130445 | 1557.497006 | 0.634901829 | 0.002372416 | 0.042929835 | Protein FAM98A |
| FANK1 | 34.47110593 | 73.9957 | 2.146600696 | 0.001433714 | 0.02951234 | Fibronectin type III and ankyrin repeat domains 1, isoform CRA_d |
| FARSB | 1544.620782 | 967.6286029 | 0.626450592 | 0.001946024 | 0.036509661 | Phenylalanine--tRNA ligase beta subunit |
| FASTKD1 | 449.379501 | 244.2696729 | 0.543571018 | 0.000187308 | 0.006360689 | FAST kinase domain-containing protein 1 |
| FAXDC2 | 161.5066556 | 384.5220376 | 2.380843293 | 2.04E-07 | 2.46E-05 | cDNA FLJ57989 |
| FBLN1 | 388.0219403 | 899.5141401 | 2.318204325 | 0.001534382 | 0.031154372 | Fibulin-1 |
| FBXL19-AS1 | 298.6215905 | 168.7292465 | 0.56502695 | 0.000717955 | 0.017766677 | |
| FBXL8 | 105.0626545 | 191.1675206 | 1.819557306 | 0.001133974 | 0.024886665 | F-box/LRR-repeat protein 8 |
| FBXO10 | 2604.965377 | 1509.267072 | 0.579380857 | 0.000983914 | 0.022339882 | F-box only protein 10 |
| FBXO16 | 88.94742685 | 46.53393875 | 0.523162281 | 0.002483391 | 0.044445622 | |
| FBXO44 | 334.3150885 | 200.5384474 | 0.599848629 | 0.002417565 | 0.04351896 | F-box only protein 44 |
| FBXW7 | 1538.584432 | 724.2519851 | 0.470726188 | 5.66E-05 | 0.002513626 | cDNA FLJ55681, highly similar to F-box/WD repeat protein 7 |
| FDFT1 | 5469.109157 | 10766.14351 | 1.968536959 | 8.53E-06 | 0.000551478 | cDNA FLJ33164 fis, clone UTERU2000542, highly similar to Squalene synthetase (EC 2.5.1.21) |
| FEM1C | 765.6673494 | 1294.569508 | 1.690772773 | 0.000570519 | 0.015052224 | Protein fem-1 homolog C |
| FERMT3 | 567.8100355 | 1086.254161 | 1.91305911 | 0.001145619 | 0.025042602 | Fermitin family homolog 3 |
| FETUB | 10.9949499 | 1.022227488 | 0.092972455 | 0.000146843 | 0.005311261 | cDNA FLJ50352, highly similar to Fetuin-B |
| FGD2 | 4.742228056 | 0 | 0 | 0.002645807 | 0.046622956 | FYVE, RhoGEF and PH domain-containing protein 2 |
| FGF1 | 106.3208464 | 17.48023007 | 0.164410186 | 1.94E-07 | 2.39E-05 | Fibroblast growth factor 1 |
| FGF14 | 227.9323795 | 129.2332126 | 0.566980492 | 0.000842765 | 0.019992182 | Fibroblast growth factor 14 |
| FGF9 | 81.39691061 | 186.0205861 | 2.285351922 | 1.13E-05 | 0.000683608 | Fibroblast growth factor 9 |
| FGFBP2 | 7.719832359 | 277.2003754 | 35.90756411 | 2.12E-21 | 3.87E-18 | Fibroblast growth factor-binding protein 2 |
| FHAD1 | 100.5770651 | 216.6697197 | 2.154265682 | 0.002062114 | 0.038363703 | |
| FHL2 | 678.6429973 | 405.860027 | 0.598046438 | 0.002420509 | 0.043546758 | Four and a half LIM domains protein 2 |
| FIBCD1 | 92.75977063 | 38.49551881 | 0.415002307 | 0.000273279 | 0.008466743 | Fibrinogen C domain-containing protein 1 |
| FIGN | 389.1887846 | 212.3720802 | 0.545678829 | 0.000575334 | 0.015141746 | Fidgetin |
| FITM1 | 345.5171454 | 715.0431176 | 2.069486644 | 0.001677556 | 0.03306847 | Fat storage-inducing transmembrane protein 1 |
| FJX1 | 351.9926241 | 149.3846991 | 0.424397242 | 8.99E-06 | 0.000574269 | Four-jointed box protein 1 |
| FKBP14 | 668.502621 | 269.4602882 | 0.403080377 | 1.48E-08 | 2.62E-06 | Peptidyl-prolyl cis-trans isomerase |
| FKBP1A | 7126.924035 | 4256.60013 | 0.597256279 | 0.000462681 | 0.01283868 | Peptidyl-prolyl cis-trans isomerase FKBP1A |
| FKBP3 | 3199.61866 | 1893.387568 | 0.591754134 | 0.000442498 | 0.012433904 | Peptidyl-prolyl cis-trans isomerase FKBP3 |
| FNDC1 | 140.7308771 | 32.15274768 | 0.228469746 | 4.85E-06 | 0.000348189 | Fibronectin type III domain-containing protein 1 |
| FNIP2 | 880.8924854 | 1536.414124 | 1.744156238 | 0.000161448 | 0.005713329 | Folliculin-interacting protein 2 |
| FOLR1 | 1705.557766 | 483.4040866 | 0.283428739 | 2.33E-06 | 0.000187763 | Folate receptor alpha |

| Gene name | **Controls** normalized mean counts | **Simvastatin** normalized mean counts | foldChange | pval | padj | Protein name |
|---|---|---|---|---|---|---|
| FOXA1 | 25.66239863 | 8.707903085 | 0.339325377 | 0.001572792 | 0.031788995 | cDNA, FLJ79259, highly similar to Hepatocyte nuclear factor 3-alpha |
| FOXD2-AS1 | 20.29234875 | 45.99419596 | 2.266578232 | 0.002479119 | 0.044395787 | |
| FOXJ2 | 623.6477683 | 998.5254 | 1.601104743 | 0.001792762 | 0.034338456 | Forkhead box protein J2 |
| FOXK1 | 702.1980522 | 1467.076787 | 2.089263538 | 7.89E-06 | 0.000513507 | Forkhead box protein K1 |
| FOXN3 | 1629.639545 | 2622.329452 | 1.609146918 | 0.001231079 | 0.026506209 | Forkhead box protein N3 |
| FOXO3 | 2370.749543 | 4015.991822 | 1.693975576 | 0.000548939 | 0.014619252 | cDNA FLJ53219, highly similar to Forkhead box protein O3A |
| FOXS1 | 21.61833269 | 3.787439322 | 0.175195718 | 0.000307072 | 0.009228782 | Forkhead box protein S1 |
| FRAS1 | 2924.861693 | 1151.157276 | 0.393576653 | 0.000909486 | 0.021199286 | Extracellular matrix protein FRAS1 |
| FRMD6 | 3819.485049 | 1689.727262 | 0.442396616 | 8.22E-08 | 1.16E-05 | FERM domain-containing protein 6 |
| FRMPD4 | 65.79083089 | 25.31079098 | 0.384716086 | 0.000962448 | 0.021980843 | FERM and PDZ domain-containing protein 4 |
| FSD1 | 28.71574431 | 5.73762004 | 0.199807464 | 1.63E-05 | 0.000927681 | Fibronectin type III and SPRY domain-containing protein 1 |
| FSHR | 99.78764564 | 11.31396186 | 0.113380387 | 6.77E-08 | 9.93E-06 | Follicle-stimulating hormone receptor |
| FSTL1 | 10607.68157 | 3941.106728 | 0.37153328 | 1.59E-09 | 3.78E-07 | cDNA FLJ50214, highly similar to Follistatin-related protein 1 |
| FXN | 214.1469788 | 117.1684148 | 0.547140172 | 0.000763304 | 0.018578507 | Frataxin, mitochondrial |
| FXYD6 | 261.1814606 | 1161.889665 | 4.448591653 | 1.64E-17 | 1.82E-14 | FXYD domain-containing ion transport regulator 6 |
| FZD7 | 319.3783635 | 580.4451302 | 1.817421581 | 0.002736442 | 0.0479278 | Frizzled-7 |
| GAB1 | 265.8139018 | 483.0693626 | 1.817321665 | 0.000295675 | 0.008972928 | GRB2-associated-binding protein 1 |
| GADD45G | 1026.852222 | 625.6975503 | 0.609335537 | 0.001322229 | 0.027827641 | Growth arrest and DNA damage-inducible protein GADD45 gamma |
| GALNT10 | 1932.137062 | 586.2963508 | 0.303444493 | 2.19E-07 | 2.62E-05 | Polypeptide N-acetylgalactosaminyltransferase 10 |
| GALNT13 | 63.18641432 | 171.7949732 | 2.718859347 | 0.001752242 | 0.033854179 | Polypeptide N-acetylgalactosaminyltransferase 13 |
| GALNT15 | 130.0220607 | 395.5732874 | 3.042355161 | 0.000364246 | 0.010648091 | Polypeptide N-acetylgalactosaminyltransferase 15 |
| GANC | 497.9509316 | 829.7240173 | 1.666276664 | 0.001275448 | 0.027099788 | Neutral alpha-glucosidase C |
| GBE1 | 3134.33265 | 5102.620761 | 1.627976775 | 0.00094284 | 0.021692293 | 1,4-alpha-glucan-branching enzyme |
| GBP1 | 193.413068 | 55.57508731 | 0.287338844 | 4.15E-09 | 9.02E-07 | Interferon-induced guanylate-binding protein 1 |
| GFM2 | 1431.950133 | 808.7099658 | 0.564761263 | 0.000153597 | 0.005498197 | Ribosome-releasing factor 2, mitochondrial |
| GFPT2 | 331.5294019 | 1033.81502 | 3.118320772 | 0.001277527 | 0.027106916 | Glutamine--fructose-6-phosphate aminotransferase [isomerizing] 2 |
| GGH | 322.5320066 | 171.4050429 | 0.531435763 | 0.000649879 | 0.016435062 | Gamma-glutamyl hydrolase |
| GINS4 | 47.24879541 | 18.08523567 | 0.382766069 | 0.000770568 | 0.018696793 | DNA replication complex GINS protein SLD5 |
| GIT1 | 1468.963362 | 2522.499102 | 1.717196744 | 0.000257064 | 0.008051218 | ARF GTPase-activating protein GIT1 |

| Gene name | Controls normalized mean counts | Simvastatin normalized mean counts | foldChange | pval | padj | Protein name |
|---|---|---|---|---|---|---|
| GIT2 | 2259.317338 | 1077.719923 | 0.47701131 | 1.02E-05 | 0.000627502 | ARF GTPase-activating protein GIT2 |
| GJD4 | 100.9645861 | 306.3434835 | 3.034167675 | 1.84E-06 | 0.000155867 | Gap junction delta-4 protein |
| GLCCI1 | 88.50875788 | 180.7065129 | 2.041679459 | 0.000153569 | 0.005498197 | Glucocorticoid-induced transcript 1 protein |
| GLDC | 44.84316783 | 17.58060753 | 0.392046512 | 0.000302806 | 0.009135872 | Glycine dehydrogenase (decarboxylating), mitochondrial |
| GLDN | 5.516277446 | 34.50005524 | 6.254227707 | 1.66E-05 | 0.000936421 | Gliomedin |
| GLIS3 | 133.4734127 | 281.0046934 | 2.105323358 | 0.000234182 | 0.00750204 | GLIS family zinc finger 3 transcript variant TS7 |
| GLRX | 2778.56895 | 1184.806524 | 0.426408898 | 2.61E-08 | 4.39E-06 | Glutaredoxin-1 |
| GLRX5 | 1451.369873 | 920.5598128 | 0.634269617 | 0.002741403 | 0.047936428 | Glutaredoxin-related protein 5, mitochondrial |
| GLS | 5344.706407 | 2740.962017 | 0.512836779 | 2.39E-05 | 0.001236461 | Glutaminase kidney isoform, mitochondrial |
| GLT8D2 | 407.6449454 | 190.0323572 | 0.466171258 | 0.000101936 | 0.003973452 | Glycosyltransferase 8 domain-containing protein 2 |
| GLUL | 1409.148076 | 3564.782515 | 2.529743024 | 2.89E-10 | 8.32E-08 | Glutamine synthetase |
| GLYCTK-AS1 | 18.51922472 | 43.12532182 | 2.328678574 | 0.002852799 | 0.049417017 | |
| GMIP | 485.8201302 | 266.1040003 | 0.547741816 | 0.000167287 | 0.005842717 | GMIP protein |
| GNAZ | 288.1027816 | 163.5756272 | 0.567768302 | 0.000826063 | 0.019683535 | Guanine nucleotide-binding protein G(z) subunit alpha |
| GNG2 | 489.0398445 | 236.994587 | 0.48461202 | 6.15E-05 | 0.002674521 | Guanine nucleotide-binding protein subunit gamma |
| GNGT2 | 2.567136018 | 21.28878429 | 8.292815083 | 0.000543992 | 0.014537304 | Guanine nucleotide-binding protein subunit gamma |
| GNLY | 2.269560694 | 14.50938281 | 6.393035818 | 0.001122614 | 0.024695924 | Granulysin |
| GNPTAB | 707.3642523 | 1352.668709 | 1.91226614 | 2.10E-05 | 0.001120874 | N-acetylglucosamine-1-phosphotransferase subunit alpha |
| GOLIM4 | 988.6798069 | 1798.200338 | 1.818789385 | 0.000121439 | 0.004540248 | Golgi integral membrane protein 4 |
| GOSR2 | 2193.32286 | 1356.826771 | 0.618616983 | 0.001358606 | 0.028401071 | Golgi SNAP receptor complex member 2 |
| GPATCH4 | 1267.230197 | 778.0056557 | 0.613941854 | 0.00148934 | 0.030398569 | G patch domain-containing protein 4 |
| GPC2 | 1452.33674 | 586.8264536 | 0.404056743 | 4.60E-05 | 0.002110623 | Glypican-2 |
| GPD2 | 257.0440772 | 472.8565961 | 1.839593432 | 0.002100583 | 0.038960929 | Glycerol-3-phosphate dehydrogenase, mitochondrial |
| GPR125 | 1545.897704 | 924.6530313 | 0.598133388 | 0.00135995 | 0.028405318 | Probable G-protein-coupled receptor 125 |
| GPR126 | 51.83045355 | 9.785659752 | 0.188801353 | 4.79E-09 | 1.01E-06 | G-protein coupled receptor 126 |
| GPR179 | 3.52616082 | 16.45640692 | 4.666947356 | 0.001762007 | 0.033956313 | Probable G-protein coupled receptor 179 |
| GPR68 | 19.06979931 | 87.58297128 | 4.592757893 | 0.000340001 | 0.010024711 | Ovarian cancer G-protein-coupled receptor 1 |
| GPSM2 | 1302.200923 | 2430.782716 | 1.866672548 | 0.000631046 | 0.016142529 | G-protein-signaling modulator 2 |
| GPX1 | 2584.563315 | 769.1340697 | 0.297587629 | 2.05E-10 | 6.26E-08 | Glutathione peroxidase 1 |
| GPX3 | 460.1995617 | 1723.755004 | 3.745668503 | 2.71E-16 | 2.63E-13 | Glutathione peroxidase |
| GRAMD3 | 6777.995204 | 2929.500909 | 0.432207581 | 0.001252877 | 0.026840216 | GRAM domain-containing protein 3 |
| GRIA1 | 85.28879302 | 34.7028273 | 0.406886134 | 0.001364977 | 0.028457754 | Glutamate receptor 1 |

| Gene name | **Controls** normalized mean counts | **Simvastatin** normalized mean counts | foldChange | pval | padj | Protein name |
|---|---|---|---|---|---|---|
| GRIA3 | 180.0808654 | 47.54826106 | 0.264038386 | 0.000462655 | 0.01283868 | cDNA FLJ51894, highly similar to Glutamate receptor 3 |
| GRK5 | 121.7691597 | 224.8799955 | 1.846772992 | 0.002303826 | 0.042080339 | G protein-coupled receptor kinase 5 |
| GS1-259H13.10 | 362.5625518 | 659.6532381 | 1.819419118 | 0.001672778 | 0.033045663 | |
| GSG1 | 1678.791066 | 94.00690681 | 0.055996788 | 7.61E-07 | 7.49E-05 | Germ cell-specific gene 1 protein |
| GSR | 1325.375292 | 2194.652956 | 1.655872845 | 0.000657618 | 0.016590332 | Glutathione reductase, mitochondrial |
| GTF2A2 | 1057.239089 | 599.8458428 | 0.567370095 | 0.000237792 | 0.007609818 | Transcription initiation factor IIA subunit 2 |
| GTF2H2C | 278.1191295 | 163.4957542 | 0.587862311 | 0.001969118 | 0.036809722 | Uncharacterized protein |
| GTPBP4 | 3041.938112 | 1727.012628 | 0.567734308 | 0.000310113 | 0.009280806 | Nucleolar GTP-binding protein 1 |
| GUCY1B3 | 83.9596871 | 33.62409283 | 0.400479015 | 0.000506245 | 0.013813389 | cDNA FLJ59711, highly similar to Guanylate cyclase soluble subunit beta-1 (EC 4.6.1.2) |
| H1FX-AS1 | 51.51924281 | 99.83616427 | 1.937842228 | 0.001773786 | 0.034037863 | |
| HARS | 1869.004709 | 1119.936622 | 0.599215516 | 0.000747077 | 0.018344996 | cDNA FLJ42841 fis, clone BRCOC2003213, highly similar to Histidyl-tRNA synthetase (EC 6.1.1.21) |
| HCG18 | 411.6372253 | 243.7884724 | 0.592241074 | 0.001665562 | 0.032981381 | |
| HCN3 | 210.9269652 | 354.6023618 | 1.681161825 | 0.001377337 | 0.028677013 | Potassium/sodium hyperpolarization-activated cyclic nucleotide-gated channel 3 |
| HCST | 23.60023247 | 5.343814949 | 0.226430606 | 0.000249956 | 0.007900081 | Hematopoietic cell signal transducer |
| HDAC5 | 917.2272329 | 2321.904311 | 2.531438478 | 4.43E-10 | 1.22E-07 | Histone deacetylase 5 |
| HDAC9 | 1851.791315 | 5138.538592 | 2.774901551 | 2.41E-09 | 5.54E-07 | Histone deacetylase 9 |
| HDDC2 | 2038.999718 | 1206.506521 | 0.591714903 | 0.000447603 | 0.012554689 | HD domain-containing protein 2 |
| HEATR3 | 710.7280776 | 415.0702501 | 0.584007109 | 0.000606157 | 0.015712607 | |
| HECA | 471.6414284 | 1119.558911 | 2.37375015 | 1.40E-08 | 2.53E-06 | Headcase protein homolog |
| HECW2 | 233.7094523 | 692.5745506 | 2.963399828 | 8.06E-07 | 7.86E-05 | E3 ubiquitin-protein ligase HECW2 |
| HEG1 | 1247.599652 | 2794.55261 | 2.239943403 | 7.79E-07 | 7.62E-05 | Protein HEG homolog 1 |
| HEPH | 1259.285159 | 398.3301576 | 0.316314502 | 9.16E-06 | 0.000583614 | Hephaestin |
| HHATL | 154.6198964 | 332.4190565 | 2.149911261 | 0.001814613 | 0.034671599 | Protein-cysteine N-palmitoyltransferase HHAT-like protein |
| HID1 | 214.0075611 | 56.36840169 | 0.26339444 | 0.000774822 | 0.018756122 | Protein HID1 |
| HIF1A | 7303.98517 | 4087.008954 | 0.559558769 | 0.000200213 | 0.00663241 | Hypoxia-inducible factor 1, alpha subunit (Basic helix-loop-helix transcription factor), isoform CRA_a |
| HIST1H2BC | 279.951321 | 157.9280998 | 0.564127003 | 0.001744975 | 0.03377672 | Histone H2B type 1-C/E/F/G/I |
| HIST1H4H | 366.7843559 | 201.3425575 | 0.548939872 | 0.001175414 | 0.025550263 | |
| HIVEP3 | 990.2331233 | 250.7626535 | 0.253235978 | 5.12E-10 | 1.37E-07 | Transcription factor HIVEP3 |
| HK3 | 14.26635763 | 2.890579356 | 0.202615091 | 0.001423762 | 0.02936839 | Hexokinase-3 |
| HMBOX1 | 960.7621507 | 1907.339277 | 1.985235654 | 9.88E-06 | 0.000612322 | Homeobox-containing protein 1 |

| Gene name | **Controls** normalized mean counts | **Simvastatin** normalized mean counts | foldChange | pval | padj | Protein name |
|---|---|---|---|---|---|---|
| HMCES | 1106.289324 | 651.9800721 | 0.589339568 | 0.0005336 | 0.014345858 | Embryonic stem cell-specific 5-hydroxymethylcytosine-binding protein |
| HMGB2 | 465.8886566 | 1415.69441 | 3.038696886 | 3.96E-11 | 1.42E-08 | High mobility group protein B2 |
| HMGCR | 2286.817119 | 5226.444177 | 2.285466613 | 7.12E-05 | 0.002993667 | 3-hydroxy-3-methylglutaryl-coenzyme A reductase |
| HMGCS1 | 5774.386649 | 18875.15944 | 3.268773046 | 5.77E-06 | 0.000400997 | Hydroxymethylglutaryl-CoA synthase, cytoplasmic |
| HN1 | 10958.99015 | 6829.907335 | 0.623224151 | 0.001896077 | 0.035919228 | Hematological and neurological expressed 1 protein |
| HNRNPA1 | 8149.893085 | 4525.857258 | 0.555327194 | 7.65E-05 | 0.003159373 | Heterogeneous nuclear ribonucleoprotein A1 |
| HNRNPM | 3350.035974 | 1965.58414 | 0.586735234 | 0.000439039 | 0.012381448 | Heterogeneous nuclear ribonucleoprotein M |
| HNRNPU-AS1 | 172.3076779 | 473.9354339 | 2.750518373 | 4.46E-08 | 6.97E-06 | |
| HOPX | 10.58797009 | 54.22960568 | 5.121813269 | 1.82E-07 | 2.26E-05 | Homeodomain-only protein |
| HPGD | 8.32230921 | 36.63579801 | 4.402119302 | 4.25E-05 | 0.001985755 | 15-hydroxyprostaglandin dehydrogenase [NAD(+)] |
| HPS5 | 1030.623988 | 443.9144777 | 0.430723991 | 6.02E-08 | 8.92E-06 | Hermansky-Pudlak syndrome 5 protein |
| HRH1 | 154.8670922 | 65.18303082 | 0.420896589 | 2.46E-05 | 0.001261659 | Histamine H1 receptor |
| hsa-mir-6080 | 417.2764788 | 710.935712 | 1.703752184 | 0.000767889 | 0.018646333 | |
| HSD11B2 | 52.94873036 | 113.6654749 | 2.146708223 | 0.000294392 | 0.008969009 | Corticosteroid 11-beta-dehydrogenase isozyme 2 |
| HSD17B1 | 131.4448226 | 36.99188802 | 0.281425219 | 0.000955784 | 0.021909069 | cDNA FLJ50217, highly similar to Estradiol 17-beta-dehydrogenase 1 (EC 1.1.1.62) |
| HSD17B10 | 1368.682706 | 793.2966857 | 0.579605983 | 0.000546673 | 0.014583891 | 3-hydroxyacyl-CoA dehydrogenase type-2 |
| HSD17B2 | 35.02331963 | 9.057212655 | 0.258605202 | 0.001650647 | 0.03285031 | Estradiol 17-beta-dehydrogenase 2 |
| HSD17B3 | 10.51939735 | 31.15631277 | 2.96179636 | 0.001481797 | 0.030284352 | Testosterone 17-beta-dehydrogenase 3 |
| HSD17B6 | 45.04140482 | 9.8446498 | 0.218568889 | 7.28E-05 | 0.003041939 | 17-beta-hydroxysteroid dehydrogenase type 6 |
| HSD17B7 | 218.4991005 | 524.7888019 | 2.401789301 | 2.65E-08 | 4.43E-06 | 3-keto-steroid reductase |
| HSD3B7 | 359.629741 | 832.0212429 | 2.313549598 | 8.79E-08 | 1.21E-05 | 3 beta-hydroxysteroid dehydrogenase type 7 |
| HSF2BP | 270.6587165 | 40.95732031 | 0.151324594 | 1.53E-09 | 3.66E-07 | Heat shock factor 2-binding protein |
| HSF4 | 150.6347193 | 428.5787489 | 2.845152505 | 4.57E-10 | 1.25E-07 | Heat shock factor protein 4 |
| HSPA1A | 1243.508012 | 2322.514285 | 1.867711557 | 0.000156378 | 0.005571921 | Heat shock 70 kDa protein 1A/1B |
| HSPA1B | 817.7996064 | 2125.431031 | 2.598963138 | 3.08E-05 | 0.001520621 | |
| HSPA4L | 1533.031791 | 921.6106998 | 0.601168681 | 0.0010821 | 0.024025566 | Heat shock 70 kDa protein 4L |
| HSPB1 | 16830.54879 | 41091.37688 | 2.441475759 | 9.49E-10 | 2.38E-07 | Heat shock protein beta-1 |
| HSPB7 | 2039.236869 | 9646.778197 | 4.730582477 | 1.43E-23 | 3.42E-20 | cDNA FLJ34956 fis, clone NTONG2003158, highly similar to Heat shock 27kD protein family, member 7 |
| HSPB8 | 8798.005561 | 15438.72838 | 1.754798662 | 0.000257021 | 0.008051218 | Heat shock protein beta-8 |
| HUNK | 512.4131362 | 254.79251 | 0.497240395 | 1.43E-05 | 0.000828648 | Hormonally up-regulated neu tumor-associated kinase |

| Gene name | Controls normalized mean counts | Simvastatin normalized mean counts | foldChange | pval | padj | Protein name |
|---|---|---|---|---|---|---|
| ID2 | 350.2521354 | 150.2191191 | 0.428888518 | 2.78E-05 | 0.001390079 | DNA-binding protein inhibitor ID-2 |
| IDI1 | 5165.583591 | 10673.04736 | 2.066184232 | 1.75E-05 | 0.000964572 | Isopentenyl-diphosphate Delta-isomerase 1 |
| IER2 | 500.6421355 | 832.0356736 | 1.66193697 | 0.000709218 | 0.017620546 | Immediate early response gene 2 protein |
| IFFO2 | 605.5064208 | 349.0313226 | 0.576428772 | 0.000715971 | 0.017731686 | Intermediate filament family orphan 2 |
| IFIH1 | 27.27825702 | 65.14396712 | 2.388127917 | 0.0026298 | 0.046452332 | Interferon-induced helicase C domain-containing protein 1 |
| IFIT1 | 5.988902101 | 33.89793468 | 5.660125029 | 2.80E-05 | 0.001402292 | Interferon-induced protein with tetratricopeptide repeats 1, isoform CRA_b |
| IFIT2 | 4.213523702 | 21.73224574 | 5.157736677 | 0.001581371 | 0.031920917 | Interferon-induced protein with tetratricopeptide repeats 2 |
| IFIT3 | 13.48202122 | 55.55241138 | 4.120480934 | 1.69E-05 | 0.000949258 | Interferon-induced protein with tetratricopeptide repeats 3 |
| IFNG-AS1 | 29.50775038 | 69.13451902 | 2.342927473 | 0.000536122 | 0.014388784 | |
| IFRD1 | 1135.407329 | 2180.437524 | 1.920401135 | 2.56E-05 | 0.00129836 | Interferon-related developmental regulator 1 |
| IGFBP1 | 16.8105565 | 710.9780382 | 42.29354561 | 1.44E-08 | 2.59E-06 | Insulin-like growth factor-binding protein 1 |
| IGFBP3 | 5194.085773 | 1571.612286 | 0.302577269 | 7.67E-08 | 1.09E-05 | Insulin-like growth factor binding protein 3, isoform CRA_b |
| IGFBP7 | 3646.057621 | 1398.418266 | 0.383542558 | 1.68E-05 | 0.000945411 | Insulin-like growth factor-binding protein 7 |
| IGIP | 137.9602668 | 236.5844304 | 1.71487368 | 0.00235521 | 0.042643284 | IgA-inducing protein homolog |
| IGSF1 | 30.44309316 | 93.16224651 | 3.060209619 | 1.21E-05 | 0.000718123 | Immunoglobulin superfamily member 1 |
| IL1RAP | 429.5102047 | 210.1655688 | 0.489314495 | 0.0013161 | 0.027717396 | Interleukin-1 receptor accessory protein |
| IL2RG | 11.59059903 | 119.3820908 | 10.29990688 | 3.91E-19 | 5.53E-16 | Cytokine receptor common subunit gamma |
| IL31RA | 7.018238039 | 23.98675387 | 3.417774338 | 0.002556291 | 0.045385842 | Interleukin-31 receptor subunit alpha |
| IL6R | 68.99701855 | 156.2049409 | 2.263937547 | 0.000644255 | 0.016399421 | Interleukin-6 receptor subunit alpha |
| ILDR2 | 54.9415844 | 17.01329172 | 0.309661469 | 7.99E-05 | 0.003268696 | Immunoglobulin-like domain-containing receptor 2 |
| INADL | 83.71051851 | 153.4594521 | 1.833215883 | 0.001576708 | 0.031847462 | InaD-like protein |
| INHBA | 155.1391882 | 29.13874778 | 0.187823258 | 3.98E-09 | 8.79E-07 | Inhibin beta A chain |
| INHBA-AS1 | 55.40317178 | 12.22180698 | 0.220597605 | 0.000291361 | 0.008888582 | |
| INHBB | 27.9572597 | 212.3153456 | 7.594283126 | 1.14E-05 | 0.000687614 | Inhibin beta B chain |
| INHBE | 98.33678209 | 258.9366309 | 2.633161523 | 5.53E-06 | 0.00039017 | Inhibin beta E chain |
| INPP5A | 472.8978662 | 811.2620838 | 1.715512253 | 0.000421896 | 0.012006873 | Type I inositol 1,4,5-trisphosphate 5-phosphatase |
| INSIG1 | 2474.908904 | 6958.740745 | 2.811715912 | 5.20E-08 | 7.93E-06 | Insulin-induced gene protein |
| INSR | 399.5988508 | 915.3731918 | 2.290730291 | 9.26E-08 | 1.26E-05 | Insulin receptor |
| INSRR | 2.251008469 | 18.66470641 | 8.291708655 | 7.69E-05 | 0.003174127 | Insulin receptor-related protein |
| INTS6-AS1 | 45.11058442 | 93.65239186 | 2.076062482 | 0.001418125 | 0.029310425 | |

| Gene name | Controls normalized mean counts | Simvastatin normalized mean counts | foldChange | pval | padj | Protein name |
|---|---|---|---|---|---|---|
| INTU | 530.7276384 | 233.2818805 | 0.439551031 | 3.44E-07 | 3.82E-05 | Protein inturned |
| IP6K3 | 1964.485063 | 1167.888831 | 0.594501252 | 0.000671764 | 0.016865136 | Inositol hexakisphosphate kinase 3 |
| IPO4 | 1637.074431 | 800.2947316 | 0.488856656 | 3.64E-06 | 0.000271315 | Importin-4 |
| IRAK2 | 74.2167802 | 231.5937739 | 3.120504194 | 4.97E-08 | 7.63E-06 | Interleukin-1 receptor-associated kinase-like 2 |
| IRF6 | 31.25361854 | 320.9780222 | 10.2701075 | 3.16E-16 | 2.89E-13 | Interferon regulatory factor 6 |
| IRX3 | 951.3084432 | 551.6251547 | 0.579859412 | 0.001967118 | 0.036794443 | Iroquois-class homeodomain protein IRX-3 |
| ITGA1 | 1418.233047 | 544.643293 | 0.384029475 | 0.000192207 | 0.006498703 | Integrin alpha-1 |
| ITGA5 | 4626.075599 | 2954.426278 | 0.638646346 | 0.002473464 | 0.044320034 | Integrin alpha-5 light chain |
| ITGA6 | 33973.94884 | 16487.49952 | 0.485298297 | 0.000112356 | 0.004301247 | Integrin alpha-6 heavy chain |
| ITGAD | 12.07914311 | 77.80567112 | 6.441323724 | 3.93E-05 | 0.001866041 | Integrin alpha-D |
| ITGAX | 8.713912677 | 29.29473191 | 3.361834459 | 0.001640024 | 0.032722629 | Integrin alpha-X |
| ITGB1BP2 | 2718.788005 | 1244.142747 | 0.457609326 | 0.000201293 | 0.00666109 | Integrin beta-1-binding protein 2 |
| ITGB7 | 9.801850681 | 28.29268712 | 2.886463795 | 0.002662201 | 0.046811999 | Integrin beta |
| ITGBL1 | 3892.808222 | 1786.734764 | 0.458983506 | 0.00166395 | 0.032981381 | Integrin beta-like protein 1 |
| ITIH1 | 3.902578423 | 35.53561345 | 9.105675683 | 0.00059066 | 0.015439561 | Inter-alpha-trypsin inhibitor heavy chain H1 |
| ITIH3 | 9.065734303 | 79.13226074 | 8.728720487 | 1.61E-11 | 6.22E-09 | Inter-alpha-trypsin inhibitor heavy chain H3 |
| ITIH4 | 244.1068039 | 1318.240332 | 5.400260506 | 1.95E-07 | 2.39E-05 | ITIH4 protein |
| ITPKA | 7.051460158 | 57.07992034 | 8.094766056 | 2.88E-09 | 6.49E-07 | Inositol-trisphosphate 3-kinase A |
| ITPKB | 115.0581964 | 281.8734808 | 2.449833993 | 6.51E-05 | 0.002780005 | Inositol-trisphosphate 3-kinase B |
| ITPR2 | 564.2169596 | 317.8902982 | 0.563418545 | 0.000376888 | 0.010946287 | Inositol 1,4,5-trisphosphate receptor type 2 |
| ITPR3 | 701.8922245 | 2024.136804 | 2.883828505 | 3.11E-06 | 0.000237672 | Inositol 1,4,5-trisphosphate receptor type 3 |
| IZUMO4 | 51.43397629 | 108.5365076 | 2.110210321 | 0.002375423 | 0.042959254 | Izumo sperm-egg fusion protein 4 |
| JAK2 | 389.7545664 | 678.3172668 | 1.740370287 | 0.000457468 | 0.012739473 | Tyrosine-protein kinase |
| JAM2 | 5512.547083 | 1879.475615 | 0.340945045 | 9.03E-07 | 8.64E-05 | Junctional adhesion molecule B |
| JHDM1D | 418.732613 | 753.684587 | 1.799918525 | 0.000397844 | 0.011437474 | |
| JMJD4 | 410.193172 | 216.0870454 | 0.526793375 | 9.95E-05 | 0.003907758 | |
| JUN | 1691.892136 | 3203.973458 | 1.893722058 | 2.47E-05 | 0.001261659 | Transcription factor AP-1 |
| JUND | 1072.658798 | 2277.544425 | 2.123270167 | 3.69E-07 | 4.03E-05 | Transcription factor jun-D |
| KAT2B | 786.3259632 | 1286.995827 | 1.636720504 | 0.001097291 | 0.024293471 | Histone acetyltransferase KAT2B |
| KCNA1 | 4.680527631 | 48.57405408 | 10.37790136 | 3.64E-05 | 0.001745022 | Potassium voltage-gated channel subfamily A member 1 |
| KCNA7 | 69.79104771 | 206.5944573 | 2.96018564 | 0.00012061 | 0.004520117 | Potassium voltage-gated channel subfamily A member 7 |
| KCNC4 | 1003.318161 | 488.5216138 | 0.486905981 | 0.001765031 | 0.033974667 | Potassium voltage-gated channel subfamily C member 4 |
| KCND3 | 466.8155894 | 115.3270572 | 0.247050569 | 3.43E-08 | 5.64E-06 | Potassium voltage-gated channel subfamily D member 3 |
| KCNE1L | 688.631645 | 88.95805579 | 0.1291809 | 9.72E-11 | 3.22E-08 | Potassium voltage-gated channel subfamily E member 1-like protein |

| Gene name | **Controls** normalized mean counts | **Simvastatin** normalized mean counts | **foldChange** | **pval** | **padj** | **Protein name** |
|---|---|---|---|---|---|---|
| KCNH1 | 523.179948 | 260.5029492 | 0.497922274 | 0.001677445 | 0.03306847 | Potassium voltage-gated channel subfamily H member 1 |
| KCNH3 | 39.78362066 | 109.3576814 | 2.748811685 | 5.38E-05 | 0.002413297 | Potassium voltage-gated channel subfamily H member 3 |
| KCNIP2 | 34.62861749 | 83.11880719 | 2.400292394 | 0.000906704 | 0.021158236 | Kv channel-interacting protein 2 |
| KCNJ12 | 1272.433332 | 519.7367312 | 0.408458909 | 6.16E-05 | 0.002674521 | ATP-sensitive inward rectifier potassium channel 12 |
| KCNJ2-AS1 | 138.6057884 | 315.7975558 | 2.278386491 | 0.000216664 | 0.007079352 | |
| KCNJ6 | 2020.588049 | 940.7237356 | 0.465569286 | 0.000211259 | 0.006961263 | G protein-activated inward rectifier potassium channel 2 |
| KCNMA1 | 652.1504404 | 1431.983569 | 2.195787169 | 2.90E-05 | 0.001446153 | Calcium-activated potassium channel subunit alpha-1 |
| KCTD17 | 730.1149349 | 336.5663057 | 0.460977155 | 1.75E-06 | 0.000149495 | BTB/POZ domain-containing protein KCTD17 |
| KDSR | 625.1026392 | 1022.022737 | 1.634967881 | 0.001268725 | 0.027039148 | 3-ketodihydrosphingosine reductase |
| KIAA0020 | 1109.848663 | 701.7332113 | 0.632278287 | 0.002823753 | 0.049042798 | Pumilio domain-containing protein KIAA0020 |
| KIAA0226 | 464.0307145 | 779.1557395 | 1.679103807 | 0.000790721 | 0.019037281 | Run domain Beclin-1 interacting and cysteine-rich containing protein |
| KIAA0232 | 2099.21948 | 3619.064285 | 1.724004716 | 0.000216609 | 0.007079352 | Uncharacterized protein KIAA0232 |
| KIAA1024 | 95.14245097 | 41.77960198 | 0.43912682 | 0.000116837 | 0.004410612 | UPF0258 protein KIAA1024 |
| KIAA1109 | 2093.057691 | 3421.983927 | 1.634920978 | 0.000916832 | 0.021314624 | Uncharacterized protein KIAA1109 |
| KIAA1199 | 2386.343023 | 639.8588929 | 0.268133662 | 1.43E-15 | 1.17E-12 | Protein KIAA1199 |
| KIAA1217 | 578.251398 | 1305.334712 | 2.257382718 | 0.001713861 | 0.033466747 | Sickle tail protein homolog |
| KIAA1407 | 43.3881853 | 114.2919692 | 2.634172608 | 7.58E-06 | 0.000501704 | |
| KIAA1467 | 795.0840822 | 232.0901285 | 0.291906395 | 3.80E-10 | 1.07E-07 | Uncharacterized protein KIAA1467 |
| KIAA1522 | 409.9057508 | 832.078369 | 2.029926068 | 9.28E-06 | 0.00058816 | |
| KIAA1549L | 75.63839931 | 21.53030803 | 0.284647854 | 2.38E-06 | 0.000190388 | UPF0606 protein KIAA1549L |
| KIAA1671 | 794.2973153 | 1370.892562 | 1.725918665 | 0.000478827 | 0.013169227 | |
| KIAA1683 | 26.56413879 | 78.85098729 | 2.968324624 | 6.61E-06 | 0.000449074 | Uncharacterized protein KIAA1683 |
| KIF13B | 1312.068957 | 2135.503945 | 1.627585147 | 0.001601318 | 0.032135878 | Kinesin-like protein KIF13B |
| KIF18B | 8.206858908 | 0.507760098 | 0.061870212 | 0.002646954 | 0.046622956 | Kinesin-like protein KIF18B |
| KIF20A | 16.26099261 | 2.726845611 | 0.167692445 | 0.001446281 | 0.029734312 | cDNA FLJ55710, highly similar to Kinesin family member 20A |
| KIF24 | 257.5371143 | 561.6820666 | 2.180975228 | 0.000258442 | 0.008079488 | Kinesin-like protein KIF24 |
| KIF6 | 34.37524439 | 10.65705236 | 0.310021137 | 0.000165522 | 0.005817787 | Kinesin-like protein KIF6 |
| KIFC2 | 377.6610016 | 739.8993129 | 1.959162608 | 1.61E-05 | 0.000917559 | Kinesin-like protein KIFC2 |
| KISS1 | 766.40718 | 54.18469408 | 0.070699617 | 1.70E-05 | 0.000950492 | Metastasis-suppressor KiSS-1 |
| KLF11 | 214.1091241 | 378.4086247 | 1.767363377 | 0.001023357 | 0.023017017 | Krueppel-like factor 11 |
| KLF13 | 1634.985987 | 3518.965528 | 2.15229094 | 6.01E-07 | 6.15E-05 | Krueppel-like factor 13 |
| KLF2 | 397.2776916 | 2168.576667 | 5.458591593 | 1.69E-12 | 8.10E-10 | Krueppel-like factor 2 |
| KLF3 | 1004.733896 | 1613.098639 | 1.605498377 | 0.001512458 | 0.030809778 | Krueppel-like factor 3 |
| KLF4 | 118.2057873 | 1544.573353 | 13.06681668 | 1.65E-52 | 1.28E-48 | Krueppel-like factor 4 |
| KLF6 | 1648.525689 | 4952.458959 | 3.004174573 | 3.42E-11 | 1.25E-08 | Krueppel-like factor 6 |
| KLF7 | 453.1581028 | 884.4653923 | 1.951781038 | 1.59E-05 | 0.000907078 | Krueppel-like factor 7 |

| Gene name | Controls normalized mean counts | Simvastatin normalized mean counts | foldChange | pval | padj | Protein name |
|---|---|---|---|---|---|---|
| KLF8 | 38.80458867 | 96.97404542 | 2.49903552 | 0.000176435 | 0.006091227 | Krueppel-like factor 8 |
| KLF9 | 427.1062313 | 1110.921836 | 2.60104338 | 3.43E-06 | 0.000257242 | Krueppel-like factor 9 |
| KLHDC9 | 1.485146765 | 14.53049771 | 9.78387998 | 0.000262054 | 0.008167779 | |
| KLHL24 | 1813.389334 | 2953.574074 | 1.628758932 | 0.000959639 | 0.021936581 | Kelch-like protein 24 |
| KLHL41 | 38010.18448 | 81063.48757 | 2.132678088 | 6.33E-07 | 6.42E-05 | Kelch-like protein 41 |
| KPNA2 | 1970.19573 | 1146.772463 | 0.582060171 | 0.000607744 | 0.015714443 | Importin subunit alpha-1 |
| KPRP | 403.6224831 | 99.10909065 | 0.245548984 | 0.001086434 | 0.024104579 | Keratinocyte proline-rich protein |
| KRAS | 167.7291568 | 626.9760766 | 3.738026761 | 5.00E-14 | 3.36E-11 | GTPase KRas |
| KREMEN1 | 1530.256782 | 4570.88567 | 2.987005661 | 1.95E-08 | 3.39E-06 | Kringle containing transmembrane protein 1, isoform CRA_c |
| KREMEN2 | 444.7033911 | 52.90300093 | 0.118962441 | 2.25E-15 | 1.75E-12 | Kremen protein 2 |
| KRT8P14 | 11.35729109 | 35.21989367 | 3.101082237 | 0.000454768 | 0.012687012 | |
| KRTAP5-AS1 | 2.690849462 | 30.56832093 | 11.36010073 | 9.21E-08 | 1.26E-05 | |
| KSR1 | 1052.401034 | 630.3054038 | 0.598921308 | 0.000958697 | 0.021936581 | Kinase suppressor of Ras 1 |
| L1CAM | 1703.747067 | 4962.419144 | 2.912650147 | 4.04E-09 | 8.84E-07 | Neural cell adhesion molecule L1 |
| L3MBTL1 | 250.7124678 | 423.1506793 | 1.687792725 | 0.001306077 | 0.027543611 | Lethal(3)malignant brain tumor-like protein 1 |
| LAMP5 | 25.50737398 | 2.272475758 | 0.089090933 | 1.86E-05 | 0.001007834 | Lysosome-associated membrane glycoprotein 5 |
| LANCL2 | 755.5543935 | 391.0078979 | 0.517511249 | 3.12E-05 | 0.001534589 | LanC-like protein 2 |
| LARP6 | 786.7429018 | 1458.026324 | 1.853243697 | 8.36E-05 | 0.003399083 | La-related protein 6 |
| LAYN | 166.2851391 | 64.17149236 | 0.385912371 | 6.94E-07 | 6.91E-05 | Layilin |
| LBP | 0.500226047 | 11.92359655 | 23.83641677 | 0.000175042 | 0.006063305 | Lipopolysaccharide-binding protein |
| LDHB | 9141.048505 | 5714.515435 | 0.625148792 | 0.001375638 | 0.028660815 | L-lactate dehydrogenase |
| LDHD | 41.62262255 | 175.640439 | 4.219831146 | 1.08E-10 | 3.51E-08 | Probable D-lactate dehydrogenase, mitochondrial |
| LDLR | 2422.412654 | 5792.166446 | 2.391073394 | 5.04E-07 | 5.33E-05 | Low-density lipoprotein receptor |
| LDOC1 | 283.2525532 | 135.5926476 | 0.478698766 | 1.80E-05 | 0.000983944 | Protein LDOC1 |
| LFNG | 382.9253347 | 51.39027619 | 0.13420443 | 2.82E-21 | 4.87E-18 | Beta-1,3-N-acetylglucosaminyltransferase lunatic fringe |
| LGALS3 | 2213.924848 | 3902.104831 | 1.762528134 | 0.00077697 | 0.0187935 | Galectin |
| LGI4 | 64.30356371 | 263.0076402 | 4.090094312 | 5.72E-09 | 1.16E-06 | Leucine-rich repeat LGI family member 4 |
| LHX4 | 200.5047199 | 66.35600562 | 0.330944856 | 0.001269117 | 0.027039148 | LIM/homeobox protein Lhx4 |
| LIMA1 | 8932.412835 | 3477.467625 | 0.389308879 | 6.84E-08 | 9.99E-06 | LIM domain and actin-binding protein 1 |
| LIMD1 | 908.9416233 | 1459.418672 | 1.605624205 | 0.002897007 | 0.049897172 | LIM domain-containing protein 1 |
| LIMK2 | 1623.084921 | 653.0905167 | 0.402376061 | 6.01E-07 | 6.15E-05 | LIM domain kinase 2 |
| LIMS3L_1 | 527.5083939 | 270.2207091 | 0.512258596 | 2.47E-05 | 0.001261659 | |
| LINC00116 | 539.9653719 | 300.116445 | 0.555806836 | 0.000826422 | 0.019683535 | Putative uncharacterized protein encoded by LINC00116 |
| LINC00152 | 1203.737286 | 542.9174938 | 0.451026565 | 2.35E-07 | 2.77E-05 | |
| LINC00299 | 6.504141499 | 0.241435365 | 0.037120251 | 0.002760946 | 0.04819415 | |
| LINC00319 | 289.8622567 | 76.57639219 | 0.264182005 | 1.39E-07 | 1.79E-05 | |
| LINC00342 | 143.5960924 | 300.4029676 | 2.091999598 | 0.001672021 | 0.033045663 | |
| LINC00346 | 31.30032104 | 81.69159595 | 2.609928372 | 4.54E-05 | 0.002088899 | |
| LINC00402 | 116.0578587 | 43.01787352 | 0.370658859 | 1.84E-06 | 0.000155867 | |
| LINC00472 | 660.3719899 | 277.9330995 | 0.420873544 | 7.39E-05 | 0.003081782 | |
| LINC00521 | 1.946545399 | 20.47807236 | 10.52021308 | 0.000629807 | 0.016127975 | |
| LINC00910 | 71.92174456 | 146.0949074 | 2.031303722 | 0.000257005 | 0.008051218 | |
| LINC00963 | 415.5987416 | 868.0905516 | 2.088770886 | 2.07E-06 | 0.000169495 | |
| LINC01013 | 169.075582 | 18.56437036 | 0.10979924 | 0.000212616 | 0.006991141 | |

| Gene name | **Controls** normalized mean counts | **Simvastatin** normalized mean counts | **foldChange** | **pval** | **padj** | **Protein name** |
|---|---|---|---|---|---|---|
| LINC01031 | 115.8505994 | 17.83335697 | 0.153934093 | 0.000152062 | 0.005461934 | |
| LIPA | 917.3040544 | 1734.955414 | 1.891363508 | 0.000270935 | 0.008410893 | Lysosomal acid lipase/cholesteryl ester hydrolase |
| LMCD1 | 393.9823456 | 153.0764713 | 0.388536372 | 0.000159932 | 0.005666108 | cDNA FLJ52480, highly similar to LIM and cysteine-rich domains protein 1 |
| LMO2 | 45.44069338 | 128.439936 | 2.826539969 | 8.87E-07 | 8.52E-05 | Rhombotin-2 |
| LNPEP | 935.2799291 | 1554.487254 | 1.662055611 | 0.000712383 | 0.017685064 | Leucyl-cystinyl aminopeptidase |
| LONRF1 | 501.0997395 | 1035.912869 | 2.067278802 | 3.12E-06 | 0.000237672 | LON peptidase N-terminal domain and RING finger protein 1 |
| LPAR4 | 99.75043699 | 22.86954589 | 0.229267626 | 1.88E-08 | 3.28E-06 | Lysophosphatidic acid receptor 4 |
| LPIN1 | 1272.716481 | 2238.023887 | 1.75846225 | 0.000101203 | 0.003959769 | Phosphatidate phosphatase LPIN1 |
| LRCH2 | 103.9544993 | 35.5017306 | 0.341512208 | 6.39E-07 | 6.45E-05 | |
| LRP2 | 145.9078516 | 7.797628535 | 0.053442145 | 5.07E-14 | 3.36E-11 | Low-density lipoprotein receptor-related protein 2 |
| LRP4 | 3923.940392 | 1084.050378 | 0.276265761 | 1.15E-07 | 1.52E-05 | Low-density lipoprotein receptor-related protein 4 |
| LRP4-AS1 | 781.8119593 | 293.7331944 | 0.375708239 | 0.000356153 | 0.010441549 | |
| LRRC1 | 661.4503755 | 205.2491334 | 0.310301636 | 3.69E-05 | 0.001767285 | Leucine-rich repeat-containing protein 1 |
| LRRC14B | 90.81569604 | 392.8910213 | 4.326245775 | 6.73E-06 | 0.000453191 | |
| LRRC2 | 254.4269043 | 866.7448246 | 3.406655547 | 9.78E-09 | 1.81E-06 | |
| LRRC3B | 51.42600365 | 131.2252819 | 2.551730108 | 0.001964625 | 0.036769937 | Leucine-rich repeat-containing protein 3B |
| LRRC56 | 42.76258492 | 116.4376463 | 2.722886058 | 2.66E-05 | 0.001341819 | |
| LRRK2 | 183.4946699 | 357.6318557 | 1.949004055 | 0.000421099 | 0.012005173 | Leucine-rich repeat serine/threonine-protein kinase 2 |
| LRRN1 | 1105.492269 | 139.9441877 | 0.126589929 | 1.29E-05 | 0.000760788 | Leucine-rich repeat neuronal protein 1 |
| LSMEM1 | 27.0782662 | 83.36076536 | 3.078511924 | 2.43E-06 | 0.000193306 | Leucine-rich single-pass membrane protein 1 |
| LSS | 3898.702937 | 10067.29867 | 2.582217428 | 8.40E-10 | 2.14E-07 | Lanosterol synthase |
| LTBP4 | 474.4594123 | 1055.910601 | 2.225502484 | 0.001528908 | 0.031083803 | Latent-transforming growth factor beta-binding protein 4 |
| LTF | 7.946235064 | 48.64203178 | 6.121393514 | 6.44E-07 | 6.45E-05 | Kaliocin-1 |
| LTK | 35.92379452 | 623.9994997 | 17.37008877 | 5.79E-10 | 1.53E-07 | Tyrosine-protein kinase receptor |
| LUZP1 | 2248.56641 | 1301.564508 | 0.578841925 | 0.000289386 | 0.008851178 | Leucine zipper protein 1 |
| LYPD6 | 637.7109261 | 184.2873681 | 0.28898261 | 0.000280196 | 0.008612436 | Ly6/PLAUR domain-containing protein 6 |
| LYPD6B | 458.5270953 | 73.92214397 | 0.161216523 | 0.000969921 | 0.022086641 | Ly6/PLAUR domain-containing protein 6B |
| LYSMD4 | 267.8889698 | 536.0154992 | 2.000886784 | 5.34E-05 | 0.002401829 | LysM and putative peptidoglycan-binding domain-containing protein 4 |
| LYST | 419.6967662 | 876.201309 | 2.087700882 | 1.70E-06 | 0.000147375 | Lysosomal-trafficking regulator |
| LZTS3 | 307.7150127 | 621.1775491 | 2.018678074 | 8.27E-06 | 0.000535873 | Leucine zipper putative tumor suppressor 3 |
| MADD | 878.2524425 | 1639.54643 | 1.866828205 | 3.05E-05 | 0.001507072 | MAP kinase-activating death domain protein |
| MAFA | 201.3471211 | 585.014116 | 2.905500276 | 0.000582057 | 0.015266 | Transcription factor MafA |
| MAFG | 2117.711211 | 3484.526889 | 1.645421184 | 0.000655216 | 0.01654313 | Transcription factor MafG |

| Gene name | Controls normalized mean counts | Simvastatin normalized mean counts | foldChange | pval | padj | Protein name |
|---|---|---|---|---|---|---|
| MAGEL2 | 16.65731484 | 2.27918305 | 0.136827758 | 0.000168176 | 0.005864664 | MAGE-like protein 2 |
| MAK16 | 736.3445083 | 414.9137475 | 0.563477751 | 0.00051879 | 0.014044798 | Protein MAK16 homolog |
| MAL | 354.5493153 | 908.7082523 | 2.562995366 | 0.000254426 | 0.008010302 | Myelin and lymphocyte protein |
| MALL | 966.0273914 | 74.89304555 | 0.077526834 | 1.43E-06 | 0.000127259 | MAL-like protein |
| MAMDC2 | 14047.16357 | 3321.456608 | 0.23645034 | 3.01E-06 | 0.000232962 | MAM domain-containing protein 2 |
| MAML2 | 295.5949204 | 147.4263522 | 0.498744538 | 0.00025496 | 0.00801901 | Mastermind-like protein 2 |
| MAMLD1 | 688.334055 | 338.0559493 | 0.49112193 | 0.002614008 | 0.046273552 | Mastermind-like domain-containing protein 1 |
| MAP2K3 | 2311.178657 | 1293.055857 | 0.559478971 | 0.000110257 | 0.004234131 | Dual-specificity mitogen-activated protein kinase kinase 3 |
| MAP3K7CL | 8609.820515 | 3149.80098 | 0.36583817 | 1.04E-05 | 0.00063448 | MAP3K7 C-terminal-like protein |
| MAP3K8 | 131.9190209 | 410.7458722 | 3.113621292 | 5.70E-07 | 5.93E-05 | Mitogen-activated protein kinase kinase kinase 8 |
| MAP4 | 16361.10089 | 28677.30138 | 1.752773336 | 0.000116234 | 0.004408937 | Microtubule-associated protein |
| MAP4K1 | 103.4944119 | 185.1359876 | 1.788850086 | 0.00229991 | 0.042033486 | Mitogen-activated protein kinase kinase kinase kinase 1 |
| MAPRE3 | 8618.745064 | 5268.842513 | 0.611323629 | 0.00219892 | 0.04042529 | Microtubule-associated protein RP/EB family member 3 |
| MAPT | 66.40546712 | 148.1067958 | 2.230340396 | 0.000112558 | 0.004301247 | Microtubule-associated protein |
| MARCH1 | 210.2493458 | 110.8894618 | 0.52741882 | 0.001656378 | 0.032895175 | E3 ubiquitin-protein ligase MARCH1 |
| MARCH4 | 253.5646014 | 95.61608351 | 0.377087665 | 5.34E-05 | 0.002401829 | E3 ubiquitin-protein ligase MARCH4 |
| MARCKS | 2484.949796 | 1034.975447 | 0.416497528 | 1.24E-08 | 2.28E-06 | Myristoylated alanine-rich C-kinase substrate |
| MARCKSL1 | 2643.032742 | 1266.411756 | 0.47915099 | 2.21E-06 | 0.000180192 | MARCKS-related protein |
| MAS1 | 12.63171194 | 0.241435365 | 0.019113432 | 2.03E-06 | 0.000167251 | Proto-oncogene Mas |
| MAZ | 2213.743269 | 3429.580123 | 1.549222157 | 0.002882234 | 0.049725329 | Myc-associated zinc finger protein |
| MB | 13.65317073 | 58.20056607 | 4.26278754 | 1.18E-05 | 0.000702196 | Myoglobin |
| MBD3L2 | 8.085885422 | 0.255427941 | 0.031589359 | 0.000507187 | 0.013814846 | |
| MCAM | 2568.124064 | 867.2657872 | 0.337704007 | 1.46E-06 | 0.000128946 | Cell surface glycoprotein MUC18 |
| MCF2 | 74.53001326 | 11.93785207 | 0.160175097 | 2.61E-12 | 1.14E-09 | DBL-transforming protein |
| MCF2L2 | 143.1491939 | 308.1043305 | 2.152330181 | 2.35E-05 | 0.001217979 | Probable guanine nucleotide exchange factor MCF2L2 |
| MDFIC | 772.0590584 | 1232.36987 | 1.596211918 | 0.001909114 | 0.036117555 | MyoD family inhibitor domain-containing protein |
| MED12L | 64.58725105 | 22.13186508 | 0.342666157 | 0.000125032 | 0.004652209 | Mediator of RNA polymerase II transcription subunit 12-like protein |
| MEF2A | 3755.907928 | 10669.84162 | 2.840815542 | 1.72E-12 | 8.10E-10 | Myocyte-specific enhancer factor 2A |
| MESDC2 | 1392.291391 | 844.9301901 | 0.606863043 | 0.001128862 | 0.0247983 | LDLR chaperone MESD |
| MEST | 276.2030051 | 29.83345558 | 0.108012784 | 1.83E-12 | 8.44E-10 | Mesoderm-specific transcript homolog protein |
| METRNL | 493.0148593 | 973.2667714 | 1.974112449 | 9.65E-05 | 0.003818538 | Meteorin-like protein |
| METTL1 | 386.7230281 | 225.356253 | 0.582732955 | 0.000937787 | 0.021608011 | tRNA (guanine-N(7)-)-methyltransferase |
| METTL7A | 521.1219362 | 1363.140521 | 2.615780351 | 7.77E-06 | 0.000512347 | Methyltransferase-like protein 7A |

| Gene name | **Controls** normalized mean counts | **Simvastatin** normalized mean counts | foldChange | pval | padj | Protein name |
|---|---|---|---|---|---|---|
| METTL7B | 345.4276176 | 82.76633368 | 0.239605432 | 1.31E-06 | 0.000118749 | Methyltransferase-like protein 7B |
| MEX3A | 515.8876263 | 182.2155301 | 0.353207793 | 1.02E-05 | 0.000624186 | RNA-binding protein MEX3A |
| MEX3B | 472.2203378 | 278.3728988 | 0.589497903 | 0.001940865 | 0.036478876 | RNA-binding protein MEX3B |
| MFAP3L | 968.580165 | 366.790161 | 0.378688491 | 5.42E-06 | 0.000383286 | Microfibrillar-associated protein 3-like |
| MFAP5 | 9561.937736 | 2925.466687 | 0.305949146 | 8.86E-06 | 0.000567374 | cDNA FLJ42377 fis, clone UTERU2035469, highly similar to Microfibrillar-associated protein 5 |
| MFI2 | 49.43473915 | 174.181859 | 3.523470782 | 1.82E-05 | 0.000992178 | Melanotransferrin |
| MFNG | 0.981038388 | 11.17622642 | 11.39224169 | 0.001761568 | 0.033956313 | Beta-1,3-N-acetylglucosaminyltransferase manic fringe |
| MFRP_2 | 210.8647435 | 97.82296561 | 0.463913331 | 0.000129598 | 0.004784586 | |
| MGC4294 | 10.84119021 | 0.720694587 | 0.066477441 | 0.000174447 | 0.006049427 | |
| MGST3 | 655.9758476 | 1322.894406 | 2.016681575 | 5.11E-06 | 0.000365252 | Microsomal glutathione S-transferase 3 |
| MID1 | 472.6060515 | 144.1013047 | 0.304907871 | 4.61E-12 | 1.97E-09 | E3 ubiquitin-protein ligase Midline-1 |
| MINA | 964.6964787 | 599.8818049 | 0.621834762 | 0.002125171 | 0.039348549 | Bifunctional lysine-specific demethylase and histidyl-hydroxylase MINA |
| MIR143HG | 7.236843164 | 0.255427941 | 0.035295492 | 0.001143429 | 0.025042602 | |
| MIR181A2HG | 12.42290498 | 1.498390926 | 0.12061518 | 0.000123332 | 0.004594433 | |
| MIR24-2 | 414.7915512 | 779.4461197 | 1.879127281 | 6.05E-05 | 0.002649302 | |
| MIR503HG | 1485.170263 | 523.0568439 | 0.352186451 | 5.61E-06 | 0.000393764 | |
| MKI67 | 45.25042774 | 1.270948138 | 0.028086986 | 3.30E-13 | 1.89E-10 | Antigen KI-67 |
| MKI67IP | 1102.372891 | 580.0623104 | 0.526194281 | 3.56E-05 | 0.001716173 | |
| MKNK2 | 943.7134293 | 2068.510026 | 2.191883639 | 1.47E-07 | 1.88E-05 | cDNA FLJ53467, highly similar to MAP kinase-interactingserine/threonine-protein kinase 2 (EC 2.7.11.1) |
| MLLT11 | 1948.877744 | 1067.810563 | 0.547910492 | 0.001297953 | 0.027465386 | Protein AF1q |
| MLLT3 | 987.4002462 | 509.7633858 | 0.516268239 | 0.000441344 | 0.012423943 | Protein AF-9 |
| MLXIPL | 49.96304093 | 287.3442822 | 5.751136777 | 0.000149742 | 0.005396328 | Carbohydrate-responsive element-binding protein |
| MMP11 | 2792.928953 | 1087.825934 | 0.389492877 | 1.00E-07 | 1.35E-05 | Stromelysin-3 |
| MMP19 | 230.8127736 | 87.77632436 | 0.380292316 | 8.32E-08 | 1.17E-05 | Matrix metalloproteinase-19 |
| MN1 | 100.499008 | 245.559935 | 2.443406556 | 0.000136119 | 0.004981314 | Probable tumor suppressor protein MN1 |
| MNT | 463.8297468 | 738.2867825 | 1.591719349 | 0.002896033 | 0.049897172 | Max-binding protein MNT |
| MOK | 350.2157674 | 157.7516926 | 0.450441434 | 5.74E-05 | 0.002534507 | MAPK/MAK/MRK overlapping kinase |
| MPC1 | 1009.236686 | 1869.988352 | 1.852873938 | 4.37E-05 | 0.002026798 | Mitochondrial pyruvate carrier 1 |
| MPDU1 | 1053.769916 | 580.4048216 | 0.550788946 | 0.000122802 | 0.004580181 | Mannose-P-dolichol utilization defect 1 protein |
| MPHOSPH6 | 481.1822235 | 279.4356525 | 0.580727298 | 0.000705558 | 0.017585799 | M-phase phosphoprotein 6 |
| MPZL1 | 1422.654082 | 876.5507166 | 0.616137632 | 0.001312402 | 0.027658251 | Myelin protein zero-like protein 1 |
| MRAP2 | 146.038242 | 547.1441161 | 3.746581091 | 0.000305473 | 0.009207397 | Melanocortin-2 receptor accessory protein 2 |
| MRC2 | 6255.600493 | 3403.211107 | 0.544026287 | 0.000383926 | 0.011109198 | C-type mannose receptor 2 |

| Gene name | **Controls** normalized mean counts | **Simvastatin** normalized mean counts | **foldChange** | **pval** | **padj** | **Protein name** |
|---|---|---|---|---|---|---|
| MRE11A | 395.2649819 | 238.2575772 | 0.602779371 | 0.002334218 | 0.042436102 | cDNA FLJ38069 fis, clone CTONG2015434, highly similar to DOUBLE-STRAND BREAK REPAIR PROTEIN MRE11A |
| MRGPRF | 24.63354574 | 115.6710914 | 4.69567364 | 9.85E-11 | 3.22E-08 | Mas-related G-protein coupled receptor member F |
| MROH1 | 415.503739 | 820.5169298 | 1.974752217 | 1.18E-05 | 0.000703609 | |
| MROH5 | 5.317509053 | 47.05819979 | 8.849669897 | 2.36E-08 | 4.02E-06 | |
| MRPL12 | 1055.151369 | 578.3791384 | 0.548148025 | 0.000334774 | 0.009917615 | 39S ribosomal protein L12, mitochondrial |
| MRPL19 | 1289.309426 | 775.7980461 | 0.60171595 | 0.000780573 | 0.018865963 | 39S ribosomal protein L19, mitochondrial |
| MRPL32 | 1087.819741 | 685.3075614 | 0.629982648 | 0.002502355 | 0.044632028 | cDNA FLJ33232 fis, clone ASTRO2002024, highly similar to 39S ribosomal protein L32, mitochondrial |
| MRPL42 | 674.2585739 | 415.8023156 | 0.616680798 | 0.001981383 | 0.037016759 | Mitochondrial ribosomal protein L42, isoform CRA_e |
| MRPS12 | 473.8926309 | 282.6371334 | 0.59641597 | 0.001228729 | 0.026506209 | 28S ribosomal protein S12, mitochondrial |
| MRPS17_2 | 78.99043952 | 33.84835002 | 0.428511985 | 0.000788708 | 0.019004093 | |
| MRTO4 | 1128.500503 | 680.068894 | 0.602630563 | 0.000920306 | 0.02135038 | mRNA turnover protein 4 homolog |
| MRVI1 | 4.821588361 | 20.67942993 | 4.288924806 | 0.001045194 | 0.023372967 | Protein MRVI1 |
| MRVI1-AS1 | 6.541763974 | 29.23152482 | 4.468446879 | 0.00011651 | 0.004408937 | |
| MSMO1 | 3306.127494 | 7557.263706 | 2.285835534 | 1.01E-05 | 0.00061844 | Methylsterol monooxygenase 1 |
| MST1 | 141.9846606 | 280.1464667 | 1.973075581 | 0.000410861 | 0.011757356 | Hepatocyte growth factor-like protein alpha chain |
| MSTN | 465.5080853 | 56.80867168 | 0.122035843 | 4.03E-13 | 2.24E-10 | Growth/differentiation factor 8 |
| MTG1 | 770.3185735 | 447.4658859 | 0.580884197 | 0.000515704 | 0.013973422 | Mitochondrial GTPase 1 |
| MTHFR | 1250.711495 | 2013.194622 | 1.609639497 | 0.001590158 | 0.032017489 | Methylenetetrahydrofolate reductase |
| MTMR11 | 28.25932251 | 75.17644688 | 2.660235285 | 0.001284325 | 0.027232598 | Myotubularin-related protein 11 |
| MTMR12 | 1917.44521 | 950.907011 | 0.495923955 | 0.002444489 | 0.043927362 | Myotubularin-related protein 12 |
| MT-RNR1 | 9570.634021 | 19739.45769 | 2.062502615 | 0.001328991 | 0.027932156 | |
| MTSS1 | 424.3381668 | 1043.997354 | 2.46029567 | 5.68E-09 | 1.16E-06 | Metastasis suppressor protein 1 |
| MUSTN1 | 812.4538574 | 1989.686796 | 2.448984368 | 9.70E-05 | 0.003824035 | Musculoskeletal embryonic nuclear protein 1 |
| MVD | 2080.057217 | 7443.745722 | 3.578625463 | 3.21E-13 | 1.89E-10 | Diphosphomevalonate decarboxylase |
| MX1 | 17.81924293 | 77.64243649 | 4.357224198 | 0.002532714 | 0.045070139 | Interferon-induced GTP-binding protein Mx1, N-terminally processed |
| MXD3 | 263.2051959 | 493.9570988 | 1.876699649 | 0.000190582 | 0.006450762 | Max dimerization protein 3 |
| MXD4 | 1057.462346 | 1852.15985 | 1.75151376 | 0.000963579 | 0.021990524 | Max dimerization protein 4 |
| MXI1 | 508.9604747 | 937.4610249 | 1.841913216 | 6.07E-05 | 0.00265329 | cDNA FLJ61042, highly similar to MAX-interacting protein 1 |
| MXRA5 | 680.4146337 | 218.9709903 | 0.321819931 | 1.64E-05 | 0.000931238 | Matrix-remodeling-associated protein 5 |
| MYCBP | 303.1657778 | 141.6781672 | 0.467329025 | 1.14E-05 | 0.000686735 | C-Myc-binding protein |

| Gene name | Controls normalized mean counts | Simvastatin normalized mean counts | foldChange | pval | padj | Protein name |
|---|---|---|---|---|---|---|
| MYEF2 | 855.5142044 | 413.2651099 | 0.483060489 | 4.82E-05 | 0.002191842 | Myelin expression factor 2 |
| MYF6 | 483.3780805 | 959.1836468 | 1.984334179 | 0.000276958 | 0.008538222 | Myogenic factor 6 |
| MYH1 | 5137.591929 | 911.4674855 | 0.177411421 | 0.000330907 | 0.009815591 | Myosin-1 |
| MYH7B | 416.2801875 | 2771.458685 | 6.657676172 | 7.30E-29 | 2.27E-25 | Myosin-7B |
| MYL10 | 14.79649946 | 2.006151024 | 0.135582813 | 0.000396607 | 0.011412445 | Myosin regulatory light chain 10 |
| MYL2 | 2642.479177 | 7853.045536 | 2.971847651 | 0.000101084 | 0.003959769 | Myosin regulatory light chain 2, ventricular/cardiac muscle isoform |
| MYL5 | 719.8869253 | 2099.564986 | 2.916520515 | 7.78E-05 | 0.003202785 | Myosin light chain 5 |
| MYL9 | 11909.85338 | 6121.8475 | 0.514015354 | 0.000166544 | 0.005833912 | Myosin regulatory light polypeptide 9 |
| MYO15B | 87.03574383 | 255.4474845 | 2.934972154 | 0.002488004 | 0.044445622 | Unconventional myosin-XVB |
| MYO1B | 6484.64001 | 3648.050622 | 0.562567948 | 0.000109179 | 0.004203106 | Unconventional myosin-Ib |
| MYO1D | 1899.18313 | 3283.401448 | 1.728849312 | 0.000610284 | 0.015727839 | Unconventional myosin-Id |
| MYO6 | 488.2405159 | 810.8983811 | 1.660858439 | 0.00097407 | 0.022164903 | Unconventional myosin-VI |
| MYOM2 | 1469.014932 | 4723.909318 | 3.21569864 | 5.38E-06 | 0.000382217 | Myomesin-2 |
| MYOM3 | 6847.513885 | 11489.4763 | 1.677904783 | 0.001910029 | 0.036117555 | Myomesin-3 |
| N4BP2L1 | 47.64897461 | 161.4374116 | 3.388056362 | 5.44E-09 | 1.12E-06 | |
| N4BP3 | 909.747025 | 458.6717967 | 0.504175099 | 0.001533841 | 0.031154372 | NEDD4-binding protein 3 |
| NACC1 | 2019.590892 | 1264.765179 | 0.626248209 | 0.002101731 | 0.038960929 | Nucleus accumbens-associated protein 1 |
| NAMA | 7.315374906 | 54.11483513 | 7.397411044 | 3.03E-05 | 0.00150313 | |
| NAMPT | 965.8548302 | 1813.901724 | 1.878027284 | 2.16E-05 | 0.001141705 | Nicotinamide phosphoribosyltransferase |
| NAP1L3 | 258.2322169 | 134.7894933 | 0.521970089 | 0.00118141 | 0.025644755 | Nucleosome assembly protein 1-like 3 |
| NAV2 | 3367.27316 | 1239.682664 | 0.368156252 | 0.000345007 | 0.010153073 | Neuron navigator 2 |
| NAV3 | 6187.397623 | 1894.830449 | 0.306240291 | 9.14E-10 | 2.31E-07 | Neuron navigator 3 |
| NBEAL2 | 159.2046084 | 311.9540182 | 1.959453444 | 7.95E-05 | 0.003263819 | Neurobeachin-like protein 2 |
| NCBP1 | 1943.187805 | 1222.938433 | 0.629346494 | 0.00241578 | 0.043512008 | Nuclear cap-binding protein subunit 1 |
| NCBP2 | 3320.023797 | 1973.509696 | 0.594426371 | 0.00048199 | 0.013244494 | Nuclear cap binding protein subunit 2, 20kDa, isoform CRA_a |
| NCLN | 2040.552922 | 1155.92272 | 0.566475247 | 0.000182249 | 0.006243417 | Nicalin |
| NCR3LG1 | 133.3515239 | 236.2299265 | 1.771482766 | 0.000909826 | 0.021199286 | Natural cytotoxicity triggering receptor 3 ligand 1 |
| NDRG1 | 487.7215915 | 1419.18473 | 2.909825513 | 2.52E-10 | 7.32E-08 | Protein NDRG1 |
| NDRG2 | 728.1573374 | 2549.520356 | 3.501331683 | 0.000371478 | 0.010799256 | cDNA FLJ55190, highly similar to Protein NDRG2 |
| NDUFB8_2 | 605.9936917 | 256.0861754 | 0.422588847 | 0.000369538 | 0.010762937 | |
| NEB | 38060.69027 | 77373.19691 | 2.032890007 | 1.33E-05 | 0.000783169 | Nebulin |
| NEBL | 36.69280959 | 14.68952478 | 0.400337967 | 0.001668711 | 0.033019681 | Nebulette |
| NEDD4 | 1334.477175 | 715.1089083 | 0.535871967 | 8.38E-05 | 0.003399083 | E3 ubiquitin-protein ligase |
| NEIL1 | 91.89913573 | 177.2379629 | 1.928614034 | 0.000428089 | 0.012160845 | Endonuclease 8-like 1 |
| NEK6 | 2120.545368 | 993.8311096 | 0.468667695 | 7.91E-06 | 0.000513507 | Serine/threonine-protein kinase Nek6 |
| NETO2 | 123.0491722 | 64.38097461 | 0.523213391 | 0.001960154 | 0.03670834 | Neuropilin and tolloid-like protein 2 |
| NEURL1B | 231.6446939 | 84.2875663 | 0.363865733 | 3.79E-05 | 0.001807613 | E3 ubiquitin-protein ligase NEURL1B |
| NEUROD2 | 20.00821884 | 1.249154554 | 0.062432072 | 0.000112472 | 0.004301247 | Neurogenic differentiation factor 2 |
| NFASC | 1505.592694 | 592.0654496 | 0.393244104 | 2.58E-08 | 4.36E-06 | Neurofascin |
| NFE2L1 | 20153.03318 | 36735.80049 | 1.822842258 | 3.89E-05 | 0.001851591 | Nuclear factor erythroid 2-related factor 1 |

| Gene name | Controls normalized mean counts | Simvastatin normalized mean counts | foldChange | pval | padj | Protein name |
|---|---|---|---|---|---|---|
| NFIL3 | 221.9758875 | 659.764979 | 2.97223715 | 1.62E-11 | 6.22E-09 | Nuclear factor interleukin-3-regulated protein |
| NGEF | 42.60545936 | 422.7967169 | 9.92353382 | 9.35E-22 | 1.82E-18 | cDNA FLJ59455, highly similar to Homo sapiens neuronal guanine nucleotide exchange factor (NGEF), mRNA |
| NGFR | 5380.615389 | 693.8336526 | 0.128950613 | 2.95E-06 | 0.000229608 | cDNA FLJ51424, highly similar to Tumor necrosis factor receptor superfamilymember 16 |
| NGFRAP1 | 2374.616589 | 1340.054043 | 0.564324384 | 0.000108964 | 0.00420174 | Protein BEX3 |
| NHLH2 | 155.8878969 | 48.19487126 | 0.30916365 | 2.84E-06 | 0.000223493 | Helix-loop-helix protein 2 |
| NHLRC3 | 339.8347814 | 553.9438726 | 1.63003878 | 0.001944884 | 0.036509661 | NHL repeat-containing protein 3 |
| NHP2 | 1422.054728 | 888.5462574 | 0.624832673 | 0.001923106 | 0.036254624 | H/ACA ribonucleoprotein complex subunit 2 |
| NHSL1 | 633.2476262 | 198.2518203 | 0.313071557 | 1.41E-10 | 4.48E-08 | |
| NINJ1 | 286.5098061 | 652.1673077 | 2.276247772 | 0.000395299 | 0.011385346 | Ninjurin-1 |
| NIPSNAP1 | 372.5996318 | 182.3095715 | 0.489290799 | 1.88E-05 | 0.00101754 | Protein NipSnap homolog 1 |
| NKX3-2 | 194.8879345 | 29.43314748 | 0.151026012 | 1.09E-10 | 3.51E-08 | Homeobox protein Nkx-3.2 |
| NME1 | 1293.407005 | 344.5620532 | 0.266398784 | 1.90E-12 | 8.59E-10 | Nucleoside diphosphate kinase A |
| NMNAT3 | 74.98054051 | 145.7177649 | 1.943407768 | 0.000684732 | 0.017149167 | Nicotinamide nucleotide adenylyltransferase 3, isoform CRA_g |
| NMRK2 | 177.5629575 | 489.1824544 | 2.754980326 | 0.000243028 | 0.007729689 | Nicotinamide riboside kinase 2 |
| NOL6 | 1682.734489 | 1066.434568 | 0.633750942 | 0.002804078 | 0.048837435 | Nucleolar protein 6 |
| NOL9 | 767.5386223 | 396.1839812 | 0.516174652 | 5.31E-05 | 0.002395782 | Polynucleotide 5'-hydroxyl-kinase NOL9 |
| NOP16 | 651.3349284 | 322.3830112 | 0.494957352 | 1.08E-05 | 0.00065663 | Nucleolar protein 16 |
| NOP2 | 1188.827905 | 745.1886232 | 0.626826322 | 0.002594159 | 0.045979429 | Putative ribosomal RNA methyltransferase NOP2 |
| NOP58 | 1030.087602 | 616.337904 | 0.598335426 | 0.000864618 | 0.020359426 | Nucleolar protein 58 |
| NOS1AP | 63.13243094 | 16.21955224 | 0.256913159 | 0.000224061 | 0.007252482 | Carboxyl-terminal PDZ ligand of neuronal nitric oxide synthase protein |
| NOX4 | 17.99084839 | 4.608612062 | 0.256164243 | 0.001646963 | 0.032818991 | NADPH oxidase 4 |
| NOXA1 | 81.36120159 | 199.9191276 | 2.457180126 | 1.37E-06 | 0.000122964 | NADPH oxidase activator 1 |
| NPAS1 | 1145.234387 | 272.6648379 | 0.238086492 | 0.000247641 | 0.007852321 | Neuronal PAS domain-containing protein 1 |
| NPC1 | 1220.565708 | 1989.345062 | 1.629854952 | 0.000944229 | 0.021708188 | cDNA FLJ51802, highly similar to Niemann-Pick C1 protein |
| NPHP3-AS1 | 121.3928716 | 23.58262483 | 0.194266966 | 2.97E-07 | 3.40E-05 | |
| NPIPA3 | 43.54996929 | 110.4308086 | 2.535726439 | 0.001709652 | 0.033425784 | |
| NPM1 | 13551.93348 | 8615.924565 | 0.635770872 | 0.002213076 | 0.040613531 | Nucleophosmin |
| NPM3 | 320.4139848 | 159.6655272 | 0.498310107 | 4.28E-05 | 0.001999846 | Nucleoplasmin-3 |
| NPPA | 68.50681376 | 18.92470126 | 0.276245533 | 2.93E-05 | 0.001458266 | Natriuretic peptides A |
| NPR3 | 189.5099471 | 31.17761817 | 0.164517054 | 9.49E-06 | 0.000597441 | Atrial natriuretic peptide receptor 3 |
| NPTX2 | 158.1904257 | 318.8397533 | 2.015543936 | 0.000558713 | 0.014828766 | Neuronal pentraxin-2 |
| NPY6R | 176.8482703 | 75.9439745 | 0.429430123 | 0.001182349 | 0.02564724 | Putative neuropeptide Y receptor type 6 |
| NQO1 | 3311.502834 | 7409.738473 | 2.237575761 | 4.54E-08 | 7.06E-06 | NAD(P)H dehydrogenase, quinone 1, isoform CRA_a |

| Gene name | Controls normalized mean counts | Simvastatin normalized mean counts | foldChange | pval | padj | Protein name |
|---|---|---|---|---|---|---|
| NR1D1 | 380.0511106 | 813.0457357 | 2.139306301 | 1.65E-05 | 0.00093288 | Nuclear receptor subfamily 1 group D member 1 |
| NR3C1 | 1177.415215 | 2546.518023 | 2.162803734 | 2.34E-07 | 2.77E-05 | Glucocorticoid receptor |
| NR4A1 | 246.4103569 | 1331.223721 | 5.402466594 | 1.11E-11 | 4.50E-09 | Nuclear receptor subfamily 4 group A member 1 |
| NR4A2 | 59.4104476 | 177.6722224 | 2.990588854 | 3.74E-08 | 6.03E-06 | Nuclear receptor subfamily 4 group A member 2 |
| NRAP | 514.5336999 | 2763.733077 | 5.371335399 | 1.90E-16 | 1.90E-13 | Nebulin-related-anchoring protein |
| NSG1 | 118.3173261 | 12.10255499 | 0.102288949 | 0.000167274 | 0.005842717 | Neuron-specific protein family member 1 |
| NT5M | 46.64741052 | 98.59226982 | 2.113563619 | 0.000589131 | 0.015412532 | 5'(3')-deoxyribonucleotidase, mitochondrial |
| NTAN1 | 599.9459561 | 1057.86631 | 1.76326934 | 0.000198084 | 0.006600877 | Protein N-terminal asparagine amidohydrolase |
| NTF4 | 6.893663113 | 29.24758028 | 4.24267618 | 0.000159141 | 0.005648091 | Neurotrophin-4 |
| NTM | 605.978823 | 242.0049159 | 0.399362002 | 1.35E-06 | 0.000121543 | Neurotrimin |
| NTN4 | 1043.41432 | 2769.922543 | 2.654671774 | 1.06E-07 | 1.43E-05 | Netrin-4 |
| NTNG2 | 153.2150224 | 373.4518917 | 2.437436525 | 0.000952131 | 0.021847765 | Netrin-G2 |
| NUAK1 | 5115.359344 | 2338.03379 | 0.457061495 | 0.000219001 | 0.007133255 | NUAK family SNF1-like kinase 1 |
| NUDT11 | 280.6898769 | 143.7889561 | 0.512269832 | 7.76E-05 | 0.003195127 | Diphosphoinositol polyphosphate phosphohydrolase 3-beta |
| NUDT8 | 29.08638616 | 84.14838223 | 2.893050439 | 4.42E-06 | 0.000319792 | Nucleoside diphosphate-linked moiety X motif 8, mitochondrial |
| NUP93 | 2787.524916 | 1321.994385 | 0.474253836 | 1.68E-06 | 0.000146075 | Nuclear pore complex protein Nup93 |
| NUPR1 | 1371.642674 | 2683.042186 | 1.956079552 | 5.53E-05 | 0.002472991 | Nuclear protein 1 |
| OASL | 0.994408343 | 22.29797821 | 22.42336196 | 2.82E-07 | 3.25E-05 | 2'-5'-oligoadenylate synthase-like protein |
| OBSCN | 7774.852612 | 13186.95602 | 1.696103667 | 0.000546488 | 0.014583891 | Obscurin |
| OCEL1 | 69.5258455 | 130.4255602 | 1.87592915 | 0.001475265 | 0.030190519 | |
| OLFM4 | 12.81959777 | 2.023755108 | 0.157864166 | 0.000603896 | 0.015678675 | Olfactomedin-4 |
| OLFML2B | 2213.565778 | 1262.739999 | 0.570455151 | 0.001750093 | 0.033833676 | Olfactomedin-like protein 2B |
| OPN3 | 2423.908949 | 755.3512381 | 0.311625252 | 0.000250163 | 0.007900081 | Opsin-3 |
| OR7E18P | 9.496966642 | 0.283928817 | 0.02989679 | 0.000108748 | 0.00420174 | |
| OR7E19P | 321.8126601 | 28.46722639 | 0.088459001 | 1.93E-05 | 0.001043356 | |
| OR7E38P | 214.4515134 | 111.6986961 | 0.520857579 | 0.000558548 | 0.014828766 | |
| ORAI3 | 220.8493148 | 422.2632294 | 1.91199701 | 6.64E-05 | 0.002821288 | Protein orai-3 |
| OVCA2 | 279.9429517 | 162.8943948 | 0.581884251 | 0.001358465 | 0.028401071 | Ovarian cancer-associated gene 2 protein |
| OVGP1 | 28.65302737 | 72.46557924 | 2.529072349 | 0.000127709 | 0.004734836 | Oviduct-specific glycoprotein |
| P2RX1 | 148.0923042 | 282.849879 | 1.909956636 | 0.002183541 | 0.040213867 | P2X purinoceptor 1 |
| P2RX7 | 11.11106915 | 35.503051 | 3.195286658 | 0.000713463 | 0.017697746 | P2X purinoceptor 7 |
| PABPC1L | 446.5733242 | 271.4681891 | 0.607891637 | 0.001781296 | 0.03416091 | Polyadenylate-binding protein 1-like |
| PACSIN1 | 519.9591586 | 124.3737629 | 0.2391991 | 0.000702271 | 0.017521859 | Protein kinase C and casein kinase substrate in neurons protein 1 |
| PAICS | 1839.715731 | 1172.367138 | 0.637254506 | 0.00282326 | 0.049042798 | Phosphoribosylaminoimidazole carboxylase |

| Gene name | **Controls** normalized mean counts | **Simvastatin** normalized mean counts | foldChange | pval | padj | Protein name |
|---|---|---|---|---|---|---|
| PAK1IP1 | 507.0630913 | 296.8436582 | 0.5854176 | 0.001747953 | 0.033813319 | p21-activated protein kinase-interacting protein 1 |
| PAK7 | 49.09712688 | 3.7490731 | 0.076360336 | 1.84E-12 | 8.44E-10 | Serine/threonine-protein kinase PAK 7 |
| PALM | 1425.917626 | 2419.865175 | 1.697058183 | 0.001470443 | 0.030111653 | Paralemmin-1 |
| PALM2 | 715.3016909 | 230.951862 | 0.322873362 | 9.28E-06 | 0.00058816 | Paralemmin-2 |
| PANK1 | 272.5325288 | 446.1897761 | 1.637198239 | 0.001725032 | 0.033540427 | Pantothenate kinase 1 |
| PAQR6 | 59.56879354 | 139.279527 | 2.338129056 | 2.75E-05 | 0.001382214 | Progestin and adipoQ receptor family member 6 |
| PAQR8 | 36.58294709 | 161.5404249 | 4.415730217 | 3.99E-10 | 1.11E-07 | Membrane progestin receptor beta |
| PARD6G | 180.9586412 | 90.4430899 | 0.499799785 | 0.000168956 | 0.005885261 | Partitioning defective 6 homolog gamma |
| PARK2 | 134.1545823 | 271.5095001 | 2.023855581 | 0.000182917 | 0.006245683 | E3 ubiquitin-protein ligase parkin |
| PARM1 | 757.1073575 | 252.9655374 | 0.334121093 | 0.001235815 | 0.026547825 | Prostate androgen-regulated mucin-like protein 1 |
| PARP1 | 3409.915029 | 1936.484143 | 0.567898064 | 0.000146446 | 0.005303072 | Poly [ADP-ribose] polymerase 1 |
| PARP12 | 86.59093374 | 201.8955303 | 2.331601261 | 4.34E-05 | 0.002022043 | Poly [ADP-ribose] polymerase 12 |
| PARP4 | 1376.094707 | 2221.944769 | 1.614674308 | 0.001332313 | 0.027945337 | Poly [ADP-ribose] polymerase 4 |
| PASK | 50.98372616 | 103.2341815 | 2.024845755 | 0.000747812 | 0.018344996 | PAS domain-containing serine/threonine-protein kinase |
| PAWR | 552.9655159 | 250.6867599 | 0.453349717 | 0.002234304 | 0.040954777 | PRKC apoptosis WT1 regulator protein |
| PCAT1 | 50.03901443 | 14.63257152 | 0.292423256 | 0.000158618 | 0.005638817 | |
| PCBP3 | 193.7178867 | 401.0939355 | 2.070505427 | 5.88E-05 | 0.002585401 | Poly(rC)-binding protein 3 |
| PCDH19 | 717.2332999 | 301.5330817 | 0.420411436 | 1.25E-06 | 0.000114365 | Protocadherin-19 |
| PCDH7 | 1086.150668 | 423.163997 | 0.389599721 | 1.48E-06 | 0.000130054 | Protocadherin-7 |
| PCDHGA2 | 9.262781788 | 40.70635849 | 4.39461486 | 0.000383771 | 0.011109198 | Protocadherin gamma-A2 |
| PCK2 | 430.5271461 | 733.5820657 | 1.703915937 | 0.000586224 | 0.015349392 | cDNA FLJ50710, highly similar to Phosphoenolpyruvate carboxykinase (GTP), mitochondrial (EC 4.1.1.32) |
| PCNT | 1072.150429 | 1954.045357 | 1.822547754 | 7.27E-05 | 0.003041939 | Pericentrin |
| PCSK4 | 39.99995393 | 115.409895 | 2.885250699 | 3.79E-06 | 0.000279111 | Proprotein convertase subtilisin/kexin type 4 |
| PCYT2 | 913.7058326 | 1953.919115 | 2.138455338 | 3.50E-07 | 3.86E-05 | Ethanolamine-phosphate cytidylyltransferase |
| PDCD1LG2 | 69.16514394 | 21.05781837 | 0.304457089 | 6.46E-05 | 0.002767772 | Programmed cell death 1 ligand 2 |
| PDE3A | 1006.181435 | 12705.39353 | 12.62733846 | 6.03E-50 | 3.13E-46 | cGMP-inhibited 3',5'-cyclic phosphodiesterase A |
| PDE4B | 21.62574222 | 53.01456413 | 2.451456398 | 0.000929004 | 0.021508601 | cAMP-specific 3',5'-cyclic phosphodiesterase 4B |
| PDF | 127.9275196 | 51.43143917 | 0.402035773 | 9.24E-05 | 0.003688584 | Peptide deformylase, mitochondrial |
| PDGFRB | 2854.388641 | 1726.446492 | 0.604839322 | 0.001722712 | 0.033540427 | Platelet-derived growth factor receptor beta |
| PDIA3 | 9577.068612 | 5562.746332 | 0.580840188 | 0.002522037 | 0.044931554 | Protein disulfide-isomerase A3 |
| PDK4 | 57.92583053 | 271.3232118 | 4.683976204 | 1.91E-06 | 0.000158586 | [Pyruvate dehydrogenase (acetyl-transferring)] kinase isozyme 4, mitochondrial |

| Gene name | Controls normalized mean counts | Simvastatin normalized mean counts | foldChange | pval | padj | Protein name |
|---|---|---|---|---|---|---|
| PDLIM2 | 1147.956698 | 466.8799282 | 0.406705174 | 2.48E-05 | 0.001266873 | PDZ and LIM domain protein 2 |
| PDLIM3 | 16496.03051 | 29205.10988 | 1.770432582 | 0.000149888 | 0.005396328 | PDZ and LIM domain protein 3 |
| PDLIM4 | 2649.558656 | 1610.692979 | 0.607909916 | 0.000976715 | 0.022208837 | PDZ and LIM domain protein 4 |
| PDPN | 600.0554246 | 1329.697131 | 2.215957187 | 3.52E-06 | 0.000263537 | Podoplanin |
| PDSS1 | 127.8442878 | 70.79865809 | 0.553788201 | 0.002539499 | 0.045162307 | Decaprenyl-diphosphate synthase subunit 1 |
| PDZK1 | 34.38146066 | 267.1764331 | 7.770944805 | 1.15E-19 | 1.79E-16 | Na(+)/H(+) exchange regulatory cofactor NHE-RF3 |
| PEA15 | 23321.11514 | 14120.57617 | 0.605484604 | 0.001582964 | 0.03193235 | cDNA FLJ38560 fis, clone HCHON2003642, highly similar to Astrocytic phosphoprotein PEA-15 |
| PEBP4 | 9.283510866 | 32.12197963 | 3.460111168 | 0.000566513 | 0.01495921 | Phosphatidylethanolamine-binding protein 4 |
| PEG10 | 2253.344831 | 5427.845442 | 2.408794858 | 2.34E-08 | 4.00E-06 | Retrotransposon-derived protein PEG10 |
| PELI2 | 142.2768361 | 246.196189 | 1.730402473 | 0.001340048 | 0.028069712 | E3 ubiquitin-protein ligase pellino homolog 2 |
| PFAS | 483.7077583 | 282.5770743 | 0.584189667 | 0.000929324 | 0.021508601 | Phosphoribosylformylglycinamidine synthase |
| PGAM1 | 5943.001696 | 3752.69671 | 0.63144803 | 0.001831031 | 0.03489955 | Phosphoglycerate mutase 1 |
| PGAM2 | 315.5443955 | 958.2524653 | 3.036822961 | 0.002381359 | 0.043016573 | Phosphoglycerate mutase 2 |
| PGM5 | 2208.026526 | 901.9070504 | 0.408467489 | 0.000166163 | 0.005827147 | Phosphoglucomutase-like protein 5 |
| PHLDA1 | 3102.812652 | 1936.87365 | 0.624231582 | 0.002116387 | 0.039209249 | Pleckstrin homology-like domain family A member 1 |
| PHLDA3 | 3558.982005 | 6293.707262 | 1.768400979 | 0.000104465 | 0.004066944 | Pleckstrin homology-like domain family A member 3 |
| PHOSPHO1 | 118.0194979 | 333.6019151 | 2.826667805 | 9.58E-07 | 9.14E-05 | Phosphoethanolamine/phosphocholine phosphatase |
| PHYHD1 | 300.9169836 | 732.1681678 | 2.433123445 | 1.64E-07 | 2.05E-05 | Phytanoyl-CoA dioxygenase domain containing 1, isoform CRA_c |
| PI15 | 103.8756453 | 5.747938839 | 0.055334808 | 4.75E-14 | 3.36E-11 | Peptidase inhibitor 15 |
| PIBF1 | 831.8117575 | 402.9130822 | 0.484380124 | 3.76E-05 | 0.001795653 | Progesterone-induced-blocking factor 1 |
| PICK1 | 99.96765415 | 255.2874371 | 2.553700387 | 6.35E-06 | 0.000432092 | PRKCA-binding protein |
| PIGQ | 740.5177162 | 1168.804812 | 1.578361715 | 0.002540797 | 0.045162307 | Phosphatidylinositol N-acetylglucosaminyltransferase subunit Q |
| PIK3IP1 | 367.5653182 | 1162.140266 | 3.161724483 | 2.37E-07 | 2.78E-05 | Phosphoinositide-3-kinase-interacting protein 1 |
| PIKFYVE | 732.4968596 | 1209.356403 | 1.651005581 | 0.00088006 | 0.020676081 | 1-phosphatidylinositol 3-phosphate 5-kinase |
| PIM3 | 271.5279013 | 459.1876258 | 1.691125014 | 0.001052898 | 0.023511442 | Serine/threonine-protein kinase pim-3 |
| PIP5K1A | 1580.280873 | 952.7035487 | 0.602869759 | 0.000872746 | 0.020519756 | Phosphatidylinositol 4-phosphate 5-kinase type-1 alpha |
| PIPOX | 232.1884282 | 87.31025159 | 0.376031882 | 0.000240431 | 0.00766276 | Peroxisomal sarcosine oxidase |

| Gene name | **Controls** normalized mean counts | **Simvastatin** normalized mean counts | **foldChange** | **pval** | **padj** | **Protein name** |
|---|---|---|---|---|---|---|
| PITPNM2 | 402.4920313 | 727.1038842 | 1.806505043 | 0.000159242 | 0.005648091 | Membrane-associated phosphatidylinositol transfer protein 2 |
| PITX2 | 1638.650285 | 2653.182372 | 1.619126666 | 0.00103418 | 0.023193363 | Pituitary homeobox 2 |
| PIWIL2 | 8.22481765 | 26.2205274 | 3.187976745 | 0.002787578 | 0.048577252 | Piwi-like protein 2 |
| PKD1 | 5110.215208 | 7874.685986 | 1.540969542 | 0.002853243 | 0.049417017 | Polycystin-1 |
| PKDCC | 562.7666544 | 984.5726237 | 1.749521966 | 0.000758372 | 0.018509072 | Protein kinase domain-containing protein, cytoplasmic |
| PKHD1 | 2102.634982 | 950.05473 | 0.451840067 | 0.000118198 | 0.00445466 | Fibrocystin |
| PKIG | 311.4269261 | 544.7381681 | 1.749168497 | 0.000410152 | 0.011747868 | cAMP-dependent protein kinase inhibitor gamma |
| PKN3 | 169.8625075 | 351.0329 | 2.066570811 | 6.71E-05 | 0.00284589 | Serine/threonine-protein kinase N3 |
| PKP4 | 2223.821165 | 1053.959536 | 0.473940779 | 3.59E-05 | 0.001725077 | Plakophilin-4 |
| PLA2G4C | 748.1655011 | 1625.113729 | 2.172131335 | 6.45E-07 | 6.45E-05 | Cytosolic phospholipase A2 gamma |
| PLAGL1 | 1079.888925 | 660.6710267 | 0.611795354 | 0.001703123 | 0.033382072 | Zinc finger protein PLAGL1 |
| PLAT | 1372.487648 | 635.7094968 | 0.463180487 | 2.24E-05 | 0.001177188 | Tissue-type plasminogen activator chain A |
| PLCD1 | 464.3924585 | 828.6973785 | 1.784476391 | 0.000148058 | 0.005342803 | 1-phosphatidylinositol 4,5-bisphosphate phosphodiesterase delta-1 |
| PLCG2 | 66.91059625 | 16.68790089 | 0.249405951 | 0.00064984 | 0.016435062 | 1-phosphatidylinositol 4,5-bisphosphate phosphodiesterase gamma-2 |
| PLCH1 | 68.41508628 | 18.75306951 | 0.274107226 | 4.81E-05 | 0.002191842 | 1-phosphatidylinositol 4,5-bisphosphate phosphodiesterase eta-1 |
| PLCH2 | 40.02547929 | 97.97974652 | 2.447934373 | 0.00022118 | 0.00718018 | 1-phosphatidylinositol 4,5-bisphosphate phosphodiesterase eta-2 |
| PLCXD2 | 158.5320241 | 79.46866218 | 0.501278291 | 0.000762484 | 0.018573084 | PI-PLC X domain-containing protein 2 |
| PLEC | 20175.38004 | 41473.27811 | 2.055638012 | 8.53E-07 | 8.24E-05 | Plectin |
| PLEKHA4 | 1565.018721 | 817.7829097 | 0.522538739 | 0.000757481 | 0.018509072 | Pleckstrin homology domain-containing family A member 4 |
| PLEKHA6 | 3939.028953 | 1926.350703 | 0.489042027 | 3.13E-06 | 0.000237672 | |
| PLEKHF1 | 142.7722504 | 360.7247433 | 2.526574613 | 6.65E-08 | 9.81E-06 | Pleckstrin homology domain-containing family F member 1 |
| PLEKHG2 | 2763.578183 | 465.6813016 | 0.16850665 | 2.43E-13 | 1.48E-10 | Pleckstrin homology domain-containing family G member 2 |
| PLEKHG5 | 6182.126334 | 2847.569897 | 0.460613346 | 2.11E-05 | 0.001120874 | Pleckstrin homology domain-containing family G member 5 |
| PLEKHO1 | 1871.912266 | 3072.835055 | 1.64154865 | 0.000755652 | 0.018480153 | Pleckstrin homology domain-containing family O member 1 |
| PLGRKT | 980.6245464 | 417.4122232 | 0.42565957 | 2.33E-05 | 0.001212845 | Plasminogen receptor (KT) |
| PLIN2 | 470.0594103 | 1181.722793 | 2.513986034 | 0.0021702 | 0.040015561 | Perilipin-2 |
| PLK2 | 4810.505908 | 1686.353313 | 0.350556333 | 1.96E-06 | 0.000162099 | Serine/threonine-protein kinase PLK2 |
| PLN | 11.75960667 | 1.540884379 | 0.131031966 | 0.00018571 | 0.006313336 | Cardiac phospholamban |

| Gene name | **Controls** normalized mean counts | **Simvastatin** normalized mean counts | foldChange | pval | padj | Protein name |
|---|---|---|---|---|---|---|
| PLOD2 | 2206.600279 | 783.2553163 | 0.354960218 | 4.06E-08 | 6.48E-06 | cDNA FLJ43712 fis, clone TESOP2004114, highly similar to Homo sapiens procollagen-lysine, 2-oxoglutarate 5-dioxygenase 2 (PLOD2), transcript variant 1, mRNA |
| PLP2 | 789.5294476 | 1384.012959 | 1.752959264 | 0.000194814 | 0.006537084 | Proteolipid protein 2 |
| PLXNA2 | 411.3347438 | 719.5740887 | 1.749363747 | 0.000339912 | 0.010024711 | Plexin-A2 |
| PMEPA1 | 1740.136197 | 999.9254568 | 0.574624824 | 0.0015498 | 0.031405915 | Transmembrane prostate androgen-induced protein |
| PNMAL1 | 1974.774804 | 1218.690429 | 0.617128812 | 0.001599391 | 0.032135878 | |
| PNO1 | 931.5632051 | 474.1560188 | 0.508989638 | 4.39E-05 | 0.002030899 | RNA-binding protein PNO1 |
| PNPLA7 | 383.5051727 | 864.4683167 | 2.254124268 | 1.83E-05 | 0.000992178 | Patatin-like phospholipase domain-containing protein 7 |
| PNRC1 | 549.0496452 | 1505.769569 | 2.74250167 | 1.66E-09 | 3.90E-07 | Proline-rich nuclear receptor coactivator 1 |
| PODXL2 | 193.9292722 | 108.5599496 | 0.559791455 | 0.000965216 | 0.022006301 | Podocalyxin-like protein 2 |
| POLA2 | 845.800285 | 388.8261248 | 0.459713873 | 4.85E-05 | 0.002203351 | cDNA FLJ60312, highly similar to DNA polymerase subunit alpha B |
| POLR3H | 1033.239367 | 602.1293253 | 0.582758792 | 0.000468549 | 0.012943762 | DNA-directed RNA polymerase III subunit RPC8 |
| POM121L9P | 48.28549083 | 12.20270423 | 0.252719896 | 2.39E-05 | 0.001236645 | |
| POP1 | 181.1794669 | 74.72403343 | 0.412431026 | 2.34E-06 | 0.000187763 | Ribonucleases P/MRP protein subunit POP1 |
| POP5 | 372.8324368 | 223.6571666 | 0.599886556 | 0.00175537 | 0.033893572 | Ribonuclease P/MRP protein subunit POP5 |
| PPDPF | 2961.49306 | 4572.206672 | 1.543885662 | 0.002780102 | 0.048501325 | Pancreatic progenitor cell differentiation and proliferation factor |
| PPIB | 2693.250948 | 1545.795484 | 0.573951523 | 0.000207725 | 0.006852055 | Peptidyl-prolyl cis-trans isomerase B |
| PPIL1 | 587.6747338 | 324.6799303 | 0.552482371 | 0.000239618 | 0.007644669 | Peptidyl-prolyl cis-trans isomerase-like 1 |
| PPIL6 | 33.68118376 | 88.83745046 | 2.637598817 | 5.54E-05 | 0.002473966 | cDNA FLJ50002, highly similar to Homo sapiens peptidylprolyl isomerase (cyclophilin)-like 6 (PPIL6), mRNA |
| PPM1A | 1317.790738 | 2091.600298 | 1.587202153 | 0.001985287 | 0.037067427 | Protein phosphatase 1A |
| PPM1B | 1215.599187 | 2047.491022 | 1.684347146 | 0.000415962 | 0.011881474 | Protein phosphatase 1B |
| PPME1 | 8627.039483 | 2127.454843 | 0.246603119 | 2.26E-12 | 1.00E-09 | Protein phosphatase methylesterase 1 |
| PPP1R14C | 14.08050119 | 65.5272152 | 4.653755879 | 1.94E-05 | 0.00104455 | Protein phosphatase 1 regulatory subunit 14C |
| PPP1R1A | 140.1444579 | 328.6666581 | 2.345199111 | 7.90E-06 | 0.000513507 | Protein phosphatase 1 regulatory subunit 1A |
| PPP1R27 | 23.15184079 | 140.332624 | 6.061402427 | 2.09E-13 | 1.30E-10 | Protein phosphatase 1 regulatory subunit 27 |
| PPP1R3C | 1625.878346 | 2863.41975 | 1.761152523 | 0.000203307 | 0.006720583 | Protein phosphatase 1 regulatory subunit 3C |
| PPP1R3E | 16.93656891 | 62.11163305 | 3.667309087 | 1.71E-06 | 0.000147375 | Protein phosphatase 1 regulatory subunit 3E |
| PPP2R2C | 27.73910676 | 68.61396365 | 2.473546255 | 0.000890308 | 0.020853427 | Serine/threonine-protein phosphatase 2A 55 kDa regulatory subunit B gamma isoform |

| Gene name | **Controls** normalized mean counts | **Simvastatin** normalized mean counts | foldChange | pval | padj | Protein name |
|---|---|---|---|---|---|---|
| PPP2R5A | 460.3534028 | 999.9777628 | 2.172195875 | 5.03E-07 | 5.33E-05 | Serine/threonine-protein phosphatase 2A 56 kDa regulatory subunit alpha isoform |
| PREX1 | 146.7228179 | 285.3527966 | 1.944842668 | 0.000101647 | 0.00396716 | Phosphatidylinositol 3,4,5-trisphosphate-dependent Rac exchanger 1 protein |
| PRICKLE1 | 881.0448561 | 464.5574787 | 0.527280167 | 0.002565526 | 0.045523825 | Prickle-like protein 1 |
| PRICKLE2 | 1477.893451 | 525.6226592 | 0.355656667 | 7.06E-07 | 7.00E-05 | Prickle-like protein 2 |
| PRKCA | 470.0178711 | 877.9831163 | 1.867978156 | 0.00148348 | 0.030298831 | Protein kinase C alpha type |
| PRKCD | 269.5510779 | 495.9333429 | 1.839849229 | 9.55E-05 | 0.003785512 | Protein kinase C delta type |
| PRKCQ | 198.225166 | 693.9264992 | 3.500698287 | 9.85E-06 | 0.000611337 | Protein kinase C theta type |
| PRMT5 | 2252.778393 | 1418.140326 | 0.629507248 | 0.002179012 | 0.040154228 | Protein arginine methyltransferase 5, isoform CRA_c |
| PROB1 | 950.1025353 | 1614.318611 | 1.699099362 | 0.000649636 | 0.016435062 | |
| PROM1 | 1.251856314 | 32.17906605 | 25.70507948 | 5.98E-10 | 1.54E-07 | Prominin-1 |
| PRR16 | 404.7415303 | 216.4107576 | 0.534688786 | 0.000108521 | 0.004198591 | |
| PRR7 | 168.5453346 | 67.54728769 | 0.400766286 | 1.09E-06 | 0.000102445 | Proline-rich protein 7 |
| PRRX1 | 1129.720914 | 3095.186196 | 2.739779496 | 5.74E-06 | 0.000399993 | Paired mesoderm homeobox protein 1 |
| PSD4 | 10.82819869 | 35.16712847 | 3.247735793 | 0.000295656 | 0.008972928 | PH and SEC7 domain-containing protein 4 |
| PSMC3IP | 119.8230157 | 229.5030375 | 1.915350203 | 0.000306052 | 0.009213844 | Homologous-pairing protein 2 homolog |
| PSMD10 | 1639.709684 | 1020.353417 | 0.622276874 | 0.001845562 | 0.03511394 | 26S proteasome non-ATPase regulatory subunit 10 |
| PSTPIP2 | 89.30885114 | 45.11113806 | 0.505113855 | 0.001187684 | 0.025727091 | Proline-serine-threonine phosphatase-interacting protein 2 |
| PTCH2 | 23.73618844 | 61.56290347 | 2.593630549 | 0.000448601 | 0.012560013 | Protein patched homolog 2 |
| PTCHD1 | 21.83839448 | 80.5107709 | 3.686661625 | 1.44E-06 | 0.000128067 | Patched domain-containing protein 1 |
| PTGER2 | 126.9205648 | 58.50508674 | 0.460958292 | 0.000843237 | 0.019992182 | Prostaglandin E2 receptor EP2 subtype |
| PTGS1 | 447.9937123 | 195.8054963 | 0.437071974 | 0.000597073 | 0.015554896 | Prostaglandin G/H synthase 1 |
| PTK2B | 256.5465166 | 425.3896232 | 1.658138372 | 0.001696489 | 0.03329101 | Protein-tyrosine kinase 2-beta |
| PTK7 | 813.9809833 | 407.9845931 | 0.501221283 | 0.000176659 | 0.006092179 | Inactive tyrosine-protein kinase 7 |
| PTP4A3 | 478.440213 | 1187.373111 | 2.481758594 | 6.15E-05 | 0.002674521 | Protein tyrosine phosphatase type IVA 3 |
| PTPDC1 | 289.3707029 | 500.1908437 | 1.728546942 | 0.000598182 | 0.015570745 | Protein tyrosine phosphatase domain-containing protein 1 |
| PTPN13 | 474.3282842 | 830.2784584 | 1.75043 | 0.000323869 | 0.009649671 | Tyrosine-protein phosphatase non-receptor type 13 |
| PTPRB | 51.4171604 | 159.8603996 | 3.109086507 | 6.87E-06 | 0.000460509 | Receptor-type tyrosine-protein phosphatase beta |
| PTPRCAP | 33.11311648 | 82.26741192 | 2.484435796 | 0.000100106 | 0.003926719 | Protein tyrosine phosphatase receptor type C-associated protein |
| PTPRQ | 180.0950725 | 68.10519112 | 0.378162435 | 0.000138507 | 0.005056816 | Phosphatidylinositol phosphatase PTPRQ |
| PTPRU | 499.3612358 | 914.7431739 | 1.831826558 | 6.22E-05 | 0.002690069 | Receptor-type tyrosine-protein phosphatase U |

| Gene name | Controls normalized mean counts | Simvastatin normalized mean counts | foldChange | pval | padj | Protein name |
|---|---|---|---|---|---|---|
| PTRF | 10858.25362 | 6864.540178 | 0.632195602 | 0.002132976 | 0.039422661 | Polymerase I and transcript release factor |
| PTRH1 | 677.620943 | 384.6119284 | 0.56759156 | 0.000299709 | 0.009060012 | Probable peptidyl-tRNA hydrolase |
| PTRH2 | 991.4898514 | 617.8820443 | 0.623185445 | 0.001843646 | 0.03511394 | Peptidyl-tRNA hydrolase 2, mitochondrial |
| PTS | 590.0274085 | 355.2819447 | 0.602144815 | 0.001620241 | 0.032412634 | 6-pyruvoyl tetrahydrobiopterin synthase |
| PUS7 | 573.0951499 | 330.5848641 | 0.576841148 | 0.000592766 | 0.015468619 | Pseudouridylate synthase 7 homolog |
| PVRL2 | 1939.666645 | 1104.99928 | 0.569685148 | 0.000817432 | 0.019529296 | Poliovirus receptor-related protein 2 |
| PVT1 | 214.3856912 | 114.5116204 | 0.534138355 | 0.000426157 | 0.012117028 | |
| PXK | 309.4291829 | 510.0472629 | 1.648348931 | 0.001601212 | 0.032135878 | PX domain-containing protein kinase-like protein |
| PYCRL | 337.1862636 | 172.0128922 | 0.510142051 | 7.52E-05 | 0.003118792 | Pyrroline-5-carboxylate reductase 3 |
| PYURF | 392.8753917 | 212.0279549 | 0.539682452 | 0.00263951 | 0.046597383 | Protein preY, mitochondrial |
| QRICH2 | 56.88379406 | 109.025979 | 1.916643937 | 0.001772608 | 0.034037863 | |
| R3HDM1 | 1072.351796 | 619.0688062 | 0.577300107 | 0.000365358 | 0.010651182 | R3H domain-containing protein 1 |
| RAB11FIP5 | 2354.936273 | 3808.478871 | 1.617232243 | 0.000990467 | 0.022472251 | Rab11 family-interacting protein 5 |
| RAB15 | 3093.796567 | 870.980199 | 0.281524716 | 6.34E-09 | 1.25E-06 | Ras-related protein Rab-15 |
| RAB30 | 329.4556958 | 115.4291151 | 0.350363088 | 7.79E-07 | 7.62E-05 | Ras-related protein Rab-30 |
| RAB33A | 6.059651694 | 23.46369457 | 3.872119348 | 0.000336342 | 0.009945094 | Ras-related protein Rab-33A |
| RAB4B-EGLN2 | 153.7533852 | 279.4524831 | 1.817537107 | 0.001122203 | 0.024695924 | |
| RAB6B | 314.0042692 | 170.2179227 | 0.542087925 | 0.000285566 | 0.008751534 | cDNA FLJ57728, highly similar to Ras-related protein Rab-6B |
| RAD21 | 10943.91374 | 4685.423173 | 0.428130492 | 9.44E-06 | 0.000595873 | Double-strand-break repair protein rad21 homolog |
| RADIL | 51.06587635 | 109.74101 | 2.149008651 | 0.000252878 | 0.007977719 | Ras-associating and dilute domain-containing protein |
| RAET1E | 204.8804309 | 58.09749344 | 0.283567802 | 1.46E-08 | 2.59E-06 | NKG2D ligand 4 |
| RAN | 6590.237015 | 4154.216687 | 0.630359223 | 0.001735719 | 0.033660399 | GTP-binding nuclear protein Ran |
| RANBP10 | 769.582961 | 1302.473865 | 1.692441141 | 0.000708031 | 0.017605118 | RAN binding protein 10, isoform CRA_b |
| RAP1GAP2 | 56.81043086 | 129.7551747 | 2.284002651 | 0.000181825 | 0.006237271 | Rap1 GTPase-activating protein 2 |
| RAPGEF5 | 456.881309 | 109.6362022 | 0.239966486 | 0.002741558 | 0.047936428 | Rap guanine nucleotide exchange factor 5 |
| RARG | 347.7180264 | 782.2320557 | 2.249616057 | 2.02E-07 | 2.45E-05 | Retinoic acid receptor gamma |
| RASA4B | 33.42811455 | 107.545316 | 3.217211543 | 1.34E-05 | 0.000788463 | Putative Ras GTPase-activating protein 4B |
| RASA4CP | 50.36820278 | 150.0826938 | 2.979711118 | 0.000272325 | 0.008445594 | |
| RASD1 | 57.05208563 | 160.6823613 | 2.816415202 | 1.45E-06 | 0.000128647 | Dexamethasone-induced Ras-related protein 1 |
| RASGEF1B | 5.475306098 | 56.30598768 | 10.28362372 | 3.34E-11 | 1.24E-08 | Ras-GEF domain-containing family member 1B |
| RASGEF1C | 7.016094106 | 24.32289085 | 3.466728137 | 0.001456539 | 0.029885948 | Ras-GEF domain-containing family member 1C |

| Gene name | **Controls** normalized mean counts | **Simvastatin** normalized mean counts | foldChange | pval | padj | Protein name |
|---|---|---|---|---|---|---|
| RASL10B | 100.6337715 | 36.2448054 | 0.360165428 | 0.000632334 | 0.016162197 | Ras-like protein family member 10B |
| RASSF2 | 586.1921032 | 73.70434983 | 0.125734123 | 1.42E-25 | 4.02E-22 | Ras association domain-containing protein 2 |
| RASSF3 | 1737.914828 | 1021.918596 | 0.588014199 | 0.000432495 | 0.01225245 | Ras association domain-containing protein 3 |
| RASSF8 | 412.9338598 | 811.6452526 | 1.965557518 | 1.65E-05 | 0.00093288 | Ras association domain-containing protein 8 |
| RBM28 | 1412.727994 | 695.8510494 | 0.492558407 | 4.34E-06 | 0.000314868 | RNA-binding protein 28 |
| RBM4 | 2932.379352 | 1723.696564 | 0.587814998 | 0.000437862 | 0.012368724 | RNA-binding protein 4 |
| RBM8A | 4815.529752 | 2958.129788 | 0.614289588 | 0.001197582 | 0.025923452 | RNA-binding protein 8A |
| RCN1 | 7121.202115 | 4210.789853 | 0.591303236 | 0.000456023 | 0.012710615 | cDNA FLJ55835, highly similar to Reticulocalbin-1 |
| RDH10 | 342.3991636 | 186.0164663 | 0.543273717 | 0.000273961 | 0.008475257 | Retinol dehydrogenase 10 |
| REEP2 | 551.5726574 | 170.2011264 | 0.308574263 | 1.17E-12 | 5.78E-10 | Receptor expression-enhancing protein 2 |
| REEP6 | 44.64651045 | 90.32123166 | 2.023030036 | 0.002132829 | 0.039422661 | Receptor expression-enhancing protein 6 |
| REM2 | 6.789285863 | 23.90267162 | 3.520645927 | 0.002223729 | 0.040784971 | GTP-binding protein REM 2 |
| RFFL | 252.6851387 | 458.6319831 | 1.815033466 | 0.000219478 | 0.007133846 | E3 ubiquitin-protein ligase rififylin |
| RGL2 | 303.4926018 | 598.9737192 | 1.973602374 | 1.76E-05 | 0.000969071 | Ral guanine nucleotide dissociation stimulator-like 2 |
| RGMA | 69.81718501 | 157.9039898 | 2.261677978 | 0.000224414 | 0.007253947 | Repulsive guidance molecule A |
| RGN | 36.15040725 | 12.94140786 | 0.357987886 | 0.002381254 | 0.043016573 | Regucalcin |
| RGS16 | 157.8452986 | 63.26453223 | 0.400800865 | 0.000452826 | 0.012644174 | Regulator of G-protein signaling 16 |
| RGS3 | 906.8700657 | 1518.067113 | 1.673963195 | 0.00133138 | 0.027944601 | cDNA FLJ39521 fis, clone PUAEN2001586, highly similar to Regulator of G-protein signaling 3 |
| RGS6 | 93.73362157 | 31.33308224 | 0.334277943 | 1.81E-05 | 0.000987503 | cDNA FLJ53896, highly similar to Regulator of G-protein signaling 6 |
| RGS9 | 24.79950337 | 5.942237799 | 0.239611161 | 0.000177362 | 0.006109648 | Regulator of G-protein-signaling 9 |
| RHOA | 12895.28786 | 38035.91023 | 2.949597608 | 1.85E-13 | 1.18E-10 | Transforming protein RhoA |
| RHOB | 1358.416002 | 7431.184856 | 5.470478001 | 9.50E-23 | 1.97E-19 | Rho-related GTP-binding protein RhoB |
| RHOQ | 1421.659433 | 2544.387094 | 1.789730392 | 8.45E-05 | 0.003414061 | Rho-related GTP-binding protein RhoQ |
| RICTOR | 596.6900993 | 1015.020385 | 1.701084677 | 0.000652454 | 0.016486798 | Rapamycin-insensitive companion of mTOR |
| RIMS3 | 125.8278513 | 245.3198496 | 1.949646656 | 0.000514683 | 0.013970105 | Regulating synaptic membrane exocytosis protein 3 |
| RMDN2 | 268.9419069 | 672.4650384 | 2.500410019 | 0.000648733 | 0.016435062 | Regulator of microtubule dynamics protein 2 |
| RMRP | 2.741746892 | 110.5094042 | 40.30620207 | 8.41E-31 | 3.27E-27 | |
| RN7SK | 24.06731186 | 416.9367614 | 17.32377774 | 6.07E-42 | 2.70E-38 | |
| RN7SL2 | 114.702405 | 803.7279812 | 7.007071745 | 8.59E-09 | 1.62E-06 | |
| RNA5-8SP6 | 133.8857898 | 3969.211044 | 29.64624587 | 5.66E-79 | 1.76E-74 | |
| RND3 | 2909.594487 | 1506.492143 | 0.517767046 | 4.24E-05 | 0.001985755 | Rho-related GTP-binding protein RhoE |
| RNF115 | 1691.677154 | 3692.292033 | 2.182622154 | 1.62E-06 | 0.00014109 | E3 ubiquitin-protein ligase RNF115 |
| RNF121 | 1176.468841 | 654.6431307 | 0.556447488 | 0.000155026 | 0.005530108 | RING finger protein 121 |
| RNF122 | 286.8548232 | 497.4140851 | 1.734027267 | 0.000695094 | 0.017394695 | RING finger protein 122 |

| Gene name | Controls normalized mean counts | Simvastatin normalized mean counts | foldChange | pval | padj | Protein name |
|---|---|---|---|---|---|---|
| RNF128 | 10.86259283 | 40.79277092 | 3.755343826 | 6.83E-05 | 0.002883672 | E3 ubiquitin-protein ligase RNF128 |
| RNF144B | 904.888101 | 364.6022837 | 0.402925271 | 0.002329353 | 0.042372432 | cDNA FLJ59797, highly similar to E3 ubiquitin ligase IBRDC2 (EC 6.3.2.-) |
| RNF152 | 82.37246121 | 29.0003807 | 0.352064031 | 0.00011363 | 0.004320982 | E3 ubiquitin-protein ligase RNF152 |
| RNF19A | 732.1476523 | 1218.80468 | 1.664697928 | 0.000751848 | 0.018414942 | E3 ubiquitin-protein ligase RNF19A |
| RNF19B | 439.4256569 | 717.8396239 | 1.633586052 | 0.001125178 | 0.024734825 | E3 ubiquitin-protein ligase RNF19B |
| RNF213 | 986.5209673 | 1995.593955 | 2.022860153 | 7.58E-06 | 0.000501704 | E3 ubiquitin-protein ligase RNF213 |
| RNF224 | 13.22017289 | 40.07763279 | 3.031551336 | 0.001270889 | 0.027058366 | RING finger protein 224 |
| RNF39 | 32.36746994 | 80.49681604 | 2.48696658 | 0.001149039 | 0.025099726 | RING finger protein 39 |
| RNF41 | 2689.077622 | 1396.876744 | 0.519463154 | 0.000189326 | 0.006415213 | E3 ubiquitin-protein ligase NRDP1 |
| RNPEPL1 | 812.2717141 | 1405.959939 | 1.730898558 | 0.000221365 | 0.00718018 | Arginyl aminopeptidase-like 1 |
| RNU12 | 2.000042707 | 13.72476772 | 6.862237324 | 0.001506943 | 0.030717547 | |
| ROBO4 | 77.11829302 | 174.6250886 | 2.264379589 | 1.77E-05 | 0.000973069 | Roundabout homolog 4 |
| RORA | 267.8015576 | 645.7284554 | 2.411219939 | 3.61E-08 | 5.89E-06 | Nuclear receptor ROR-alpha |
| RP11-1055B8.4 | 19.21510331 | 61.89733257 | 3.221285442 | 6.16E-05 | 0.002674521 | |
| RP11-108K14.8 | 78.30251147 | 205.5230445 | 2.624731194 | 3.34E-05 | 0.00162944 | |
| RP11-108K3.1 | 221.7827542 | 1089.414413 | 4.912079015 | 1.42E-07 | 1.82E-05 | |
| RP11-111F5.4 | 154.9942026 | 79.54745896 | 0.513228609 | 0.000402301 | 0.011554917 | |
| RP11-114F10.2 | 104.7152349 | 7.491376637 | 0.071540465 | 5.11E-10 | 1.37E-07 | |
| RP11-114H24.2 | 3.684396323 | 37.87816205 | 10.28069695 | 8.85E-05 | 0.00355082 | |
| RP11-1223D19.1 | 440.678138 | 29.89338378 | 0.06783496 | 4.55E-06 | 0.000328439 | |
| RP11-122A3.2 | 82.53642697 | 40.43116513 | 0.489858437 | 0.001414132 | 0.029286283 | |
| RP11-137H2.6 | 86.44440525 | 152.3648018 | 1.762575628 | 0.001928262 | 0.036329807 | |
| RP11-1407O15.2 | 1090.002357 | 1808.91451 | 1.659551008 | 0.000613253 | 0.015791273 | |
| RP11-141O15.1 | 49.75449441 | 102.9800457 | 2.069763685 | 0.001762986 | 0.033956313 | |
| RP11-143J24.1 | 8.665081669 | 0.944525868 | 0.109003689 | 0.001635301 | 0.03264934 | |
| RP11-161M6.2 | 693.9581303 | 1627.094279 | 2.344657708 | 4.72E-07 | 5.04E-05 | |
| RP11-166D19.1 | 2737.617475 | 1337.315231 | 0.488496017 | 0.000362602 | 0.010610625 | |
| RP11-175P13.2 | 3.729267489 | 21.41472981 | 5.742342128 | 0.000196863 | 0.006570394 | |
| RP11-176H8.1 | 122.0063642 | 61.04212715 | 0.500319205 | 0.000847993 | 0.020059063 | |
| RP11-178L8.4 | 428.1571462 | 847.3218086 | 1.978997235 | 1.35E-05 | 0.000789097 | |
| RP11-191N8.2 | 114.4691492 | 346.8266 | 3.029869642 | 6.15E-09 | 1.22E-06 | |
| RP11-203E8.1 | 10.71429884 | 129.9548668 | 12.12910604 | 7.98E-13 | 4.07E-10 | |
| RP11-203J24.9 | 501.2083248 | 890.2679869 | 1.776243416 | 0.000215564 | 0.007058261 | |
| RP11-216B9.6 | 76.00370098 | 32.79145763 | 0.431445538 | 0.000196202 | 0.006561974 | |
| RP11-236B18.2 | 37.06488686 | 9.458192264 | 0.25517931 | 1.11E-05 | 0.000670596 | |
| RP11-244K5.8 | 281.7876551 | 153.2562475 | 0.543871404 | 0.000609791 | 0.015727839 | |
| RP11-244O19.1 | 110.2906864 | 259.3786223 | 2.351772673 | 0.001229717 | 0.026506209 | |
| RP11-245M24.1 | 264.5252471 | 630.6573268 | 2.384110151 | 5.87E-06 | 0.000405848 | |
| RP11-24F11.2 | 33.9700184 | 2.988980845 | 0.087988791 | 5.90E-10 | 1.54E-07 | |
| RP11-255A11.21 | 146.7992429 | 296.4991229 | 2.019759211 | 0.001120453 | 0.024695924 | |
| RP11-266E14.1 | 557.291074 | 251.0798494 | 0.450536284 | 0.000869633 | 0.020462021 | |
| RP11-279O9.4 | 12.32543296 | 40.29268729 | 3.269068715 | 0.001707081 | 0.033417539 | |
| RP11-286H14.6 | 140.6107104 | 56.59939002 | 0.402525454 | 0.000127011 | 0.004720205 | |
| RP11-293P20.2 | 693.1326937 | 1383.487734 | 1.99599261 | 6.23E-05 | 0.002690069 | |
| RP11-305L7.3 | 1.509304285 | 14.63357687 | 9.695577637 | 0.000766492 | 0.018638239 | |
| RP11-307P5.1 | 28.66513031 | 6.20598247 | 0.216499364 | 2.11E-05 | 0.001120874 | |
| RP11-319E12.1 | 15.45448002 | 0.766799547 | 0.049616651 | 1.28E-06 | 0.000116836 | |
| RP11-319E12.2 | 96.35732525 | 2.790554655 | 0.028960483 | 5.17E-11 | 1.79E-08 | |
| RP11-320G24.1 | 9.767346111 | 1.480786842 | 0.151605853 | 0.002325892 | 0.042359517 | |
| RP11-332H18.4 | 38.10995961 | 85.14818955 | 2.234276563 | 0.00141352 | 0.029286283 | |
| RP11-334C17.5 | 27.62013507 | 60.7604431 | 2.199860462 | 0.001715871 | 0.033474995 | |
| RP11-345J4.5 | 438.8376104 | 213.8662077 | 0.487347034 | 0.000116371 | 0.004408937 | |

| Gene name | Controls normalized mean counts | Simvastatin normalized mean counts | foldChange | pval | padj | Protein name |
|---|---|---|---|---|---|---|
| RP11-351I21.11 | 244.1676011 | 86.19819881 | 0.353028815 | 0.002083029 | 0.038660331 | |
| RP11-352D13.6 | 17.34838754 | 4.285334074 | 0.247016275 | 0.001433966 | 0.02951234 | |
| RP11-359D14.3 | 259.2645595 | 87.23748073 | 0.336480547 | 9.44E-05 | 0.003748387 | |
| RP11-373D23.3 | 5.806509079 | 21.97724196 | 3.784931989 | 0.001450148 | 0.029794115 | |
| RP11-377D9.3 | 222.3909572 | 115.7057802 | 0.520280958 | 0.00188542 | 0.03576089 | |
| RP11-383J24.6 | 51.97692957 | 100.5399227 | 1.934318235 | 0.001829067 | 0.034883484 | |
| RP11-38L15.3 | 27.77850261 | 5.716342179 | 0.205782949 | 0.000129163 | 0.004777337 | |
| RP11-400N13.3 | 51.53300238 | 177.1302107 | 3.437218918 | 0.000154736 | 0.005526085 | |
| RP11-412H8.2 | 18.77349476 | 2.684352159 | 0.142986279 | 9.18E-06 | 0.000583953 | |
| RP11-417L19.2 | 0.718846604 | 14.22275084 | 19.78551578 | 2.84E-05 | 0.001418865 | |
| RP11-419C5.2 | 12.25680981 | 38.09576704 | 3.108130714 | 0.001228936 | 0.026506209 | |
| RP11-434D12.1 | 19.70025393 | 69.07665886 | 3.506384188 | 5.42E-06 | 0.000383286 | |
| RP11-436K8.1 | 4.825470691 | 20.14110461 | 4.173915024 | 0.001610274 | 0.032253174 | |
| RP11-43N5.1 | 10.16489833 | 37.45905974 | 3.685138654 | 0.0001066 | 0.004129377 | |
| RP11-441O15.3 | 53.88977942 | 123.2882449 | 2.287785296 | 6.40E-05 | 0.002744519 | |
| RP11-443P15.2 | 261.8554104 | 514.3921442 | 1.964412893 | 0.001301234 | 0.027496984 | |
| RP11-446F17.3 | 6.052308003 | 22.96606122 | 3.794595584 | 0.000562003 | 0.014858114 | |
| RP11-446H18.5 | 283.8022083 | 85.69426312 | 0.301950656 | 6.41E-07 | 6.45E-05 | |
| RP11-469H8.6 | 7.978269733 | 0.507760098 | 0.063642884 | 0.001682556 | 0.033103901 | |
| RP11-47I22.2 | 16.239187 | 3.976515888 | 0.244871611 | 0.002210408 | 0.040612497 | |
| RP11-47I22.3 | 160.0937182 | 38.37418646 | 0.239698265 | 2.46E-05 | 0.001261659 | |
| RP11-496B10.3 | 28.00328491 | 78.21467851 | 2.793053699 | 0.000162837 | 0.005749395 | |
| RP11-509J21.1 | 134.6454924 | 238.4821958 | 1.771185886 | 0.001179385 | 0.025618678 | |
| RP11-514P8.7 | 261.2936553 | 560.8565082 | 2.146460494 | 1.88E-06 | 0.000157978 | |
| RP11-514P8.8 | 265.0661468 | 515.6441409 | 1.945341369 | 0.001144615 | 0.025042602 | |
| RP11-520B13.4 | 37.38680765 | 6.036524382 | 0.161461349 | 0.000176196 | 0.006089738 | |
| RP11-524D16__A.3 | 156.2755211 | 34.410132 | 0.22018888 | 1.09E-13 | 7.07E-11 | |
| RP11-540A21.2 | 330.7477992 | 53.81389704 | 0.162703719 | 3.87E-08 | 6.20E-06 | |
| RP11-545I5.3 | 80.53634977 | 165.2639175 | 2.05204132 | 0.000275737 | 0.008517439 | |
| RP11-571M6.8 | 32.06724966 | 7.159354564 | 0.223260636 | 0.00111067 | 0.024517354 | |
| RP11-589P10.7 | 134.3737611 | 69.5481294 | 0.517572247 | 0.002864769 | 0.049533915 | |
| RP11-597D13.9 | 271.8680323 | 115.1272895 | 0.42346755 | 3.36E-05 | 0.001631552 | |
| RP11-59D5__B.2 | 93.2367392 | 29.09822885 | 0.312089731 | 0.002489046 | 0.044445622 | |
| RP11-600F24.7 | 39.15509492 | 114.7329107 | 2.930216641 | 9.86E-07 | 9.35E-05 | |
| RP11-603J24.9 | 39.14540138 | 91.72845631 | 2.343275406 | 0.002310706 | 0.042181235 | |
| RP11-61L23.2 | 78.04850616 | 157.8824251 | 2.022875682 | 0.001232173 | 0.026506209 | |
| RP11-661A12.14 | 55.72146249 | 122.3932196 | 2.196518436 | 8.81E-05 | 0.00353961 | |
| RP11-66N24.4 | 241.21926 | 137.7048817 | 0.570870177 | 0.00109886 | 0.024310901 | |
| RP11-67L2.2 | 79.01722984 | 149.4096218 | 1.890848642 | 0.000850773 | 0.020109544 | |
| RP11-686D22.8 | 56.86234686 | 19.93038365 | 0.350502305 | 0.000238859 | 0.007636126 | |
| RP11-774D14.1 | 55.63847037 | 148.6329581 | 2.671406261 | 6.64E-06 | 0.000449074 | |
| RP11-779O18.3 | 269.1548834 | 96.6658129 | 0.359145677 | 1.63E-07 | 2.05E-05 | |
| RP11-795F19.5 | 95.60021336 | 207.6944084 | 2.172530804 | 0.00100105 | 0.022662772 | |
| RP11-798K3.2 | 35.29719512 | 8.91930273 | 0.252691544 | 0.001478575 | 0.030238374 | |
| RP11-798M19.3 | 54.1110465 | 203.6534083 | 3.763619842 | 6.24E-07 | 6.34E-05 | |
| RP11-798M19.6 | 32.91184321 | 69.25418605 | 2.104232984 | 0.002340365 | 0.042498195 | |
| RP11-802E16.3 | 19.85377272 | 54.62317691 | 2.751274414 | 0.001010047 | 0.022783554 | |
| RP11-815M8.1 | 36.66224532 | 98.00034126 | 2.673058904 | 2.09E-05 | 0.001114559 | |
| RP11-863P13.4 | 18.50178243 | 55.72137349 | 3.011675967 | 0.002861727 | 0.049533915 | |
| RP11-879F14.2 | 98.01117733 | 48.31057547 | 0.492908837 | 0.000956987 | 0.021920509 | |
| RP11-88H9.2 | 41.37903546 | 12.67519389 | 0.306319221 | 2.50E-05 | 0.001274552 | |
| RP11-88I21.2 | 208.7382658 | 25.28764791 | 0.121145243 | 8.14E-08 | 1.15E-05 | |
| RP11-903H12.3 | 12.52418387 | 36.85229921 | 2.942491072 | 0.000221753 | 0.007185268 | |
| RP1-191J18.66 | 50.64326259 | 18.31383445 | 0.361624301 | 0.000278567 | 0.008579321 | |
| RP11-93L9.1 | 21.46537822 | 104.8812951 | 4.886067883 | 5.99E-07 | 6.15E-05 | |
| RP11-950C14.7 | 162.5573653 | 68.12626162 | 0.419090587 | 0.000901344 | 0.021048958 | |
| RP11-96C23.10 | 11.28051524 | 33.66755691 | 2.984576165 | 0.00141635 | 0.029310425 | |
| RP11-96C23.5 | 13.51784322 | 47.7317674 | 3.531019456 | 0.000474716 | 0.013067723 | |
| RP11-977G19.10 | 103.9294868 | 45.89360559 | 0.441584068 | 0.001336169 | 0.028007331 | |
| RP11-99E15.2 | 80.53361235 | 29.42562174 | 0.36538311 | 0.000268908 | 0.008356307 | |
| RP1-34H18.1 | 141.0776896 | 28.54999114 | 0.202370702 | 3.65E-08 | 5.92E-06 | |
| RP13-516M14.1 | 23.60419231 | 60.73275852 | 2.572964908 | 0.000327732 | 0.00974611 | |

| Gene name | Controls normalized mean counts | Simvastatin normalized mean counts | foldChange | pval | padj | Protein name |
|---|---|---|---|---|---|---|
| RP1-66C13.4 | 44.49904091 | 112.8153444 | 2.53523092 | 5.68E-05 | 0.002514863 | |
| RP3-514A23.2 | 25.97196597 | 7.67587252 | 0.295544532 | 0.00269088 | 0.047209545 | |
| RP4-769N13.6 | 641.6190925 | 377.138007 | 0.587791123 | 0.000614012 | 0.015791705 | |
| RP4-773N10.6 | 38.68249908 | 6.511078381 | 0.168321038 | 0.001620319 | 0.032412634 | |
| RP5-1039K5.13 | 0.98966453 | 16.38957599 | 16.56073901 | 2.32E-05 | 0.001207491 | |
| RP5-1070A16.1 | 23.31751293 | 6.476963929 | 0.277772503 | 0.002248884 | 0.041173502 | |
| RP5-1103G7.4 | 384.3551628 | 608.6812305 | 1.583642655 | 0.002785562 | 0.048569339 | |
| RP5-1120P11.1 | 7.534433921 | 24.80348578 | 3.292017163 | 0.00180214 | 0.034485844 | |
| RP5-1165K10.2 | 58.71457657 | 110.8045997 | 1.887173614 | 0.002351137 | 0.04261916 | |
| RP5-1177I5.3 | 22.24717478 | 4.784777442 | 0.215073486 | 0.000129666 | 0.004784586 | |
| RP5-882C2.2 | 33.5108587 | 82.91918659 | 2.474397548 | 0.000282149 | 0.008663891 | |
| RP5-940J5.3 | 6.713792457 | 23.14896862 | 3.447972032 | 0.002196528 | 0.040405204 | |
| RP5-966M1.6 | 1358.99606 | 4011.506198 | 2.951815914 | 9.67E-06 | 0.000602702 | |
| RP5-977B1.11 | 190.0650386 | 62.37371464 | 0.328170373 | 3.33E-07 | 3.71E-05 | |
| RPL10A | 7773.812425 | 4960.245741 | 0.63807119 | 0.002081326 | 0.038660331 | 60S ribosomal protein L10a |
| RPL11 | 11809.94916 | 7388.540301 | 0.625619992 | 0.001304933 | 0.027538162 | 60S ribosomal protein L11 |
| RPL13A | 24816.10555 | 15227.52593 | 0.61361465 | 0.000817449 | 0.019529296 | 60S ribosomal protein L13a |
| RPL17 | 11587.5962 | 7334.693547 | 0.632978007 | 0.001740936 | 0.033719516 | 60S ribosomal protein L17 |
| RPL18A | 5601.058666 | 3002.262743 | 0.536017014 | 0.000421451 | 0.012005173 | 60S ribosomal protein L18a |
| RPL18AP3 | 132.2538521 | 53.60741553 | 0.405337271 | 0.000991912 | 0.022488649 | |
| RPL22L1 | 675.2083909 | 211.5456465 | 0.313304232 | 1.38E-12 | 6.72E-10 | 60S ribosomal protein L22-like 1 |
| RPL27 | 9624.267311 | 6096.131215 | 0.6334125 | 0.001756502 | 0.033894391 | 60S ribosomal protein L27 |
| RPL32 | 10287.50084 | 6526.908406 | 0.63445034 | 0.001951313 | 0.036586824 | 60S ribosomal protein L32 |
| RPL35A | 6034.579305 | 3716.423432 | 0.615854601 | 0.000952407 | 0.021847765 | 60S ribosomal protein L35a |
| RPL36A | 2162.011687 | 878.7636145 | 0.406456459 | 7.45E-05 | 0.003099886 | 60S ribosomal protein L36a |
| RPL39P | 24.62501254 | 3.865656666 | 0.156980901 | 1.95E-06 | 0.000161861 | |
| RPL5 | 15007.02436 | 9651.275029 | 0.643117169 | 0.002580284 | 0.045759594 | 60S ribosomal protein L5 |
| RPL7P3 | 114.9778347 | 268.6450515 | 2.336494267 | 0.000472138 | 0.013008267 | |
| RPPH1 | 5.740941755 | 69.81279337 | 12.1605124 | 6.69E-16 | 5.95E-13 | |
| RPS10 | 9024.819822 | 5088.832133 | 0.563870773 | 0.000336224 | 0.009945094 | 40S ribosomal protein S10 |
| RPS12 | 11156.8903 | 7211.185136 | 0.646343644 | 0.00272021 | 0.047697203 | 40S ribosomal protein S12 |
| RPS15 | 5964.434354 | 3814.082816 | 0.639471002 | 0.002399692 | 0.043306934 | 40S ribosomal protein S15 |
| RPS16 | 7262.305498 | 4343.90693 | 0.59814434 | 0.000538727 | 0.014421372 | 40S ribosomal protein S16 |
| RPS17 | 9506.563961 | 5441.740506 | 0.572419281 | 0.001364792 | 0.028457754 | 40S ribosomal protein S17 |
| RPS18 | 17233.67507 | 10467.46042 | 0.607384111 | 0.001258761 | 0.026892194 | Ribosomal protein S18, isoform CRA_a |
| RPS19 | 10095.10366 | 6335.907112 | 0.627621798 | 0.001600892 | 0.032135878 | 40S ribosomal protein S19 |
| RPS2 | 24741.76108 | 13696.20437 | 0.553566269 | 0.000241106 | 0.007676415 | 40S ribosomal protein S2 |
| RPS26 | 943.4504679 | 432.7799119 | 0.458720332 | 5.58E-06 | 0.000392519 | 40S ribosomal protein S26 |
| RPS27A | 12608.71969 | 8181.702663 | 0.648892423 | 0.002847526 | 0.049372985 | Ubiquitin-40S ribosomal protein S27a |
| RPS3 | 12381.97578 | 7174.952019 | 0.579467457 | 0.000199793 | 0.006631129 | 40S ribosomal protein S3 |
| RPS6KA2 | 2323.941338 | 4282.717085 | 1.842867983 | 0.001273387 | 0.027074494 | Ribosomal protein S6 kinase, 90kDa, polypeptide 2, isoform CRA_a |
| RPSA | 16937.59132 | 9394.394438 | 0.554647604 | 0.000384581 | 0.011117816 | 40S ribosomal protein SA |
| RPTN | 19.3314885 | 3336.902734 | 172.6148886 | 1.60E-59 | 2.49E-55 | Repetin |
| RRAD | 166.8587797 | 583.765173 | 3.498558327 | 6.95E-07 | 6.91E-05 | GTP-binding protein RAD |
| RRAS | 814.2315381 | 1538.70954 | 1.889769026 | 0.000120124 | 0.004518114 | Ras-related protein R-Ras |
| RRM2 | 30.13121595 | 4.665098092 | 0.154826081 | 3.99E-05 | 0.001884916 | Ribonucleoside-diphosphate reductase subunit M2 |
| RRP1B | 1180.356238 | 721.6480835 | 0.611381598 | 0.001683331 | 0.033103901 | Ribosomal RNA processing protein 1 homolog B |
| RRP9 | 510.3617607 | 297.3770561 | 0.582678953 | 0.000853445 | 0.020157378 | U3 small nucleolar RNA-interacting protein 2 |

| Gene name | **Controls** normalized mean counts | **Simvastatin** normalized mean counts | **foldChange** | **pval** | **padj** | **Protein name** |
|---|---|---|---|---|---|---|
| RRS1 | 679.1194458 | 336.5916312 | 0.4956295 | 1.04E-05 | 0.00063257 | Ribosome biogenesis regulatory protein homolog |
| RSAD2 | 0.756374079 | 13.96171077 | 18.45873775 | 8.67E-05 | 0.003494266 | Radical S-adenosyl methionine domain-containing protein 2 |
| RTKN2 | 243.53052 | 137.7844616 | 0.565779031 | 0.001773729 | 0.034037863 | Rhotekin-2 |
| RTN4R | 53.38936494 | 147.8865818 | 2.769963306 | 0.000364909 | 0.010648091 | Reticulon-4 receptor |
| RUNX1T1 | 118.6790672 | 28.90259708 | 0.243535762 | 7.03E-09 | 1.37E-06 | Protein CBFA2T1 |
| RUNX2 | 182.8945493 | 376.5405298 | 2.058784864 | 7.34E-05 | 0.003063181 | Runt-related transcription factor 2 |
| RXRA | 778.3588379 | 1684.073317 | 2.16362073 | 2.49E-07 | 2.91E-05 | cDNA FLJ16020 fis, clone BRAMY2003287, highly similar to RETINOIC ACID RECEPTOR RXR-ALPHA |
| S100A1 | 70.80339731 | 592.882421 | 8.373643688 | 4.91E-13 | 2.68E-10 | Protein S100-A1 |
| S100A13 | 2645.716568 | 5414.16049 | 2.046387189 | 1.17E-06 | 0.000108304 | Protein S100-A13 |
| S100A16 | 4711.779904 | 2940.287608 | 0.624029065 | 0.001571641 | 0.031786382 | Protein S100-A16 |
| SAMD4A | 3531.042267 | 6254.577449 | 1.771311974 | 0.000116653 | 0.004409014 | Protein Smaug homolog 1 |
| SAMD8 | 1144.53238 | 1830.34906 | 1.599211252 | 0.001734628 | 0.033660222 | Sphingomyelin synthase-related protein 1 |
| SAP30L | 552.7554123 | 1218.102509 | 2.20369169 | 2.15E-07 | 2.58E-05 | Histone deacetylase complex subunit SAP30L |
| SATB2 | 289.4762396 | 483.3747766 | 1.669825397 | 0.001662008 | 0.032970933 | DNA-binding protein SATB2 |
| SBF2-AS1 | 477.6300947 | 66.11510265 | 0.138423235 | 8.04E-23 | 1.79E-19 | |
| SBK1 | 93.52876658 | 326.2413366 | 3.488138982 | 0.000147347 | 0.005323331 | Serine/threonine-protein kinase SBK1 |
| SCAF11 | 5437.117191 | 2845.244926 | 0.523300276 | 1.73E-05 | 0.000959701 | Protein SCAF11 |
| SCD5 | 473.6769646 | 276.2750593 | 0.583256269 | 0.001733342 | 0.033660222 | Stearoyl-CoA desaturase 5 |
| SCN10A | 12.77260016 | 1.203565317 | 0.094230251 | 6.06E-05 | 0.002649302 | Sodium channel protein type 10 subunit alpha |
| SCN1B | 128.475265 | 437.0951766 | 3.402173769 | 3.09E-10 | 8.73E-08 | Sodium channel subunit beta-1 |
| SCN3A | 89.80629845 | 10.09279246 | 0.112384016 | 2.11E-06 | 0.000172627 | Sodium channel protein type 3 subunit alpha |
| SCN3B | 257.9466416 | 487.2227742 | 1.888851009 | 0.000181869 | 0.006237271 | Sodium channel subunit beta-3 |
| SCN4A | 5385.124599 | 12751.97145 | 2.367999332 | 1.31E-08 | 2.38E-06 | Sodium channel protein type 4 subunit alpha |
| SCN5A | 6482.660474 | 2863.699702 | 0.441747599 | 1.73E-05 | 0.000959701 | Cardiac sodium channel alpha subunit |
| SCO1 | 1586.799984 | 799.9108083 | 0.504103111 | 1.44E-05 | 0.000832984 | Protein SCO1 homolog, mitochondrial |
| SCUBE2 | 197.0358383 | 606.5922892 | 3.078588618 | 3.23E-05 | 0.001583652 | Signal peptide, CUB and EGF-like domain-containing protein 2 |
| SDHA | 2171.947882 | 4241.338867 | 1.952781143 | 0.00046648 | 0.012921041 | Succinate dehydrogenase [ubiquinone] flavoprotein subunit, mitochondrial |
| SDPR | 1668.649323 | 335.3938678 | 0.200997216 | 3.68E-11 | 1.33E-08 | Serum deprivation-response protein |
| SEC61G | 1261.407632 | 796.2652974 | 0.631251371 | 0.002464247 | 0.044231309 | Protein transport protein Sec61 subunit gamma |
| SECTM1 | 220.2131637 | 556.0906396 | 2.525237957 | 8.28E-05 | 0.003370271 | Secreted and transmembrane protein 1 |
| SEMA3B | 214.5433706 | 476.6276432 | 2.221591102 | 2.91E-06 | 0.000228135 | Semaphorin-3B |
| SEMA3C | 6177.043674 | 2374.2911 | 0.384373371 | 6.70E-06 | 0.000451977 | Semaphorin-3C |
| SEMA3F | 216.1751891 | 76.36184673 | 0.353240569 | 5.81E-08 | 8.73E-06 | Semaphorin-3F |
| SEMA4A | 14.3478773 | 43.19836671 | 3.010784508 | 0.000308359 | 0.009240677 | Semaphorin-4A |
| SEMA4C | 1063.275623 | 1852.587532 | 1.74233989 | 0.000154586 | 0.005526085 | Semaphorin-4C |
| SEMA5A | 463.1514335 | 157.4465721 | 0.339946205 | 1.36E-05 | 0.000792085 | Semaphorin-5A |

| Gene name | Controls normalized mean counts | Simvastatin normalized mean counts | foldChange | pval | padj | Protein name |
|---|---|---|---|---|---|---|
| SEMA6A | 1325.998988 | 381.8276118 | 0.287954678 | 0.000386578 | 0.011154818 | Semaphorin-6A |
| SEMA6D | 212.827509 | 66.23145236 | 0.311197799 | 2.26E-06 | 0.000182744 | Semaphorin-6D |
| SERPINB7 | 78.13213304 | 19.12828757 | 0.244819728 | 6.01E-09 | 1.21E-06 | Serpin B7 |
| SERPINF2 | 13.50313834 | 81.96051298 | 6.069738078 | 2.52E-10 | 7.32E-08 | Alpha-2-antiplasmin |
| SERPINH1 | 8335.059826 | 3655.503447 | 0.438569551 | 8.88E-08 | 1.22E-05 | Serpin H1 |
| SERTAD1 | 400.3171348 | 230.9142779 | 0.576828364 | 0.00116479 | 0.025355381 | SERTA domain-containing protein 1 |
| SERTAD4 | 48.70480988 | 14.58442977 | 0.299445369 | 1.70E-05 | 0.000950492 | |
| SESN2 | 467.235393 | 919.8893483 | 1.968792095 | 3.10E-05 | 0.001526859 | Sestrin-2 |
| SESN3 | 2287.033055 | 788.1559602 | 0.3446194 | 0.001164818 | 0.025355381 | cDNA FLJ58707, highly similar to Sestrin-3 |
| SESTD1 | 525.1831224 | 972.1661435 | 1.851099363 | 5.79E-05 | 0.002554589 | SEC14 domain and spectrin repeat-containing protein 1 |
| SEZ6 | 10.64846557 | 0.749195463 | 0.070357129 | 0.000185582 | 0.006313336 | Seizure protein 6 homolog |
| SFRP4 | 2024.207131 | 997.0068326 | 0.492541903 | 0.000511579 | 0.013910109 | Secreted frizzled-related protein 4 |
| SFTA1P | 1062.960573 | 243.1303653 | 0.228729429 | 2.93E-06 | 0.000229159 | |
| SGK223 | 960.3551436 | 431.9639468 | 0.449796046 | 0.000175706 | 0.006079549 | Tyrosine-protein kinase SgK223 |
| SH3D19 | 705.4273396 | 1320.207545 | 1.871500395 | 0.000245868 | 0.007804066 | SH3 domain-containing protein 19 |
| SH3GLB2 | 1133.916714 | 2495.993952 | 2.201214534 | 5.55E-07 | 5.79E-05 | Endophilin-B2 |
| SH3TC2 | 135.8868391 | 46.07722075 | 0.339085235 | 0.000420752 | 0.012005173 | SH3 domain and tetratricopeptide repeat-containing protein 2 |
| SHROOM2 | 549.5770515 | 70.6261672 | 0.128510037 | 7.69E-06 | 0.000507944 | Protein Shroom2 |
| SHROOM3 | 634.2772535 | 199.9835853 | 0.315293642 | 5.70E-06 | 0.000398425 | Protein Shroom3 |
| SIX5 | 130.9151637 | 263.2579518 | 2.010904958 | 6.31E-05 | 0.002713179 | Homeobox protein SIX5 |
| SKA2 | 779.4800491 | 449.7012521 | 0.576924647 | 0.001855646 | 0.035260684 | Spindle and kinetochore-associated protein 2 |
| SKAP2 | 809.4669769 | 1311.233545 | 1.619872808 | 0.001434422 | 0.02951234 | Src kinase-associated phosphoprotein 2 |
| SKP2 | 1785.909315 | 365.4233381 | 0.204614722 | 5.86E-08 | 8.77E-06 | S-phase kinase-associated protein 2 |
| SLC16A4 | 399.940555 | 191.5057207 | 0.478835463 | 7.45E-05 | 0.003099886 | Monocarboxylate transporter 5 |
| SLC16A8 | 9.967396955 | 34.5966314 | 3.47097959 | 0.000430799 | 0.01221554 | Monocarboxylate transporter 3 |
| SLC17A7 | 10.27625832 | 55.13134188 | 5.364923704 | 0.000600945 | 0.015629598 | Vesicular glutamate transporter 1 |
| SLC19A1 | 219.9239622 | 123.2165023 | 0.560268654 | 0.001155927 | 0.025214773 | Folate transporter 1 |
| SLC1A2 | 15.59887213 | 53.91859492 | 3.456570094 | 5.70E-05 | 0.002522699 | Excitatory amino acid transporter 2 |
| SLC1A3 | 208.3766972 | 77.70775716 | 0.372919612 | 8.16E-07 | 7.93E-05 | Excitatory amino acid transporter 1 |
| SLC1A4 | 1101.46528 | 3058.96058 | 2.777173858 | 9.71E-09 | 1.81E-06 | Neutral amino acid transporter A |
| SLC22A17 | 200.5275475 | 90.7943698 | 0.452777541 | 0.000382484 | 0.011088105 | Solute carrier family 22 member 17 |
| SLC22A23 | 186.9808648 | 349.8255354 | 1.870916233 | 0.000828081 | 0.019707937 | Solute carrier family 22 member 23 |
| SLC24A2 | 2180.656749 | 1175.43819 | 0.539029442 | 0.000937029 | 0.021606889 | Sodium/potassium/calcium exchanger 2 |
| SLC25A28 | 464.3415373 | 782.6099369 | 1.685418758 | 0.000761044 | 0.018552543 | Mitoferrin-2 |
| SLC25A29 | 910.791729 | 1439.382594 | 1.580364147 | 0.00188047 | 0.035688764 | cDNA FLJ38213 fis, clone FCBBF1000574, highly similar to Mitochondrial carnitine/acylcarnitine carrier protein CACL |
| SLC26A4 | 96.77195519 | 31.70443971 | 0.327620121 | 1.17E-05 | 0.000698275 | Pendrin |

| Gene name | **Controls** normalized mean counts | **Simvastatin** normalized mean counts | **foldChange** | **pval** | **padj** | **Protein name** |
|---|---|---|---|---|---|---|
| SLC29A4 | 336.2871002 | 144.4879512 | 0.429656538 | 2.31E-05 | 0.001206008 | Equilibrative nucleoside transporter 4 |
| SLC34A3 | 40.64454499 | 110.7560539 | 2.724991851 | 2.93E-05 | 0.001458003 | Sodium-dependent phosphate transport protein 2C |
| SLC35E3 | 1782.96716 | 894.2856751 | 0.501571591 | 1.36E-05 | 0.000792085 | Solute carrier family 35 member E3 |
| SLC37A1 | 19.18714097 | 68.81884941 | 3.586717245 | 2.26E-05 | 0.001183557 | Glycerol-3-phosphate transporter |
| SLC38A4 | 72.05977277 | 26.59531666 | 0.369073002 | 0.00172102 | 0.033540427 | Sodium-coupled neutral amino acid transporter 4 |
| SLC43A1 | 114.0990974 | 201.2849597 | 1.764124032 | 0.001929591 | 0.036332843 | Large neutral amino acids transporter small subunit 3 |
| SLC43A2 | 574.5426073 | 1237.984682 | 2.154730852 | 3.47E-05 | 0.001680396 | cDNA FLJ55865, highly similar to Homo sapiens solute carrier family 43, member 2 (SLC43A2), mRNA |
| SLC44A2 | 4450.825253 | 7015.96475 | 1.576328962 | 0.001709106 | 0.033425784 | Choline transporter-like protein 2 |
| SLC46A3 | 652.5192423 | 336.5060359 | 0.515702855 | 0.000442152 | 0.012433904 | Solute carrier family 46 member 3 |
| SLC48A1 | 174.8870323 | 365.4806442 | 2.089809859 | 5.55E-05 | 0.002473966 | Heme transporter HRG1 |
| SLC4A11 | 271.1805 | 680.7084327 | 2.510167334 | 7.18E-09 | 1.39E-06 | Sodium bicarbonate transporter-like protein 11 |
| SLC4A3 | 967.3640791 | 484.8351718 | 0.501192035 | 9.82E-06 | 0.000610915 | Anion exchange protein 3 |
| SLC4A4 | 1978.13686 | 822.4968549 | 0.415793705 | 0.002889429 | 0.04982183 | Electrogenic sodium bicarbonate cotransporter 1 |
| SLC5A1 | 44.29785557 | 3.785326486 | 0.085451687 | 2.11E-05 | 0.001120874 | Sodium/glucose cotransporter 1 |
| SLC6A1 | 181.2991279 | 46.87285348 | 0.25853877 | 8.12E-05 | 0.003319528 | Transporter |
| SLC6A17 | 1339.535253 | 116.6413591 | 0.087075991 | 4.37E-05 | 0.002026798 | Sodium-dependent neutral amino acid transporter SLC6A17 |
| SLC7A11 | 1787.484258 | 5250.208376 | 2.937205378 | 5.27E-09 | 1.09E-06 | Cystine/glutamate transporter |
| SLC7A8 | 101.8159033 | 209.0912528 | 2.053620761 | 0.00191832 | 0.036186329 | Large neutral amino acids transporter small subunit 2 |
| SLC9A3R1 | 476.2493079 | 805.5731559 | 1.691494649 | 0.000675994 | 0.016957627 | cDNA FLJ46618 fis, clone TLIVE2007736, highly similar to Ezrin-radixin-moesin-binding phosphoprotein 50 |
| SLC9A4 | 25.22563851 | 7.492484126 | 0.297018612 | 0.001069484 | 0.023779391 | Sodium/hydrogen exchanger 4 |
| SLCO4C1 | 18.17890467 | 66.39453801 | 3.652284843 | 0.000501765 | 0.013715209 | Solute carrier organic anion transporter family member 4C1 |
| SLFN12 | 181.5224511 | 69.66892364 | 0.383803343 | 9.41E-05 | 0.003743602 | Schlafen family member 12 |
| SLIRP | 1098.190798 | 592.7347544 | 0.539737499 | 5.54E-05 | 0.002473966 | SRA stem-loop-interacting RNA-binding protein, mitochondrial |
| SLK | 2081.82362 | 3508.748498 | 1.685420639 | 0.000429676 | 0.0121948 | STE20-like serine/threonine-protein kinase |
| SLN | 5838.103103 | 11904.28811 | 2.039067811 | 0.000802338 | 0.019227678 | Sarcolipin |

| Gene name | **Controls** normalized mean counts | **Simvastatin** normalized mean counts | **foldChange** | **pval** | **padj** | **Protein name** |
|---|---|---|---|---|---|---|
| SMAD3 | 8043.742136 | 3298.470131 | 0.410066618 | 0.000291466 | 0.008888582 | Mothers against decapentaplegic homolog 3 |
| SMARCA2 | 1194.449526 | 1960.67793 | 1.641490819 | 0.000825119 | 0.019682628 | Probable global transcription activator SNF2L2 |
| SNAI3 | 0.766300161 | 19.21636512 | 25.07681206 | 3.01E-06 | 0.000232962 | Zinc finger protein SNAI3 |
| SNAI3-AS1 | 187.1640524 | 461.7188514 | 2.466920574 | 2.19E-08 | 3.76E-06 | |
| SNAP91 | 75.56953286 | 28.06880589 | 0.371430189 | 2.23E-05 | 0.001175879 | Clathrin coat assembly protein AP180 |
| SNED1 | 564.4037627 | 2150.123471 | 3.809548436 | 3.10E-16 | 2.89E-13 | Sushi, nidogen and EGF-like domain-containing protein 1 |
| SNHG15 | 387.1180989 | 222.1527528 | 0.573862998 | 0.000649095 | 0.016435062 | |
| SNHG16 | 2681.689185 | 1305.872029 | 0.486958756 | 1.72E-06 | 0.000147767 | |
| SNHG8 | 793.3708267 | 466.4221905 | 0.587899346 | 0.000574226 | 0.01513717 | |
| SNN | 778.9293654 | 1500.314793 | 1.926124319 | 1.75E-05 | 0.000965897 | Stannin |
| SNORD3A | 3.437735916 | 25.81677402 | 7.50981886 | 1.45E-05 | 0.000836824 | |
| SNRPD1 | 747.9815559 | 401.5701878 | 0.536871778 | 7.84E-05 | 0.003219908 | Small nuclear ribonucleoprotein Sm D1 |
| SNRPE | 891.8465872 | 552.7532243 | 0.619785098 | 0.001872771 | 0.035564351 | Small nuclear ribonucleoprotein E |
| SNRPF | 457.290888 | 279.2959476 | 0.61076211 | 0.001938076 | 0.036448487 | Small nuclear ribonucleoprotein F |
| SNRPN | 4338.68531 | 2688.678317 | 0.619698855 | 0.001141197 | 0.02501626 | Small nuclear ribonucleoprotein-associated protein |
| SNTA1 | 410.7907309 | 1013.493019 | 2.467175966 | 4.61E-09 | 9.88E-07 | Alpha-1-syntrophin |
| SNX21 | 2323.425418 | 4207.6688 | 1.810976487 | 5.61E-05 | 0.00249356 | Sorting nexin-21 |
| SNX29 | 594.9334375 | 1000.674979 | 1.681994852 | 0.000811597 | 0.019434599 | Sorting nexin-29 |
| SNX7 | 469.9150325 | 764.7119304 | 1.627340854 | 0.001903316 | 0.036034416 | Sorting nexin-7 |
| SNX9 | 1073.725145 | 1887.720609 | 1.758104127 | 0.000198279 | 0.006600877 | Sorting nexin-9 |
| SOAT2 | 11.73018731 | 48.70870129 | 4.152423145 | 0.001845666 | 0.03511394 | Sterol O-acyltransferase 2 |
| SOD2 | 4435.343249 | 9394.79573 | 2.118166555 | 1.46E-05 | 0.000836824 | Superoxide dismutase |
| SOD3 | 707.6789107 | 301.0860901 | 0.42545579 | 5.35E-05 | 0.002403955 | Extracellular superoxide dismutase [Cu-Zn] |
| SOGA1 | 1941.062704 | 3121.510264 | 1.60814499 | 0.001330668 | 0.027944601 | C-terminal 80 kDa form |
| SORBS1 | 3077.192175 | 7212.040506 | 2.343708191 | 7.79E-09 | 1.50E-06 | Sorbin and SH3 domain-containing protein 1 |
| SORBS2 | 3844.272261 | 1406.500224 | 0.365869046 | 6.28E-11 | 2.10E-08 | Sorbin and SH3 domain-containing protein 2 |
| SORCS3 | 96.51079158 | 35.17190974 | 0.364434994 | 0.000755698 | 0.018480153 | VPS10 domain-containing receptor SorCS3 |
| SORD | 478.8089131 | 263.986123 | 0.5513392 | 0.000196399 | 0.006561974 | cDNA FLJ50165, moderately similar to Sorbitol dehydrogenase (EC 1.1.1.14) |
| SOX4 | 177.4744278 | 460.8933167 | 2.596956206 | 0.000758666 | 0.018509072 | Transcription factor SOX-4 |
| SOX8 | 998.0403329 | 1561.643771 | 1.564710081 | 0.002727184 | 0.047792558 | Transcription factor SOX-8 |
| SOX9 | 29.46427621 | 68.95500644 | 2.340291882 | 0.001065325 | 0.023720834 | Transcription factor SOX-9 |
| SP7 | 0.495482235 | 30.13643068 | 60.8224242 | 1.44E-09 | 3.47E-07 | Transcription factor Sp7 |
| SPAG11B | 117.4701469 | 14.11859746 | 0.120188812 | 5.85E-07 | 6.05E-05 | Sperm-associated antigen 11B |
| SPAG9 | 2932.769009 | 5094.527973 | 1.73710509 | 0.00016603 | 0.005827147 | C-Jun-amino-terminal kinase-interacting protein 4 |
| SPOCK2 | 37.23454044 | 79.57220478 | 2.137053495 | 0.000847066 | 0.020052376 | Testican-2 |
| SPRED2 | 1094.320873 | 663.7431678 | 0.606534321 | 0.00117473 | 0.025550263 | Sprouty-related, EVH1 domain-containing protein 2 |
| SPRR2G | 8.182608444 | 0 | 0 | 2.01E-05 | 0.001079782 | Small proline-rich protein 2G |

| Gene name | **Controls** normalized mean counts | **Simvastatin** normalized mean counts | foldChange | pval | padj | Protein name |
|---|---|---|---|---|---|---|
| SPSB1 | 268.4203789 | 546.3616474 | 2.03547007 | 8.79E-05 | 0.003535334 | SPRY domain-containing SOCS box protein 1 |
| SPTAN1 | 18427.08986 | 9640.989899 | 0.523196553 | 0.000120266 | 0.004518114 | Spectrin alpha chain, non-erythrocytic 1 |
| SQLE | 1118.613033 | 2643.392594 | 2.363098333 | 4.55E-09 | 9.82E-07 | Squalene monooxygenase |
| SRGAP1 | 898.9383067 | 461.3687762 | 0.513237419 | 0.000311511 | 0.00930821 | SLIT-ROBO Rho GTPase activating protein 1, isoform CRA_a |
| SRL | 6729.818318 | 2950.288239 | 0.438390474 | 7.25E-05 | 0.003040047 | Sarcalumenin |
| SRM | 1554.520453 | 925.8982698 | 0.595616653 | 0.000547502 | 0.01459349 | Spermidine synthase |
| SRP19 | 545.279054 | 276.0794334 | 0.506308525 | 0.000646065 | 0.016405316 | Signal recognition particle 19 kDa protein |
| SRPK3 | 4971.892707 | 12471.66953 | 2.508434969 | 4.99E-09 | 1.04E-06 | SRSF protein kinase 3 |
| SRPX | 1205.002254 | 711.4415758 | 0.590406842 | 0.002295668 | 0.041980626 | Sushi repeat-containing protein SRPX |
| SRPX2 | 1763.118038 | 428.7058902 | 0.243152121 | 1.83E-16 | 1.89E-13 | Sushi repeat-containing protein SRPX2 |
| SRSF1 | 2732.099041 | 1592.470556 | 0.582874388 | 0.000337779 | 0.009978107 | Serine/arginine-rich-splicing factor 1 |
| SRSF3 | 3356.103764 | 2144.573605 | 0.639006942 | 0.002550042 | 0.045300754 | Serine/arginine-rich splicing factor 3 |
| SRXN1 | 985.8599454 | 2446.000521 | 2.481083172 | 3.09E-06 | 0.00023721 | Sulfiredoxin-1 |
| SSB | 3318.84069 | 1969.407751 | 0.593402316 | 0.00052005 | 0.014066666 | Lupus La protein |
| SSH2 | 670.5216224 | 1253.386613 | 1.869270984 | 3.35E-05 | 0.001630053 | Protein phosphatase Slingshot homolog 2 |
| SSH3 | 281.1423819 | 465.0363729 | 1.654095586 | 0.001716473 | 0.033474995 | Protein phosphatase Slingshot homolog 3 |
| SSR3 | 7341.173211 | 4173.013194 | 0.56843955 | 0.000604344 | 0.015678675 | Translocon-associated protein subunit gamma |
| ST3GAL1 | 270.456527 | 580.2256399 | 2.145356395 | 3.00E-06 | 0.000232962 | CMP-N-acetylneuraminate-beta-galactosamide-alpha-2,3-sialyltransferase 1 |
| ST3GAL6 | 47.06359865 | 142.3194367 | 3.023981183 | 0.000257279 | 0.008051218 | Type 2 lactosamine alpha-2,3-sialyltransferase |
| ST5 | 1459.911175 | 798.3698688 | 0.546861948 | 7.97E-05 | 0.003266694 | Suppression of tumorigenicity 5 protein |
| ST6GAL2 | 9.187804351 | 0.283928817 | 0.030902793 | 0.000213343 | 0.007007652 | Beta-galactoside alpha-2,6-sialyltransferase 2 |
| ST6GALNAC2 | 111.1277157 | 286.6690796 | 2.579636213 | 0.000163103 | 0.00575224 | Alpha-N-acetylgalactosaminide alpha-2,6-sialyltransferase 2 |
| STARD13 | 3953.343059 | 1849.555032 | 0.467845822 | 3.27E-05 | 0.001598646 | StAR-related lipid transfer protein 13 |
| STARD3NL | 982.6169293 | 586.9827826 | 0.597366853 | 0.000766956 | 0.018638239 | MLN64 N-terminal domain homolog |
| STARD9 | 202.0083664 | 606.5525553 | 3.00261106 | 1.61E-11 | 6.22E-09 | StAR-related lipid transfer protein 9 |
| STAT5A | 314.3491866 | 715.1346655 | 2.274969034 | 1.60E-07 | 2.01E-05 | Signal transducer and activator of transcription 5A |
| STIP1 | 4267.906517 | 2620.051578 | 0.613896196 | 0.001343231 | 0.028117455 | Stress-induced-phosphoprotein 1 |
| STK10 | 2143.20095 | 945.7042572 | 0.441257856 | 6.52E-05 | 0.002780005 | Serine/threonine-protein kinase 10 |
| STK17B | 1769.204037 | 985.1038528 | 0.556806243 | 0.000729122 | 0.018000047 | Serine/threonine-protein kinase 17B |
| STK24 | 996.9889476 | 1639.627195 | 1.644579109 | 0.000799523 | 0.019197327 | Serine/threonine-protein kinase 24 12 kDa subunit |
| STK31 | 117.9978257 | 4.593574267 | 0.038929313 | 1.16E-12 | 5.78E-10 | Serine/threonine-protein kinase 31 |

| Gene name | **Controls** normalized mean counts | **Simvastatin** normalized mean counts | **foldChange** | **pval** | **padj** | **Protein name** |
|---|---|---|---|---|---|---|
| STK32C | 413.3096603 | 695.385987 | 1.682481814 | 0.002082657 | 0.038660331 | Serine/threonine-protein kinase 32C |
| STRBP | 139.6250291 | 74.11781102 | 0.530834704 | 0.001121411 | 0.024695924 | Spermatid perinuclear RNA-binding protein |
| STRC | 52.43746466 | 113.4293801 | 2.163136239 | 0.000740072 | 0.018227584 | Stereocilin |
| STRN | 990.1237953 | 1903.346614 | 1.922331958 | 1.40E-05 | 0.000812752 | Striatin |
| STT3B | 4672.709071 | 2984.596253 | 0.638729313 | 0.002662198 | 0.046811999 | Dolichyl-diphosphooligosaccharide--protein glycosyltransferase subunit STT3B |
| SULF1 | 1545.74606 | 796.47167 | 0.515266828 | 0.000539333 | 0.014425198 | Extracellular sulfatase Sulf-1 |
| SUMF1 | 6144.897923 | 2137.762315 | 0.347892242 | 9.86E-05 | 0.003875658 | Sulfatase-modifying factor 1 |
| SVIL | 7390.577287 | 11700.11746 | 1.583112794 | 0.00191658 | 0.036175444 | Supervillin |
| SYDE1 | 1517.538753 | 908.0289033 | 0.59835632 | 0.000818063 | 0.019529296 | Synapse defective 1, Rho GTPase, homolog 1 (C. elegans), isoform CRA_a |
| SYNE2 | 409.7109458 | 1873.705407 | 4.573237366 | 4.40E-19 | 5.95E-16 | Nesprin-2 |
| SYNGR1 | 188.6435163 | 324.0999332 | 1.718054983 | 0.001094925 | 0.024258356 | Synaptogyrin-1 |
| SYNJ2 | 1474.948667 | 600.6456716 | 0.407231577 | 3.68E-06 | 0.000273009 | Synaptojanin-2 |
| SYNM | 86.29560031 | 368.3552414 | 4.268528639 | 5.88E-05 | 0.002585401 | Synemin |
| SYNPO | 476.407168 | 1675.291964 | 3.516512926 | 6.09E-13 | 3.21E-10 | Synaptopodin |
| SYNPO2L | 5304.527022 | 10519.69677 | 1.983154525 | 6.21E-05 | 0.002688578 | Synaptopodin 2-like protein |
| SYT1 | 188.7328627 | 87.237991 | 0.462229999 | 0.002404727 | 0.043338038 | Synaptotagmin-1 |
| SYT15 | 136.2452756 | 23.03784799 | 0.169090986 | 7.25E-12 | 3.01E-09 | Synaptotagmin-15 |
| SYT3 | 51.39994467 | 150.0621487 | 2.919500199 | 0.001942604 | 0.036489523 | Synaptotagmin-3 |
| SYT8 | 78.85038979 | 22.89904349 | 0.290411291 | 3.05E-07 | 3.45E-05 | Synaptotagmin-8 |
| SYTL2 | 7096.375225 | 3544.937453 | 0.499541997 | 0.001652638 | 0.032868895 | Synaptotagmin-like protein 2 |
| SYTL4 | 694.5206847 | 235.0060614 | 0.33837158 | 1.92E-10 | 5.92E-08 | Synaptotagmin-like protein 4 |
| TACR2 | 80.90683029 | 193.4163086 | 2.39060544 | 1.24E-05 | 0.000731296 | Substance-K receptor |
| TAF13 | 1384.270626 | 888.1409216 | 0.641594862 | 0.00280567 | 0.048837806 | Transcription initiation factor TFIID subunit 13 |
| TAGLN | 10131.94474 | 1387.196651 | 0.136913168 | 1.96E-11 | 7.43E-09 | Transgelin |
| TANGO6 | 421.1275522 | 232.9361014 | 0.55312482 | 0.000307658 | 0.009237465 | Transport and Golgi organization protein 6 homolog |
| TAS1R1 | 973.2201784 | 316.5167925 | 0.325226295 | 1.86E-07 | 2.30E-05 | Taste receptor type 1 member 1 |
| TBC1D3F | 77.85330576 | 161.9132004 | 2.079721585 | 0.001630996 | 0.032605244 | TBC1 domain family member 3F |
| TCEA3 | 686.9881481 | 2670.101164 | 3.886677189 | 1.17E-11 | 4.66E-09 | Transcription elongation factor A protein 3 |
| TCEAL7 | 519.8473679 | 3234.416483 | 6.221857958 | 1.20E-16 | 1.28E-13 | Transcription elongation factor A protein-like 7 |
| TCF4 | 1718.325173 | 925.8712171 | 0.53882189 | 0.000941679 | 0.021681612 | Transcription factor 4 |
| TCF7 | 40.64946131 | 136.5378493 | 3.358909194 | 1.35E-07 | 1.75E-05 | Transcription factor 7 (T-cell specific, HMG-box), isoform CRA_c |
| TCP11L2 | 239.4257462 | 395.2148164 | 1.650678019 | 0.002342505 | 0.04251223 | |
| TCTN2 | 68.51028299 | 141.4370995 | 2.064465265 | 0.000496921 | 0.013606709 | Tectonic-2 |
| TDG | 1219.312832 | 594.3036602 | 0.487408682 | 0.000204322 | 0.006746975 | cDNA FLJ60553, highly similar to G/T mismatch-specific thymine DNA glycosylase(EC 3.2.2.-) |
| TDP2 | 752.6562068 | 1291.623886 | 1.716087471 | 0.000714648 | 0.017713028 | Tyrosyl-DNA phosphodiesterase 2 |

| Gene name | Controls normalized mean counts | Simvastatin normalized mean counts | foldChange | pval | padj | Protein name |
|---|---|---|---|---|---|---|
| TDRD1 | 51.94200386 | 15.17092294 | 0.292074271 | 0.000109008 | 0.00420174 | Tudor domain-containing protein 1 |
| TEAD2 | 1647.785102 | 993.6638094 | 0.603029975 | 0.000835016 | 0.019842644 | Transcriptional enhancer factor TEF-4 |
| TECRL | 3.720623858 | 18.28162137 | 4.91359032 | 0.00180343 | 0.034485844 | Trans-2,3-enoyl-CoA reductase-like |
| TENM4 | 242.8869199 | 123.2048783 | 0.50725201 | 0.001061806 | 0.023676371 | Teneurin-4 |
| TEX14 | 63.31727293 | 20.68729176 | 0.326724301 | 1.81E-05 | 0.000988703 | Inactive serine/threonine-protein kinase TEX14 |
| TEX2 | 2977.982145 | 1862.228776 | 0.625332418 | 0.002027439 | 0.037781556 | Testis-expressed sequence 2 protein |
| TEX26-AS1 | 2.262234491 | 15.05280597 | 6.653954767 | 0.000644779 | 0.016399421 | |
| TFCP2L1 | 3.39028236 | 25.40935832 | 7.494761682 | 0.001803783 | 0.034485844 | Transcription factor CP2-like protein 1 |
| TGFBR2 | 3920.61194 | 2304.063662 | 0.587679601 | 0.001673501 | 0.033045663 | TGF-beta receptor type-2 |
| TGIF2-C20orf24 | 279.8346227 | 128.3449282 | 0.458645635 | 0.000404211 | 0.011599075 | |
| TGM1 | 88.82640721 | 31.24634758 | 0.351768675 | 4.24E-05 | 0.001985755 | Protein-glutamine gamma-glutamyltransferase K |
| THBS1 | 27242.30423 | 10114.05269 | 0.371262747 | 0.002616517 | 0.046273552 | Thrombospondin-1 |
| THOP1 | 1233.67175 | 751.7184326 | 0.609334235 | 0.002158775 | 0.039852132 | cDNA FLJ36073 fis, clone TESTI2019697, highly similar to THIMET OLIGOPEPTIDASE (EC 3.4.24.15) |
| THSD1 | 1186.080151 | 462.4689225 | 0.389913719 | 1.51E-06 | 0.000132387 | Thrombospondin type-1 domain-containing protein 1 |
| TICAM2 | 93.42673434 | 273.5133884 | 2.927570896 | 0.000562207 | 0.014858114 | TIR domain-containing adapter molecule 2 |
| TIPARP | 977.4984741 | 1770.017393 | 1.810762308 | 0.001609772 | 0.032253174 | TCDD-inducible poly [ADP-ribose] polymerase |
| TJP2 | 452.2276106 | 743.7616604 | 1.644662208 | 0.001693616 | 0.033260952 | Tight junction protein ZO-2 |
| TKT | 1351.432343 | 2307.221779 | 1.707241795 | 0.002526351 | 0.044982637 | Transketolase |
| TLE1 | 132.124268 | 412.5682529 | 3.122577397 | 6.01E-06 | 0.000414115 | Transducin-like enhancer protein 1 |
| TLK1 | 911.1216679 | 1598.951568 | 1.754926509 | 0.000193233 | 0.006512156 | cDNA FLJ34656 fis, clone KIDNE2018352, highly similar to Serine/threonine-protein kinase tousled-like 1 (EC 2.7.11.1) |
| TMCO3 | 1302.98188 | 2179.743807 | 1.672888811 | 0.000538231 | 0.014421372 | Transmembrane and coiled-coil domain-containing protein 3 |
| TMCO4 | 98.63832012 | 211.7345925 | 2.146575411 | 0.001100105 | 0.02432115 | Transmembrane and coiled-coil domain-containing protein 4 |
| TMED3 | 2155.606812 | 1199.031029 | 0.556238282 | 8.17E-05 | 0.003330814 | Transmembrane emp24 domain-containing protein 3 |
| TMEFF1 | 468.6037966 | 756.6394557 | 1.614667788 | 0.00173819 | 0.033687309 | Tomoregulin-1 |
| TMEM106C | 802.4943719 | 452.571498 | 0.563955978 | 0.000227909 | 0.007346463 | Transmembrane protein 106C |
| TMEM126A | 471.440275 | 284.3360857 | 0.603122179 | 0.00158492 | 0.031951073 | Transmembrane protein 126A |
| TMEM132A | 5495.931867 | 1970.837807 | 0.358599389 | 1.01E-09 | 2.52E-07 | Transmembrane protein 132A |
| TMEM132B | 25.60282671 | 111.2292892 | 4.344414408 | 0.000742439 | 0.018256377 | Transmembrane protein 132B |
| TMEM140 | 72.92252722 | 261.982946 | 3.592620222 | 0.000105884 | 0.004106769 | Transmembrane protein 140 |

| Gene name | **Controls** normalized mean counts | **Simvastatin** normalized mean counts | foldChange | pval | padj | Protein name |
|---|---|---|---|---|---|---|
| TMEM144 | 174.0668223 | 96.26427576 | 0.55303058 | 0.001258538 | 0.026892194 | Transmembrane protein 144 |
| TMEM173 | 699.6697125 | 1439.474097 | 2.057362312 | 1.94E-05 | 0.00104455 | Stimulator of interferon genes protein |
| TMEM177 | 161.6910429 | 86.40745428 | 0.534398522 | 0.00089096 | 0.020853427 | Transmembrane protein 177 |
| TMEM182 | 8043.38882 | 4460.179864 | 0.554515014 | 0.002028393 | 0.037781556 | Transmembrane protein 182 |
| TMEM2 | 1435.762138 | 861.6020346 | 0.600100819 | 0.00126392 | 0.026965356 | Transmembrane protein 2 |
| TMEM25 | 238.4232329 | 89.03636563 | 0.373438295 | 6.90E-08 | 1.00E-05 | Transmembrane protein 25 |
| TMEM56 | 88.64593798 | 28.93551423 | 0.326416696 | 0.000188369 | 0.006389746 | Transmembrane protein 56 |
| TMEM59L | 65.65460211 | 25.47054813 | 0.387947643 | 0.000949066 | 0.021803295 | Transmembrane protein 59-like |
| TMEM74B | 1.762008443 | 14.58965216 | 8.280126136 | 0.000485474 | 0.013328471 | Transmembrane protein 74B |
| TMEM97 | 581.2389349 | 1852.179025 | 3.186605221 | 4.23E-12 | 1.83E-09 | Transmembrane protein 97 |
| TMOD4 | 2.783156696 | 37.21538475 | 13.37164551 | 1.24E-08 | 2.28E-06 | Tropomodulin-4 |
| TMPO | 1277.851297 | 615.8258144 | 0.481922909 | 0.000799848 | 0.019197327 | Thymopoietin zeta isoform |
| TMX2 | 1174.366641 | 704.3186182 | 0.599743379 | 0.000863883 | 0.020357528 | Thioredoxin-related transmembrane protein 2 |
| TMX4 | 970.0459478 | 1927.921254 | 1.987453542 | 5.64E-06 | 0.000395371 | Thioredoxin-related transmembrane protein 4 |
| TNC | 193.5610763 | 25.0943572 | 0.129645679 | 1.13E-07 | 1.50E-05 | Tenascin |
| TNFAIP8 | 317.0671271 | 148.3910484 | 0.468011458 | 0.000490837 | 0.013463832 | Tumor necrosis factor alpha-induced protein 8 |
| TNFAIP8L3 | 35.58085366 | 213.4828008 | 5.999934763 | 4.61E-08 | 7.13E-06 | |
| TNFRSF12A | 2338.628374 | 980.4480097 | 0.41924062 | 1.47E-05 | 0.00084355 | Tumor necrosis factor receptor superfamily member 12A |
| TNFRSF1B | 6.773739054 | 35.85533326 | 5.293285285 | 0.00037794 | 0.010966605 | Tumor necrosis factor receptor superfamily member 1B |
| TNFRSF25 | 931.0718471 | 511.5100038 | 0.549377586 | 0.000437749 | 0.012368724 | Tumor necrosis factor receptor superfamily member 25 |
| TNFSF4 | 339.8445768 | 131.3366846 | 0.386461028 | 4.98E-08 | 7.63E-06 | Tumor necrosis factor ligand superfamily member 4 |
| TNS1 | 12318.88029 | 23694.40866 | 1.923422267 | 1.71E-05 | 0.000953171 | Tensin-1 |
| TNS3 | 3196.10989 | 5431.392953 | 1.69937616 | 0.000244043 | 0.007754022 | Tensin-3 |
| TOM1L2 | 8463.964439 | 4890.317622 | 0.577780975 | 0.000263909 | 0.008217383 | cDNA FLJ60511, highly similar to Mus musculus target of myb1-like 2 (chicken) (Tom1l2), transcript variant 1, mRNA |
| TOMM40 | 1086.840048 | 599.1709861 | 0.551296382 | 0.000144878 | 0.005264689 | Mitochondrial import receptor subunit TOM40 homolog |
| TOP2A | 68.69575355 | 11.7674994 | 0.171298789 | 6.24E-05 | 0.002690069 | DNA topoisomerase 2-alpha |
| TP53I3 | 2524.644411 | 1173.321309 | 0.464747156 | 0.000213915 | 0.007019018 | Quinone oxidoreductase PIG3 |
| TPPP3 | 479.790853 | 1000.452155 | 2.085183886 | 4.38E-05 | 0.002028567 | Tubulin polymerization-promoting protein family member 3 |
| TPRG1 | 731.7111984 | 342.8905414 | 0.468614588 | 3.72E-05 | 0.001779569 | Tumor protein p63-regulated gene 1 protein |

| Gene name | Controls normalized mean counts | Simvastatin normalized mean counts | foldChange | pval | padj | Protein name |
|---|---|---|---|---|---|---|
| TRAF1 | 75.45409127 | 173.7441409 | 2.302647053 | 0.000723562 | 0.017891185 | TNF receptor-associated factor 1 |
| TRERF1 | 62.48629246 | 134.6460517 | 2.154809421 | 0.000164244 | 0.005785934 | Transcriptional-regulating factor 1 |
| TRIB3 | 551.3226244 | 1229.407533 | 2.229923966 | 2.45E-06 | 0.000193734 | Tribbles homolog 3 |
| TRIM25 | 529.207826 | 927.3973929 | 1.752425696 | 0.001258175 | 0.026892194 | E3 ubiquitin/ISG15 ligase TRIM25 |
| TRIM47 | 58.01460095 | 200.9471312 | 3.463733749 | 0.000515319 | 0.013973422 | Tripartite motif-containing protein 47 |
| TRIM5 | 1335.575849 | 664.6973461 | 0.497685958 | 2.55E-05 | 0.001296712 | Tripartite motif-containing protein 5 |
| TRIM59 | 161.1482494 | 72.11986451 | 0.447537375 | 0.000307957 | 0.009237524 | Tripartite motif-containing protein 59 |
| TRIM7 | 75.49201884 | 210.1537558 | 2.78378773 | 0.000356701 | 0.010447769 | Tripartite motif-containing protein 7 |
| TRMT10C | 721.6394401 | 423.2969943 | 0.586576857 | 0.000629724 | 0.016127975 | Mitochondrial ribonuclease P protein 1 |
| TRPM4 | 1323.176907 | 2327.670731 | 1.75915308 | 0.000297389 | 0.009007375 | cDNA FLJ61284, highly similar to Homo sapiens transient receptor potential cation channel, subfamily M, member 4 (TRPM4), mRNA |
| TRPM8 | 11.62200524 | 37.46509913 | 3.223634678 | 0.000347258 | 0.010200018 | TRPM8 protein |
| TSC22D1 | 1650.237004 | 2720.886776 | 1.648785459 | 0.000841664 | 0.019985349 | TSC22 domain family protein 1 |
| TSC22D3 | 1798.000426 | 10682.6768 | 5.941420618 | 1.68E-30 | 5.82E-27 | TSC22 domain family protein 3 |
| TSPAN12 | 762.7212916 | 403.1105092 | 0.528516135 | 0.000219402 | 0.007133846 | Tetraspanin-12 |
| TSPAN15 | 137.2154879 | 290.3652489 | 2.116125908 | 2.05E-05 | 0.001097397 | Tetraspanin-15 |
| TSPAN2 | 42.02910013 | 8.618347822 | 0.205056682 | 0.000580646 | 0.015254707 | Tetraspanin-2 |
| TSPAN32 | 5.792277641 | 28.9640665 | 5.00046239 | 0.000180504 | 0.006204143 | Tetraspanin-32 |
| TSPAN6 | 576.2709768 | 344.1481197 | 0.597198425 | 0.001076565 | 0.023919731 | Tetraspanin-6 |
| TSPAN9 | 8686.360517 | 4440.587971 | 0.511213869 | 0.000728109 | 0.017989321 | Tetraspanin-9 |
| TSPYL4 | 2018.178038 | 1088.835 | 0.539513848 | 3.51E-05 | 0.001700064 | Testis-specific Y-encoded-like protein 4 |
| TSSC1 | 564.3676718 | 337.1425055 | 0.597380967 | 0.001248138 | 0.026775563 | |
| TTBK2 | 470.4280314 | 957.3118767 | 2.034980513 | 3.77E-06 | 0.000278875 | Tau-tubulin kinase 2 |
| TTC18 | 106.297893 | 204.4214623 | 1.923099853 | 0.000524932 | 0.014137253 | Tetratricopeptide repeat protein 18 |
| TTL | 1500.095831 | 2842.454191 | 1.894848404 | 1.76E-05 | 0.000969071 | Tubulin--tyrosine ligase |
| TTLL3 | 340.6582078 | 663.1255887 | 1.946600944 | 2.32E-05 | 0.001207491 | Tubulin monoglycylase TTLL3 |
| TUSC3 | 951.6927524 | 539.8713907 | 0.567274879 | 0.0012472 | 0.026773919 | Tumor suppressor candidate 3 |
| TXLNB | 2504.467391 | 4692.661654 | 1.873716412 | 0.001029367 | 0.023118758 | Beta-taxilin |
| TXNL4B | 201.0017775 | 113.0656107 | 0.562510502 | 0.001418109 | 0.029310425 | Thioredoxin-like protein 4B |
| TYMP | 39.46694628 | 154.5040805 | 3.914771603 | 6.22E-06 | 0.000425282 | Thymidine phosphorylase |
| TYRO3 | 260.9832594 | 473.043183 | 1.812542246 | 0.000230212 | 0.007405343 | Tyrosine-protein kinase receptor TYRO3 |
| U91328.21 | 327.9805705 | 766.4081604 | 2.336748666 | 6.06E-07 | 6.18E-05 | |
| UBALD2 | 347.6068528 | 579.0230367 | 1.665741144 | 0.001465669 | 0.030053457 | |
| UBE2S | 905.4008263 | 548.4350667 | 0.605737316 | 0.001276802 | 0.027106916 | Ubiquitin-conjugating enzyme E2 S |
| UBE2V1 | 1953.962425 | 1006.140697 | 0.514923258 | 1.15E-05 | 0.000690149 | Ubiquitin-conjugating enzyme E2 variant 1 |
| UHRF1 | 384.5051594 | 102.1935465 | 0.265779389 | 0.00023924 | 0.007640445 | E3 ubiquitin-protein ligase UHRF1 |
| UNC119 | 1121.946173 | 544.4396208 | 0.485263584 | 9.34E-05 | 0.003721378 | Unc-119 homolog (C. elegans), isoform CRA_b |
| UNC13A | 34.93687987 | 115.4391669 | 3.304220851 | 0.000661291 | 0.016642503 | Protein unc-13 homolog A |

| Gene name | **Controls** normalized mean counts | **Simvastatin** normalized mean counts | **foldChange** | **pval** | **padj** | **Protein name** |
|---|---|---|---|---|---|---|
| UNC13D | 56.42202281 | 127.6139234 | 2.261775048 | 0.000255489 | 0.00802752 | Protein unc-13 homolog D |
| UPRT | 804.9233264 | 345.5173579 | 0.429254994 | 3.15E-07 | 3.53E-05 | Uracil phosphoribosyltransferase homolog |
| USP18 | 701.7032333 | 369.1627471 | 0.526095263 | 4.68E-05 | 0.002137683 | Ubl carboxyl-terminal hydrolase 18 |
| USP41 | 70.89326255 | 11.73684582 | 0.165556576 | 3.70E-06 | 0.000274398 | Putative ubiquitin carboxyl-terminal hydrolase 41 |
| USP54 | 559.7808925 | 1088.403119 | 1.944337746 | 4.15E-05 | 0.001952148 | Inactive ubiquitin carboxyl-terminal hydrolase 54 |
| UTP15 | 442.2118399 | 247.7329193 | 0.560213221 | 0.001260979 | 0.026921085 | U3 small nucleolar RNA-associated protein 15 homolog |
| VASH1 | 4045.836112 | 519.7168091 | 0.128457208 | 9.55E-16 | 8.03E-13 | Vasohibin-1 |
| VAX2 | 477.3674749 | 219.5936953 | 0.460009755 | 7.15E-06 | 0.000476466 | Ventral anterior homeobox 2 |
| VDR | 2804.48603 | 548.6400039 | 0.19562943 | 5.59E-08 | 8.45E-06 | cDNA FLJ51539, highly similar to Vitamin D3 receptor |
| VGLL2 | 4435.432439 | 1541.091306 | 0.347450069 | 0.002402026 | 0.043314446 | Transcription cofactor vestigial-like protein 2 |
| VGLL3 | 9905.650786 | 4584.555049 | 0.462822196 | 0.000575373 | 0.015141746 | Transcription cofactor vestigial-like protein 3 |
| VLDLR | 542.516143 | 1236.671066 | 2.279510171 | 8.46E-08 | 1.18E-05 | Very low-density lipoprotein receptor |
| VNN1 | 240.2316632 | 57.11185164 | 0.23773657 | 8.74E-09 | 1.64E-06 | Pantetheinase |
| VPS13C | 1516.411636 | 2383.169119 | 1.571584563 | 0.002264235 | 0.041430168 | Vacuolar protein sorting-associated protein 13C |
| VPS37B | 2156.676094 | 1130.580724 | 0.524223701 | 2.48E-05 | 0.001266873 | Vacuolar protein sorting-associated protein 37B |
| VSTM2L | 287.5538233 | 1539.9186 | 5.355236047 | 5.48E-07 | 5.74E-05 | V-set and transmembrane domain-containing protein 2-like protein |
| VWA2 | 102.1619582 | 27.91764065 | 0.273268457 | 1.19E-06 | 0.000110292 | von Willebrand factor A domain-containing protein 2 |
| VWA5A | 519.7502434 | 1278.731432 | 2.460280583 | 1.04E-05 | 0.000633781 | |
| VWF | 30.64425394 | 218.939584 | 7.14455586 | 6.89E-09 | 1.35E-06 | von Willebrand factor |
| WDR19 | 380.2757945 | 609.6311652 | 1.603129029 | 0.002320614 | 0.042337247 | cDNA FLJ57049, highly similar to WD repeat protein 19 |
| WDR31 | 107.2767391 | 201.9389091 | 1.882410957 | 0.002839233 | 0.049284142 | |
| WDR77 | 2324.312607 | 1360.812589 | 0.585468833 | 0.000364781 | 0.010648091 | cDNA FLJ55120, moderately similar to Methylosome protein 50 |
| WDR81 | 753.2898092 | 1333.537013 | 1.770284154 | 0.000113166 | 0.004313894 | WD repeat-containing protein 81 |
| WFDC1 | 10.41813595 | 52.64497044 | 5.053204404 | 0.000142945 | 0.005200532 | WAP four-disulfide core domain protein 1 |
| WFDC10B | 11.28225363 | 42.81120809 | 3.79456175 | 8.94E-05 | 0.003580375 | Protein WFDC10B |
| WIPI1 | 1911.493969 | 4052.353181 | 2.119992658 | 4.71E-07 | 5.04E-05 | WD repeat domain phosphoinositide-interacting protein 1 |
| WSCD1 | 572.8622418 | 996.0388679 | 1.73870574 | 0.000629959 | 0.016127975 | WSC domain-containing protein 1 |
| WWC2 | 1761.270718 | 1110.586361 | 0.630559714 | 0.002350274 | 0.04261916 | |
| XIRP2 | 1390.932317 | 6612.470072 | 4.753984066 | 8.66E-08 | 1.20E-05 | Xin actin-binding repeat-containing protein 2 |
| XKR8 | 218.0844828 | 421.1640012 | 1.931196552 | 5.45E-05 | 0.002443748 | XK-related protein 8 |
| XRCC4 | 897.3692851 | 424.0373562 | 0.472533842 | 6.04E-05 | 0.002649164 | DNA repair protein XRCC4 |
| XXbac-BPG299F13.17 | 40.20188652 | 101.903655 | 2.534797838 | 2.77E-05 | 0.001390079 | |
| YIPF7 | 439.4071983 | 771.4952092 | 1.755763702 | 0.000892481 | 0.020873321 | Protein YIPF7 |

| Gene name | Controls normalized mean counts | Simvastatin normalized mean counts | foldChange | pval | padj | Protein name |
|---|---|---|---|---|---|---|
| YPEL1 | 206.1513033 | 416.6645798 | 2.021159086 | 0.000120414 | 0.00451821 | Protein yippee-like 1 |
| YWHAH | 1161.710507 | 4551.567867 | 3.917988036 | 1.37E-19 | 2.03E-16 | 14-3-3 protein eta |
| ZBTB16 | 4.062614403 | 31.54136145 | 7.763808799 | 0.000192848 | 0.006512156 | Zinc finger and BTB domain-containing protein 16 |
| ZBTB4 | 1653.388806 | 2570.190282 | 1.554498417 | 0.002672958 | 0.046921569 | Zinc finger and BTB domain-containing protein 4 |
| ZC3H12C | 610.8368189 | 1203.268396 | 1.969868807 | 9.54E-06 | 0.000599305 | cDNA FLJ55575, moderately similar to Homo sapiens zinc finger CCCH-type containing 12A (ZC3H12A), mRNA |
| ZC3H6 | 166.2208671 | 322.427842 | 1.939755505 | 8.42E-05 | 0.003411992 | Zinc finger CCCH domain-containing protein 6 |
| ZDHHC15 | 159.5096632 | 70.96129812 | 0.444871469 | 2.60E-05 | 0.001315189 | Palmitoyltransferase |
| ZDHHC22 | 15.33285805 | 2.906622498 | 0.189568213 | 0.00142029 | 0.029316215 | Palmitoyltransferase ZDHHC22 |
| ZER1 | 1832.280398 | 2894.082746 | 1.57949774 | 0.001655765 | 0.032895175 | Protein zer-1 homolog |
| ZFAND5 | 4705.207059 | 7680.413822 | 1.632322175 | 0.000919142 | 0.02135038 | AN1-type zinc finger protein 5 |
| ZFP36L2 | 125.6035163 | 434.3527621 | 3.458125815 | 1.66E-05 | 0.000936421 | Zinc finger protein 36, C3H1 type-like 2 |
| ZFYVE21 | 563.1168581 | 891.3485695 | 1.582883831 | 0.002751373 | 0.048081014 | Zinc finger FYVE domain-containing protein 21 |
| ZIC4 | 19.31735206 | 45.4147129 | 2.350980236 | 0.002169401 | 0.040015561 | Zinc finger protein ZIC 4 |
| ZMIZ1 | 1033.36732 | 1681.200459 | 1.626914677 | 0.0010401 | 0.023275782 | Zinc finger MIZ domain-containing protein 1 |
| ZNF146 | 967.3101898 | 555.0307382 | 0.573787751 | 0.000330159 | 0.009808916 | Zinc finger protein OZF |
| ZNF192P1 | 57.1403785 | 132.8828081 | 2.325550015 | 0.000249143 | 0.00789191 | |
| ZNF239 | 130.7200689 | 45.55730565 | 0.348510416 | 3.01E-07 | 3.41E-05 | Zinc finger protein 239 |
| ZNF280B | 434.393124 | 173.8934978 | 0.400313652 | 9.64E-06 | 0.000602271 | Zinc finger protein 280B |
| ZNF280C | 199.5394938 | 115.8176522 | 0.580424707 | 0.00273851 | 0.047936428 | Zinc finger protein 280C |
| ZNF300 | 254.3107188 | 133.1316561 | 0.52349998 | 0.000173288 | 0.006015964 | Zinc finger protein 300 |
| ZNF33A | 422.5352477 | 820.3550482 | 1.94150678 | 2.12E-05 | 0.001125635 | Zinc finger protein 33A |
| ZNF365 | 90.73007603 | 178.3153195 | 1.965338587 | 0.000890097 | 0.020853427 | Protein ZNF365 |
| ZNF385A | 58.94391129 | 126.0449502 | 2.138387959 | 0.001047112 | 0.023399049 | Zinc finger protein 385A |
| ZNF395 | 138.7566902 | 440.8735222 | 3.177313624 | 5.14E-11 | 1.79E-08 | Zinc finger protein 395 |
| ZNF436 | 345.5127107 | 767.1318179 | 2.220270902 | 0.000524011 | 0.014136931 | Zinc finger protein 436 |
| ZNF449 | 207.3841482 | 360.1972984 | 1.736860322 | 0.000748434 | 0.018345784 | Zinc finger protein 449 |
| ZNF462 | 456.9687414 | 164.5669179 | 0.360127298 | 5.98E-09 | 1.21E-06 | Zinc finger protein 462 |
| ZNF471 | 347.9740465 | 579.6997935 | 1.665928247 | 0.001379298 | 0.028679451 | cDNA FLJ59573, moderately similar to Zinc finger protein 471 |
| ZNF503 | 210.0835324 | 101.2397355 | 0.481902291 | 6.11E-05 | 0.002663968 | Zinc finger protein 503 |
| ZNF514 | 144.8390812 | 271.7029114 | 1.875895022 | 0.001285331 | 0.027235352 | Zinc finger protein 514 |
| ZNF517 | 111.4444055 | 214.9432892 | 1.928704167 | 0.000231185 | 0.007421302 | Zinc finger protein 517 |
| ZNF542 | 391.7643082 | 183.9075237 | 0.469434095 | 5.78E-06 | 0.000400997 | Putative zinc finger protein 542 |
| ZNF622 | 1985.182206 | 1239.367146 | 0.624309014 | 0.001724576 | 0.033540427 | Zinc finger protein 622 |
| ZNF655 | 1052.204903 | 610.1927745 | 0.579918201 | 0.000625281 | 0.016047837 | Zinc finger protein 655 |
| ZNF699 | 388.9772953 | 148.9339404 | 0.382885948 | 0.00035415 | 0.010392639 | Zinc finger protein 699 |
| ZNF703 | 453.5388712 | 1061.177447 | 2.339771769 | 4.19E-08 | 6.65E-06 | Zinc finger protein 703 |
| ZNF705G | 351.5931465 | 28.91208736 | 0.082231658 | 0.001034024 | 0.023193363 | Putative zinc finger protein 705G |
| ZNF711 | 281.5036433 | 133.6702724 | 0.474843845 | 9.69E-05 | 0.003824035 | Zinc finger protein 711 |
| ZNF862 | 277.9189845 | 457.9008104 | 1.647605367 | 0.001933664 | 0.036387505 | Zinc finger protein 862 |
| ZRANB1 | 1298.416409 | 2052.449783 | 1.58073309 | 0.001766307 | 0.033978197 | Ubiquitin thioesterase ZRANB1 |
| ZYX | 5441.485475 | 2802.487079 | 0.515022431 | 1.00E-05 | 0.00061844 | Zyxin |

Supplementary Table S3. Differentially expressed genes at RNA-level from rosuvastatin-treated primary human myotubes (FDR<0.05)

| Gene name | Controls normalized mean counts | Rosuvastatin normalized mean counts | foldChange | pval | padj | Protein name |
|---|---|---|---|---|---|---|
| ABCG1 | 138.7478852 | 48.68837108 | 0.350912528 | 2.82566E-07 | 0.000242756 | ATP-binding cassette sub-family G member 1 |
| AC010970.2 | 1182.337019 | 26537.0642 | 22.44458541 | 3.94518E-72 | 6.10083E-68 | |
| AC093388.3 | 5525.579029 | 11996.09997 | 2.171012288 | 8.10E-06 | 0.005216046 | |
| ACAT2 | 1064.936878 | 4111.547115 | 3.860836449 | 2.54045E-10 | 3.92855E-07 | Acetyl-CoA acetyltransferase, cytosolic |
| ACLY | 4858.020226 | 8604.21452 | 1.771136002 | 1.68E-05 | 0.010411877 | cDNA FLJ55447, highly similar to ATP-citrate synthase (EC 2.3.3.8) |
| ACSS2 | 664.1220082 | 1843.596227 | 2.775990261 | 1.11069E-09 | 1.41095E-06 | Acetyl-coenzyme A synthetase, cytoplasmic |
| ANKRD30BL | 10121.40241 | 208273.1356 | 20.57749777 | 2.74875E-69 | 2.83377E-65 | |
| C14orf1 | 205.5251562 | 716.9780251 | 3.488517115 | 4.41E-13 | 1.04991E-09 | Probable ergosterol biosynthetic protein 28 |
| CDHR1 | 133.7425541 | 1189.765868 | 8.895940979 | 3.33091E-13 | 8.9712E-10 | Cadherin-related family member 1 |
| CENPI | 62.99448449 | 135.7300572 | 2.154633986 | 6.66614E-05 | 0.033253265 | Centromere protein I |
| CTD-2328D6.1 | 11606.69888 | 24752.11441 | 2.132571427 | 1.33353E-05 | 0.00841703 | |
| CTD-2636A23.2 | 82.66326051 | 237.24156 | 2.869975834 | 9.10338E-05 | 0.043315266 | |
| CYP51A1 | 1586.891945 | 4559.494552 | 2.873223074 | 3.38E-14 | 1.04659E-10 | Lanosterol 14-alpha demethylase |
| DHCR24 | 2540.165571 | 9176.576659 | 3.612589968 | 3.48E-13 | 8.97E-10 | cDNA FLJ53870, highly similar to 24-dehydrocholesterol reductase (EC1.3.1.-) |
| DHCR7 | 1753.126494 | 4579.788796 | 2.61235502 | 5.42E-10 | 7.50E-07 | 7-dehydrocholesterol reductase |
| EBP | 479.1687988 | 1217.637684 | 2.541145599 | 1.43485E-09 | 1.70681E-06 | 3-beta-hydroxysteroid-Delta(8),Delta(7)-isomerase |
| ELOVL6 | 485.0140297 | 1111.187235 | 2.291041428 | 9.4443E-09 | 1.04319E-05 | Elongation of very long chain fatty acids protein 6 |
| FABP3 | 9386.496306 | 18603.0356 | 1.981893455 | 9.80453E-05 | 0.045944601 | Fatty acid-binding protein, heart |
| FDFT1 | 5469.109157 | 14033.62533 | 2.565980112 | 3.31337E-10 | 4.87981E-07 | cDNA FLJ33164 fis, clone UTERU2000542, highly similar to Squalene synthetase (EC 2.5.1.21) |
| FDPS | 9684.352645 | 24184.32169 | 2.497257439 | 4.22E-06 | 0.003107623 | Farnesyl pyrophosphate synthase |
| GPR20 | 4.774134804 | 27.6185635 | 5.785040564 | 2.56E-05 | 0.014726735 | G-protein coupled receptor 20 |
| GPR85 | 83.78629075 | 201.8628334 | 2.409258502 | 2.80652E-06 | 0.002170004 | Probable G-protein-coupled receptor 85 |
| HFM1 | 46547.60798 | 94742.51865 | 2.035389631 | 3.95E-05 | 0.020811386 | Probable ATP-dependent DNA helicase HFM1 |
| HMGCR | 2286.817119 | 6726.493963 | 2.941421903 | 1.76424E-08 | 1.70514E-05 | 3-hydroxy-3-methylglutaryl-coenzyme A reductase |
| HMGCS1 | 5774.386649 | 22593.23366 | 3.912663808 | 1.75774E-08 | 1.70514E-05 | Hydroxymethylglutaryl-CoA synthase, cytoplasmic |
| HSD17B7 | 218.4991005 | 580.3248778 | 2.655960031 | 1.14052E-09 | 1.41095E-06 | 3-keto-steroid reductase |
| IDI1 | 5165.583591 | 12847.56104 | 2.487146091 | 5.57912E-10 | 7.50222E-07 | Isopentenyl-diphosphate Delta-isomerase 1 |
| INSIG1 | 2474.908904 | 6176.492882 | 2.495644536 | 2.69773E-05 | 0.015170083 | Insulin-induced gene protein |
| ITIH1 | 3.902578423 | 118.6985637 | 30.41542048 | 1.13E-07 | 1.00E-04 | Inter-alpha-trypsin inhibitor heavy chain H1 |
| KIF1A | 148.8518228 | 391.1984222 | 2.62810636 | 7.73345E-06 | 0.005199572 | Kinesin-like protein KIF1A |

| Gene name | Controls normalized mean counts | Rosuvastatin normalized mean counts | foldChange | pval | padj | Protein name |
|---|---|---|---|---|---|---|
| LDHD | 41.62262255 | 106.0011671 | 2.546720043 | 2.35204E-05 | 0.014188937 | Probable D-lactate dehydrogenase, mitochondrial |
| LDLR | 2422.412654 | 5982.836237 | 2.469784092 | 9.24812E-08 | 8.41253E-05 | Low-density lipoprotein receptor |
| LINC00578 | 90.94656948 | 275.7969393 | 3.032516134 | 7.44248E-05 | 0.036536685 | |
| LPIN1 | 1272.716481 | 2461.057609 | 1.933704518 | 4.25462E-05 | 0.021931164 | Phosphatidate phosphatase LPIN1 |
| LSS | 3898.702937 | 10872.63607 | 2.788782896 | 1.23E-10 | 2.00E-07 | Lanosterol synthase |
| LUC7L2 | 50.16147423 | 16.13287188 | 0.321618775 | 2.39E-05 | 0.014188937 | Putative RNA-binding protein Luc7-like 2 |
| MIR663A | 221.4538393 | 861.2448224 | 3.889048956 | 1.88E-06 | 1.53E-03 | |
| MSMO1 | 3306.127494 | 9929.671565 | 3.003414594 | 1.58E-11 | 2.88E-08 | Methylsterol monooxygenase 1 |
| MVD | 2080.057217 | 10021.38283 | 4.817839984 | 5.9961E-19 | 2.06053E-15 | Diphosphomevalonate decarboxylase |
| MVK | 669.4153473 | 1515.027268 | 2.263209641 | 4.62774E-06 | 0.003328529 | Mevalonate kinase (Mevalonic aciduria), isoform CRA_a |
| NSDHL | 829.6917563 | 2197.956151 | 2.649123768 | 4.28698E-12 | 8.28672E-09 | Sterol-4-alpha-carboxylate 3-dehydrogenase, decarboxylating |
| P2RY6 | 38.24712052 | 88.8489559 | 2.323023399 | 8.1157E-05 | 0.039219097 | P2Y purinoceptor 6 |
| PCYT2 | 913.7058326 | 1759.18786 | 1.925332856 | 7.04235E-06 | 0.004840132 | Ethanolamine-phosphate cytidylyltransferase |
| PDE3A | 1006.181435 | 19334.94578 | 19.21616232 | 7.77356E-66 | 6.01051E-62 | cGMP-inhibited 3',5'-cyclic phosphodiesterase A |
| PEG10 | 2253.344831 | 4553.969673 | 2.020982146 | 3.9701E-05 | 0.020811386 | Retrotransposon-derived protein PEG10 |
| PLA2G3 | 35.84308641 | 311.5735133 | 8.692708819 | 2.97761E-06 | 0.002246134 | Group 3 secretory phospholipase A2 |
| RAB33A | 6.059651694 | 34.16203733 | 5.637623918 | 6.07E-06 | 4.27E-03 | Ras-related protein Rab-33A |
| RMRP | 2.741746892 | 154.3091441 | 56.28132362 | 7.31658E-38 | 3.77145E-34 | |
| RN7SK | 24.06731186 | 460.6775993 | 19.14121536 | 4.72685E-45 | 2.92384E-41 | |
| RN7SL2 | 114.702405 | 1030.284013 | 8.982235489 | 6.67246E-11 | 1.14648E-07 | |
| RNA5-8SP6 | 133.8857898 | 6487.839015 | 48.45801054 | 2.5348E-99 | 7.83975E-95 | |
| RP11-613D13.5 | 1105.300328 | 1913.204419 | 1.730936263 | 6.63918E-05 | 0.033253265 | |
| RP11-903H12.3 | 12.52418387 | 39.86502214 | 3.183043507 | 1.01E-04 | 0.046527088 | |
| RPPH1 | 5.740941755 | 133.3021685 | 23.21956469 | 2.02969E-26 | 8.96775E-23 | |
| RPTN | 19.3314885 | 57.81197599 | 2.990559987 | 1.63911E-06 | 0.001370121 | Repetin |
| SC5D | 2061.544089 | 3787.659416 | 1.837292463 | 7.99077E-06 | 0.005216046 | Lathosterol oxidase |
| SLC29A2 | 163.2396168 | 318.000853 | 1.948061746 | 3.79832E-05 | 0.02060957 | Solute carrier family 29 (Nucleoside transporters), member 2, isoform CRA_d |
| SLC2A6 | 394.6721725 | 788.4401852 | 1.997709086 | 2.64E-06 | 2.09E-03 | Solute carrier family 2, facilitated glucose transporter member 6 |
| SLCO4C1 | 18.17890467 | 235.1459111 | 12.93509787 | 3.51E-12 | 7.24123E-09 | Solute carrier organic anion transporter family member 4C1 |
| SLCO5A1 | 25337.05006 | 51971.89451 | 2.051221211 | 3.26E-05 | 1.80E-02 | Solute carrier organic anion transporter family member 5A1 |
| SNAI3-AS1 | 187.1640524 | 492.8956621 | 2.633495353 | 1.58266E-08 | 1.63161E-05 | |
| SNORD3A | 3.437735916 | 24.09131611 | 7.007901913 | 2.57127E-05 | 0.014726735 | |
| SQLE | 1118.613033 | 2632.955052 | 2.353767545 | 2.86E-09 | 3.27761E-06 | Squalene monooxygenase |
| STARD4 | 1382.749885 | 3389.159783 | 2.451028794 | 1.05E-08 | 1.1246E-05 | StAR-related lipid transfer protein 4 |
| TM7SF2 | 349.0162126 | 1109.615297 | 3.179265767 | 3.00E-12 | 6.6342E-09 | Delta(14)-sterol reductase |

| Gene name | Controls normalized mean counts | Rosuvastatin normalized mean counts | foldChange | pval | padj | Protein name |
|---|---|---|---|---|---|---|
| TMEM97 | 581.2389349 | 3258.604654 | 5.606308281 | 1.42E-25 | 5.51E-22 | Transmembrane protein 97 |
| TOP2A | 68.69575355 | 18.74356972 | 0.27284903 | 1.08E-04 | 4.91E-02 | DNA topoisomerase 2-alpha |
| VWF | 30.64425394 | 222.6817284 | 7.266671554 | 3.12E-08 | 2.92295E-05 | von Willebrand factor |

Supplementary Table S4. Protein coding transcript variants differentially expressed for at least one of the statin treatments in human primary myotubes.

| Feature ID | Transcript ID | UniProt protein ID | **Controls vs Simvastatin** P-value | **Controls vs Simvastatin** Fold change | **Controls vs Simvastatin** FDR p-value | **Controls vs Rosuvastatin** P-value | **Controls vs Rosuvastatin** Fold change | **Controls vs Rosuvastatin** FDR p-value | Control 1 Total transcript reads | Control 2 Total transcript reads | Control 3 Total transcript reads | Control 4 Total transcript reads | Simvastatin 1 Total transcript reads | Simvastatin 2 Total transcript reads | Simvastatin 3 Total transcript reads | Simvastatin 4 Total transcript reads | Rosuvastatin 1 Total transcript reads | Rosuvastatin 2 Total transcript reads | Rosuvastatin 3 Total transcript reads | Rosuvastatin 4 Total transcript reads |
|---|---|---|---|---|---|---|---|---|---|---|---|---|---|---|---|---|---|---|---|---|
| HMGCR_1 | ENST00000287936 | P04035 | 3.36749E-06 | 2.433430181 | 0.00240361 | 1.93E-09 | 3.115096027 | 2.57432E-05 | 1565 | 2794 | 1519 | 1563 | 3885 | 6455 | 3015 | 4444 | 7410 | 8224 | 3626 | 5142 |
| HMGCR_13 | ENST00000343975 | P04035 | 0.034220124 | 2.484450281 | 1 | 0.049068173 | 2.355005058 | 1 | 40 | 188 | 204 | 100 | 213 | 553 | 393 | 202 | 220 | 446 | 451 | 204 |
| LDLR_10 | ENST00000558518 | P01130 | 0.009514698 | 2.566483179 | 1 | 0.135573308 | 1.699352706 | 1 | 379 | 556 | 96 | 106 | 534 | 1185 | 195 | 970 | 658 | 577 | 350 | 386 |
| LDLR_7 | ENST00000558013 | P01130 | 0.000113457 | 3.624581565 | 0.039916158 | 0.00086141 | 3.081628722 | 1 | 90 | 397 | 98 | 197 | 1030 | 329 | 549 | 861 | 1030 | 739 | 305 | 516 |
| LDLR_4 | ENST00000535915 | P01130 | 0.044513608 | 2.217466699 | 1 | 0.010063786 | 2.705647472 | 1 | 182 | 97 | 83 | 181 | 135 | 330 | 208 | 584 | 324 | 406 | 251 | 628 |
| LDLR_8 | ENST00000545707 | P01130 | 0.019095917 | 1.769959496 | 1 | 2.37804E-05 | 2.579406724 | 0.082492331 | 294 | 493 | 423 | 286 | 767 | 572 | 489 | 754 | 828 | 1441 | 922 | 829 |
| LDLR_2 | ENST00000557933 | H0YMD1 | 0.017150341 | 3.207146641 | 1 | 0.001317772 | 4.184779005 | 1 | 94 | 86 | 91 | 63 | 273 | 306 | 116 | 425 | 338 | 675 | 173 | 396 |
| LDLR_13 | ENST00000560467 | H0YM92 | 0.000260201 | 2.870292149 | 0.074719808 | 1.29222E-05 | 3.557701867 | 0.046693677 | 101 | 56 | 97 | 32 | 232 | 211 | 156 | 213 | 364 | 358 | 215 | 144 |
| PCSK9_1 | ENST00000302118 | Q8NBP7 | 0.755200327 | 1.154694263 | 1 | 0.008298369 | 2.223215634 | 1 | 290 | 204 | 633 | 134 | 297 | 473 | 273 | 411 | 1089 | 714 | 635 | 541 |

Supplementary Table S5. Differentially expressed proteins from simvastatin-treated primary human myotubes

| Protein.names | Gene.names | Simvastatin (Ratio.H.L.normalized) |
|---|---|---|
| Zinc finger protein 292 | ZNF292 | 0.011419 |
| Bisphosphoglycerate mutase | BPGM | 0.061013 |
| Actin-related protein 3B | ACTR3B | 0.11137 |
| E3 ubiquitin-protein ligase listerin | LTN1 | 0.12766 |
| Transient receptor potential cation channel subfamily V member 2 | TRPV2 | 0.13894 |
| Ubiquitin Specific Peptidase 39 | USP39 | 0.15494 |
| Baculoviral IAP repeat-containing protein 6 | BIRC6 | 0.18097 |
| Glycerate kinase | GLYCTK | 0.19235 |
| Dipeptidyl peptidase 1;Dipeptidyl peptidase 1 exclusion domain chain;Dipeptidyl peptidase 1 heavy chain;Dipeptidyl peptidase 1 light chain | CTSC | 0.19786 |
| Nucleotide exchange factor SIL1 | SIL1 | 0.20189 |
| WD repeat-containing protein 75 | WDR75 | 0.21071 |
| Protein lin-7 homolog C;Protein lin-7 homolog A | LIN7C;LIN7A | 0.23254 |
| Actin, aortic smooth muscle;Actin, gamma-enteric smooth muscle | ACTA2;ACTG2 | 0.25349 |
| Host cell factor 1;HCF N-terminal chain 1;HCF N-terminal chain 2;HCF N-terminal chain 3;HCF N-terminal chain 4;HCF N-terminal chain 5;HCF N-terminal chain 6;HCF C-terminal chain 1;HCF C-terminal chain 2;HCF C-terminal chain 3;HCF C-terminal chain 4;HCF C-terminal chain 5;HCF C-terminal chain 6 | HCFC1 | 0.26369 |
| Conserved oligomeric Golgi complex subunit 7 | COG7 | 0.26838 |
| RNA-binding protein 12 | RBM12 | 0.27592 |
| Angio-associated migratory cell protein | AAMP | 0.28235 |
| Centrosomal protein of 170 kDa | CEP170 | 0.28318 |
| Uncharacterized protein C1orf198 | C1orf198 | 0.28357 |
| Group XV phospholipase A2 | PLA2G15 | 0.28664 |
| Myosin-6 | MYH6 | 0.28888 |
| Protein C10 | C12orf57 | 0.29716 |
| Myosin-1 | MYH1 | 0.31068 |
| Uroporphyrinogen decarboxylase | UROD | 0.31083 |
| Transportin-3 | TNPO3 | 0.31368 |
| Alpha/beta hydrolase domain-containing protein 11 | ABHD11 | 0.32104 |
| GDP-Man:Man(3)GlcNAc(2)-PP-Dol alpha-1,2-mannosyltransferase | ALG11 | 0.32198 |
| Myosin-2 | MYH2 | 0.32317 |
| cGMP-dependent 3,5-cyclic phosphodiesterase | PDE2A | 0.32506 |
| Uncharacterized protein C1orf50 | C1orf50 | 0.34367 |
| Suppressor of SWI4 1 homolog | PPAN | 0.34573 |
| Titin | TTN | 0.34961 |
| Tubulin beta-2B chain | TUBB2B | 0.35348 |
| Breast carcinoma-amplified sequence 1 | BCAS1 | 0.3551 |
| WD repeat and FYVE domain-containing protein 3 | WDFY3 | 0.36039 |
| RNA-binding protein 42 | RBM42 | 0.36075 |
| Cell division cycle and apoptosis regulator protein 1 | CCAR1 | 0.36535 |
| Ras-specific guanine nucleotide-releasing factor 2 | RASGRF2 | 0.36607 |
| CD276 antigen | CD276 | 0.36763 |
| EF-hand domain-containing protein D1 | EFHD1 | 0.36825 |
| Inositol monophosphatase 3 | IMPAD1 | 0.37054 |
| Porphobilinogen deaminase | HMBS | 0.37192 |
| Epidermal growth factor receptor kinase substrate 8-like protein 2 | EPS8L2 | 0.37823 |
| MMS19 nucleotide excision repair protein homolog | MMS19 | 0.38107 |
| Junctophilin-2 | JPH2 | 0.38398 |
| Obscurin-like protein 1 | OBSL1 | 0.38442 |
| Transcription activator BRG1;Probable global transcription activator SNF2L2 | SMARCA4;SMARCA2 | 0.3858 |
| RNA polymerase II-associated protein 3 | RPAP3 | 0.3883 |
| Integrin beta-1-binding protein 2 | ITGB1BP2 | 0.39459 |
| Integrin beta;Integrin beta-8 | ITGB8 | 0.39703 |
| Calsyntenin-2 | CLSTN2 | 0.39749 |
| Lysosomal alpha-glucosidase;76 kDa lysosomal alpha-glucosidase;70 kDa lysosomal alpha-glucosidase | GAA | 0.40241 |
| 28S ribosomal protein S35, mitochondrial | MRPS35 | 0.40296 |
| Ancient ubiquitous protein 1 | AUP1 | 0.40335 |
| Vacuolar protein sorting-associated protein 37B | VPS37B | 0.40485 |
| Trimethyllysine dioxygenase, mitochondrial | TMLHE | 0.41198 |
| N-acetylgalactosamine-6-sulfatase | GALNS | 0.4121 |
| RNA polymerase I-specific transcription initiation factor RRN3;Putative RRN3-like protein RRN3P1 | RRN3;RRN3P1 | 0.41254 |

| Protein.names | Gene.names | Simvastatin (Ratio.H.L.normalized) |
|---|---|---|
| Protein LSM12 homolog | LSM12 | 0.41257 |
| Dystrobrevin alpha | DTNA | 0.41305 |
| Ras-related protein Rab-24 | RAB24 | 0.41383 |
| Troponin I, fast skeletal muscle | TNNI2 | 0.41598 |
| Myosin light chain 1/3, skeletal muscle isoform | MYL1 | 0.41839 |
| Ras association domain-containing protein 4 | RASSF4 | 0.42751 |
| NFX1-type zinc finger-containing protein 1 | ZNFX1 | 0.43639 |
| Mitochondrial import inner membrane translocase subunit Tim23;Putative mitochondrial import inner membrane translocase subunit Tim23B | TIMM23;TIMM23B | 0.44209 |
| HEAT repeat-containing protein 3 | HEATR3 | 0.44254 |
| Vacuolar protein sorting-associated protein 11 homolog | VPS11 | 0.4465 |
| 39S ribosomal protein L46, mitochondrial | MRPL46 | 0.44703 |
| Interferon regulatory factor 2-binding protein-like | IRF2BPL | 0.45466 |
| Protein pelota homolog | PELO | 0.45778 |
| Sarcalumenin | SRL | 0.45975 |
| Conserved oligomeric Golgi complex subunit 5 | COG5 | 0.46137 |
| Tripeptidyl-peptidase 1 | TPP1 | 0.46374 |
| Chromodomain-helicase-DNA-binding protein 4 | CHD4 | 0.46595 |
| AP2-associated protein kinase 1 | AAK1 | 0.46606 |
| StAR-related lipid transfer protein 13 | STARD13 | 0.46666 |
| Niemann-Pick C1 protein | NPC1 | 0.46701 |
| Tumor protein p53-inducible protein 11 | TP53I11 | 0.46731 |
| Protein sel-1 homolog 1 | SEL1L | 0.47029 |
| ATP-dependent Clp protease ATP-binding subunit clpX-like, mitochondrial | CLPX | 0.47244 |
| GRAM domain-containing protein 3 | GRAMD3 | 0.473 |
| L-aminoadipate-semialdehyde dehydrogenase-phosphopantetheinyl | AASDHPPT | 0.4738 |
| Serine/arginine-rich splicing factor 11 | SRSF11 | 0.47396 |
| Probable ATP-dependent RNA helicase DDX46 | DDX46 | 0.47768 |
| Nucleolar protein 6 | NOL6 | 0.47941 |
| Integrator complex subunit 2 | INTS2 | 0.48434 |
| Lysosomal acid phosphatase | ACP2 | 0.48459 |
| Coiled-coil-helix-coiled-coil-helix domain-containing protein 3, mitochondrial | CHCHD3 | 0.4846 |
| Transmembrane 9 superfamily member 4 | TM9SF4 | 0.48474 |
| LanC-like protein 2 | LANCL2 | 0.4863 |
| NudC domain-containing protein 1 | NUDCD1 | 0.48908 |
| Kelch-like protein 31 | KLHL31 | 0.4903 |
| 1-acyl-sn-glycerol-3-phosphate acyltransferase gamma | AGPAT3 | 0.49041 |
| Cleft lip and palate transmembrane protein 1-like protein | CLPTM1L | 0.49122 |
| SAM domain and HD domain-containing protein 1 | SAMHD1 | 0.49175 |
| Myosin-binding protein H | MYBPH | 0.49201 |
| Mitotic checkpoint protein BUB3 | BUB3 | 0.49346 |
| Methionine aminopeptidase;Methionine aminopeptidase 1 | METAP1 | 0.49411 |
| Ras-related protein Rab-12 | RAB12 | 0.49589 |
| Leucine carboxyl methyltransferase 1 | LCMT1 | 0.50351 |
| Pescadillo homolog | PES1 | 0.5044 |
| HD domain-containing protein 2 | HDDC2 | 0.51347 |
| Myosin-3 | MYH3 | 0.51497 |
| NudC domain-containing protein 3 | NUDCD3 | 0.51709 |
| Negative elongation factor B | NELFB | 0.51933 |
| Serine/threonine-protein kinase ULK3 | ULK3 | 0.52114 |
| ELAV-like protein 2;ELAV-like protein 4 | ELAVL2;ELAVL4 | 0.52148 |
| Neurofibromin;Neurofibromin truncated | NF1 | 0.52158 |
| SCY1-like protein 2 | SCYL2 | 0.52166 |
| E3 ubiquitin-protein ligase UBR1 | UBR1 | 0.52378 |
| Serine/threonine-protein kinase SIK3 | SIK3;KIAA0999 | 0.526 |
| Splicing factor 1 | SF1 | 0.52713 |
| ATP-binding cassette sub-family B member 7, mitochondrial | ABCB7 | 0.52741 |
| La-related protein 4 | LARP4 | 0.53013 |
| Cadherin-15 | CDH15 | 0.53035 |
| ATP-dependent RNA helicase DHX8 | DHX8 | 0.53045 |
| Cleavage stimulation factor subunit 3 | CSTF3 | 0.53315 |
| Ribonucleases P/MRP protein subunit POP1 | POP1 | 0.53406 |
| ATP-dependent RNA helicase DDX50 | DDX50 | 0.53434 |
| Guanidinoacetate N-methyltransferase | GAMT | 0.53578 |

| Protein.names | Gene.names | Simvastatin (Ratio.H.L.normalized) |
|---|---|---|
| Myosin light chain 3 | MYL3 | 0.53859 |
| WD repeat-containing protein 18 | WDR18 | 0.53868 |
| Coagulation factor XIII A chain | F13A1 | 0.53888 |
| DNA topoisomerase 2;DNA topoisomerase 2-beta | TOP2B | 0.53947 |
| Probable ATP-dependent RNA helicase DDX5 | DDX5 | 0.54572 |
| Ribosomal RNA processing protein 1 homolog A | RRP1 | 0.5478 |
| Ankyrin repeat domain-containing protein 2 | ANKRD2 | 0.55037 |
| Pre-mRNA-processing factor 6 | PRPF6 | 0.55077 |
| 5-3 exoribonuclease 1 | XRN1 | 0.55097 |
| Myomesin-3 | MYOM3 | 0.55109 |
| Eukaryotic translation initiation factor 3 subunit K | EIF3K | 0.55158 |
| WD repeat-containing protein C2orf44 | C2orf44 | 0.55331 |
| CDKN2A-interacting protein | CDKN2AIP | 0.55465 |
| Putative heat shock protein HSP 90-beta 2 | HSP90AB2P | 0.55482 |
| Translation initiation factor eIF-2B subunit delta | EIF2B4 | 0.55812 |
| ATP-dependent zinc metalloprotease YME1L1 | YME1L1 | 0.55814 |
| Casein kinase I isoform alpha | CSNK1A1 | 0.55941 |
| Nucleolar RNA helicase 2 | DDX21 | 0.56133 |
| Peptidyl-prolyl cis-trans isomerase-like 4 | PPIL4 | 0.56162 |
| 28S ribosomal protein S36, mitochondrial | MRPS36 | 0.5626 |
| ATPase family AAA domain-containing protein 1 | ATAD1 | 0.56372 |
| Golgin subfamily A member 2 | GOLGA2 | 0.56372 |
| RAC-beta serine/threonine-protein kinase | AKT2 | 0.56389 |
| 39S ribosomal protein L39, mitochondrial | MRPL39 | 0.57011 |
| Calcium-binding mitochondrial carrier protein Aralar2 | SLC25A13 | 0.57067 |
| 28S ribosomal protein S15, mitochondrial | MRPS15 | 0.57266 |
| tRNA (cytosine(34)-C(5))-methyltransferase | NSUN2 | 0.57272 |
| TBC1 domain family member 13 | TBC1D13 | 0.57288 |
| Tumor protein D53 | TPD52L1 | 0.57311 |
| Myosin-4 | MYH4 | 0.57363 |
| OCIA domain-containing protein 1 | OCIAD1 | 0.57369 |
| SH3 domain-binding glutamic acid-rich protein | SH3BGR | 0.57379 |
| Microtubule-associated protein RP/EB family member 3 | MAPRE3 | 0.57394 |
| Folate receptor alpha | FOLR1 | 0.57458 |
| BAG family molecular chaperone regulator 5 | BAG5 | 0.57529 |
| Diphthine synthase | DPH5 | 0.57529 |
| Death-associated protein kinase 3 | DAPK3 | 0.57583 |
| WD repeat-containing protein 82 | WDR82 | 0.57616 |
| 39S ribosomal protein L50, mitochondrial | MRPL50 | 0.57643 |
| Myosin regulatory light chain 2, skeletal muscle isoform | MYLPF | 0.57651 |
| Echinoderm microtubule-associated protein-like 1 | EML1 | 0.57717 |
| Ubiquinone biosynthesis protein COQ9, mitochondrial | COQ9 | 0.57746 |
| MAP7 domain-containing protein 1 | MAP7D1 | 0.57886 |
| Mitogen-activated protein kinase 12 | MAPK12 | 0.57951 |
| U2 snRNP-associated SURP motif-containing protein | U2SURP | 0.58009 |
| Ribosomal L1 domain-containing protein 1 | RSL1D1 | 0.58016 |
| Myosin-8 | MYH8 | 0.58107 |
| Sorbin and SH3 domain-containing protein 2 | SORBS2 | 0.58191 |
| Sortilin | SORT1 | 0.58244 |
| Zinc finger CCCH domain-containing protein 15 | ZC3H15 | 0.58671 |
| Aldehyde dehydrogenase family 16 member A1 | ALDH16A1 | 0.58836 |
| Platelet glycoprotein 4 | CD36 | 0.59306 |
| CCA tRNA nucleotidyltransferase 1, mitochondrial | TRNT1 | 0.59395 |
| Phytanoyl-CoA dioxygenase, peroxisomal | PHYH | 0.59476 |
| N-acetyltransferase 10 | NAT10 | 0.59478 |
| Gamma-aminobutyric acid receptor-associated protein-like 1 | GABARAPL1 | 0.59487 |
| 28S ribosomal protein S9, mitochondrial | MRPS9 | 0.59646 |
| Phospholysine phosphohistidine inorganic pyrophosphate phosphatase | LHPP | 0.5976 |
| Interferon-inducible double stranded RNA-dependent protein kinase activator A | PRKRA | 0.59771 |
| DNA-directed RNA polymerases I, II, and III subunit RPABC5 | POLR2L | 0.59787 |
| Inositol polyphosphate 1-phosphatase | INPP1 | 0.59896 |
| Protein FAM195B | FAM195B | 0.59918 |
| YTH domain family protein 3;YTH domain family protein 1 | YTHDF3;YTHDF1 | 0.60154 |
| Translation initiation factor eIF-2B subunit beta | EIF2B2 | 0.60172 |

| Protein.names | Gene.names | Simvastatin (Ratio.H.L.normalized) |
|---|---|---|
| Serine/threonine-protein kinase 3;Serine/threonine-protein kinase 3 36kDa subunit;Serine/threonine-protein kinase 3 20kDa subunit | STK3 | 0.60217 |
| Hematological and neurological expressed 1 protein | HN1 | 0.60283 |
| Coiled-coil domain-containing protein 22 | CCDC22 | 0.60295 |
| RNA 3-terminal phosphate cyclase-like protein | RCL1 | 0.60411 |
| Brefeldin A-inhibited guanine nucleotide-exchange protein 2 | ARFGEF2 | 0.60459 |
| cAMP-regulated phosphoprotein 21 | ARPP21 | 0.60727 |
| DDB1- and CUL4-associated factor 7 | DCAF7 | 0.60913 |
| Peptidyl-prolyl cis-trans isomerase FKBP8;Peptidyl-prolyl cis-trans isomerase | FKBP8 | 0.61151 |
| Catenin delta-1 | CTNND1 | 0.61198 |
| Cytochrome c oxidase assembly protein 3 homolog, mitochondrial | COA3 | 0.61274 |
| Iron-sulfur protein NUBPL | NUBPL | 0.61625 |
| Immunoglobulin-binding protein 1 | IGBP1 | 0.61832 |
| Spartin | SPG20 | 0.62114 |
| Protein phosphatase methylesterase 1 | PPME1 | 0.62139 |
| Myosin light chain 6B | MYL6B | 0.6224 |
| Cyclin-D1-binding protein 1 | CCNDBP1 | 0.62272 |
| UPF0452 protein C7orf41 | C7orf41 | 0.62292 |
| Importin subunit alpha;Importin subunit alpha-7;Importin subunit alpha-6;Importin subunit alpha-1 | KPNA6;KPNA5;KPNA1 | 0.62357 |
| NHP2-like protein 1 | NHP2L1 | 0.62362 |
| Ataxin-2-like protein | ATXN2L | 0.62371 |
| Small muscular protein | SMPX | 0.6289 |
| Cell division cycle 5-like protein | CDC5L | 0.62893 |
| ARF GTPase-activating protein GIT2 | GIT2 | 0.63079 |
| Biogenesis of lysosome-related organelles complex 1 subunit 1 | BLOC1S1 | 0.63157 |
| Heat shock protein beta-11 | HSPB11 | 0.63207 |
| Myosin-7 | MYH7 | 0.63228 |
| Adenosine deaminase-like protein | ADAL | 0.63334 |
| Allograft inflammatory factor 1-like | AIF1L | 0.63533 |
| Methylthioribose-1-phosphate isomerase | MRI1 | 0.63554 |
| Isochorismatase domain-containing protein 1 | ISOC1 | 0.636 |
| Glutamine--fructose-6-phosphate aminotransferase [isomerizing] 2 | GFPT2 | 0.63616 |
| Junction Plakoglobin | JUP | 0.63631 |
| 28S ribosomal protein S7, mitochondrial | MRPS7 | 0.63704 |
| ATP-binding cassette sub-family F member 2 | ABCF2 | 0.63726 |
| Oligoribonuclease, mitochondrial | REXO2 | 0.63847 |
| Bifunctional 3-phosphoadenosine 5-phosphosulfate synthase 1;Sulfate adenylyltransferase;Adenylyl-sulfate kinase | PAPSS1 | 0.63853 |
| Golgi SNAP receptor complex member 2 | GOSR2 | 0.64063 |
| N-alpha-acetyltransferase 25, NatB auxiliary subunit | NAA25 | 0.64065 |
| Myosin light chain 4 | MYL4 | 0.64081 |
| Transcription initiation factor IIB | GTF2B | 0.64174 |
| Galactocerebrosidase | GALC | 0.64227 |
| Actin-binding LIM protein 3 | ABLIM3 | 0.64597 |
| Splicing factor 3B subunit 1 | SF3B1 | 0.64607 |
| 40S ribosomal protein S10 | RPS10 | 0.64684 |
| Intraflagellar transport protein 27 homolog | IFT27 | 0.64702 |
| Elongation factor 2 | EEF2 | 0.64704 |
| Ubiquitin-conjugating enzyme E2 R2 | UBE2R2 | 0.64707 |
| Serine/threonine-protein kinase N1 | PKN1 | 0.64738 |
| Serine/threonine-protein phosphatase 6 regulatory ankyrin repeat subunit B | ANKRD44 | 0.64772 |
| HEAT repeat-containing protein 5A | HEATR5A | 0.64955 |
| Phosphoglucomutase-like protein 5 | PGM5 | 0.64959 |
| TOM1-like protein 2 | TOM1L2 | 0.64965 |
| Erythrocyte band 7 integral membrane protein | STOM | 0.6502 |
| Golgi to ER traffic protein 4 homolog | GET4 | 0.65101 |
| DNA repair protein XRCC4 | XRCC4 | 0.65142 |
| Peptidyl-prolyl cis-trans isomerase-like 3;Peptidyl-prolyl cis-trans isomerase | PPIL3 | 0.65471 |
| U6 snRNA-associated Sm-like protein LSm1 | LSM1 | 0.65545 |
| Myosin-7B | MYH7B | 0.65591 |
| Glomulin | GLMN | 0.65689 |
| Translation initiation factor eIF-2B subunit gamma | EIF2B3 | 0.65714 |
| 39S ribosomal protein L44, mitochondrial | MRPL44 | 0.65765 |

| Protein.names | Gene.names | Simvastatin (Ratio.H.L.normalized) |
|---|---|---|
| Sorting nexin-17 | SNX17 | 0.65789 |
| Vacuolar protein sorting-associated protein 16 homolog | VPS16 | 0.658 |
| Propionyl-CoA carboxylase beta chain, mitochondrial | PCCB | 0.65824 |
| Exosome complex exonuclease RRP44 | DIS3 | 0.66015 |
| Protein phosphatase 1G | PPM1G | 0.66032 |
| Chromobox protein homolog 1 | CBX1 | 0.66061 |
| Pyridoxal-dependent decarboxylase domain-containing protein 1;Putative pyridoxal-dependent decarboxylase domain-containing protein 2 | PDXDC1;PDXDC2P | 0.66244 |
| Melanoma-associated antigen D2 | MAGED2 | 0.66286 |
| GPI transamidase component PIG-S | PIGS | 0.66294 |
| NHL repeat-containing protein 2 | NHLRC2 | 0.66426 |
| Cell surface glycoprotein MUC18 | MCAM | 0.66539 |
| Serine/threonine-protein phosphatase 6 catalytic subunit | PPP6C | 0.66632 |
| Ubiquitin carboxyl-terminal hydrolase 10 | USP10 | 0.66804 |
| Serine/threonine-protein kinase mTOR | MTOR | 0.66878 |
| Myosin regulatory light polypeptide 9 | MYL9 | 0.66996 |
| Long-chain-fatty-acid--CoA ligase 4 | ACSL4 | 0.67143 |
| Myelin expression factor 2 | MYEF2 | 0.67363 |
| Microtubule-actin cross-linking factor 1, isoforms 1/2/3/5 | MACF1 | 0.67406 |
| Ubiquitin carboxyl-terminal hydrolase;Ubiquitin carboxyl-terminal hydrolase 24 | USP24 | 0.67573 |
| Zinc finger FYVE domain-containing protein 1 | ZFYVE1 | 0.6763 |
| Methylenetetrahydrofolate reductase | MTHFR | 0.67715 |
| Myomesin-2 | MYOM2 | 0.67839 |
| Poly [ADP-ribose] polymerase 1 | PARP1 | 0.67916 |
| Cullin-associated NEDD8-dissociated protein 2 | CAND2 | 0.6793 |
| Replication protein A 70 kDa DNA-binding subunit | RPA1 | 0.67968 |
| Thiosulfate sulfurtransferase | TST | 0.68004 |
| GrpE protein homolog 1, mitochondrial | GRPEL1 | 0.68009 |
| SUN domain-containing protein 1 | SUN1;UNC84A | 0.68026 |
| Armadillo repeat-containing X-linked protein 2 | ARMCX2 | 0.68101 |
| UPF0687 protein C20orf27 | C20orf27 | 0.68124 |
| Protein ATP1B4 | ATP1B4 | 0.68226 |
| ADP-ribosylation factor-like protein 2 | ARL2 | 0.6829 |
| EH domain-binding protein 1-like protein 1 | EHBP1L1 | 0.68338 |
| Sorting nexin-5 | SNX5 | 0.68493 |
| Protein CLEC16A | CLEC16A | 0.68507 |
| Mitochondrial-processing peptidase subunit alpha | PMPCA | 0.68817 |
| Voltage-dependent L-type calcium channel subunit alpha-1S | CACNA1S | 0.68998 |
| Histone deacetylase 6 | HDAC6 | 0.69024 |
| Propionyl-CoA carboxylase alpha chain, mitochondrial | PCCA | 0.69073 |
| Tumor suppressor p53-binding protein 1 | TP53BP1 | 0.69117 |
| Ubiquitin-protein ligase E3A | UBE3A | 0.69155 |
| Tripartite motif-containing protein 55 | TRIM55 | 0.69163 |
| Protein transport protein Sec24A | SEC24A | 0.69306 |
| Alpha-protein kinase 3 | ALPK3 | 0.69383 |
| Myosin IF | MYO1F | 0.69418 |
| Protein DEK | DEK | 0.69451 |
| Tropomodulin-1 | TMOD1 | 0.69513 |
| WW domain-containing transcription regulator protein 1 | WWTR1 | 0.69553 |
| SLIT-ROBO Rho GTPase-activating protein 2C;SLIT-ROBO Rho GTPase-activating protein 2 | SRGAP2C;SRGAP2 | 0.69585 |
| Developmentally-regulated GTP-binding protein 1 | DRG1 | 0.6959 |
| Pre-mRNA-processing factor 19 | PRPF19 | 0.69646 |
| Nuclease EXOG, mitochondrial | EXOG | 0.69658 |
| Pannexin-1 | PANX1 | 0.69856 |
| Actin-binding LIM protein 1 | ABLIM1 | 0.69942 |
| Mitochondrial pyruvate carrier 2 | MPC2 | 0.6995 |
| Reticulon-3 | RTN3 | 0.69975 |
| Protein arginine N-methyltransferase 1 | PRMT1 | 0.69979 |
| 40S ribosomal protein S27 | RPS27 | 0.70005 |
| Coiled-coil domain-containing protein 109B | CCDC109B | 0.70053 |
| Rap guanine nucleotide exchange factor 1 | RAPGEF1;DKFZp781P1719 | 0.70103 |
| pre-rRNA processing protein FTSJ3 | FTSJ3 | 0.70142 |
| Ryanodine receptor 1 | RYR1 | 0.70187 |

| Protein.names | Gene.names | Simvastatin (Ratio.H.L.normalized) |
|---|---|---|
| Poly(U)-binding-splicing factor PUF60 | PUF60 | 0.70234 |
| Tripartite motif-containing protein 54 | TRIM54 | 0.70265 |
| RNA-binding protein 25 | RBM25 | 0.70282 |
| IST1 homolog | IST1 | 0.70289 |
| Splicing factor U2AF 65 kDa subunit | U2AF2 | 0.70337 |
| Ubiquitin-conjugating enzyme E2 D2;Ubiquitin-conjugating enzyme E2 D3 | UBE2D3;UBE2D2 | 0.70343 |
| Kynurenine--oxoglutarate transaminase 3 | CCBL2 | 0.70347 |
| Proteasome activator complex subunit 3 | PSME3 | 0.70355 |
| Ran-binding protein 9 | RANBP9 | 0.70365 |
| Stromal interaction molecule 1 | STIM1 | 0.70379 |
| Ragulator complex protein LAMTOR5 | LAMTOR5 | 0.70485 |
| 28S ribosomal protein S29, mitochondrial | DAP3 | 0.70535 |
| Mitochondrial inner membrane protein OXA1L | OXA1L | 0.70571 |
| Regulator of nonsense transcripts 2 | UPF2 | 0.70583 |
| Elongator complex protein 1 | IKBKAP | 0.70637 |
| Eukaryotic translation initiation factor 3 subunit G | EIF3G | 0.70656 |
| Superkiller viralicidic activity 2-like 2 | SKIV2L2 | 0.70658 |
| Cytochrome c oxidase subunit 6C | COX6C | 0.70659 |
| Titin | TTN | 0.70667 |
| Histone-lysine N-methyltransferase SMYD1 | SMYD1 | 0.70679 |
| FH1/FH2 domain-containing protein 1 | FHOD1 | 0.70685 |
| Calponin-2 | CNN2 | 0.70742 |
| 60S ribosomal protein L24 | RPL24 | 0.70899 |
| Elongation factor Ts, mitochondrial;Elongation factor Ts | TSFM | 0.70912 |
| FACT complex subunit SPT16 | SUPT16H | 0.71023 |
| Ribosomal protein L19;60S ribosomal protein L19 | RPL19 | 0.71188 |
| TFIIH basal transcription factor complex helicase XPB subunit | ERCC3 | 0.71244 |
| CLIP-associating protein 1 | CLASP1 | 0.71254 |
| Cytochrome c oxidase subunit 7A1, mitochondrial | COX7A1 | 0.71286 |
| 60S ribosomal protein L9 | RPL9 | 0.7132 |
| Rho guanine nucleotide exchange factor 6 | ARHGEF6 | 0.71344 |
| EGF-like repeat and discoidin I-like domain-containing protein 3 | EDIL3 | 0.71365 |
| 60S ribosomal protein L36a-like;60S ribosomal protein L36a | RPL36AL;RPL36A | 0.7143 |
| LIM and calponin homology domains-containing protein 1 | LIMCH1 | 0.71447 |
| UPF0364 protein C6orf211 | C6orf211 | 0.71519 |
| Calsequestrin-2;Calsequestrin | CASQ2 | 0.71567 |
| Neurochondrin | NCDN | 0.71634 |
| Ankyrin repeat domain-containing protein 1 | ANKRD1 | 0.71714 |
| X-ray repair cross-complementing protein 5 | XRCC5 | 0.71714 |
| E3 ubiquitin-protein ligase UBR3 | UBR3 | 0.71797 |
| Cofilin-2 | CFL2 | 0.71874 |
| Myomesin-1 | MYOM1 | 0.71882 |
| Trafficking protein particle complex subunit 4 | TRAPPC4 | 0.71936 |
| Rho GTPase-activating protein 12 | ARHGAP12 | 0.72044 |
| 28S ribosomal protein S17, mitochondrial | MRPS17 | 0.7206 |
| 40S ribosomal protein S24 | RPS24 | 0.72089 |
| Putative 60S ribosomal protein L39-like 5;60S ribosomal protein L39 | RPL39P5;RPL39 | 0.72177 |
| Cleavage stimulation factor subunit 2 | CSTF2 | 0.72263 |
| Apoptotic protease-activating factor 1 | APAF1 | 0.7228 |
| Ankyrin repeat domain-containing protein 13A;Ankyrin repeat domain-containing protein 13D | ANKRD13A;ANKRD13D | 0.72314 |
| E3 ubiquitin-protein ligase;E3 ubiquitin-protein ligase NEDD4-like | NEDD4L | 0.72443 |
| Zinc finger protein 512 | ZNF512 | 0.72516 |
| ER membrane protein complex subunit 3 | EMC3 | 0.72571 |
| 5-phosphohydroxy-L-lysine phospho-lyase | AGXT2L2 | 0.72592 |
| THUMP domain-containing protein 3 | THUMPD3 | 0.7264 |
| Type II inositol 3,4-bisphosphate 4-phosphatase | INPP4B | 0.72693 |
| Protein kish-A | TMEM167A | 0.72759 |
| Syntaxin-8 | STX8 | 0.72767 |
| Drebrin | DBN1 | 0.72776 |
| Oxysterol-binding protein;Oxysterol-binding protein-related protein 11;Oxysterol-binding protein-related protein 10 | OSBPL10;OSBPL11 | 0.72816 |
| Homer protein homolog 1 | HOMER1 | 0.72835 |
| Mitochondrial import receptor subunit TOM22 homolog | TOMM22 | 0.72841 |

| Protein.names | Gene.names | Simvastatin (Ratio.H.L.normalized) |
|---|---|---|
| 60S ribosomal protein L31 | RPL31 | 0.72857 |
| Interferon-induced, double-stranded RNA-activated protein kinase | EIF2AK2 | 0.72894 |
| Semaphorin-3C | SEMA3C | 0.72922 |
| Musculoskeletal embryonic nuclear protein 1 | MUSTN1;TMEM110-MUSTN1 | 0.72935 |
| Integrin alpha-6;Integrin alpha-6 heavy chain;Integrin alpha-6 light chain | ITGA6 | 0.72945 |
| Transcription factor BTF3 | BTF3 | 0.7305 |
| Beta-enolase;Enolase | ENO3 | 0.73067 |
| MTSS1-like protein | MTSS1L | 0.73132 |
| 40S ribosomal protein S4, X isoform;40S ribosomal protein S4, Y isoform 2 | RPS4X;RPS4Y2 | 0.73164 |
| Cysteine-rich protein 1 | CRIP1 | 0.73287 |
| Coatomer subunit epsilon | COPE | 0.73296 |
| Tyrosine--tRNA ligase, mitochondrial | YARS2 | 0.73299 |
| Double-stranded RNA-binding protein Staufen homolog 1 | STAU1 | 0.73318 |
| Probable leucine--tRNA ligase, mitochondrial | LARS2 | 0.73325 |
| Interleukin enhancer-binding factor 2 | ILF2 | 0.73349 |
| U2 small nuclear ribonucleoprotein A | SNRPA1 | 0.73405 |
| Protein diaphanous homolog 1 | DIAPH1 | 0.73413 |
| Cob(I)yrinic acid a,c-diamide adenosyltransferase, mitochondrial | MMAB | 0.73451 |
| Leiomodin-2 | LMOD2 | 0.73488 |
| 40S ribosomal protein S16 | RPS16 | 0.73499 |
| 40S ribosomal protein S13 | RPS13 | 0.73527 |
| Protein CREG1 | CREG1 | 0.73547 |
| 60S ribosomal protein L38 | RPL38 | 0.7355 |
| 40S ribosomal protein S26 | RPS26 | 0.73631 |
| Nesprin-1 | SYNE1 | 0.7366 |
| Serine/arginine-rich splicing factor 6;Serine/arginine-rich splicing factor 4 | SRSF6;SRSF4 | 0.73687 |
| LIM domain and actin-binding protein 1 | LIMA1 | 0.73712 |
| RILP-like protein 1 | RILPL1 | 0.73736 |
| Beta-sarcoglycan | SGCB | 0.73766 |
| Protein LZIC | LZIC | 0.73806 |
| Sister chromatid cohesion protein PDS5 homolog B | PDS5B | 0.73842 |
| WD repeat-containing protein 47 | WDR47 | 0.73864 |
| NADH dehydrogenase [ubiquinone] 1 alpha subcomplex assembly factor 4 | NDUFAF4 | 0.73902 |
| AP-1 complex subunit beta-1 | AP1B1 | 0.7394 |
| 60S ribosomal protein L11 | RPL11 | 0.73976 |
| 60S ribosomal protein L34 | RPL34 | 0.74021 |
| cGMP-dependent protein kinase 1 | PRKG1 | 0.74036 |
| Nuclear fragile X mental retardation-interacting protein 2 | NUFIP2 | 0.74046 |
| 39S ribosomal protein L41, mitochondrial | MRPL41 | 0.74049 |
| Junctional adhesion molecule C | JAM3 | 0.74249 |
| Mitochondrial 2-oxoglutarate/malate carrier protein | SLC25A11 | 0.74257 |
| TBC1 domain family member 24 | TBC1D24 | 0.7429 |
| Armadillo repeat-containing protein 8 | ARMC8 | 0.74313 |
| Serine/threonine-protein phosphatase 4 regulatory subunit 3A | SMEK1 | 0.74313 |
| Cyclin-dependent kinase 18 | CDK18 | 0.74314 |
| LIM and cysteine-rich domains protein 1 | LMCD1 | 0.74328 |
| Troponin I, slow skeletal muscle | TNNI1 | 0.7433 |
| Zinc finger and BTB domain-containing protein 20 | ZBTB20 | 0.74369 |
| Far upstream element-binding protein 1 | FUBP1 | 0.74408 |
| Acyl-protein thioesterase 1 | LYPLA1 | 0.74462 |
| ER lumen protein retaining receptor 1;ER lumen protein retaining receptor;ER lumen protein retaining receptor 2 | KDELR1;KDELR2 | 0.74464 |
| Chitinase domain-containing protein 1 | CHID1 | 0.74553 |
| Ras-related protein Rab-34 | RAB34 | 0.74588 |
| BolA-like protein 2 | BOLA2 | 0.74594 |
| Troponin C, slow skeletal and cardiac muscles | TNNC1 | 0.74677 |
| Dolichyl-diphosphooligosaccharide--protein glycosyltransferase 48 kDa subunit | DDOST | 0.74699 |
| Isocitrate dehydrogenase [NADP], mitochondrial;Isocitrate dehydrogenase [NADP] | IDH2 | 0.7474 |
| Transforming growth factor-beta receptor-associated protein 1 | TGFBRAP1 | 0.74769 |
| Phosphatidylinositol 3,4,5-trisphosphate 5-phosphatase 2 | INPPL1 | 0.74823 |
| Histone-arginine methyltransferase CARM1 | CARM1 | 0.7483 |
| RNA-binding motif protein, X chromosome;RNA-binding motif protein, X chromosome, N-terminally processed;RNA binding motif protein, X-linked-like-1 | RBMX;RBMXL1 | 0.74877 |

| Protein.names | Gene.names | Simvastatin (Ratio.H.L.normalized) |
|---|---|---|
| Serine/threonine-protein phosphatase 2A activator | PPP2R4 | 0.74899 |
| Protein unc-45 homolog B | UNC45B | 0.74944 |
| 3-ketoacyl-CoA thiolase, peroxisomal | ACAA1 | 1.2504 |
| Inhibitor of nuclear factor kappa-B kinase-interacting protein | IKBIP | 1.2513 |
| Nicotinamide phosphoribosyltransferase | NAMPT;NAMPTL | 1.2523 |
| Alpha-crystallin B chain | CRYAB | 1.255 |
| Cathepsin L1;Cathepsin L1 heavy chain;Cathepsin L1 light chain;Putative inactive cathepsin L-like protein CTSL3P;Cathepsin K;Cathepsin L2 | CTSL1;CTSL3P;CTSK;CTSL2 | 1.2551 |
| Large neutral amino acids transporter small subunit 1 | SLC7A5 | 1.2553 |
| UTP--glucose-1-phosphate uridylyltransferase | UGP2 | 1.2557 |
| Argininosuccinate synthase | ASS1 | 1.2564 |
| Heme oxygenase 2 | HMOX2 | 1.2567 |
| Elongation factor Tu GTP-binding domain-containing protein 1 | EFTUD1 | 1.2576 |
| Histone H1.2;Histone H1.4;Histone H1.3 | HIST1H1C;HIST1H1E;HIST1H1D | 1.2593 |
| Glutathione reductase, mitochondrial | GSR | 1.2594 |
| Ras-related protein Rap-1A | RAP1A | 1.2615 |
| Vacuolar protein-sorting-associated protein 25 | VPS25 | 1.2617 |
| Ethanolamine-phosphate cytidylyltransferase | PCYT2 | 1.2625 |
| Protein FAM35A | FAM35A | 1.2626 |
| Conserved oligomeric Golgi complex subunit 3 | COG3 | 1.2628 |
| Prelamin-A/C;Lamin-A/C | LMNA | 1.2629 |
| Reticulon 4 | RTN4 | 1.2631 |
| Inorganic pyrophosphatase 2, mitochondrial | PPA2 | 1.2636 |
| Cytoplasmic aconitate hydratase | IRP1;ACO1 | 1.2639 |
| N-acetylglucosamine-6-sulfatase | GNS | 1.2646 |
| Proteasome activator complex subunit 2 | PSME2 | 1.2653 |
| Coiled-coil domain-containing protein 6 | CCDC6 | 1.2659 |
| Fatty acid-binding protein, epidermal | FABP5 | 1.2659 |
| Protein disulfide-isomerase A5 | PDIA5 | 1.2679 |
| Collagen alpha-1(V) chain | COL5A1 | 1.269 |
| Oxysterol-binding protein-related protein 8 | OSBPL8 | 1.2691 |
| Anthrax toxin receptor 1 | ANTXR1 | 1.272 |
| UDP-glucose 6-dehydrogenase | UGDH | 1.2728 |
| Neutral cholesterol ester hydrolase 1 | NCEH1 | 1.2729 |
| Ras-related protein Rab-2A;Ras-related protein Rab-2B | RAB2A;RAB2B | 1.2733 |
| Calumenin | CALU | 1.2743 |
| Dynamin-1-like protein | DNM1L | 1.2745 |
| Eukaryotic translation initiation factor 2 subunit 2 | EIF2S2 | 1.2747 |
| BTB/POZ domain-containing protein KCTD12 | KCTD12 | 1.2749 |
| Clathrin light chain B | CLTB | 1.2754 |
| Serine beta-lactamase-like protein LACTB, mitochondrial | LACTB | 1.2759 |
| Focadhesin | FOCAD | 1.2764 |
| Protein SEC13 homolog | SEC13 | 1.2773 |
| Switch-associated protein 70 | SWAP70 | 1.2785 |
| cAMP-dependent protein kinase type I-alpha regulatory subunit;cAMP-dependent protein kinase type I-alpha regulatory subunit, N-terminally | FHADHR1AR1A | 1.2808 |
| Annexin A2;Annexin;Putative annexin A2-like protein | ANXA2;ANXA2P2 | 1.2816 |
| 45 kDa calcium-binding protein | SDF4 | 1.2823 |
| Beta-2-microglobulin;Beta-2-microglobulin form pI 5.3 | B2M | 1.2832 |
| Adenylyl cyclase-associated protein 1 | CAP1 | 1.2836 |
| Voltage-dependent calcium channel gamma-1 subunit | CACNG1 | 1.2839 |
| Glucosidase 2 subunit beta | PRKCSH | 1.286 |
| 39S ribosomal protein L11, mitochondrial | MRPL11 | 1.2862 |
| Sodium/potassium-transporting ATPase subunit alpha-2 | ATP1A2 | 1.2866 |
| B-cell receptor-associated protein 31 | BCAP31 | 1.2877 |
| Isocitrate dehydrogenase [NAD] subunit beta, mitochondrial | IDH3B | 1.2879 |
| Isochorismatase domain-containing protein 2, mitochondrial | ISOC2 | 1.2894 |
| Cytochrome b-c1 complex subunit 9 | UQCR10 | 1.2894 |
| Urotensin-2 | UTS2 | 1.2902 |
| 4-aminobutyrate aminotransferase, mitochondrial | ABAT | 1.2904 |
| Glutamate dehydrogenase 1, mitochondrial;Glutamate dehydrogenase;Glutamate dehydrogenase 2, mitochondrial | GLUD1;GLUD2 | 1.2908 |
| Glutaryl-CoA dehydrogenase, mitochondrial | GCDH | 1.2909 |
| Guanine nucleotide-binding protein G(I)/G(S)/G(T) subunit beta-2 | GNB2 | 1.2909 |

| Protein.names | Gene.names | Simvastatin (Ratio.H.L.normalized) |
|---|---|---|
| Redox-regulatory protein FAM213A | FAM213A | 1.2923 |
| Myotrophin | MTPN | 1.2923 |
| HLA class I histocompatibility antigen, alpha chain G | HLA-G | 1.2928 |
| Aldose reductase | AKR1B1 | 1.293 |
| Very-long-chain (3R)-3-hydroxyacyl-[acyl-carrier protein] dehydratase 3 | PTPLAD1 | 1.2933 |
| Osteoclast-stimulating factor 1 | OSTF1 | 1.2956 |
| Active breakpoint cluster region-related protein | ABR | 1.2966 |
| Ras-related GTP-binding protein C;Ras-related GTP-binding protein D | RRAGC;RRAGD | 1.297 |
| Transmembrane glycoprotein NMB | GPNMB | 1.2982 |
| Exocyst complex component 1 | EXOC1 | 1.299 |
| Guanine nucleotide-binding protein G(q) subunit alpha | GNAQ | 1.2995 |
| Protein S100-A16 | S100A16 | 1.2995 |
| UPF0600 protein C5orf51 | C5orf51 | 1.2997 |
| DNA dC->dU-editing enzyme APOBEC-3C | APOBEC3C | 1.3 |
| Beta-hexosaminidase subunit beta;Beta-hexosaminidase subunit beta chain B;Beta-hexosaminidase subunit beta chain A | HEXB | 1.3002 |
| Transketolase | TKT | 1.3002 |
| Laminin subunit gamma-1 | LAMC1 | 1.3006 |
| Saccharopine dehydrogenase-like oxidoreductase | SCCPDH | 1.3006 |
| Ras-related protein R-Ras2 | RRAS2 | 1.3015 |
| Lysosomal Pro-X carboxypeptidase | PRCP | 1.3017 |
| Tyrosine-protein phosphatase non-receptor type 11 | PTPN11 | 1.3032 |
| Alpha-enolase;Enolase | ENO1 | 1.304 |
| Insulin-like growth factor 2 mRNA-binding protein 2 | IGF2BP2 | 1.3045 |
| Pseudouridine-5-monophosphatase | HDHD1 | 1.3065 |
| Dynactin subunit 4 | DCTN4 | 1.3072 |
| Protein S100-A4 | S100A4 | 1.3075 |
| Filamin-A | FLNA | 1.3079 |
| KDEL motif-containing protein 2 | KDELC2 | 1.308 |
| LIM and SH3 domain protein 1 | LASP1 | 1.3109 |
| Endoplasmic reticulum aminopeptidase 1 | ERAP1 | 1.3118 |
| Sterol-4-alpha-carboxylate 3-dehydrogenase, decarboxylating | NSDHL | 1.3122 |
| Zinc transporter 1 | SLC30A1 | 1.3127 |
| Thioredoxin | TXN | 1.3128 |
| Sodium-coupled neutral amino acid transporter 2 | SLC38A2 | 1.313 |
| Creatine kinase B-type | CKB | 1.3132 |
| Zinc finger CCCH-type antiviral protein 1 | ZC3HAV1 | 1.3132 |
| NADH dehydrogenase [ubiquinone] 1 alpha subcomplex subunit 13 | NDUFA13;YJEFN3 | 1.3135 |
| Ferritin heavy chain;Ferritin | FTH1 | 1.3137 |
| cAMP-regulated phosphoprotein 19;Alpha-endosulfine | ARPP19;ENSA | 1.3142 |
| Gem-associated protein 5 | GEMIN5 | 1.315 |
| Kinesin light chain 2 | KLC2 | 1.315 |
| Tyrosine-protein phosphatase non-receptor type 1;Tyrosine-protein phosphatase non-receptor type | PTPN1 | 1.3162 |
| mRNA cap guanine-N7 methyltransferase | RNMT | 1.3163 |
| L-lactate dehydrogenase A chain | LDHA | 1.3164 |
| Mevalonate kinase | MVK | 1.3167 |
| RNA-binding motif, single-stranded-interacting protein 1;RNA-binding motif, single-stranded-interacting protein 3 | RBMS1;RBMS3 | 1.3167 |
| TLD domain-containing protein KIAA1609 | TLDC1;KIAA1609 | 1.3169 |
| Ubiquitin-like-conjugating enzyme ATG3 | ATG3 | 1.3188 |
| Protein kinase C and casein kinase substrate in neurons protein 2 | PACSIN2 | 1.3192 |
| NIF3-like protein 1 | NIF3L1 | 1.3193 |
| Vimentin | VIM | 1.3196 |
| Sideroflexin-1 | SFXN1 | 1.3219 |
| Cleavage stimulation factor subunit 1 | CSTF1 | 1.3226 |
| Choline transporter-like protein 2 | SLC44A2 | 1.3239 |
| LIM and senescent cell antigen-like-containing domain protein 1;LIM and senescent cell antigen-like-containing domain protein 2 | LIMS1;LIMS2 | 1.324 |
| Serine protease HTRA3 | HTRA3 | 1.3248 |
| Procollagen-lysine,2-oxoglutarate 5-dioxygenase 3 | PLOD3 | 1.3251 |
| GDP-fucose protein O-fucosyltransferase 1 | POFUT1 | 1.3259 |
| Pterin-4-alpha-carbinolamine dehydratase | PCBD1 | 1.3275 |
| Lanosterol synthase | LSS | 1.3277 |

| Protein.names | Gene.names | Simvastatin (Ratio.H.L.normalized) |
|---|---|---|
| Paralemmin-2 | PALM2;PALM2-AKAP2 | 1.3284 |
| Acetyl-coenzyme A transporter 1 | SLC33A1 | 1.3284 |
| Coproporphyrinogen-III oxidase, mitochondrial | CPOX | 1.3293 |
| Sorting nexin-2 | SNX2 | 1.3303 |
| Actin, alpha skeletal muscle | ACTA1 | 1.3304 |
| Myristoylated alanine-rich C-kinase substrate | MARCKS | 1.3316 |
| Fibrillin-1 | FBN1 | 1.3317 |
| FERM, RhoGEF and pleckstrin domain-containing protein 1 | FARP1 | 1.3322 |
| NAD(P)H-hydrate epimerase | APOA1BP | 1.3329 |
| Cytoplasmic dynein 1 light intermediate chain 1 | DYNC1LI1 | 1.3333 |
| Ras-related protein Rab-1B;Putative Ras-related protein Rab-1C | RAB1B;RAB1C | 1.3347 |
| Dihydropyrimidinase-related protein 2 | DPYSL2 | 1.335 |
| Signal peptidase complex subunit 2 | SPCS2 | 1.3351 |
| Adapter molecule crk | CRK | 1.3356 |
| 6-phosphofructokinase type C | PFKP | 1.3359 |
| Acyl-CoA-binding domain-containing protein 5 | ACBD5 | 1.3367 |
| Calpain small subunit 1 | CAPNS1 | 1.3378 |
| FK506-binding protein 15 | FKBP15 | 1.3385 |
| Ubiquitin-like modifier-activating enzyme ATG7 | ATG7 | 1.3388 |
| Atlastin-3 | ATL3 | 1.3389 |
| CD59 glycoprotein | CD59 | 1.3411 |
| Kinase D-interacting substrate of 220 kDa | KIDINS220 | 1.342 |
| Tumor protein D54 | TPD52L2 | 1.3435 |
| Aldo-keto reductase family 1 member C3 | AKR1C3 | 1.3439 |
| Sorcin | SRI | 1.345 |
| CD82 antigen | CD82 | 1.3476 |
| Protein disulfide-isomerase A4 | PDIA4 | 1.3479 |
| Collagen type IV alpha-3-binding protein | COL4A3BP | 1.3481 |
| Protein-glutamine gamma-glutamyltransferase 2 | TGM2 | 1.3493 |
| Regulator of microtubule dynamics protein 1 | RMDN1 | 1.3502 |
| Phostensin | PPP1R18 | 1.3507 |
| Long-chain-fatty-acid--CoA ligase 3 | ACSL3 | 1.3508 |
| FUN14 domain-containing protein 2 | FUNDC2 | 1.3528 |
| UPF0556 protein C19orf10 | C19orf10 | 1.3532 |
| Integrin alpha-V;Integrin alpha-V heavy chain;Integrin alpha-V light chain | ITGAV | 1.3539 |
| Xaa-Pro dipeptidase | PEPD | 1.3545 |
| Ankyrin repeat and SAM domain-containing protein 1A | ANKS1A | 1.3547 |
| Nucleobindin-2;Nesfatin-1 | NUCB2 | 1.3557 |
| 28S ribosomal protein S31, mitochondrial | MRPS31 | 1.3561 |
| Glutathione S-transferase omega-1 | GSTO1 | 1.3574 |
| Microtubule-associated protein 1B;MAP1B heavy chain;MAP1 light chain LC1 | MAP1B | 1.359 |
| 39S ribosomal protein L1, mitochondrial | MRPL1 | 1.3597 |
| GTPase KRas;GTPase KRas, N-terminally processed | KRAS | 1.3617 |
| Protein S100-A11 | S100A11 | 1.3621 |
| Discoidin domain-containing receptor 2 | DDR2 | 1.3623 |
| Leucine-rich repeat serine/threonine-protein kinase 1 | LRRK1 | 1.3624 |
| Synaptic vesicle membrane protein VAT-1 homolog | VAT1 | 1.3624 |
| Vacuolar protein sorting-associated protein 33A | VPS33A | 1.3631 |
| Receptor expression-enhancing protein 6 | REEP6 | 1.3648 |
| Tyrosine-protein phosphatase non-receptor type 12 | PTPN12 | 1.3654 |
| Plasma membrane calcium-transporting ATPase 1 | ATP2B1 | 1.3658 |
| Fructose-bisphosphate aldolase C;Fructose-bisphosphate aldolase | ALDOC | 1.366 |
| Macrophage-capping protein | CAPG | 1.3677 |
| Hydroxysteroid dehydrogenase-like protein 2 | HSDL2 | 1.3685 |
| Glia-derived nexin | SERPINE2 | 1.3691 |
| Calcium/calmodulin-dependent protein kinase type II subunit gamma;Calcium/calmodulin-dependent protein kinase type II subunit beta | CAMK2G;CAMK2B | 1.3694 |
| TBC1 domain family member 15 | TBC1D15 | 1.3697 |
| Glucose-6-phosphate 1-dehydrogenase | G6PD | 1.3715 |
| Beta-taxilin | TXLNB | 1.3722 |
| Geranylgeranyl transferase type-1 subunit beta | PGGT1B | 1.3747 |
| PRA1 family protein 3 | ARL6IP5 | 1.377 |
| Acyl-CoA synthetase family member 2, mitochondrial | ACSF2 | 1.3786 |
| Myosin light polypeptide 6 | MYL6 | 1.379 |

| Protein.names | Gene.names | Simvastatin (Ratio.H.L.normalized) |
|---|---|---|
| Twinfilin-1 | TWF1 | 1.3796 |
| Signal transducer and activator of transcription 6 | STAT6 | 1.3797 |
| Gamma-enolase;Enolase | ENO2 | 1.38 |
| DNA mismatch repair protein Msh2 | MSH2 | 1.3826 |
| Tensin-1 | TNS1 | 1.3829 |
| Radixin | RDX | 1.3836 |
| Coronin-1C;Coronin | CORO1C | 1.3842 |
| 7-dehydrocholesterol reductase | DHCR7 | 1.385 |
| Protein NOXP20 | FAM114A1;DKFZp686F20250 | 1.3853 |
| Fermitin family homolog 3 | FERMT3 | 1.3853 |
| Ankycorbin | RAI14 | 1.3858 |
| Neutral amino acid transporter B(0) | SLC1A5 | 1.387 |
| Multivesicular body subunit 12A | MVB12A | 1.3891 |
| Myotonin-protein kinase | DMPK | 1.3913 |
| E3 ubiquitin-protein ligase KCMF1 | KCMF1 | 1.3914 |
| Peripherin | PRPH | 1.3929 |
| UDP-N-acetylglucosamine--dolichyl-phosphate N-acetylglucosaminephosphotransferase | DPAGT1 | 1.3935 |
| Sorting and assembly machinery component 50 homolog | SAMM50 | 1.3944 |
| Acyl-CoA-binding protein | DBI | 1.3945 |
| Plexin-B1;Plexin-B3 | PLXNB1;PLXNB3 | 1.3947 |
| Pinin | PNN | 1.398 |
| Leucine zipper transcription factor-like protein 1 | LZTFL1 | 1.3989 |
| Actin-related protein 2/3 complex subunit 1B | ARPC1B | 1.4004 |
| Calpain-2 catalytic subunit | CAPN2 | 1.4009 |
| Actin-related protein 2/3 complex subunit 5 | ARPC5 | 1.4014 |
| Nucleolar GTP-binding protein 1 | GTPBP4 | 1.4017 |
| Gelsolin | GSN | 1.4036 |
| Deoxycytidylate deaminase | DCTD | 1.4044 |
| Ras-related protein R-Ras | RRAS | 1.4049 |
| Zinc finger CCCH domain-containing protein 6 | ZC3H6 | 1.4069 |
| Signal peptidase complex subunit 1 | SPCS1 | 1.4072 |
| Tropomodulin-3 | TMOD3 | 1.4072 |
| Golgi-associated plant pathogenesis-related protein 1 | GLIPR2 | 1.408 |
| Vesicle-associated membrane protein 3;Vesicle-associated membrane protein 2 | VAMP3;VAMP2 | 1.4088 |
| Pro-cathepsin H;Cathepsin H mini chain;Cathepsin H;Cathepsin H heavy chain;Cathepsin H light chain | CTSH | 1.4095 |
| Acetyl-coenzyme A synthetase, cytoplasmic | ACSS2 | 1.4103 |
| A-kinase anchor protein 12 | AKAP12 | 1.4118 |
| Ras GTPase-activating-like protein IQGAP1 | IQGAP1 | 1.4123 |
| Chloride intracellular channel protein 1 | CLIC1 | 1.4128 |
| 3-oxo-5-beta-steroid 4-dehydrogenase | AKR1D1 | 1.4139 |
| Presequence protease, mitochondrial | PITRM1 | 1.4154 |
| Myosin light chain 5 | MYL5 | 1.4156 |
| Acidic leucine-rich nuclear phosphoprotein 32 family member E | ANP32E | 1.4159 |
| Na(+)/H(+) exchange regulatory cofactor NHE-RF1 | SLC9A3R1 | 1.4172 |
| Protein kinase C alpha type | PRKCA | 1.4198 |
| Serum deprivation-response protein | SDPR | 1.4203 |
| Protein S100-A10 | S100A10 | 1.4207 |
| Cytoplasmic dynein 1 intermediate chain 2 | DYNC1I2 | 1.4214 |
| Protein S100-A9 | S100A9 | 1.422 |
| StAR-related lipid transfer protein 9 | STARD9 | 1.4251 |
| Epoxide hydrolase 1 | EPHX1 | 1.4276 |
| Transcription elongation factor SPT5 | SUPT5H | 1.4276 |
| Adenosine deaminase | ADA | 1.4284 |
| Serine/arginine-rich splicing factor 10;Serine/arginine-rich splicing factor 12 | SRSF10;SRSF12 | 1.4295 |
| Optineurin | OPTN | 1.4297 |
| Bifunctional ATP-dependent dihydroxyacetone kinase/FAD-AMP lyase (cyclizing);ATP-dependent dihydroxyacetone kinase;FAD-AMP lyase (cyclizing) | DAK | 1.4314 |
| Glutathione S-transferase P | GSTP1 | 1.4324 |
| Plasma membrane calcium-transporting ATPase 3 | ATP2B3 | 1.4339 |
| Trafficking protein particle complex subunit 3 | TRAPPC3 | 1.437 |
| Galactokinase | GALK1 | 1.4376 |
| Tubulin polymerization-promoting protein family member 3 | TPPP3 | 1.4378 |

| Protein.names | Gene.names | Simvastatin (Ratio.H.L.normalized) |
|---|---|---|
| SH3 domain-binding glutamic acid-rich-like protein 3 | SH3BGRL3 | 1.44 |
| Mitogen-activated protein kinase 14 | MAPK14 | 1.4434 |
| Methionine aminopeptidase;Methionine aminopeptidase 2 | METAP2 | 1.4451 |
| Calmodulin | CALM1;CALM2;CALM3 | 1.4464 |
| Aspartyl/asparaginyl beta-hydroxylase | ASPH | 1.4473 |
| Tropomyosin 2 (Beta) | TPM2 | 1.448 |
| A-kinase anchor protein 2 | AKAP2 | 1.4498 |
| Polypeptide N-acetylgalactosaminyltransferase 1;Polypeptide N-acetylgalactosaminyltransferase 1 soluble form | GALNT1 | 1.4502 |
| 4F2 cell-surface antigen heavy chain | SLC3A2 | 1.4525 |
| Striatin | STRN | 1.4536 |
| Protein S100-A13 | S100A13 | 1.4558 |
| ATP synthase-coupling factor 6, mitochondrial | ATP5J | 1.4572 |
| Histone-binding protein RBBP7 | RBBP7 | 1.4576 |
| Acetylcholine receptor subunit alpha | CHRNA1 | 1.461 |
| Rapamycin-insensitive companion of mTOR | RICTOR | 1.4611 |
| Serine/threonine-protein kinase 38 | STK38 | 1.4626 |
| TBC1 domain family member 17 | TBC1D17 | 1.4646 |
| Exportin-5 | XPO5 | 1.4673 |
| T-complex protein 11-like protein 1 | TCP11L1 | 1.4695 |
| Extended synaptotagmin-2 | ESYT2 | 1.4707 |
| Methyltransferase-like protein 7A | METTL7A | 1.4728 |
| Tricarboxylate transport protein, mitochondrial | SLC25A1 | 1.4751 |
| Sorbitol dehydrogenase | SORD | 1.4778 |
| STAM-binding protein | STAMBP | 1.4783 |
| Tenascin | TNC | 1.4832 |
| Carnitine O-acetyltransferase | CRAT | 1.4847 |
| Delta(24)-sterol reductase | DHCR24 | 1.4864 |
| Tropomyosin alpha-4 chain | TPM4 | 1.4884 |
| Tubulin beta-3 chain | TUBB3 | 1.4886 |
| Putative ataxin-7-like protein 3B | ATXN7L3B | 1.4897 |
| Acetyl-CoA acetyltransferase, cytosolic | ACAT2 | 1.4903 |
| Cathepsin B;Cathepsin B light chain;Cathepsin B heavy chain | CTSB | 1.4911 |
| Polypyrimidine tract-binding protein 1 | PTBP1 | 1.4931 |
| Peroxiredoxin-5, mitochondrial | PRDX5 | 1.4934 |
| Protein cornichon homolog 4 | CNIH4 | 1.4949 |
| UDP-N-acetylhexosamine pyrophosphorylase;UDP-N-acetylgalactosamine pyrophosphorylase;UDP-N-acetylglucosamine pyrophosphorylase | UAP1 | 1.4981 |
| Acetyl-CoA carboxylase 1;Biotin carboxylase | ACACA | 1.4982 |
| Heat shock 70 kDa protein 1A/1B | HSPA1A | 1.5009 |
| Calcium/calmodulin-dependent protein kinase type II subunit delta | CAMK2D | 1.5011 |
| Autophagy-related protein 9A | ATG9A | 1.5049 |
| Dual specificity mitogen-activated protein kinase kinase 1 | MAP2K1 | 1.5062 |
| Raftlin | RFTN1 | 1.5075 |
| Integrin alpha-3;Integrin alpha-3 heavy chain;Integrin alpha-3 light chain | ITGA3 | 1.5082 |
| Phosphatidylethanolamine-binding protein 1;Hippocampal cholinergic neurostimulating peptide | PEBP1 | 1.5146 |
| Ras-related protein Rab-5A | RAB5A | 1.515 |
| Collagen alpha-2(I) chain | COL1A2 | 1.5163 |
| Myotubularin-related protein 6 | MTMR6 | 1.5186 |
| Phosphatidylinositide phosphatase SAC1 | SACM1L | 1.5195 |
| Thy-1 membrane glycoprotein | THY1 | 1.5195 |
| NAD(P)H dehydrogenase [quinone] 1 | NQO1 | 1.5199 |
| Receptor-type tyrosine-protein phosphatase O | PTPRO | 1.5215 |
| Leupaxin | LPXN | 1.5236 |
| Transgelin-2 | TAGLN2 | 1.5243 |
| Unconventional myosin-VI | MYO6 | 1.5301 |
| Galectin-3-binding protein | LGALS3BP | 1.5392 |
| AP-4 complex subunit epsilon-1 | AP4E1 | 1.5425 |
| Serine/threonine-protein kinase Nek9 | NEK9 | 1.5458 |
| Solute carrier family 35 member E2;Solute carrier family 35 member E2B | SLC35E2;SLC35E2B | 1.5474 |
| Retinoid-inducible serine carboxypeptidase | SCPEP1 | 1.5477 |
| Monoglyceride lipase | MGLL | 1.5566 |
| Clathrin light chain A | CLTA | 1.5569 |

| Protein.names | Gene.names | Simvastatin (Ratio.H.L.normalized) |
|---|---|---|
| Regulator of microtubule dynamics protein 3 | RMDN3 | 1.5596 |
| Heat shock protein beta-1 | HSPB1 | 1.5607 |
| Vacuolar protein sorting-associated protein 29 | VPS29 | 1.5659 |
| Calcineurin-like phosphoesterase domain-containing protein 1 | CPPED1 | 1.5663 |
| Niban-like protein 1 | FAM129B | 1.5669 |
| ERO1-like protein alpha | ERO1L | 1.5714 |
| Putative adenosylhomocysteinase 2;Adenosylhomocysteinase;Putative adenosylhomocysteinase 3 | AHCYL1;AHCYL2 | 1.5758 |
| Vacuolar protein sorting-associated protein 52 homolog | VPS52 | 1.5789 |
| Probable threonine--tRNA ligase 2, cytoplasmic | TARSL2 | 1.5815 |
| Dihydropyrimidinase-related protein 3 | DPYSL3 | 1.5818 |
| A-kinase anchor protein 8 | AKAP8 | 1.5828 |
| PDZ domain-containing protein GIPC1 | GIPC1 | 1.5833 |
| CD44 antigen | CD44 | 1.586 |
| V-type proton ATPase subunit S1 | ATP6AP1 | 1.5872 |
| Transforming growth factor-beta-induced protein ig-h3 | TGFBI | 1.5888 |
| Tropomyosin 1 (Alpha) | TPM1 | 1.6052 |
| Protein AHNAK2 | AHNAK2 | 1.6092 |
| TOM1-like protein 1 | TOM1L1 | 1.6101 |
| KIF1-binding protein | KIAA1279 | 1.6126 |
| Protein NipSnap homolog 3A | NIPSNAP3A | 1.6134 |
| Triosephosphate isomerase | TPI1 | 1.6135 |
| Cadherin-13 | CDH13 | 1.624 |
| Eukaryotic translation initiation factor 4B | EIF4B | 1.6255 |
| Exocyst complex component 5 | EXOC5 | 1.6258 |
| BAG family molecular chaperone regulator 1 | BAG1 | 1.6268 |
| Squalene synthase | FDFT1 | 1.6322 |
| Nuclear RNA export factor 1 | NXF1 | 1.6329 |
| GTP:AMP phosphotransferase AK4, mitochondrial | AK4 | 1.6375 |
| Transmembrane emp24 domain-containing protein 5 | TMED5 | 1.6385 |
| Putative ribosomal RNA methyltransferase NOP2 | NOP2 | 1.6417 |
| ELKS/Rab6-interacting/CAST family member 1 | ERC1 | 1.6442 |
| Nephronectin | NPNT | 1.6452 |
| SRA stem-loop-interacting RNA-binding protein, mitochondrial | SLIRP | 1.6461 |
| Cysteine and glycine-rich protein 1 | CSRP1 | 1.6468 |
| Dephospho-CoA kinase domain-containing protein | DCAKD | 1.6486 |
| Nuclear receptor-binding protein 2 | NRBP2 | 1.6486 |
| Annexin;Annexin A4 | ANXA4 | 1.6547 |
| Leucine-rich repeat-containing protein 17 | LRRC17 | 1.6558 |
| CD97 antigen;CD97 antigen subunit alpha;CD97 antigen subunit beta | CD97 | 1.6572 |
| Tax1-binding protein 1 | TAX1BP1 | 1.6622 |
| Nucleoporin Nup43 | NUP43 | 1.6761 |
| Cell division control protein 42 homolog | CDC42 | 1.6793 |
| Collagen alpha-1(I) chain | COL1A1 | 1.6822 |
| Zinc finger RNA-binding protein | ZFR | 1.6845 |
| DCN1-like protein 1 | DCUN1D1 | 1.6921 |
| Glucosamine-6-phosphate isomerase 2 | GNPDA2 | 1.6932 |
| Phosphoglycerate kinase 2 | PGK2 | 1.7015 |
| CD9 antigen | CD9 | 1.7035 |
| Metalloproteinase inhibitor 2 | TIMP2 | 1.7065 |
| SWI/SNF-related matrix-associated actin-dependent regulator of chromatin subfamily E member 1 | SMARCE1 | 1.7178 |
| Ezrin | EZR | 1.7212 |
| E3 ubiquitin-protein ligase RNF123 | RNF123 | 1.7302 |
| Atlastin-2 | ATL2 | 1.7338 |
| Transient receptor potential cation channel subfamily M member 4 | TRPM4 | 1.7362 |
| COP9 signalosome complex subunit 6 | COPS6 | 1.7398 |
| Sarcoplasmic/endoplasmic reticulum calcium ATPase 1 | ATP2A1 | 1.746 |
| Protein NDRG1 | NDRG1 | 1.7462 |
| Enoyl-CoA delta isomerase 1, mitochondrial | ECI1;DCI | 1.748 |
| 5-nucleotidase | NT5E | 1.7496 |
| Insulin;Insulin B chain;Insulin A chain | INS;INS-IGF2 | 1.7511 |
| Acyl-coenzyme A thioesterase 1;Acyl-coenzyme A thioesterase 2, mitochondrial | ACOT1;ACOT2 | 1.7519 |

| Protein.names | Gene.names | Simvastatin (Ratio.H.L.normalized) |
|---|---|---|
| Syntaxin-7 | STX7 | 1.7533 |
| Unconventional myosin-Id | MYO1D | 1.754 |
| Fatty aldehyde dehydrogenase | ALDH3A2 | 1.7557 |
| Solute carrier family 2, facilitated glucose transporter member 5 | SLC2A5 | 1.7607 |
| CD81 antigen | CD81 | 1.7654 |
| Ras-related protein Rab-1A | RAB1A | 1.768 |
| Integrin Subunit Alpha 7 | ITGA7 | 1.7712 |
| Coactosin-like protein | COTL1 | 1.7766 |
| Protein OS-9 | OS9 | 1.7784 |
| Filamin-A-interacting protein 1 | FILIP1 | 1.7787 |
| Sarcosine dehydrogenase, mitochondrial | SARDH | 1.7788 |
| Transmembrane protein 97 | TMEM97 | 1.7811 |
| Integrin beta-5;Integrin beta | ITGB5 | 1.7887 |
| Neutral amino acid transporter A | SLC1A4 | 1.7931 |
| Pre-mRNA-processing factor 40 homolog A | PRPF40A | 1.8088 |
| Ubiquitin-conjugating enzyme E2 G1 | UBE2G1 | 1.8123 |
| Integral membrane protein 2B;BRI2, membrane form;BRI2 intracellular domain;BRI2C, soluble form;Bri23 peptide | ITM2B | 1.8149 |
| Chondroitin sulfate proteoglycan 4 | CSPG4 | 1.8238 |
| Isocitrate dehydrogenase [NADP] cytoplasmic | IDH1 | 1.8243 |
| Coronin;Coronin-6 | CORO6 | 1.8251 |
| E3 ubiquitin-protein ligase TRIM23 | TRIM23 | 1.8345 |
| Alkaline phosphatase, tissue-nonspecific isozyme | ALPL | 1.8399 |
| Fibronectin;Anastellin;Ugl-Y1;Ugl-Y2;Ugl-Y3 | FN1 | 1.8434 |
| Lactoylglutathione lyase | GLO1 | 1.8445 |
| Pleckstrin homology domain-containing family O member 2 | PLEKHO2 | 1.8477 |
| Synaptosomal-associated protein 29;Synaptosomal-associated protein | SNAP29 | 1.8529 |
| Protein-arginine deiminase type-2 | PADI2 | 1.8534 |
| 14-3-3 protein eta | YWHAH | 1.8543 |
| Coiled-coil domain-containing protein 80 | CCDC80 | 1.8725 |
| Heat shock protein beta-7 | HSPB7;DKFZp779D0968 | 1.8805 |
| Endoplasmic reticulum metallopeptidase 1 | ERMP1 | 1.8813 |
| Alpha-N-acetylglucosaminidase;Alpha-N-acetylglucosaminidase 82 kDa form;Alpha-N-acetylglucosaminidase 77 kDa form | NAGLU | 1.8961 |
| Superoxide dismutase [Cu-Zn] | SOD1 | 1.8974 |
| Sequestosome-1 | SQSTM1 | 1.8976 |
| Inositol polyphosphate 5-phosphatase K | INPP5K | 1.8989 |
| Nucleolar transcription factor 1 | UBTF | 1.9249 |
| Histone deacetylase;Histone deacetylase 2;Histone deacetylase 1 | HDAC2;HDAC1 | 1.9373 |
| Ras-related protein Rab-21 | RAB21 | 1.9374 |
| E3 ubiquitin-protein ligase RNF14 | RNF14 | 1.9474 |
| CD166 antigen | ALCAM | 1.9481 |
| Olfactomedin-like protein 2A | OLFML2A | 1.9625 |
| Aldo-keto reductase family 1 member C1 | AKR1C1 | 1.9637 |
| Twinfilin-2 | TWF2 | 1.9912 |
| Amyloid beta A4 protein;N-APP;Soluble APP-alpha;Soluble APP-beta;C99;Beta-amyloid protein 42;Beta-amyloid protein 40;C83;P3(42);P3(40);C80;Gamma-secretase C-terminal fragment 59;Gamma-secretase C-terminal fragment 57;Gamma-secretase C-terminal fragment 50;C31 | APP | 2.023 |
| Calcineurin B homologous protein 1 | CHP1 | 2.0725 |
| Prolyl endopeptidase | PREP | 2.0739 |
| Matrix-remodeling-associated protein 7 | MXRA7 | 2.0748 |
| Aldehyde dehydrogenase, dimeric NADP-preferring | ALDH3A1 | 2.0821 |
| Heme-binding protein 2 | HEBP2 | 2.0862 |
| Heat shock-related 70 kDa protein 2 | HSPA2 | 2.0968 |
| Isopentenyl-diphosphate Delta-isomerase 1 | IDI1 | 2.1346 |
| Valacyclovir hydrolase | BPHL | 2.196 |
| Coiled-coil domain-containing protein 90B, mitochondrial | CCDC90B | 2.1966 |
| Proline-rich basic protein 1 | PROB1 | 2.2036 |
| Transcription elongation factor SPT6 | SUPT6H | 2.2121 |
| C-terminal-binding protein 1 | CTBP1 | 2.2171 |
| Lanosterol 14-alpha demethylase | CYP51A1 | 2.2315 |

| **Protein.names** | **Gene.names** | **Simvastatin** (Ratio.H.L.normalized) |
|---|---|---|
| HLA class I histocompatibility antigen, A-36 alpha chain;HLA class I histocompatibility antigen, A-1 alpha chain;HLA class I histocompatibility antigen, A-11 alpha chain;HLA class I histocompatibility antigen, A-3 alpha chain;HLA class I histocompatibility antigen, A-80 alpha chain | HLA-A | 2.2493 |
| Heme oxygenase 1 | HMOX1 | 2.2992 |
| GMP reductase 2;GMP reductase | GMPR2 | 2.3517 |
| Integral membrane protein 2C;CT-BRI3 | ITM2C | 2.353 |
| HLA class I histocompatibility antigen, A-66 alpha chain;HLA class I histocompatibility antigen, A-34 alpha chain;HLA class I histocompatibility antigen, A-26 alpha chain;HLA class I histocompatibility antigen, A-25 alpha chain;HLA class I histocompatibility antigen, A-43 alpha chain;HLA class I histocompatibility antigen, A-33 alpha chain;HLA class I histocompatibility antigen, A-31 alpha chain | HLA-A | 2.3717 |
| Alpha- and gamma-adaptin-binding protein p34 | AAGAB | 2.4019 |
| Heterogeneous nuclear ribonucleoprotein U-like protein 1 | HNRNPUL1 | 2.5168 |
| 72 kDa type IV collagenase;PEX | MMP2 | 2.588 |
| SPARC | SPARC | 2.6067 |
| Matrix metalloproteinase-14 | MMP14 | 2.6678 |
| Pigment epithelium-derived factor | SERPINF1 | 2.7256 |
| Alpha-adducin | ADD1 | 2.8051 |
| Fragile X mental retardation syndrome-related protein 2 | FXR2 | 2.8239 |
| RRP12-like protein | RRP12 | 2.8581 |
| Prosalusin;Salusin-alpha;Salusin-beta;Torsin-2A | TOR2A | 2.8747 |
| Transcription factor 25 | TCF25 | 2.9221 |
| Protein farnesyltransferase subunit beta | CHURC1-FNTB;FNTB | 2.9336 |
| Transforming protein RhoA;Rho-related GTP-binding protein RhoC;Rho-related GTP-binding protein RhoB | RHOA;RHOC;RHOB | 2.9516 |
| Argininosuccinate lyase | ASL | 2.9943 |
| Coiled-coil domain-containing protein 3 | CCDC3 | 3.057 |
| N-alpha-acetyltransferase 20 | NAA20 | 3.1405 |
| Parafibromin | CDC73 | 3.1673 |
| 5-nucleotidase domain-containing protein 3 | NT5DC3 | 3.2032 |
| Prostaglandin-H2 D-isomerase | PTGDS | 3.295 |
| Heat shock 70 kDa protein 14 | HSPA14 | 3.4512 |
| Desmoglein-1 | DSG1 | 3.5184 |
| Ubiquitin domain-containing protein UBFD1 | UBFD1 | 3.5361 |
| WD repeat-containing protein 26 | WDR26 | 3.6259 |
| 5-AMP-activated protein kinase catalytic subunit alpha-2 | PRKAA2 | 3.6778 |
| Receptor tyrosine-protein kinase erbB-2;Receptor tyrosine-protein kinase erbB-4;ERBB4 intracellular domain | ERBB2;ERBB4 | 3.7316 |
| Serine/threonine-protein phosphatase 2A 56 kDa regulatory subunit gamma isoform | PPP2R5C | 3.747 |
| U5 small nuclear ribonucleoprotein 40 kDa protein | SNRNP40;DKFZp434D199 | 3.8353 |
| Zinc finger protein ZPR1 | ZNF259 | 4.1119 |
| Symplekin | SYMPK | 4.2357 |
| Neural cell adhesion molecule L1 | L1CAM | 4.3601 |
| Carnitine O-palmitoyltransferase 1, liver isoform | CPT1A | 4.787 |
| 1-phosphatidylinositol 4,5-bisphosphate phosphodiesterase gamma-1 | PLCG1 | 4.8345 |
| Phosphoglycerate mutase 2 | PGAM2 | 4.8948 |
| Periodic tryptophan protein 2 homolog | PWP2 | 5.7269 |
| Alpha-1,3/1,6-mannosyltransferase ALG2 | ALG2 | 6.8716 |
| Sodium bicarbonate cotransporter 3 | SLC4A7 | 7.7591 |
| Arf-GAP domain and FG repeat-containing protein 1 | AGFG1 | 12.427 |
| V-type proton ATPase 116 kDa subunit a isoform 2 | ATP6V0A2 | 22.944 |

Supplementary Table S6. Differentially expressed proteins from rosuvastatin-treated primary human myotubes.

| Protein.names | Gene.names | Rosuvastatin (Ratio.M.L.normalized) |
|---|---|---|
| Transient receptor potential cation channel subfamily V member 2 | TRPV2 | 0.11993 |
| Dipeptidyl peptidase 1;Dipeptidyl peptidase 1 exclusion domain chain;Dipeptidyl peptidase 1 heavy chain;Dipeptidyl peptidase 1 light chain | CTSC | 0.14899 |
| Glycerate kinase | GLYCTK | 0.15514 |
| E3 ubiquitin-protein ligase listerin | LTN1 | 0.19845 |
| Peptidyl-prolyl cis-trans isomerase-like 3;Peptidyl-prolyl cis-trans isomerase | PPIL3 | 0.22733 |
| Host cell factor 1;HCF N-terminal chain 1;HCF N-terminal chain 2;HCF N-terminal chain 3;HCF N-terminal chain 4;HCF N-terminal chain 5;HCF N-terminal chain 6;HCF C-terminal chain 1;HCF C-terminal chain 2;HCF C-terminal chain 3;HCF C-terminal chain 4;HCF C-terminal chain 5;HCF C-terminal chain 6 | HCFC1 | 0.23896 |
| Carboxymethylenebutenolidase homolog | CMBL | 0.25821 |
| Conserved oligomeric Golgi complex subunit 7 | COG7 | 0.29189 |
| Short-chain specific acyl-CoA dehydrogenase, mitochondrial | ACADS | 0.29531 |
| Protein lin-7 homolog C;Protein lin-7 homolog A | LIN7C;LIN7A | 0.30127 |
| Calsyntenin-2 | CLSTN2 | 0.30642 |
| Group XV phospholipase A2 | PLA2G15 | 0.31443 |
| cGMP-dependent 3,5-cyclic phosphodiesterase | PDE2A | 0.32153 |
| GDP-Man:Man(3)GlcNAc(2)-PP-Dol alpha-1,2-mannosyltransferase | ALG11 | 0.32949 |
| Mediator of DNA damage checkpoint protein 1 | MDC1 | 0.33283 |
| Charged multivesicular body protein 7 | CHMP7 | 0.33723 |
| Bisphosphoglycerate mutase | BPGM | 0.35981 |
| Myosin-6 | MYH6 | 0.36672 |
| EF-hand domain-containing protein D1 | EFHD1 | 0.38834 |
| Immunoglobulin-binding protein 1 | IGBP1 | 0.39996 |
| Nucleotide exchange factor SIL1 | SIL1 | 0.41658 |
| FAS-associated factor 1 | FAF1 | 0.4214 |
| GRB2-associated-binding protein 2 | GAB2 | 0.43554 |
| Protein diaphanous homolog 1 | DIAPH1 | 0.44116 |
| Fatty acyl-CoA reductase 1 | FAR1 | 0.44349 |
| Cell division cycle and apoptosis regulator protein 1 | CCAR1 | 0.46081 |
| Programmed cell death protein 2-like | PDCD2L | 0.46853 |
| Angio-associated migratory cell protein | AAMP | 0.47056 |
| Epidermal growth factor receptor kinase substrate 8-like protein 2 | EPS8L2 | 0.47779 |
| Ragulator complex protein LAMTOR3 | LAMTOR3 | 0.48562 |
| WD repeat-containing protein C2orf44 | C2orf44 | 0.48623 |
| Chromodomain-helicase-DNA-binding protein 4 | CHD4 | 0.48929 |
| Neurofibromin;Neurofibromin truncated | NF1 | 0.49521 |
| Phospholysine phosphohistidine inorganic pyrophosphate phosphatase | LHPP | 0.51634 |
| Integrator complex subunit 2 | INTS2 | 0.51648 |
| Gamma-aminobutyric acid receptor-associated protein-like 1 | GABARAPL1 | 0.5217 |
| 39S ribosomal protein L46, mitochondrial | MRPL46 | 0.52337 |
| Titin | TTN | 0.52449 |
| Nidogen-1 | NID1 | 0.52563 |
| Troponin I, fast skeletal muscle | TNNI2 | 0.52684 |
| LEM domain-containing protein 2 | LEMD2 | 0.53201 |
| Poly [ADP-ribose] polymerase 4 | PARP4 | 0.53432 |
| Tax1-binding protein 1 | TAX1BP1 | 0.53588 |
| SUN domain-containing protein 1 | SUN1;UNC84A | 0.53796 |
| Putative 60S ribosomal protein L39-like 5;60S ribosomal protein L39 | RPL39P5;RPL39 | 0.538 |
| Zinc finger FYVE domain-containing protein 1 | ZFYVE1 | 0.53904 |
| 5-AMP-activated protein kinase subunit gamma-3 | PRKAG3 | 0.54228 |
| Ras-specific guanine nucleotide-releasing factor 2 | RASGRF2 | 0.54367 |
| Ubiquitin Specific Peptidase 39 | USP39 | 0.54418 |
| LanC-like protein 2 | LANCL2 | 0.55522 |
| Small nuclear ribonucleoprotein E | SNRPE | 0.55534 |
| Serine/arginine-rich splicing factor 11 | SRSF11 | 0.55733 |
| Glomulin | GLMN | 0.55753 |
| Ferritin light chain | FTL | 0.55982 |
| Transcription activator BRG1;Probable global transcription activator SNF2L2 | SMARCA4;SMARCA2 | 0.56538 |
| 28S ribosomal protein S36, mitochondrial | MRPS36 | 0.57128 |
| Putative ATP-dependent RNA helicase DHX30 | DHX30 | 0.57915 |
| Ancient ubiquitous protein 1 | AUP1 | 0.58 |
| Probable ATP-dependent RNA helicase DDX46 | DDX46 | 0.58422 |

| Protein.names | Gene.names | Rosuvastatin (Ratio.M.L.normalized) |
|---|---|---|
| Probable threonine--tRNA ligase 2, cytoplasmic | TARSL2 | 0.58558 |
| Nuclear pore complex protein Nup160 | NUP160 | 0.58979 |
| Cleavage stimulation factor subunit 3 | CSTF3 | 0.59081 |
| Serine/threonine-protein phosphatase 4 regulatory subunit 3A | SMEK1 | 0.591 |
| Ribonucleases P/MRP protein subunit POP1 | POP1 | 0.59186 |
| FH1/FH2 domain-containing protein 1 | FHOD1 | 0.59645 |
| E3 ubiquitin-protein ligase HECTD3 | HECTD3 | 0.59897 |
| Interferon regulatory factor 2-binding protein-like | IRF2BPL | 0.60009 |
| Ras association domain-containing protein 4 | RASSF4 | 0.60011 |
| NudC domain-containing protein 1 | NUDCD1 | 0.60016 |
| Phytanoyl-CoA dioxygenase, peroxisomal | PHYH | 0.6041 |
| DNA-directed RNA polymerase II subunit RPB7 | POLR2G | 0.60416 |
| Glutamate--cysteine ligase regulatory subunit | GCLM | 0.60445 |
| Glutamine--fructose-6-phosphate aminotransferase [isomerizing] 2 | GFPT2 | 0.60707 |
| AFG3-like protein 2 | AFG3L2 | 0.60967 |
| Methylthioribose-1-phosphate isomerase | MRI1 | 0.60972 |
| Ras-related protein Rap-1b;Ras-related protein Rap-1b-like protein | RAP1B | 0.61431 |
| DNA topoisomerase 2;DNA topoisomerase 2-beta | TOP2B | 0.61459 |
| Alanyl-tRNA editing protein Aarsd1 | AARSD1;PTGES3L-AARSD1 | 0.61605 |
| RNA-binding protein 34 | RBM34 | 0.61863 |
| Vacuolar protein sorting-associated protein 37B | VPS37B | 0.62377 |
| Endonuclease G, mitochondrial | ENDOG | 0.62577 |
| ATP-binding cassette sub-family B member 7, mitochondrial | ABCB7 | 0.62585 |
| Leukocyte surface antigen CD47 | CD47 | 0.62587 |
| RNA 3-terminal phosphate cyclase-like protein | RCL1 | 0.62653 |
| 39S ribosomal protein L39, mitochondrial | MRPL39 | 0.62674 |
| THO complex subunit 2 | THOC2 | 0.62676 |
| Cysteine-rich protein 1 | CRIP1 | 0.62745 |
| Oxysterol-binding protein;Oxysterol-binding protein-related protein 11;Oxysterol-binding protein-related protein 10 | OSBPL10;OSBPL11 | 0.62881 |
| Disco-interacting protein 2 homolog C | DIP2C | 0.62911 |
| Conserved oligomeric Golgi complex subunit 5 | COG5 | 0.62921 |
| Replication protein A 70 kDa DNA-binding subunit | RPA1 | 0.63014 |
| Spartin | SPG20 | 0.63703 |
| Nuclear pore complex protein Nup85 | NUP85 | 0.63947 |
| Mitochondrial import inner membrane translocase subunit Tim23;Putative mitochondrial import inner membrane translocase subunit Tim23B | TIMM23;TIMM23B | 0.64257 |
| ATP-dependent zinc metalloprotease YME1L1 | YME1L1 | 0.64611 |
| KN motif and ankyrin repeat domain-containing protein 1 | KANK1 | 0.64791 |
| Zinc transporter 1 | SLC30A1 | 0.65069 |
| COMM domain-containing protein 5 | COMMD5 | 0.65261 |
| Zinc finger protein 512 | ZNF512 | 0.65361 |
| Ran-binding protein 6 | RANBP6 | 0.65587 |
| NHP2-like protein 1 | NHP2L1 | 0.65615 |
| Uncharacterized protein C2orf47, mitochondrial | C2orf47 | 0.65639 |
| 1-acyl-sn-glycerol-3-phosphate acyltransferase gamma | AGPAT3 | 0.65742 |
| Serine/threonine-protein kinase SIK3 | SIK3;KIAA0999 | 0.6583 |
| Rho GTPase-activating protein 12 | ARHGAP12 | 0.65859 |
| Elongation factor Ts, mitochondrial;Elongation factor Ts | TSFM | 0.66055 |
| Gamma-glutamyltransferase 5;Gamma-glutamyltransferase 5 heavy chain;Gamma-glutamyltransferase 5 light chain | GGT5 | 0.66196 |
| N-acetyltransferase 10 | NAT10 | 0.665 |
| GTP-binding protein 1 | GTPBP1 | 0.66559 |
| Zinc finger and BTB domain-containing protein 20 | ZBTB20 | 0.66636 |
| A-kinase anchor protein 13 | AKAP13 | 0.6678 |
| Myomesin-2 | MYOM2 | 0.66898 |
| 39S ribosomal protein L4, mitochondrial | MRPL4 | 0.67031 |
| Uroporphyrinogen decarboxylase | UROD | 0.67056 |
| Nucleolar protein 6 | NOL6 | 0.67157 |
| NFU1 iron-sulfur cluster scaffold homolog, mitochondrial | NFU1 | 0.67293 |
| SCY1-like protein 2 | SCYL2 | 0.67401 |
| Acyl-CoA-binding domain-containing protein 5 | ACBD5 | 0.67734 |
| Transmembrane 9 superfamily member 4 | TM9SF4 | 0.67841 |

| Protein.names | Gene.names | **Rosuvastatin** (Ratio.M.L.normalized) |
|---|---|---|
| Heat shock protein beta-11 | HSPB11 | 0.6787 |
| Transportin-3 | TNPO3 | 0.68233 |
| Nischarin | NISCH | 0.68377 |
| DNA repair protein XRCC4 | XRCC4 | 0.68663 |
| Cell division cycle 5-like protein | CDC5L | 0.68818 |
| ATPase family AAA domain-containing protein 1 | ATAD1 | 0.68887 |
| NudC domain-containing protein 3 | NUDCD3 | 0.69 |
| Pre-mRNA-splicing factor SPF27 | BCAS2 | 0.69199 |
| Syntaxin-8 | STX8 | 0.69266 |
| 39S ribosomal protein L14, mitochondrial | MRPL14 | 0.69364 |
| RNA-binding protein 42 | RBM42 | 0.69377 |
| Acidic leucine-rich nuclear phosphoprotein 32 family member B | ANP32B | 0.69701 |
| Trimethyllysine dioxygenase, mitochondrial | TMLHE | 0.69765 |
| Alpha-1,3/1,6-mannosyltransferase ALG2 | ALG2 | 0.69783 |
| IST1 homolog | IST1 | 0.69809 |
| Pantothenate kinase 4 | PANK4 | 0.69865 |
| Pannexin-1 | PANX1 | 0.70008 |
| 39S ribosomal protein L50, mitochondrial | MRPL50 | 0.70009 |
| Eukaryotic translation initiation factor 3 subunit K | EIF3K | 0.70205 |
| Alpha-2-macroglobulin | A2M | 0.70222 |
| 28S ribosomal protein S29, mitochondrial | DAP3 | 0.70315 |
| Protein sel-1 homolog 1 | SEL1L | 0.70349 |
| 28S ribosomal protein S7, mitochondrial | MRPS7 | 0.70485 |
| Protein FAM162A | FAM162A | 0.70491 |
| Diphthine synthase | DPH5 | 0.70525 |
| Serine/threonine-protein kinase N2 | PKN2 | 0.70602 |
| Histidine triad nucleotide-binding protein 2, mitochondrial | HINT2 | 0.70718 |
| OCIA domain-containing protein 1 | OCIAD1 | 0.70762 |
| NFX1-type zinc finger-containing protein 1 | ZNFX1 | 0.709 |
| Putative pre-mRNA-splicing factor ATP-dependent RNA helicase DHX32 | DHX32 | 0.71052 |
| Vacuolar protein sorting-associated protein 41 homolog | VPS41 | 0.71065 |
| Golgin subfamily A member 4 | GOLGA4 | 0.7122 |
| Prostaglandin reductase 2 | PTGR2 | 0.71248 |
| WD repeat-containing protein 75 | WDR75 | 0.71259 |
| WD repeat-containing protein 5 | WDR5 | 0.71284 |
| CD109 antigen | CD109 | 0.71378 |
| Ras-related protein Ral-B | RALB | 0.71829 |
| Huntingtin | HTT | 0.71871 |
| Vacuolar protein sorting-associated protein 16 homolog | VPS16 | 0.71904 |
| Splicing factor 3B subunit 1 | SF3B1 | 0.7191 |
| Solute carrier family 12 member 9 | SLC12A9 | 0.72082 |
| 39S ribosomal protein L33, mitochondrial | MRPL33 | 0.72105 |
| Vacuolar protein sorting-associated protein 13A | VPS13A | 0.72183 |
| Programmed cell death protein 10 | PDCD10 | 0.72284 |
| NEDD8 | NEDD8;NEDD8-MDP1 | 0.72323 |
| Serine/threonine-protein phosphatase 2A 56 kDa regulatory subunit epsilon isoform | PPP2R5E | 0.72389 |
| CDKN2A-interacting protein | CDKN2AIP | 0.72551 |
| Protein transport protein Sec24B | SEC24B | 0.72603 |
| Lamina-associated polypeptide 2, isoforms beta/gamma;Thymopoietin;Thymopentin | TMPO | 0.72608 |
| Nuclear pore glycoprotein p62 | NUP62 | 0.72676 |
| Tensin-3 | TNS3 | 0.72684 |
| Negative elongation factor B | NELFB | 0.72689 |
| Retinol dehydrogenase 13 | RDH13 | 0.72694 |
| Double-stranded RNA-binding protein Staufen homolog 2 | STAU2 | 0.72853 |
| Uncharacterized protein C1orf50 | C1orf50 | 0.72927 |
| Collagen alpha-1(V) chain | COL5A1 | 0.72929 |
| Isochorismatase domain-containing protein 1 | ISOC1 | 0.72975 |
| NEDD8-activating enzyme E1 catalytic subunit | UBA3 | 0.73048 |
| Protein phosphatase 1G | PPM1G | 0.73049 |
| TFIIH basal transcription factor complex helicase XPB subunit | ERCC3 | 0.73074 |
| Pescadillo homolog | PES1 | 0.73236 |

| Protein.names | Gene.names | Rosuvastatin (Ratio.M.L.normalized) |
|---|---|---|
| Focadhesin | FOCAD | 0.73289 |
| Coagulation factor XIII A chain | F13A1 | 0.73339 |
| UPF0364 protein C6orf211 | C6orf211 | 0.73572 |
| Dystrobrevin alpha | DTNA | 0.73652 |
| BH3-interacting domain death agonist;BH3-interacting domain death agonist p15;BH3-interacting domain death agonist p13;BH3-interacting domain death agonist p11 | BID | 0.73733 |
| COMM domain-containing protein 7 | COMMD7 | 0.73734 |
| Ubiquitin-conjugating enzyme E2 O | UBE2O | 0.738 |
| Methylmalonyl-CoA mutase, mitochondrial | MUT | 0.73825 |
| Tigger transposable element-derived protein 1 | TIGD1 | 0.7384 |
| Peptidyl-prolyl cis-trans isomerase C | PPIC | 0.73913 |
| Translation initiation factor eIF-2B subunit delta | EIF2B4 | 0.74 |
| L-aminoadipate-semialdehyde dehydrogenase-phosphopantetheinyl transferase | AASDHPPT | 0.741 |
| Rho GTPase-activating protein 17 | ARHGAP17 | 0.74103 |
| Propionyl-CoA carboxylase beta chain, mitochondrial | PCCB | 0.74135 |
| UPF0687 protein C20orf27 | C20orf27 | 0.74144 |
| Cell division cycle protein 23 homolog | CDC23 | 0.74257 |
| Acylglycerol kinase, mitochondrial | AGK | 0.74297 |
| Solute carrier family 12 member 4;Solute carrier family 12 member 6 | SLC12A4;SLC12A6 | 0.74326 |
| Translation initiation factor eIF-2B subunit beta | EIF2B2 | 0.74381 |
| Synapse-associated protein 1 | SYAP1 | 0.74484 |
| RAC-alpha serine/threonine-protein kinase | AKT1 | 0.74495 |
| Cytochrome c oxidase subunit 7A1, mitochondrial | COX7A1 | 0.74522 |
| Golgi to ER traffic protein 4 homolog | GET4 | 0.74568 |
| Reticulocalbin-3 | RCN3 | 0.74616 |
| Glycerol-3-phosphate dehydrogenase, mitochondrial | GPD2 | 0.74775 |
| Trafficking protein particle complex subunit 4 | TRAPPC4 | 0.74779 |
| Polymerase delta-interacting protein 2 | POLDIP2 | 0.74809 |
| Thioredoxin domain-containing protein 12 | TXNDC12 | 0.74907 |
| Signal peptidase complex catalytic subunit SEC11A | SEC11L1;SEC11A | 1.2512 |
| Inactive dual specificity phosphatase 27 | DUSP27 | 1.2514 |
| Zinc finger protein 292 | ZNF292 | 1.2518 |
| Leupaxin | LPXN | 1.2519 |
| Ras-related protein Rab-1A | RAB1A | 1.2535 |
| BTB/POZ domain-containing protein KCTD12 | KCTD12 | 1.2536 |
| ER membrane protein complex subunit 4 | EMC4 | 1.2538 |
| Serine beta-lactamase-like protein LACTB, mitochondrial | LACTB | 1.2555 |
| Splicing factor U2AF 35 kDa subunit | U2AF1 | 1.2555 |
| Dehydrogenase/reductase SDR family member 7B | DHRS7B | 1.2563 |
| Retinoid-inducible serine carboxypeptidase | SCPEP1 | 1.2566 |
| Caveolin-3 | CAV3 | 1.2583 |
| Stimulator of interferon genes protein | TMEM173 | 1.2588 |
| Coactosin-like protein | COTL1 | 1.2595 |
| mRNA cap guanine-N7 methyltransferase | RNMT | 1.2597 |
| Neuropathy target esterase | PNPLA6 | 1.2611 |
| Fibronectin;Anastellin;Ugl-Y1;Ugl-Y2;Ugl-Y3 | FN1 | 1.2615 |
| Tumor protein p53-inducible protein 11 | TP53I11 | 1.2617 |
| Protein ATP1B4 | ATP1B4 | 1.2621 |
| Myosin light polypeptide 6 | MYL6 | 1.2629 |
| Catalase | CAT | 1.2639 |
| WW domain-containing transcription regulator protein 1 | WWTR1 | 1.2643 |
| Tyrosine-protein phosphatase non-receptor type 12 | PTPN12 | 1.2644 |
| 3-oxo-5-beta-steroid 4-dehydrogenase | AKR1D1 | 1.2665 |
| Fatty acid-binding protein, heart | FABP3 | 1.2669 |
| Oxysterol-binding protein-related protein 8 | OSBPL8 | 1.2679 |
| Pro-cathepsin H;Cathepsin H mini chain;Cathepsin H;Cathepsin H heavy chain;Cathepsin H light chain | CTSH | 1.2686 |
| Phosphatidylinositol 4-kinase type 2-alpha | PI4K2A | 1.2697 |
| Endophilin-A2;Endophilin-A1 | SH3GL1;SH3GL2 | 1.2698 |
| Ras GTPase-activating protein 1 | RASA1 | 1.2716 |
| Insulin-like growth factor 2 mRNA-binding protein 2 | IGF2BP2 | 1.2718 |

| Protein.names | Gene.names | Rosuvastatin (Ratio.M.L.normalized) |
|---|---|---|
| Amyloid beta A4 protein;N-APP;Soluble APP-alpha;Soluble APP-beta;C99;Beta-amyloid protein 42;Beta-amyloid protein 40;C83;P3(42);P3(40);C80;Gamma-secretase C-terminal fragment 59;Gamma-secretase C-terminal fragment 57;Gamma-secretase C-terminal fragment 50;C31 | APP | 1.2723 |
| COP9 signalosome complex subunit 6 | COPS6 | 1.2729 |
| Putative heat shock protein HSP 90-beta 4 | HSP90AB4P | 1.2743 |
| 1-phosphatidylinositol 4,5-bisphosphate phosphodiesterase gamma-1 | PLCG1 | 1.2751 |
| Acetyl-CoA carboxylase 1;Biotin carboxylase | ACACA | 1.2755 |
| Rapamycin-insensitive companion of mTOR | RICTOR | 1.2757 |
| Tyrosine-protein phosphatase non-receptor type 1;Tyrosine-protein phosphatase non-receptor type | PTPN1 | 1.2759 |
| Beta-2-microglobulin;Beta-2-microglobulin form pI 5.3 | B2M | 1.2775 |
| Caspase-7;Caspase-7 subunit p20;Caspase-7 subunit p11 | CASP7 | 1.2777 |
| Cytoplasmic protein NCK1 | NCK1 | 1.278 |
| Ubiquitin carboxyl-terminal hydrolase isozyme L3 | UCHL3 | 1.2782 |
| Collagen alpha-2(I) chain | COL1A2 | 1.2783 |
| Fatty acid synthase;[Acyl-carrier-protein] S-acetyltransferase;[Acyl-carrier-protein] S-malonyltransferase;3-oxoacyl-[acyl-carrier-protein] synthase;3-oxoacyl-[acyl-carrier-protein] reductase;3-hydroxyacyl-[acyl-carrier-protein] dehydratase;Enoyl-[acyl-carrier-protein] reductase;Oleoyl-[acyl-carrier-protein] hydrolase | FASN | 1.2788 |
| Phosphatidylinositide phosphatase SAC1 | SACM1L | 1.2789 |
| Thioredoxin | PDIA3 | 1.28 |
| Heat shock 70 kDa protein 6;Putative heat shock 70 kDa protein 7 | HSPA6;HSPA7 | 1.2819 |
| Histone-lysine N-methyltransferase SETD7 | SETD7 | 1.2831 |
| Ferrochelatase, mitochondrial | FECH | 1.2838 |
| Discoidin domain-containing receptor 2 | DDR2 | 1.2843 |
| Neudesin | NENF | 1.2848 |
| Calcium-binding mitochondrial carrier protein Aralar1 | SLC25A12 | 1.2854 |
| Extracellular matrix protein 1 | ECM1 | 1.2897 |
| Cytochrome b5 | CYB5A | 1.2903 |
| Aspartyl/asparaginyl beta-hydroxylase | ASPH | 1.2906 |
| Nexilin F-Actin Binding Protein | NEXN | 1.2924 |
| 7-dehydrocholesterol reductase | DHCR7 | 1.2925 |
| Enoyl-CoA delta isomerase 1, mitochondrial | ECI1;DCI | 1.2975 |
| KRR1 small subunit processome component homolog | KRR1 | 1.2982 |
| Eukaryotic translation initiation factor 6 | EIF6 | 1.2987 |
| Endoplasmic reticulum metallopeptidase 1 | ERMP1 | 1.2994 |
| E3 ubiquitin-protein ligase TRIM23 | TRIM23 | 1.3 |
| Myosin-7B | MYH7B | 1.3016 |
| Mitogen-activated protein kinase 3 | MAPK3 | 1.302 |
| 1-acyl-sn-glycerol-3-phosphate acyltransferase epsilon | AGPAT5 | 1.3057 |
| NADH dehydrogenase [ubiquinone] 1 alpha subcomplex subunit 11 | NDUFA11 | 1.3065 |
| Heme oxygenase 1 | HMOX1 | 1.3068 |
| Phosphoglycerate kinase 2 | PGK2 | 1.309 |
| Disks large-associated protein 4 | DLGAP4 | 1.3093 |
| Proteasome activator complex subunit 4 | PSME4 | 1.3094 |
| Protein SEC13 homolog | SEC13 | 1.3111 |
| Serine incorporator 1 | SERINC1 | 1.313 |
| Transmembrane protein 97 | TMEM97 | 1.3156 |
| Basigin | BSG | 1.3164 |
| Sortilin | SORT1 | 1.3165 |
| BET1 homolog | BET1;DKFZp781C0425 | 1.3167 |
| DCN1-like protein 1 | DCUN1D1 | 1.3171 |
| Delta(24)-sterol reductase | DHCR24 | 1.318 |
| Proteasome subunit beta type-4 | PSMB4 | 1.3192 |
| Transmembrane protein 87A | TMEM87A | 1.3204 |
| Cystatin-C | CST3 | 1.3219 |
| WD repeat and FYVE domain-containing protein 3 | WDFY3 | 1.325 |
| Peroxisomal acyl-coenzyme A oxidase 3 | ACOX3 | 1.3256 |
| DDB1- and CUL4-associated factor 7 | DCAF7 | 1.3257 |
| WD repeat-containing protein 26 | WDR26 | 1.3259 |
| DnaJ homolog subfamily A member 3, mitochondrial | DNAJA3 | 1.326 |

| Protein.names | Gene.names | Rosuvastatin (Ratio.M.L.normalized) |
|---|---|---|
| Electron transfer flavoprotein-ubiquinone oxidoreductase, mitochondrial | ETFDH | 1.3283 |
| Annexin;Annexin A4 | ANXA4 | 1.3287 |
| NADH dehydrogenase [ubiquinone] 1 alpha subcomplex subunit 4 | NDUFA4 | 1.3296 |
| Cytochrome b5 type B | CYB5B | 1.3328 |
| Calcineurin-like phosphoesterase domain-containing protein 1 | CPPED1 | 1.3343 |
| Leucine-rich repeat-containing protein 40 | LRRC40 | 1.335 |
| Transmembrane emp24 domain-containing protein 5 | TMED5 | 1.3351 |
| Retinol dehydrogenase 11 | RDH11 | 1.3358 |
| GDP-fucose protein O-fucosyltransferase 1 | POFUT1 | 1.3371 |
| Cysteine protease ATG4B | ATG4B | 1.3407 |
| Aminoacyl tRNA synthase complex-interacting multifunctional protein 2 | AIMP2 | 1.3445 |
| Carnitine O-acetyltransferase | CRAT | 1.3448 |
| Alpha-N-acetylglucosaminidase;Alpha-N-acetylglucosaminidase 82 kDa form;Alpha-N-acetylglucosaminidase 77 kDa form | NAGLU | 1.3464 |
| Allograft inflammatory factor 1-like | AIF1L | 1.3482 |
| ER membrane protein complex subunit 7 | EMC7 | 1.3484 |
| B-cell receptor-associated protein 29 | BCAP29 | 1.3494 |
| ATP-citrate synthase | ACLY | 1.3517 |
| N-acetylglucosamine-6-sulfatase | GNS | 1.3537 |
| Monocarboxylate transporter 4 | SLC16A3 | 1.3538 |
| Bifunctional ATP-dependent dihydroxyacetone kinase/FAD-AMP lyase (cyclizing);ATP-dependent dihydroxyacetone kinase;FAD-AMP lyase (cyclizing) | DAK | 1.3545 |
| Vesicle-associated membrane protein 7 | VAMP7 | 1.357 |
| Tight junction protein ZO-1 | TJP1 | 1.3582 |
| Vacuolar protein sorting-associated protein 52 homolog | VPS52 | 1.3582 |
| Leucine-rich repeat flightless-interacting protein 1 | LRRFIP1 | 1.36 |
| Signal transducer and activator of transcription 3 | STAT3 | 1.3607 |
| Superoxide dismutase;Superoxide dismutase [Mn], mitochondrial | SOD2 | 1.3609 |
| Cysteine dioxygenase type 1 | CDO1 | 1.3642 |
| Serrate RNA effector molecule homolog | SRRT | 1.366 |
| Pyruvate dehydrogenase phosphatase regulatory subunit, mitochondrial | PDPR | 1.3662 |
| Vesicle-trafficking protein SEC22b | SEC22B | 1.3671 |
| ERO1-like protein alpha | ERO1L | 1.3708 |
| V-type proton ATPase subunit G 1 | ATP6V1G1 | 1.3716 |
| Tricarboxylate transport protein, mitochondrial | SLC25A1 | 1.3771 |
| Cathepsin B;Cathepsin B light chain;Cathepsin B heavy chain | CTSB | 1.3798 |
| Sorbitol dehydrogenase | SORD | 1.3798 |
| Actin, alpha skeletal muscle | ACTA1 | 1.3799 |
| Myotonin-protein kinase | DMPK | 1.3812 |
| Ras-related protein Rab-5A | RAB5A | 1.3849 |
| NADH dehydrogenase [ubiquinone] 1 beta subcomplex subunit 5, mitochondrial | NDUFB5 | 1.3854 |
| Secretory carrier-associated membrane protein 1 | SCAMP1 | 1.3856 |
| Zinc finger CCCH domain-containing protein 6 | ZC3H6 | 1.3862 |
| Transmembrane protein 205 | TMEM205 | 1.3879 |
| 2-oxoisovalerate dehydrogenase subunit alpha, mitochondrial | BCKDHA | 1.3932 |
| Hydroxysteroid dehydrogenase-like protein 2 | HSDL2 | 1.3956 |
| Putative ataxin-7-like protein 3B | ATXN7L3B | 1.3972 |
| Translocation protein SEC63 homolog | SEC63 | 1.399 |
| Protein cornichon homolog 4 | CNIH4 | 1.3991 |
| Long-chain-fatty-acid--CoA ligase 1 | ACSL1 | 1.3993 |
| U5 small nuclear ribonucleoprotein 40 kDa protein | SNRNP40;DKFZp434D199 | 1.401 |
| Proteolipid protein 2 | PLP2 | 1.4023 |
| Developmentally-regulated GTP-binding protein 2 | DRG2 | 1.4051 |
| Peptide-N(4)-(N-acetyl-beta-glucosaminyl)asparagine amidase | NGLY1 | 1.4109 |
| Methylcrotonoyl-CoA carboxylase beta chain, mitochondrial | MCCC2 | 1.4111 |
| Mitogen-activated protein kinase 14 | MAPK14 | 1.4157 |
| V-type proton ATPase 116 kDa subunit a isoform 2 | ATP6V0A2 | 1.416 |
| UDP-N-acetylhexosamine pyrophosphorylase;UDP-N-acetylgalactosamine pyrophosphorylase;UDP-N-acetylglucosamine pyrophosphorylase | UAP1 | 1.4175 |
| Acetyl-CoA acetyltransferase, cytosolic | ACAT2 | 1.4176 |
| Protein OS-9 | OS9 | 1.4216 |
| Lanosterol synthase | LSS | 1.4312 |

| Protein.names | Gene.names | Rosuvastatin (Ratio.M.L.normalized) |
|---|---|---|
| Serine/threonine-protein kinase 3;Serine/threonine-protein kinase 3 36kDa subunit;Serine/threonine-protein kinase 3 20kDa subunit | STK3 | 1.4334 |
| Adenosine deaminase | ADA | 1.4353 |
| Microsomal glutathione S-transferase 3 | MGST3 | 1.4353 |
| UBX domain-containing protein 6 | UBXN6 | 1.4358 |
| 72 kDa type IV collagenase;PEX | MMP2 | 1.4372 |
| Tubulin-specific chaperone D | TBCD | 1.4373 |
| Cdc42-interacting protein 4 | TRIP10 | 1.4385 |
| Adipocyte plasma membrane-associated protein | APMAP | 1.4415 |
| Arylsulfatase A;Arylsulfatase A component B;Arylsulfatase A component C | ARSA | 1.4444 |
| Armadillo repeat-containing protein 8 | ARMC8 | 1.4462 |
| 39S ribosomal protein L16, mitochondrial | MRPL16 | 1.4474 |
| STE20-like serine/threonine-protein kinase | SLK | 1.4491 |
| HLA class I histocompatibility antigen, alpha chain G | HLA-G | 1.4529 |
| Choline-phosphate cytidylyltransferase A;Choline-phosphate cytidylyltransferase B | PCYT1A;PCYT1B | 1.4543 |
| Methylcrotonoyl-CoA carboxylase subunit alpha, mitochondrial | MCCC1 | 1.4546 |
| Signal peptidase complex subunit 1 | SPCS1 | 1.4563 |
| Cullin-2 | CUL2 | 1.4603 |
| Ankyrin repeat domain-containing protein 2 | ANKRD2 | 1.4613 |
| Alpha-ketoglutarate-dependent dioxygenase FTO | FTO | 1.4656 |
| Solute carrier family 12 member 7;Solute carrier family 12 member 5 | SLC12A7;SLC12A5 | 1.4662 |
| Sarcoplasmic/endoplasmic reticulum calcium ATPase 1 | ATP2A1 | 1.468 |
| Eukaryotic translation initiation factor 1A, X-chromosomal;Eukaryotic translation initiation factor 1A, Y-chromosomal | EIF1AX;EIF1AY | 1.4683 |
| Sodium bicarbonate cotransporter 3 | SLC4A7 | 1.4713 |
| Receptor expression-enhancing protein 6 | REEP6 | 1.4733 |
| ATP-dependent RNA helicase DDX18 | DDX18 | 1.4739 |
| Fructose-bisphosphate aldolase C;Fructose-bisphosphate aldolase | ALDOC | 1.4772 |
| Proliferating cell nuclear antigen | PCNA | 1.4773 |
| RNA-binding protein 39 | RBM39 | 1.4784 |
| Calcineurin B homologous protein 1 | CHP1 | 1.4864 |
| Methyltransferase-like protein 7B | METTL7B | 1.4901 |
| Creatine kinase S-type, mitochondrial | CKMT2 | 1.4926 |
| Tripeptidyl-peptidase 1 | TPP1 | 1.4957 |
| Thiosulfate sulfurtransferase | TST | 1.4988 |
| Histone deacetylase;Histone deacetylase 2;Histone deacetylase 1 | HDAC2;HDAC1 | 1.4994 |
| Vacuolar protein sorting-associated protein 26B | VPS26B | 1.5037 |
| Mevalonate kinase | MVK | 1.5086 |
| Pleckstrin homology domain-containing family O member 2 | PLEKHO2 | 1.5096 |
| Interferon-induced guanylate-binding protein 2;Guanylate-binding protein 3 | GBP2;GBP3 | 1.5115 |
| Acetyl-coenzyme A synthetase, cytoplasmic | ACSS2 | 1.5118 |
| Lactoylglutathione lyase | GLO1 | 1.5147 |
| Peptidyl-prolyl cis-trans isomerase-like 4 | PPIL4 | 1.5189 |
| Myosin light chain 3 | MYL3 | 1.5209 |
| Golgi apparatus protein 1 | GLG1 | 1.5231 |
| HLA class I histocompatibility antigen, A-36 alpha chain;HLA class I histocompatibility antigen, A-1 alpha chain;HLA class I histocompatibility antigen, A-11 alpha chain;HLA class I histocompatibility antigen, A-3 alpha chain;HLA class I histocompatibility antigen, A-80 alpha chain | HLA-A | 1.5231 |
| Farnesyl pyrophosphate synthase | FDPS | 1.5318 |
| Heat shock protein beta-7 | HSPB7;DKFZp779D0968 | 1.5325 |
| Integrin Subunit Alpha 7 | ITGA7 | 1.5384 |
| Inositol polyphosphate 5-phosphatase K | INPP5K | 1.5406 |
| Phosphatidylethanolamine-binding protein 1;Hippocampal cholinergic neurostimulating peptide | PEBP1 | 1.5417 |
| AT-hook DNA-binding motif-containing protein 1 | AHDC1 | 1.5532 |
| Derlin-1 | DERL1 | 1.5548 |
| Tubulin beta-2B chain | TUBB2B | 1.5695 |
| KIF1-binding protein | KIAA1279 | 1.57 |
| Zinc finger CCCH-type antiviral protein 1 | ZC3HAV1 | 1.5742 |
| Isopentenyl-diphosphate Delta-isomerase 1 | IDI1 | 1.5817 |
| Reticulon 4 | RTN4 | 1.5819 |

| Protein.names | Gene.names | Rosuvastatin (Ratio.M.L.normalized) |
|---|---|---|
| Ribulose-phosphate 3-epimerase | RPE | 1.5837 |
| Propionyl-CoA carboxylase alpha chain, mitochondrial | PCCA | 1.5843 |
| Coiled-coil domain-containing protein 90B, mitochondrial | CCDC90B | 1.5847 |
| Kinase D-interacting substrate of 220 kDa | KIDINS220 | 1.5878 |
| COP9 signalosome complex subunit 7a | COPS7A | 1.594 |
| Tubulin beta-3 chain | TUBB3 | 1.5963 |
| Procollagen galactosyltransferase 1 | COLGALT1 | 1.6022 |
| General transcription factor II-I | GTF2I | 1.607 |
| Fat storage-inducing transmembrane protein 2 | FITM2 | 1.6103 |
| Geranylgeranyl transferase type-2 subunit alpha | RABGGTA | 1.6104 |
| Prostaglandin E synthase 3 | PTGES3 | 1.6116 |
| Cyclin-Y | CCNY | 1.6165 |
| Collagen type IV alpha-3-binding protein | COL4A3BP | 1.6184 |
| StAR-related lipid transfer protein 13 | STARD13 | 1.6187 |
| Vacuolar-sorting protein SNF8 | SNF8 | 1.6288 |
| Transient receptor potential cation channel subfamily M member 4 | TRPM4 | 1.6311 |
| Serine/threonine-protein kinase Nek9 | NEK9 | 1.6394 |
| RNA-binding motif, single-stranded-interacting protein 1;RNA-binding motif, single-stranded-interacting protein 3 | RBMS1;RBMS3 | 1.6504 |
| Prolyl endopeptidase | PREP | 1.6688 |
| Cytoplasmic aconitate hydratase | IRP1;ACO1 | 1.6746 |
| Prosalusin;Salusin-alpha;Salusin-beta;Torsin-2A | TOR2A | 1.6805 |
| Dihydropyrimidinase-related protein 3 | DPYSL3 | 1.6822 |
| Xin actin-binding repeat-containing protein 2 | XIRP2 | 1.6831 |
| Alpha- and gamma-adaptin-binding protein p34 | AAGAB | 1.6971 |
| Calmodulin | CALM1;CALM2;CALM3 | 1.7125 |
| Alpha/beta hydrolase domain-containing protein 11 | ABHD11 | 1.7165 |
| Diphosphomevalonate decarboxylase | MVD | 1.7217 |
| TBC1 domain family member 15 | TBC1D15 | 1.7335 |
| Sterol-4-alpha-carboxylate 3-dehydrogenase, decarboxylating | NSDHL | 1.7437 |
| Solute carrier family 35 member E2;Solute carrier family 35 member E2B | SLC35E2;SLC35E2B | 1.7487 |
| Glucosamine-6-phosphate isomerase 2 | GNPDA2 | 1.7579 |
| Nephronectin | NPNT | 1.7609 |
| Tropomyosin 1 (Alpha) | TPM1 | 1.7673 |
| Myotubularin-related protein 6 | MTMR6 | 1.7802 |
| HLA class I histocompatibility antigen, A-66 alpha chain;HLA class I histocompatibility antigen, A-34 alpha chain;HLA class I histocompatibility antigen, A-26 alpha chain;HLA class I histocompatibility antigen, A-25 alpha chain;HLA class I histocompatibility antigen, A-43 alpha chain;HLA class I histocompatibility antigen, A-33 alpha chain;HLA class I histocompatibility antigen, A-31 alpha chain | HLA-A | 1.7808 |
| Hydroxymethylglutaryl-CoA synthase, cytoplasmic | HMGCS1 | 1.7971 |
| 4-aminobutyrate aminotransferase, mitochondrial | ABAT | 1.7987 |
| Cleft lip and palate transmembrane protein 1-like protein | CLPTM1L | 1.8039 |
| Anoctamin-10;Anoctamin | ANO10 | 1.8044 |
| Parathymosin | PTMS | 1.8106 |
| Geranylgeranyl transferase type-1 subunit beta | PGGT1B | 1.8122 |
| Parafibromin | CDC73 | 1.8157 |
| C-terminal-binding protein 1 | CTBP1 | 1.8186 |
| Fatty acid desaturase 2 | FADS2 | 1.8317 |
| A-kinase anchor protein 2 | AKAP2 | 1.8361 |
| Guanine nucleotide-binding protein-like 1 | GNL1 | 1.8412 |
| Ras-related protein Rab-22A | RAB22A | 1.8429 |
| Matrix metalloproteinase-14 | MMP14 | 1.8524 |
| 5-3 exoribonuclease 1 | XRN1 | 1.865 |
| Paralemmin-2 | PALM2;PALM2-AKAP2 | 1.8715 |
| Neural cell adhesion molecule L1 | L1CAM | 1.886 |
| Matrix-remodeling-associated protein 7 | MXRA7 | 1.8913 |
| Arf-GAP with SH3 domain, ANK repeat and PH domain-containing protein 2 | ASAP2 | 1.8984 |
| Cytochrome b-c1 complex subunit 8 | UQCRQ | 1.9459 |
| WD40 repeat-containing protein SMU1 | SMU1 | 1.9524 |
| Thiopurine S-methyltransferase | TPMT | 1.9594 |
| Cytochrome c oxidase subunit 7A2, mitochondrial | COX7A2 | 1.9639 |

| Protein.names | Gene.names | **Rosuvastatin** (Ratio.M.L.normalized) |
|---|---|---|
| Proline-rich basic protein 1 | PROB1 | 1.9701 |
| E3 ubiquitin-protein ligase RNF123 | RNF123 | 1.9959 |
| Voltage-dependent calcium channel gamma-1 subunit | CACNG1 | 2.0283 |
| SRA stem-loop-interacting RNA-binding protein, mitochondrial | SLIRP | 2.0421 |
| N-alpha-acetyltransferase 20 | NAA20 | 2.0442 |
| GMP reductase 2;GMP reductase | GMPR2 | 2.0684 |
| NIF3-like protein 1 | NIF3L1 | 2.0725 |
| Uracil phosphoribosyltransferase homolog | UPRT | 2.0826 |
| Receptor tyrosine-protein kinase erbB-2;Receptor tyrosine-protein kinase erbB-4;ERBB4 intracellular domain | ERBB2;ERBB4 | 2.0937 |
| PERQ amino acid-rich with GYF domain-containing protein 2 | GIGYF2 | 2.1312 |
| FUN14 domain-containing protein 2 | FUNDC2 | 2.1415 |
| Ankyrin repeat and SAM domain-containing protein 1A | ANKS1A | 2.1653 |
| ER lumen protein retaining receptor 1;ER lumen protein retaining receptor;ER lumen protein retaining receptor 2 | KDELR1;KDELR2 | 2.1745 |
| Glutaryl-CoA dehydrogenase, mitochondrial | GCDH | 2.2083 |
| Ras-related protein Rab-24 | RAB24 | 2.22 |
| Serine/threonine-protein phosphatase 2A 56 kDa regulatory subunit gamma isoform | PPP2R5C | 2.2712 |
| V-type proton ATPase subunit S1 | ATP6AP1 | 2.2956 |
| Squalene monooxygenase | SQLE | 2.2973 |
| Xaa-Pro dipeptidase | PEPD | 2.3284 |
| Methyltransferase-like protein 7A | METTL7A | 2.335 |
| Squalene synthase | FDFT1 | 2.3409 |
| CD97 antigen;CD97 antigen subunit alpha;CD97 antigen subunit beta | CD97 | 2.3436 |
| Nuclear RNA export factor 1 | NXF1 | 2.3686 |
| Actin-related protein 3B | ACTR3B | 2.4102 |
| Striatin | STRN | 2.4552 |
| Cleavage stimulation factor subunit 1 | CSTF1 | 2.4724 |
| Receptor-type tyrosine-protein phosphatase O | PTPRO | 2.5766 |
| Pre-mRNA-processing factor 40 homolog A | PRPF40A | 2.6525 |
| Pterin-4-alpha-carbinolamine dehydratase | PCBD1 | 2.7346 |
| Polypeptide N-acetylgalactosaminyltransferase 1;Polypeptide N-acetylgalactosaminyltransferase 1 soluble form | GALNT1 | 2.7515 |
| Alpha-adducin | ADD1 | 2.7586 |
| Protein farnesyltransferase subunit beta | CHURC1-FNTB;FNTB | 2.866 |
| Protein FAM65B | FAM65B | 2.8929 |
| Ubiquitin domain-containing protein UBFD1 | UBFD1 | 2.9823 |
| Sorting nexin-17 | SNX17 | 3.0627 |
| Desmoglein-1 | DSG1 | 3.0841 |
| RILP-like protein 1 | RILPL1 | 3.1002 |
| Conserved oligomeric Golgi complex subunit 3 | COG3 | 3.1377 |
| Phosphoglycerate mutase 2 | PGAM2 | 3.1387 |
| Lanosterol 14-alpha demethylase | CYP51A1 | 3.1904 |
| Activating signal cointegrator 1 | TRIP4 | 3.2051 |
| pre-rRNA processing protein FTSJ3 | FTSJ3 | 3.3368 |
| Heat shock 70 kDa protein 14 | HSPA14 | 3.3799 |
| Probable leucine--tRNA ligase, mitochondrial | LARS2 | 3.4207 |
| RNA polymerase II-associated protein 3 | RPAP3 | 3.4247 |
| Protein unc-45 homolog A | UNC45A | 3.6378 |
| Nucleoporin NUP188 homolog | NUP188 | 3.6421 |
| Heterogeneous nuclear ribonucleoprotein U-like protein 1 | HNRNPUL1 | 3.6598 |
| Redox-regulatory protein FAM213A | FAM213A | 3.9479 |
| Glutaredoxin-related protein 5, mitochondrial | GLRX5 | 4.5337 |
| Arf-GAP domain and FG repeat-containing protein 1 | AGFG1 | 4.6811 |
| Protein S100-A8;Protein S100-A8, N-terminally processed | S100A8 | 8.3202 |
| Cysteine-rich with EGF-like domain protein 2 | CRELD2 | 8.6815 |
| Suppressor of SWI4 1 homolog | PPAN | 10.224 |
| Carnitine O-palmitoyltransferase 1, liver isoform | CPT1A | 12.818 |
| Protein S100-A9 | S100A9 | 30.406 |
